# Mitochondrial Dysfunction and Fatigue in Sjögren’s Disease

**DOI:** 10.1101/2024.06.17.598269

**Authors:** Biji T. Kurien, John A. Ice, Rebecca Wood, Gavin Pharaoh, Joshua Cavett, Valerie Lewis, Shylesh Bhaskaran, Astrid Rasmussen, Christopher J. Lessard, Amy Darise Farris, Kathy L. Sivils, Kristi A. Koelsch, Holly Van Remmen, R. Hal Scofield

## Abstract

**Objectives:** Sjögren’s disease (SjD) is a common exocrine disorder typified by chronic inflammation and dryness, but also profound fatigue, suggesting a pathological basis in cellular bioenergetics. In healthy states, damaged or dysfunctional mitochondrial components are broken down and recycled by mitophagy, a specialized form of autophagy. In many autoimmune disorders, however, evidence suggests that dysfunctional mitophagy allows poorly functioning mitochondria to persist and contribute to a cellular milieu with elevated reactive oxygen species. We hypothesized that mitophagic processes are dysregulated in SjD and that dysfunctional mitochondria contribute to overall fatigue. We sought to link fatigue with mitochondrial dysfunction directly in SjD, heretofore unexamined, and further sought to assess the pathogenic extent and implications of dysregulated mitophagy in SjD.

**Methods:** We isolated pan T cells via negative selection from the peripheral blood mononuclear cells of 17 SjD and 8 age-matched healthy subjects, all of whom completed fatigue questionnaires prior to phlebotomy. Isolated T cells were analyzed for mitochondrial oxygen consumption rate (OCR) and glycolysis using Seahorse, and linear correlations with fatigue measures were assessed. A mitophagy transcriptional signature in SjD was identified by reanalysis of whole-blood microarray data from 190 SjD and 32 healthy subjects. Differential expression analyses were performed by case/control and subgroup analyses comparing SjD patients by mitophagy transcriptional cluster against healthy subjects followed by bioinformatic interpretation using gene set enrichment analysis.

**Results:** Basal OCR, ATP-linked respiration, maximal respiration, and reserve capacity were significantly lower in SjD compared to healthy subjects with no observed differences in non-mitochondrial respiration, basal glycolysis, or glycolytic stress. SjD lymphocytic mitochondria show structural alterations compared to healthy subjects. Fatigue scores related to pain/discomfort in SjD correlated with the altered OCR. Results from subgroup analyses by mitophagic SjD clusters revealed highly variable inter-cluster differentially expressed genes (DEGs) and expanded the number of SjD-associated gene targets by tenfold within the same dataset.

**Conclusion:** Mitochondrial dysfunction, associated with fatigue, is a significant problem in SjD and warrants further investigation.

## Introduction

Sjögren’s disease (SjD) is a common autoimmune exocrinopathy typified by both subjective and objective features of dryness, most frequently involving the eyes (keratoconjunctivitis sicca) and mouth (xerostomia). Yet other mucosal surfaces including the airways, digestive tract, and/or vagina may become involved, resulting in “sicca syndrome”.^1^ While extraglandular manifestations like central/peripheral nervous system involvement, arthritis, interstitial lung disease, and cutaneous vasculitis are not infrequent, severe and persistent fatigue is common and debilitating in SjD. Indeed, with up to 90% of SjD patients reporting fatigue, it remains one of the most troubling aspects of the disease.^2–4^

Despite being a common disorder, the etiology and underlying pathogenesis of SjD are incompletely understood. This has resulted in frequent under- or misdiagnosis and undertreatment. Current meta-analyses estimate a pooled incidence of 6.92 per 100,000 people/year and a prevalence of 60.82 cases per 100,000 for SjD.^5^ Affecting predominantly middle-aged women and displaying a pronounced sex bias with an average female-to-male ratio of >9:1,^6^ SjD is commonly diagnosed between the ages of 51.6 and 62 years, although symptoms often start years before diagnosis.^7,8^

As with other autoimmune diseases, the precise causes of SjD remain unknown. Environmental exposures in individuals with genetic susceptibility are thought to play critical roles in immune system dysregulation and SjD occurrence.^9^ Accumulating evidence from genetic and epigenetic studies has revealed the association of numerous risk loci across the genome with allele variants contributing to SjD risk.^10^ These and other data point to the critical role that imbalance of innate immune responses plays in SjD pathogenesis (particularly early in the disease) through sustained activation of interferon (IFN) responses, particularly Type I IFNs (IFN-I).^11^ Although infectious agents have long been thought to act as triggers, viral infections like Epstein-Barr Virus (EBV), hepatitis C virus (HCV), and Coxsackie A have not shown association with SjD^12^ despite evidence showing SjD-specific genetic intervals containing likely binding sites for EBV transcription factors.^13^

Several studies have shown association between chronic inflammatory responses and autoimmune diseases.^14,15^ Increased oxidative stress biomarkers and proinflammatory cytokines like tumor necrosis factor- alpha (TNF-α), interleukin-6 (IL-6), IL-12, IL-18, and IFN-γ have been widely reported in SjD.^16^ While several studies have emphasized the role of chronic inflammatory responses in SjD^17^, the precise molecular mechanisms involved are incompletely known. In addition, classical stimuli like infection and injury alone do not appear to cause chronic inflammatory conditions, which instead arises from tissue malfunction due to disrupted cellular homeostatis.^18,19^

Mitochondria serve as the main source of ATP and reactive oxygen species (ROS) and act as the ultimate regulators of the redox state.^20^ In addition, these organelles buffer Ca^2+^ signaling by uptake through an inner mitochondrial membrane uniporter and take part in lipid and amino acid metabolism.^21^ The respiratory chain within the inner membrane generates an electrochemical gradient to power complex V, ATP synthase, which makes most cellular ATP.^20^ In addition to cellular bioenergetics, mitochondria are involved in apoptotic cell death, cell signaling, autophagy, and immune response regulation.^22,23^ Despite this, mitochondrial dysfunction has not been fully explored in autoimmune disease pathogenesis. Fatigue and other symptoms occur in several chronic diseases where imbalance between biogenesis and mitophagic processes contribute to progressive pathological conditions linked by mitochondrial dysfunction.^24^ Changes in electrical and chemical transmembrane inner mitochondrial membrane potential, decreases in crucial metabolites transported into mitochondria, and functional changes in the electron transport chain that diminish ATP synthesis are all mechanisms by which mitochondrial molecular dysfunction occurs.^25^

In this study, we investigated the relationship between fatigue and mitochondrial dysfunction in SjD, first by assessing mitochondrial dysfunction in pan T cells isolated from the peripheral blood mononuclear cells (PBMCs) of SjD and healthy subjects and then by reanalysis of a large public GEP dataset to identify mitophagy transcriptional signatures in SjD. As PBMCs are composed of 70-90% lymphocytes, which are made up of 70- 85% CD3^+^ T cells^26^, this immune cell population is likely susceptible to pathogenic influences resulting from SjD and likely reflective of broader mitochondrial dysfunction in SjD. We report clear perturbations in mitochondrial function amongst SjD patients and show the direct correlation between fatigue measures and assessed mitochondrial activity.

Furthermore, we undertook the reanalysis of a public gene expression profiling (GEP) dataset to understand the scope of mitochondrial dysfunction and altered mitophagy on the SjD transcriptional landscape. Our study goals were to be able to: 1) stratify SjD patients by transcriptional differences in key mitophagy genes to contextualize mitochondrial dysfunction; 2) identify transcriptional signatures suggestive of SjD endotypes for future clinical correlation; and 3) expand and clarify the identities and possible roles of dysregulated SjD genes for future investigation. Exploring differences in mitochondrial function in the T cells of SjD patients has the potential to provide new insights and perspectives to understand the underlying causes of fatigue in SjD. Indeed, efforts to characterize the global transcriptomic phenotypes and endotypes of SjD patients has the potential to identify novel molecular targets for therapeutics and opportunities for clinical management using precision-medicine approaches.

## Methods

### Patients

Seventeen (17) SjD subjects, all of whom met the American/European Consensus Group (AECG) for primary SjD classification criteria, were recruited from the Oklahoma Medical Research Foundation (OMRF) Sjögren’s Research Clinic^27^ ; eight (8) healthy subjects were recruited from the University of Oklahoma Health Sciences Center (OUHSC) campus. Studies were approved by the Institutional Review Boards of the OUHSC and OMRF. After obtaining written, informed consent from all subjects, patients completed fatigue questionnaires immediately followed by phlebotomy. Blood samples were obtained in Vacuette Blood Collection Tubes with heparin (Thomas Scientific, Swedesboro, NJ).

### T cell isolation

Blood samples were subjected to density gradient centrifugation to obtain peripheral blood mononuclear cells (PBMCs). T cells were isolated by negative selection from PBMCs using the Pan T cell isolation kit from Miltenyi (Cat #130-096-535) according to manufacturer’s instructions.

### Coating with Cell-Tak

Seahorse XF Cell Mito Stress kits were purchased from Agilent Technologies, Inc., Santa Clara, CA. Wells of XF24 Seahorse cell culture plates were coated with 50 µl/well of Cell-Tak (22.4 µg/ml sodium bicarbonate buffer, pH 8) according to the manufacturer’s instructions. Cell-Tak Cell and Tissue Adhesive was purchased from Corning Inc., One Riverfront Plaza Corning, NY. The Cell-Tak was allowed to sit at room temperature for 20 min, and the wells were subsequently rinsed twice with 200 µl of distilled water. The plate was stored at 4°C for use the following day.

### Coating of T cells to Cell-Tak Coated Plate

We added one million T cells in 100 µl of Seahorse running media to each well of the XF24 Seahorse cell culture plate coated with Cell-Tak. The plate was spun at 1000 g for 1 minute followed by incubation at 37°C for 30 minutes. Then, 400 µl of Seahorse running media was gently added along the sides of each well of the plated cells.

### Seahorse XF24 Extracellular Flux Analyzer Assay of T cell mitochondrial respiration and glycolysis

The sensor cartridge was hydrated overnight at 37°C with the manufacturer-supplied calibrant. The following day immediately prior to running the assay, we loaded the antibiotics oligomycin, FCCP (carbonyl cyanide-p-trifluoromethoxy phenylhydrazone), and antimycin A into the wells of the sensor cartridge. The cartridge was calibrated using the Seahorse XF24 Extracellular Flux Analyzer. The plate with the cells was then added to the cartridge in the analyzer. Mitochondrial respiration was assessed by monitoring changes in oxygen consumption rate while glycolysis was determined by measuring changes in extracellular acidification rate using the Seahorse XF24 Extracellular Flux Analyzer as described previously.^46^ Briefly, at the time of the assay run the Extracellular Flux Analyzer recorded three measurements of basal oxygen consumption. Then, the Flux Analyzer injected and mixed the complex V inhibitor oligomycin (1 μM) and recorded two measurements to determine ATP-linked oxygen consumption. Following the oligomycin, the analyzer injected and mixed the proton uncoupler FCCP (1 μM) and recorded another two measurements to determine maximal respiration capacity. Finally, the analyzer injected and mixed the complex III inhibitor antimycin A (1 μM) and recorded two more measurements to determine non-mitochondrial oxygen consumption. The oxygen consumption rate of each well was normalized to total cells coated per well.^28^

### Fatigue Assessment

We administered several previously validated questionnaires to the SjD and control subjects to assess fatigue during their phlebotomy visit. The questionnaires used to assess fatigue in SjD and healthy subjects included Bowman’s SjD,^29^ Fatigue Impact Scale,^30^ Multidimensional assessment of fatigue (provides a global score which encompasses fatigue severity, distress, QoL, and timing of fatigue),^31^ Centers for Epidemiological Studies - Depression Scale, Fatigue Severity Scale,^32^ The Rheumatology Attitudes Index or Helplessness Score, Modified Health Assessment,^33^ Validation of the Functioning in Chronic Illness Scale (FCIS; evaluates disease impact on the patient, the patient’s impact on the disease and the effect of the disease on the attitudes of the patient),^34^ SLEEP 1, and SLEEP 2. None of the subjects studied had coexisting medical conditions that could account for the assessed fatigue symptoms.

### Transmission Electron Microscopy

PBMCs were isolated from SjD and healthy subjects by density-gradient centrifugation and were subjected to transmission electron microscopy. Cells were initially fixed in 1% paraformaldehyde/3% glutaraldehyde in 0.1M sodium cacodylate buffer (pH 7.4) and were post-fixed in 1% osmium tetroxide + 1.5% potassium ferrocyanide in a 0.1M sodium cacodylate buffer. Dehydration was obtained through gradient ethanol series followed by propylene oxide. Epon/Araldite resin infiltration was assisted by three changes in decreasing gradients of propylene oxide/resin prior to incubation in pure resin, and then complete resin (with accelerator). Cell pellets were spun into complete resin in BEEM capsules and polymerized in an oven at 60°C prior to sectioning.

Sectioning was performed on a Leica UC6 ultramicrotome. Approximately 100nm ultrathin sections were placed on 400 mesh copper grids and lead stained for contrast using standard methods. Samples were viewed using a Hitachi H-7600 transmission electron microscope; digital images were acquired using a Kodak AMT 4Kx4K camera.

### Statistics

Averages with standard error are presented unless otherwise stated. Categorical data were analyzed by chi square. Continuous data were assessed by two-tailed Student’s T test. P-values of *p*<0.05 were considered significant. We analyzed fatigue scores using simple linear regression executed in GraphPad Prism version 10.2.3 for Windows, GraphPad Software, Boston, Massachusetts USA, www.graphpad.com.

### Gene Expression Profiling

All analyses related to gene expression profiling were executed in R Statistical Software (v4.3.3)^35^ as implemented in RStudio^36^. Using the package *GEOquery*^37^, we downloaded a public SjD GEP dataset (GSE51092) consisting of *log_2_*-transformed, background-normalized, and batch-corrected data for 15,063 transcript probes assessed across 222 samples (SjD=190; healthy=32). As many gene IDs have changed since the original deposition of the dataset^38^, we undertook gene reannotation using QIAGEN Ingenuity Pathway Analysis (IPA).^39,40^ Briefly, 15,063 transcript probe Illumina IDs (GPL6884, Illumina HumanWG-6 v3.0 expression beadchip) comprising GSE51092 were uploaded to IPA wherein they were converted to official gene symbols using pre-matched entries corresponding to array-specific Illumina transcript probe IDs. Following transcript reannotation, we extracted those mitophagy genes belonging to the KEGG pathway for mitophagy in humans (hsa04137). Out of the 103 genes comprising hsa04137, we identified 85 overlapping transcript probes from 15,063 available to define our mitophagy analysis set.

Differential expression analysis (DEA) was performed using the package *limma*^41^, designing the model matrix to contrast all SjD (S_1_; n=190) vs. healthy subjects (H_0_; n=32). We used the functions *lmfit* and *arrayWeights* to fit our model to this data using weighted-array approaches^42^. Finally, we applied the empirical Bayes’ step to obtain differential expression statistics, including fold change (FC) and Benjamini-Hochberg-adjusted p-values (*p_adj_*). Differentially expressed (DE) transcripts were defined by *p_adj_*<0.05 and |log_2_(FC)|>1 unless otherwise stated, and the resulting gene sets were visualized using heatmaps and volcano plots generated with the packages *pheatmap*^43^ and *ggplot2*^44^, respectively.

### Clustering Approaches & Subgroup DEA

To identify transcriptionally distinct mitophagy clusters in SjD, we applied unsupervised hierarchical clustering using average linkage to a gene expression matrix consisting of the 28 DE probes across 20 distinct mitophagy genes within all SjD patients (S_1_; n=190). Following clustering, SjD patients readily stratified into discrete mitophagy clusters, and we created gene matrices for each cluster group (M_1_-M_5_) consisting exclusively of within-cluster SjD peers and healthy subjects (H_0_). We then repeated DEA by contrasting each SjD mitophagy cluster against all healthy subjects (e.g., M_1_ vs. H_0_, M_2_ vs. H_0_, etc.), treating these as subset analyses of SjD patients with a shared mitophagy phenotype. DE transcripts for each cluster were defined and visualized as above.

### Gene Set Enrichment Analysis (GSEA)

Gene Set Enrichment Analysis (GSEA) was performed using the package *clusterProfiler*^45^ using a FC-ranked list derived from DEA results comparing: 1) all SjD subjects (S_1_) vs. healthy subjects (H_0_); and 2) SjD subjects by mitophagy cluster (M_1_-M_5_) vs. healthy subjects (H_0_). Significantly enriched gene sets were defined by *p_adj_*<0.05, and results are presented for the overall analysis (S_1_) and by mitophagy cluster (M_1_-M_5_). Using the *clusterProfiler* functions *gseGO* and *gseKEGG*, we assessed the Gene Ontology (GO) and Kyoto Encyclopedia of Genes and Genomes (KEGG) pathways using gene set size thresholds of 15/450 (min/max) over 1000 iterations.

## Results

### Metabolic Profiling of T cells in SjD and Healthy Subjects

We found significantly decreased basal oxygen consumption rate (OCR), namely mitochondrial function, in T cells comparing SjD vs. healthy subjects (70.09±5.6 vs. 146.31±14.07 pmol/O_2_/min/million cells, respectively, *p*<3x10^-6^; figure 1A). We found no significant differences in non-mitochondrial respiration (figure 1B). ATP-linked respiration, representing the proportion of basal respiration linked to energy production after adding oligomycin, was calculated as the difference between basal respiration and proton leak in live T cells. We found significantly decreased ATP-linked respiration in SjD T cells compared to healthy subjects (66.37±4.5 vs. 121.1±15.42 pmol/O_2_/min/million cells, respectively, *p*<2x10^-4^; figure 1C). Maximal respiration was also significantly decreased in SjD vs. healthy subjects (176.29±18 vs. 418±64.34 pmol/O_2_/min/million cells, respectively, *p*<3.6x10^-4^; figure 1D). Finally, we added antimycin A to determine the cellular reserve capacity (i.e. ability to increase ATP production in response to stress), which was decreased in T cells from SjD compared to healthy subjects (107.47±13.1 vs. 300.65±68.66 pmol/O_2_/min/million cells, respectively, *p*<4x10^-4^; figure 1E). Interestingly, we found no difference in the extracellular acidification rate (glycolysis) in T cells between SjD and healthy subjects (figure 1F). To rule out the contribution of age to the observed differences, we compared ages between healthy and SjD subjects and found no significant difference (52.25±3.21 vs. 59.88±3.17 years, respectively, *p*=0.17; online supplemental figure 1 and online supplemental table 1). Visualization of mitochondria using transmission electron microscopy shows clearly swollen mitochondria with prominently disorganized cristae in lymphocytes from an SjD subject (figure 1G) compared to those from a healthy subject (figure 1H).

**Figure 1:**
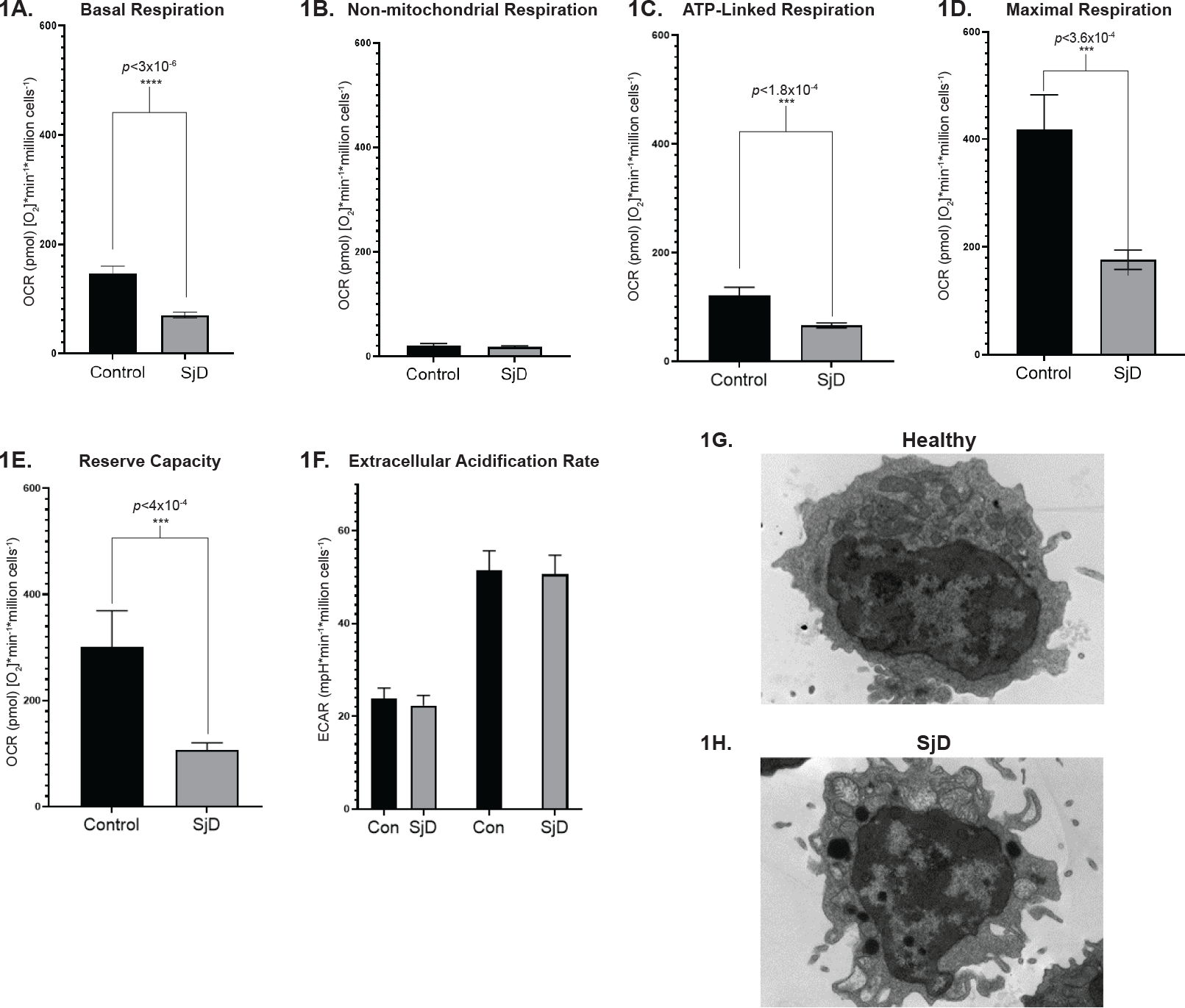
Mitochondrial metabolic characterization in T cells from SjD and healthy subjects fixed to the wells of tissue culture plates and analyzed using the Seahorse XF24 Extracellular Flux Analyzer. 1A: Mitochondrial basal respiration (mitochondrial function under normal physiological conditions); 1B: Non-mitochondrial respiration (the lowest oxygen consumption reading after adding antimycin A); 1C: ATP-linked respiration (the proportion of basal respiration linked to energy production); 1D: Maximal respiration (assesses the maximum phosphorylation capability of electron transport chain, after uncoupling electron transport chain from oxidative phosphorylation with FCCP; 1E: Mitochondrial reserve capacity (the ability of the cell to increase ATP production in response to stress); 1F: Basal (glycolytic function under normal physiological conditions) and stressed (representing highest extracellular acidification measured after adding FCCP); 1G: Representative transmission electron micrographs showing lymphocytes obtained from one each of SjD and healthy subjects. Values are expressed as the mean ± standard error of the mean.

We next identified those fatigue measures (using validated questionnaires; see methods) showing linear correlation with mitochondrial measures in SjD T cells. Scores from the Bowman fatigue questionnaire showed the best correlation with basal OCR (*r^2^*=0.34, *p*=0.014), ATP-linked respiration (*r^2^*=0.36, *p*=0.011), maximal respiration (*r^2^*=0.491, *p*=0.0017), and reserve capacity (*r^2^*=0.486, *p*=0.0019) in SjD (figure 2). These scores from the general fatigue category (pain/discomfort) related to the question “the worst problem with pains, the worst discomfort I’ve experienced in 2 weeks”. Fatigue Severity Scale (FSS) scores correlated only with mitochondrial reserve capacity in SjD (*r^2^*=0.244, *p*=0.044). Scores from Bowman mental fatigue questionnaire showed correlation trending significant correlation with ATP-linked respiration (*r^2^*=0.22, *p*=0.059; figure 2).

**Figure 2:**
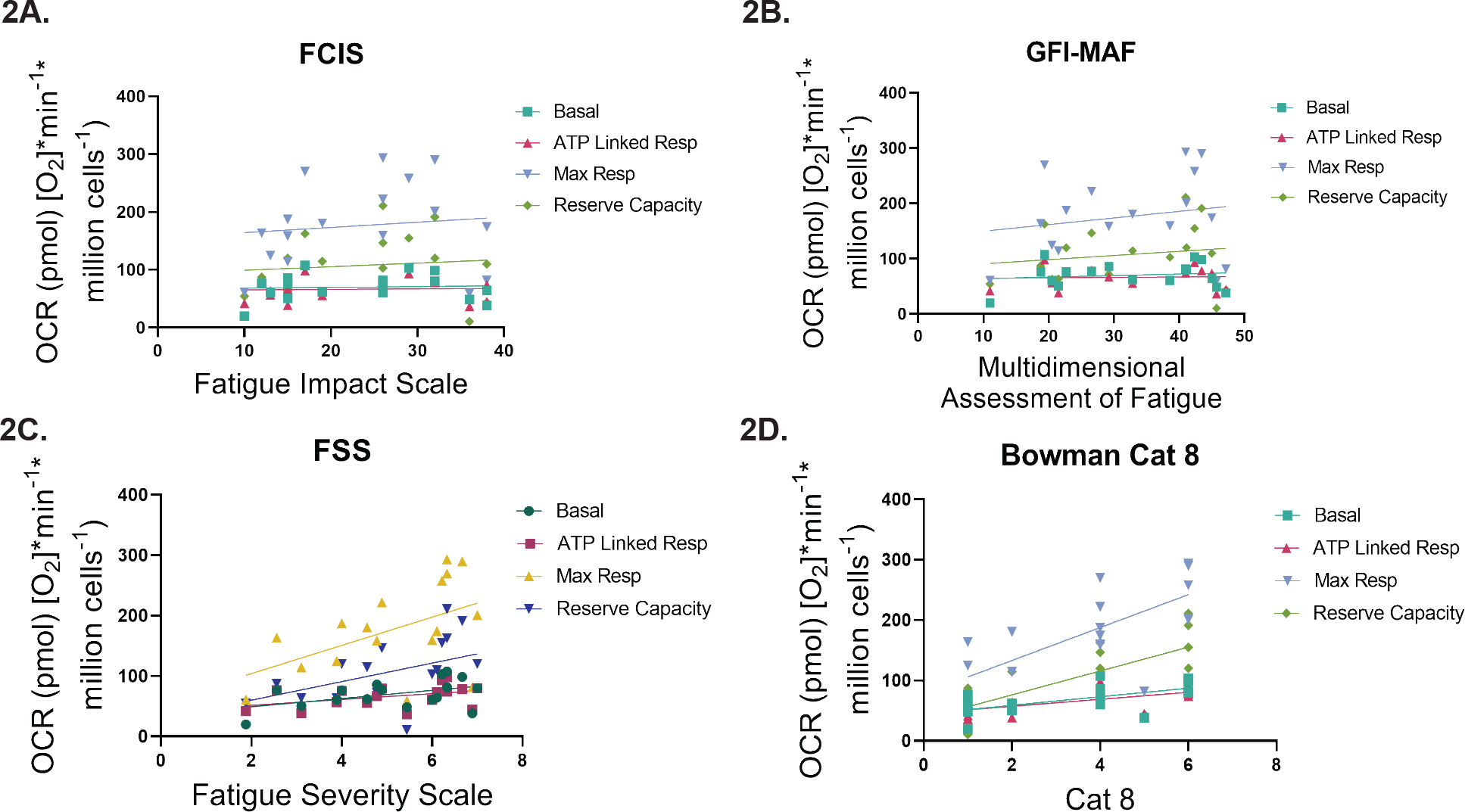
Correlative relationships between fatigue scores assessed using validated questions and mitochondrial metabolic measures (basal oxygen consumption rate, ATP linked respiration, maximal respiration, and reserve capacity). 2A: Correlation between oxygen consumption rate (OCR) and FCIS fatigue questionnaire scores; 2B: Correlation between OCR and GFI-MAF questionnaire scores; 2C: Correlation between OCR and FSS questionnaire scores; 2D: Correlation between OCR and Bowman category eight questionnaire.

### Transcriptional Landscape of SjD: IFNs and Mitophagy

Using a large, preprocessed whole-blood microarray dataset downloaded directly through the Gene Expression Omnibus (Accession No. GSE51092), we initially compared all samples from SjD patients (S_1_; n=190) against those from healthy subjects (H_0_; n=32). In total, we identified 56 differentially expressed (DE) transcripts (44 up / 12 down with *p_adj_*<0.05 and |log_2_FC|>1), dominated as one would anticipate by upregulated IFN-stimulated genes (ISGs; table 1, figure 3A and online supplemental figure 2).

**Figure 3:**
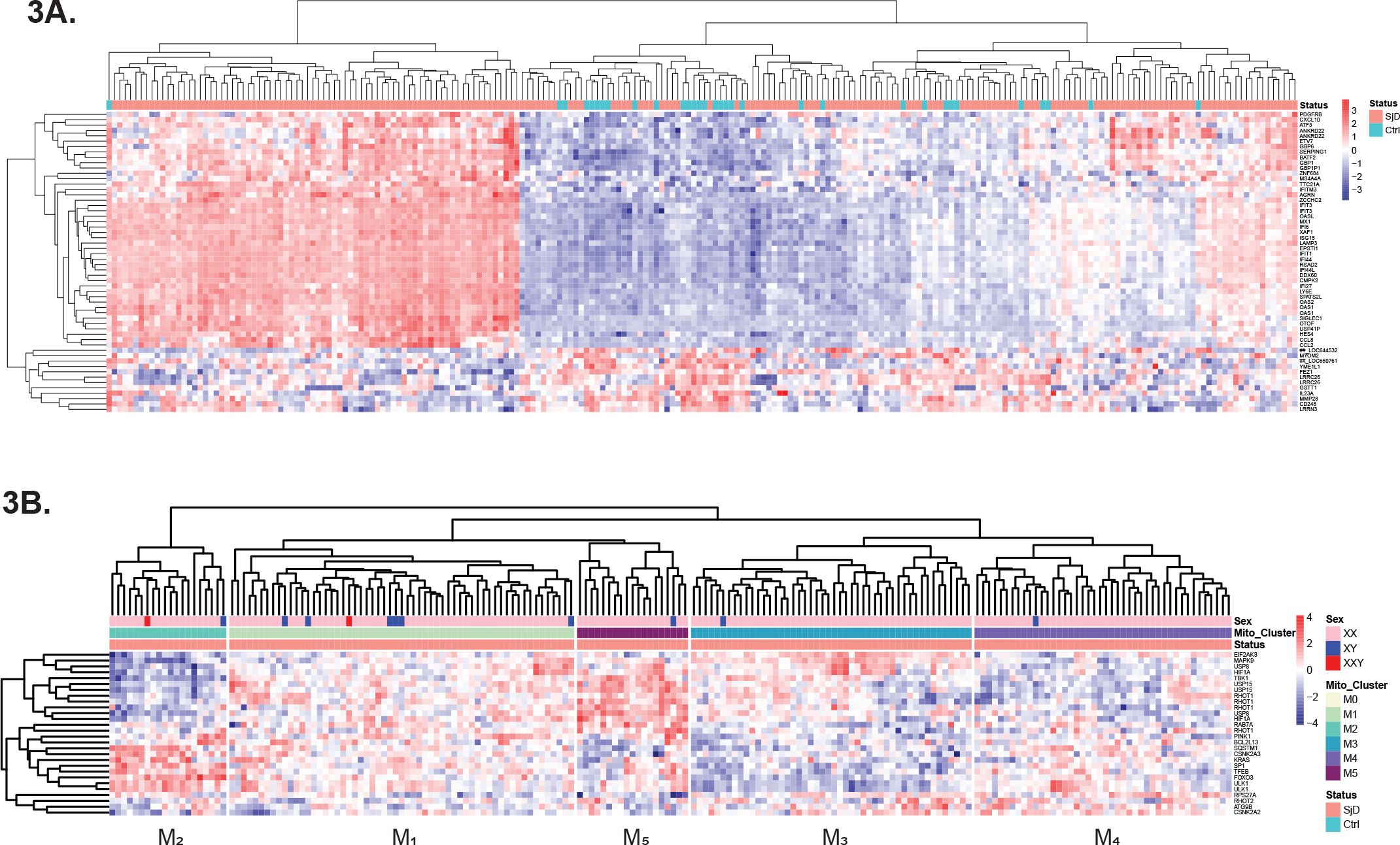
Heatmaps showing differentially expressed transcripts from the comparisons between SjD and healthy subject for: 3A) all transcripts meeting *p_adj_*<0.05 and |log_2_FC|>1 from results of overall analysis (S_1_); 3B) extracted mitophagy transcripts meeting *p_adj_*<0.05 for SjD cases only revealed 5 discrete SjD mitophagy clusters (as indicated) employing unsupervised average hierarchical clustering.

**Table 1.**
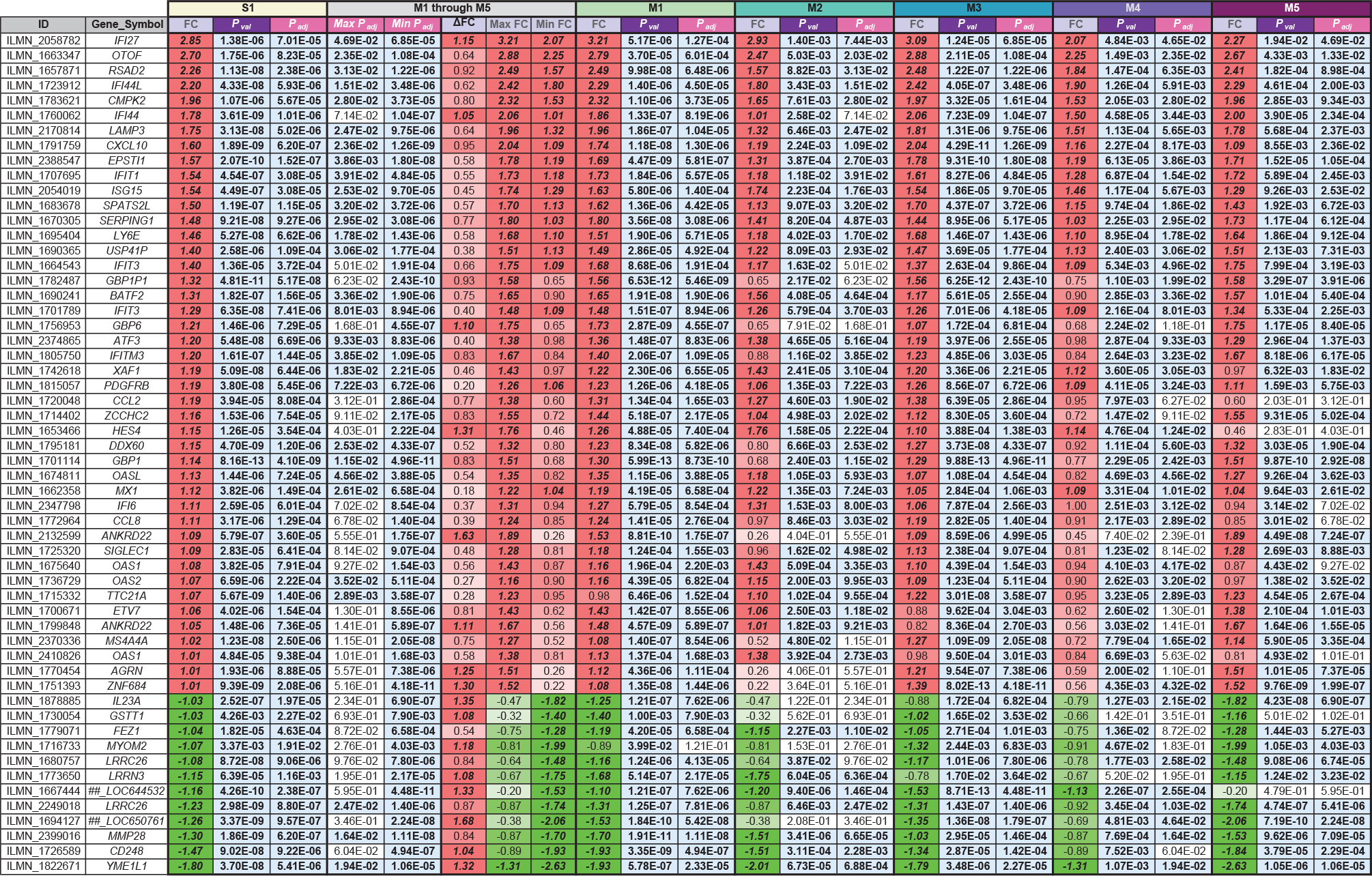
All differentially expressed probes in the overall S1 analysis (SjD vs. Healthy).

**Table 2.**
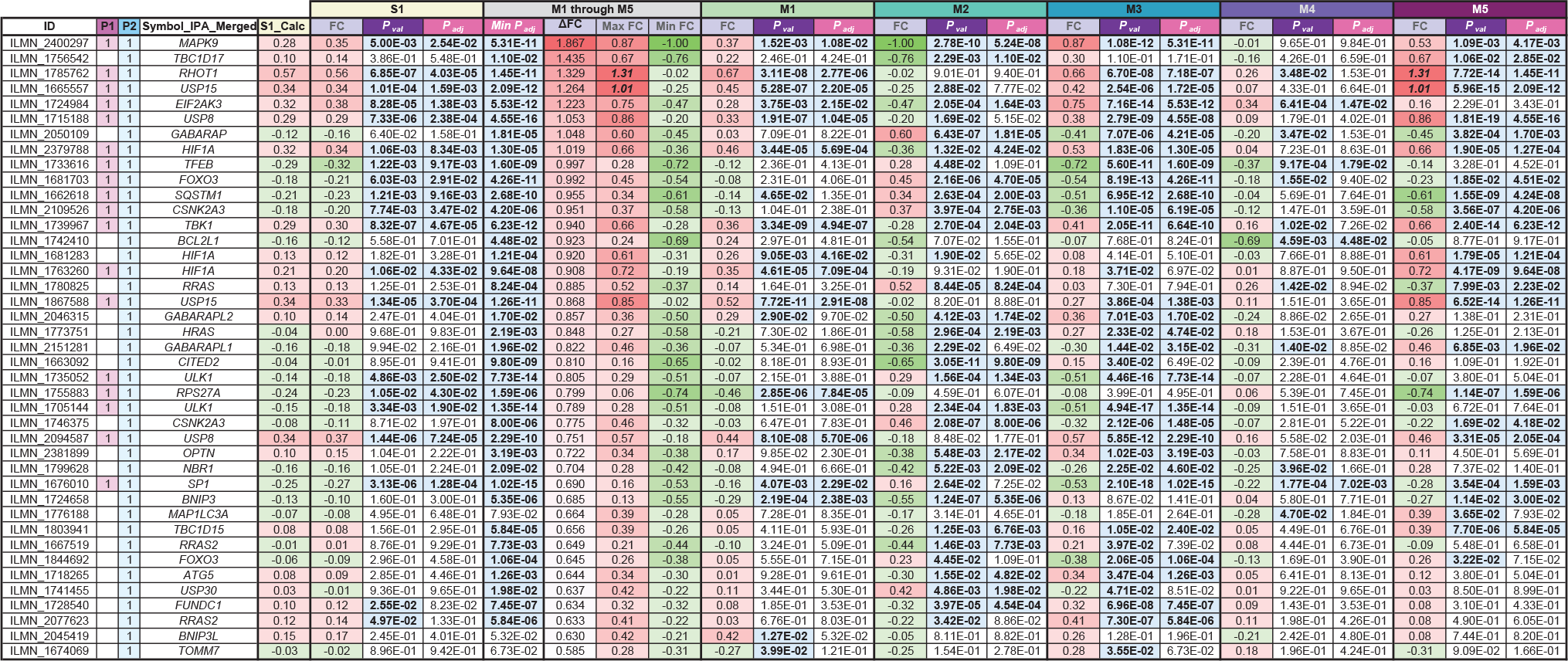
Mitophagy probes across S_1_ and M_1_ through M_5_ DEA Results.

In pursuit of mitophagic transcriptional signatures in SjD, we first sorted all transcript probes showing *p_adj_*<0.05 in the S_1_ analysis (n=371) by descending *p_adj_* and log_2_FC and hit the first mitophagy probes a rank of 46^th^ and 125^th^, respectively. To better understand the global impact of mitophagic processes in SjD, we isolated and extracted expression data for those mitophagy probes showing *p_adj_*<0.05 in the overall S_1_ analysis (28 out of 85 available). Heatmap visualization of these probes in SjD subjects alone followed by unsupervised average hierarchical clustering led us to identify discrete patient clusters of shared whole-blood mitophagic transcriptional signatures (figure 3B). Notably, we observed that healthy subjects appeared more likely to cluster near SjD patients belonging to clusters M_2_ and M_4_ (online supplemental figure 3A), and healthy subjects alone do not appear to have mitophagic transcriptional signatures as variable as those seen in SjD subjects (online supplemental figure 3B).

Notably, some clusters showed widely opposing patterns of mitophagy gene expression (figure 4). Given the degree of statistical difference between those mitophagy clusters showing oppositional transcription, we hypothesized that transcripts owing their dysregulation to mitophagic differences might influence the composition of DEGs amongst SjD subjects. To test this, we performed subgroup DEA by comparing each group of SjD patients stratified according to mitophagy cluster involvement against all healthy subjects (e.g., M_1_ vs. H_0_, M_2_ vs. H_0_, etc.). A total of 565 transcript probes showed *p_adj_*<0.05 & |log_2_FC|>1 within at least 1 mitophagy cluster, and we noted that large numbers of DE transcripts overlapping across mitophagy clusters (figure 5; online supplemental table S4).

**Figure 4:**
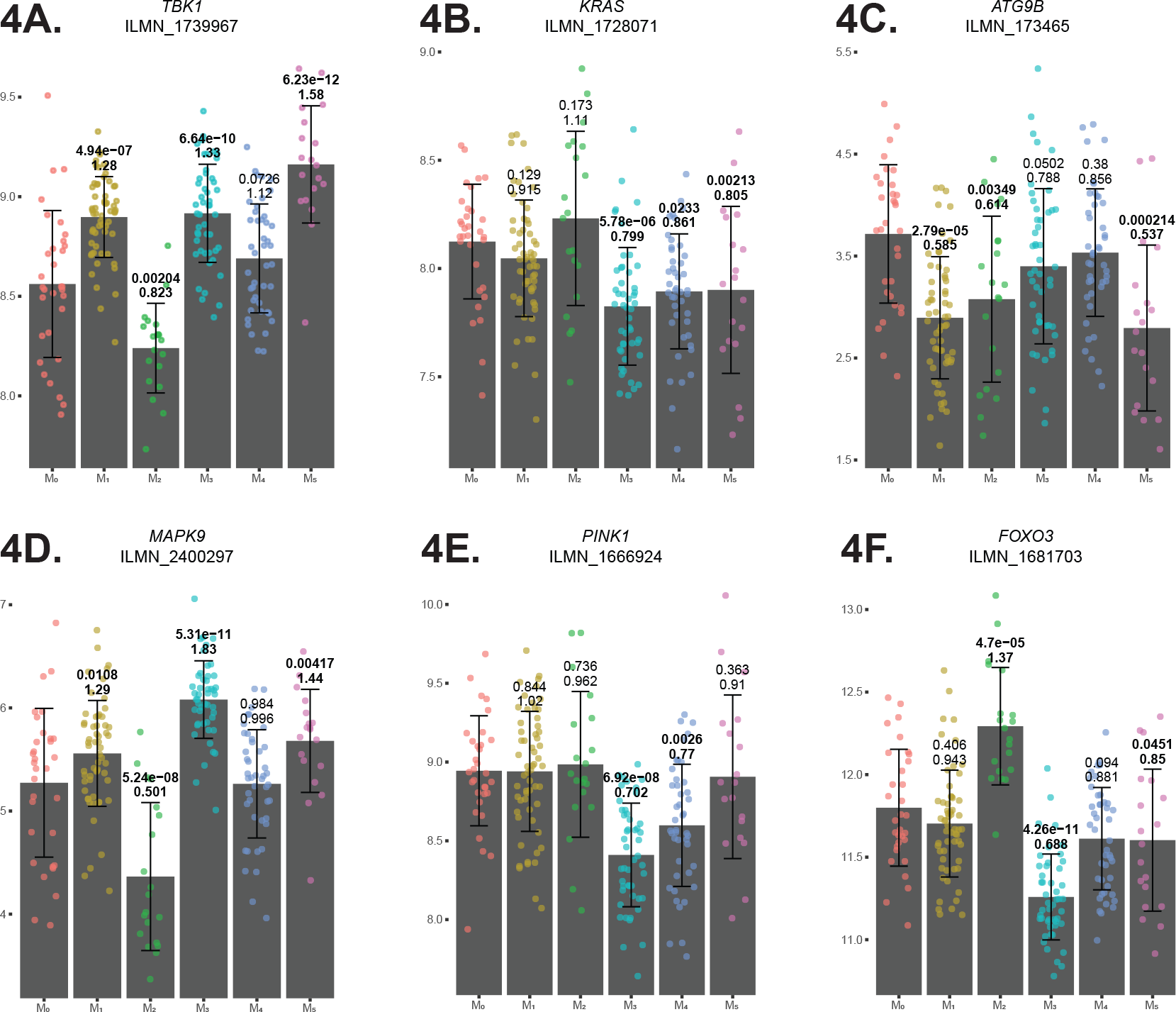
Plots for selected mitophagy transcripts showing *p_adj_*<0.05 stratified by group according to mitophagy cluster. The displayed log_2_FC results and *p_adj_* values were taken from the differential expression analysis results specific to each mitophagy cluster subgroup analysis when compared to all healthy subjects. Results are displayed for *TBK1* (4A), *KRAS* (4B), *ATG9B* (4C), *MAPK9* (4D), *PINK1* (4E), and *FOXO3* (4F). All results with *p_adj_*<0.05 are shown in bold, with the mean +/- standard deviation shown for each cluster.

**Figure 5:**
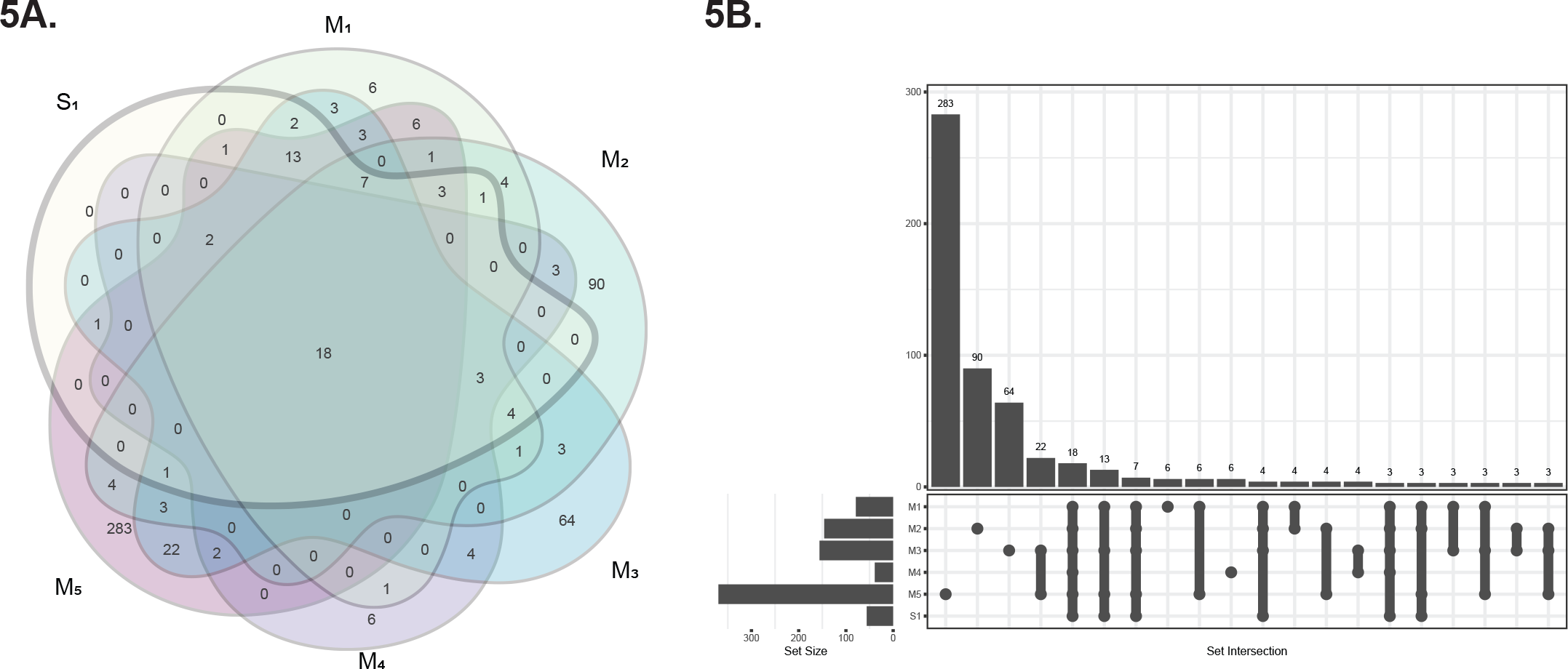
Venn diagram (figure 5A) and UpSet plot (figure 5B) showing overlap off differentially expressed transcripts (*p_adj_*<0.05 and |log_2_FC|>1) between the S_1_ and M_1_-M_5_ differential expression analyses. Few transcripts were unique to S_1_, suggesting that the M_1_-M_5_ analyses adequately capture most represented transcripts from the overall results.

From results of bioinformatic assessments using gene set enrichment analysis (GSEA) applied to these DEGs, we observed the enrichment of highly similar pathways between clusters, but also distinctions. For all mitophagy clusters, GSEA to assess for enrichment of KEGG and GO pathways overwhelmingly show enriched cellular and biological processes related to viral replication and immune responses, primarily driven by ISGs (online supplemental figures S4 and S5). Yet we observed several instances wherein the same pathways are significantly enriched in multiple mitophagy clusters, but with sometimes varying gene targets and normalized enrichment scores (NES) that flip from positive to negative, highlighting the dramatic nature of these transcriptional shifts in related pathways between clusters (online supplemental tables S2 and S3). Notably, in M_5_ we observed the significantly down-regulated enrichment of processes related to mitochondrial gene expression and respiratory chain assembly (online supplemental figures S4E and S5E).

To better discern those transcriptional differences attributable to mitophagic over IFN signatures in SjD, we calculated ΔFC, which defines maximal inter-cluster differences (see methods), sorted in descending fashion (tables 1 and 2; online supplemental table S4). Although the vast majority of ISGs were pushed to the bottom of the list owing to minimal differences conferred by mitophagy stratification, striking differences were noted in select ISGs, which show opposing differential expression. Most notably, *IFNG*, the Type II IFN (IFN-II) is down in M_2_ despite being up in M_1_/M_3_. Of further interest, several transcripts exhibit adjacency or frank overlap with immune-relevant transcripts and will require further scrutiny to determine if any enhancer/promoter elements are shared and demonstrate coordinated dysregulation: *RNF175* (up in M_2_, down in M_3_/M_5_), lies head-to-head with *TLR2*; *IL24* (up in M_2_, down in M_1_/M_3_/M_5_), neighbors *FCMR* (Fc mu receptor); *LYSMD1* (up in M_2_) overlaps directly with *TNFAIP8L2. CD69* is down in M_2_, up in M_3_/M_5_; *IL5RA* trends up in M_2_, down in M_5_. *DDX11* is down in M_2_, up in M_3_. *NOCT* is up in M2 (CCR4L). While *TBK1* does not show frank DE using the most stringent *p_adj_*and FC cutoffs, its binding partner *TBKBP1* shows DE in M_5_ and shows FC patterns highlighting its dynamic activity as well.

### Differential Expression of Sex-Specific Transcripts in SjD

One incidental finding from the present study involves the frank emergence of two sex-specific transcript probes, corresponding to *XIST* and *RPS4Y1*, that showed visually striking anticorrelated transcription and altered transcription relative to most SjD patients (data not shown). Using probes tagging these sex-specific transcripts to infer the biological sex of study participants, we identified 2 individuals out of 190 SjD patients who appear to be males with Klinefelter syndrome (47,XXY) owing to *XIST* expression at levels matching female patients (46,XX, n=174) while having *RPS4Y1* that matched males (46,XY, n=14; figure 6A). Removing all individuals with transcriptional evidence suggesting the presence of a Y chromosome (n=16) and stratifying again by mitophagy cluster to ascertain any potential differences, we found the differential expression of *XIST* in the M_2_ cluster, which was down compared to M_3_/M_4_/M_5_ and tighter compared to other clusters (figure 6B).

**Figure 6:**
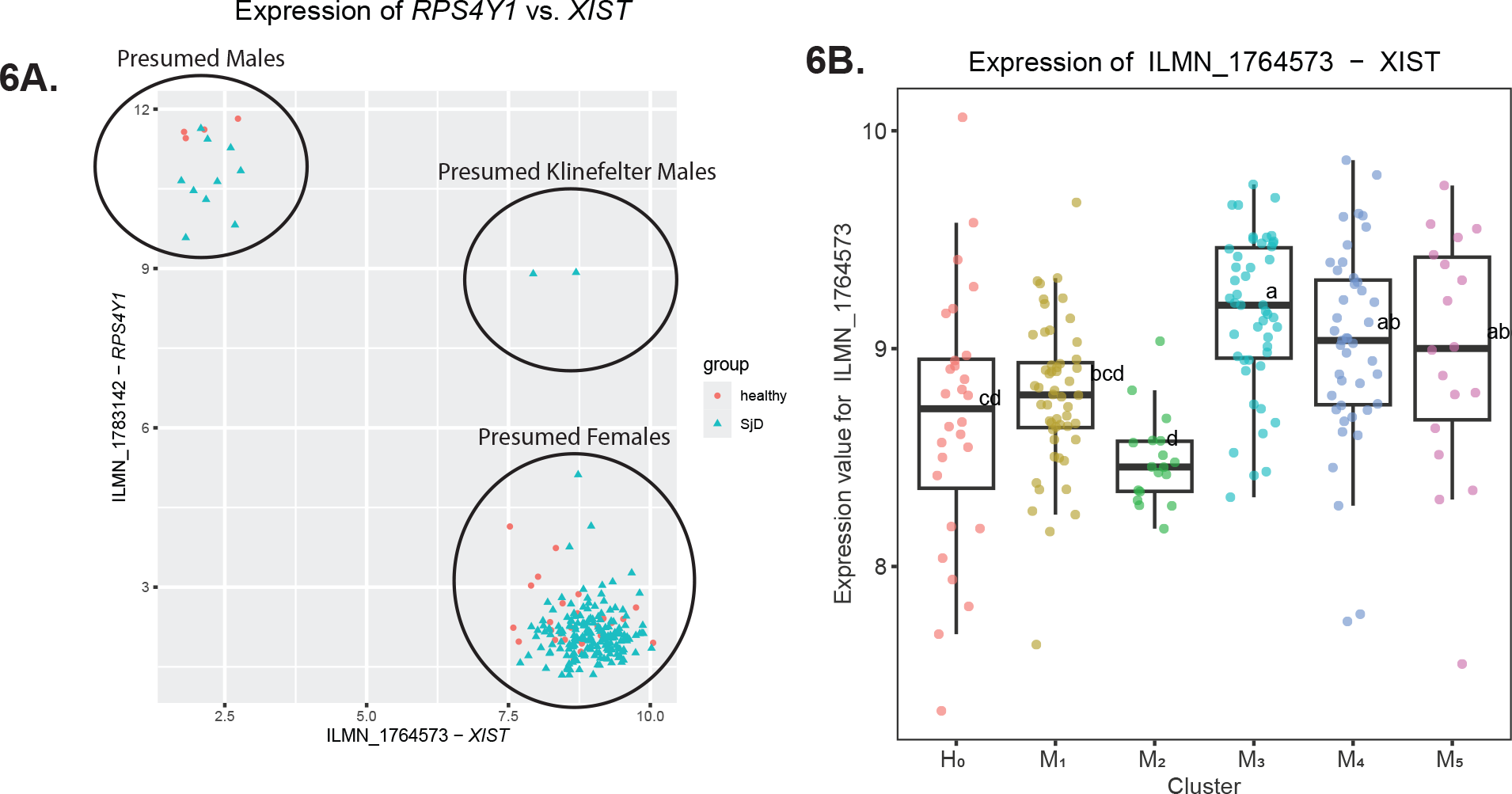
Figure 6A: Plot of transcriptional data for *RPS4Y1* (on chromosome Y) versus *XIST* (on chromosome X) using all healthy and SjD subjects. Two individuals were found to transcribe high levels of both *RPS4Y1* and *XIST*, suggesting that these individuals are Klinefelter syndrome males (47,XXY; n=2). Those expressing *XIST* exclusively were presumed to be female (46,XX; n=206) while those who expressed *RPS4Y1* exclusively were presumed to be male (46,XY; n=14). Figure 6B: Boxplot of *XIST* transcriptional levels in all subjects after removal of all individuals with any evidence for Y transcription and stratified by mitophagy cluster. We observed transcriptional differences between SjD mitophagy clusters, with SjD patients in M_2_ showing significantly lower and tighter *XIST* transcription compared to their peers in M_3_, M_4_, and M_5_. Compact letter display (CLD) showing results of one-way ANOVA with post-hoc Tukey’s test for multiple treatments is shown.

## Discussion

While the generation of ATP by oxidative phosphorylation is the main physiological function, mitochondria play additional physiologic roles including: 1) producing and detoxifying reactive oxygen species; 2) partaking in some types of apoptosis; 3) regulating calcium in the cytoplasm and mitochondrial matrix; 4) metabolite synthesis/catabolism; and 5) the transport of the organelles to specific intracellular sites. Mitochondrial dysfunction occurs when any of these processes becomes abnormal^23^ and contributes to the pathogenesis of autoimmune disorders like multiple sclerosis,^46^ rheumatoid arthritis,^47^ type I diabetes,^48^ and SLE.^49,50^

Increased oxidative damage has been reported in autoimmune diseases like SLE and SjD.^27,51–58^ As mitochondria represent a significant source of free radicals in the human body, we hypothesized that mitochondrial function would be affected in SjD. Furthermore, we chose to study T cells in SjD given prior work in SLE showing significant mitochondrial dysfunction in these cells.^59,60^ Studies have shown that free radical-mediated oxidative damage occurs in SLE, an autoimmune condition with many immunologic similarities to _SjD._52,54,61-63

One characteristic of SLE includes exclusive expansion of a CD4^+^ T helper subpopulation in children with SLE resulting from the accumulation of succinate-driven mitochondrial ROS (mtROS).^60,64^ Lymphocytes in SLE are also severely depleted in reduced glutathione (a tripeptide consisting of cysteine, glycine, and glutamic acid; abbrev: GSH), which leaves them susceptible to oxidative damage^49,60,65^ and likely contributes to mitochondrial dysfunction. Oxidized mitochondrial DNA in SLE neutrophils can be extruded into the cytoplasm where they can serve as powerful IFN stimuli.^63,66^ Excess IFN-I production (especially IFN-α) promotes SLE pathogenesis by activating and expanding autoreactive immune cells to induce production of autoantibodies and inflammatory cytokines, ultimately leading to nucleic acid-protein immune complex formation, deposition within tissues like the kidneys, and perpetuation of autoreactive inflammatory immune responses.^67,68^ By similar mechanisms, IFN production has been implicated in the pathogenesis of SjD^69^.

Oxidative stress-related biomarkers and proinflammatory cytokines are overexpressed in SjD.^16^ The glandular inflammatory activity, in particular, has been shown to increase linearly higher for IL-6 and IL-17 levels.^70^ Studies using deep bioinformatics data mining and pathway analyses of differentially expressed transcripts in whole blood point to the regulatory impairment of long noncoding RNAs, especially *LINC01871,* as an effective mechanism triggering the IFN-related and/or T cell-associated pathology of SjD.^71^ NK cells from the SG of an animal model of SjD were shown to upregulate IFN-γ with the gland and play an essential role in developing autoimmune lesions.^72^ Yet another study used transcriptomic datasets from SjD patients’ salivary gland samples to identify three subtypes of salivary gland tissues having distinct cellular and molecular characteristics: i) oxidative phosphorylation-dominant; ii) weak inflammatory with Type I IFN signatures; or iii) B cell receptor signaling pathway-dominant.^73^

Prior work has associated the autophagy-related genes *ATG5*, *ATG7*, and *IRGM* with autoimmunity^74–77^. Naïve *Irgm1*^−/−^ mice, for example, exhibit several SjD characteristics, including lymphocytic infiltration of the lungs, salivary and lacrimal glands, and the exocrine pancreas in addition to the presence of increased serum autoantibodies/cytokines^78^. Deficiency of IRGM1 incites IFN-I interferonopathy involving several organs and having autoimmune features. Mice without the GTPase IRGM1 (an IRGM homologue) also displayed similar features, including decreased mitophagic activity. Damage to mitochondria by free radicals releases PAMPs like mtDNA into the cytoplasm, which can activate the cGAS–STING-dependent IFN-I pathway, as observed in the fibroblasts of *Irgm1*^−/−^ mice; in macrophages, however, this activates lysosomal TLR7. Deletion of cGAS and STING abolishes the salivary and lacrimal gland autoimmune pathology and attenuates lung pathology.^16,79^

Diminished levels of reduced glutathione can also bring about mitochondrial hyperpolarization of human T cells in SLE.^80–82^ Mitochondrial hyperpolarization also happens during T cell activation. CD3/CD28 stimulation temporarily increased mitochondrial transmembrane potential and ATP exhaustion and made the cells sensitive to hydrogen peroxide-mediated necrosis. Thus, repeated SLE T cell activation *in vivo* can bring about continued mitochondrial hyperpolarization and subsequent decreases in ATP.^49^ Consequently, abnormal T cell activation and cell death in patients with SLE are mainly attributable to mitochondrial hyperpolarization and depletion of ATP.^49^ That mitochondrial dysfunction occurs in SLE is also seen from the enlarged mitochondria observed in the peripheral blood lymphocytes from SLE patients. In addition, mitophagy, the programmed removal of damaged mitochondria, is defective in SLE.^83,84^ Since oxidative stress and mitochondrial damage have been reported in the pathogenesis of SLE and SjD^16,17,85^, we investigated mitochondrial dysfunction in the T cells of SjD and healthy subjects.

Studies have shown that morphological changes of salivary epithelial mitochondria are involved in altered cellular bioenergetics in SjD. Transmission electron microscopy work has shown cytoplasmic lipid droplets and swollen mitochondria salivary gland epithelial cells in SjD.^14^ In addition, this study showed altered levels of the genes associated with the degree of salivary gland immune cell infiltration, mitochondrial respiratory chain complexes, and the TCA cycle in SjD.^14^ Autophagy has also been shown to be impaired in SjD.^15,86^ Using this technique, we observed that SjD lymphocytic mitochondria underwent mitochondrial matrix swelling and loss/disorganization of crests compared to mitochondria in lymphocytes from the healthy subject. We further observed decreased basal respiration, ATP-linked respiration, maximal respiration, and reserve capacity in SjD subjects compared to control subjects, values derived from the induced inhibition of complex 1, complex III, and the use of the uncoupler FCCP. Studies with mitochondria isolated from SLE PBMCs showed decreased activity of Complex I, IV, and V in SLE compared to healthy subjects.^87^

Under normal physiological conditions, damaged/dysfunctional mitochondria generating ROS at high levels are removed by mitophagy. Yet changes in mitophagy can result in the buildup of damaged, ROS-producing mitochondria and give rise to inflammation.^88^ Though the authors of a recent study observed changes in autophagy in the salivary glands of SjD patients, this work did not assess mitophagy-specific markers.^89^ Since dysfunction could be inferred from a changed mitochondrial morphology, damaged mitochondria are thought to trigger inflammation by releasing mitochondrial DAMPs. Some peculiar characteristics make the mitochondria behave like highly immunogenic organelles. These include the following; a) The circular mtDNA genome does not contain histones, has fewer introns and polycistronic genes, and features hypomethylated CpG motifs;^90^ b) The phospholipid cardiolipin is found in prokaryotes but in the eukaryotic cells is found only in the inner mitochondrial membrane;^90,91^ and c) Like in bacteria, the mitochondrial protein translation starts with a formylated methionine, an N-terminal modification that does not occur on proteins encoded in the nuclear genome.^90^

Considering the prominent role of mitochondria in generating ROS,^20^ antimitochondrial antibodies and oxidative stress markers cut across many autoimmune diseases.^62,92–95^ Interestingly, 3-27% SjD subjects have anti-mitochondrial antibodies,^96^ and many also show elevated salivary mitochondrial glutamic oxaloacetic transaminase (m-GOT) levels.^85^ Chronic fatigue is a serious concern in SjD, bringing with it severe declines in quality of life and highlighting the possibility of a mitochondrial chemical energy disorder.^97^ These observations point to the contribution of mitochondrial dysfunction to inflammation observed in autoimmune diseases like SjD.

Fatigue may involve both mental or physical signs and symptoms and is akin to a lack of energy (derived from the elemental metabolic oxidation of food). While the brain is a crucial user of resting cellular energy, mental fatigue is a subjective sensation exemplified by diminished motivation and alertness. When metabolism works optimally, fatigue can result from energy wasted secondary to the psychological and physical nature of stress, apprehension, tension, and depression.^98^ With compromised mitochondrial function, however, as supported by our present study, fatigue could be a direct consequence of diminished energy production. The fatigue scores that showed significant correlation with mitochondrial function involved pain/discomfort the subjects experienced within the prior 2 weeks, as well as some correlation with fatigue severity and mental attitude. Bowman general fatigue questionnaire (questions related to pains/discomfort) had a sensitivity of 85% and a specificity of 55% for SjD. Bowman mental fatigue questions about concentration had a sensitivity of 70% and a specificity of 73% for SjD, while questions about memory had a sensitivity of 67% and a specificity of 70% for SjD.^47^ Brain fog and diminished cognition are salient issues in SjD.^99^ A decrease in ATP production and impaired mitophagic recycling resulting from mitochondrial dysfunction has been shown to contribute to chronic fatigue syndrome in a subset of SjD patients.^107^ Hyperthyroidism uncouples the mitochondrial oxidation-phosphorylation process and can lead to fatigue.^100^

While the dominance of the IFN signature over the SjD transcriptional landscape has broadly resulted in concerted efforts to elucidate concrete functional mechanisms of these clearly dysregulated pathways, ancillary pathways with potentially profound pathogenic influence warrant deeper attention and investigation. With this study, we demonstrate the usefulness of extracting gene sets (as with mitophagy) from larger GEP datasets to contextualize transcriptional relationships and delineate the biological implications of any clusters that emerge. By performing DEA in SjD mitophagic subgroups (M_1_-M_5_), we reorient the data to highlight critical transcriptional differences between the groups, but also remove the counter. Indeed, rather than observing coordinated, homogeneous transcriptional activity across all DEGs, we find a high degree of transcriptional variability with groups of SjD patients showing coordinated transcriptional patterns in key mitophagic genes that we hypothesize have mechanisms of control whose elucidation would improve our understanding of SjD pathogenesis and etiology. Indeed, those transcripts with the greatest between-cluster differences in mitophagic activity show tremendous potential for helping us to elucidate pathogenic mechanisms in SjD beyond the IFN signature, particularly when faced with a paucity of available clinical information.

While we expected that stratification of SjD patients by mitophagy signature would show the opposing directional transcription of key mitophagic transcripts, the degree by which we observed divergence from the original FC and *p_adj_*values in those mitophagy transcripts that moved more with the overall SjD patterns was striking (e.g., *TBK1)*. TANK-binding kinase 1 (TBK1) is an IKK-related serine/threonine kinase with involvement in innate antiviral responses that operates downstream of the RIG-like receptors (RLRs) and the DNA-sensing receptors (DSRs) through induction of IFN-I with an increasingly defined relationship to humoral autoimmunity.^101^ In IFN-I-positive SjD, SLE, and SSc patients, whole-blood RNA showed increased transcription of *TBK1* and phosphorylation of TBK1 (pTBK1) in plasmacytoid dendritic cells (pDCs), the major producers of IFNs, and PBMCs from these patients treated with the inhibitor BX795 downregulated IFN-I pathways.^102^ We can readily establish a relationship with TBK1 and METTL3 to define a role in autoimmunity. Activation of IRF3. TBK1 was previously shown to be upregulated in IFN-I SjD. The gene METTL3 is involved in post-transcriptional m^6^A modifications to mRNA molecules to stabilize them by preventing their degradation. TBK1 phosphorylates METTL3 at Ser67 which leads to METTL3 interaction with the protein translational complex to facilitate antiviral responses and results in stabilized IRF3 mRNA via m^6^A modification.^103^ CD69 is a cell surface molecule expressed on activated T cells, interaction with Myl9 is required for induction of inflammatory diseases ^104^. Based on CD69 and CD103 expression, CD8^+^ T cells with a tissue-resident memory phenotype were predominant amongst infiltrating cells adjacent to salivary duct epithelial cells.^105^

Clearly, when taking an overall “case vs. control”, lump-sum approach to identifying DEGs, certain transcripts, perhaps resulting from distinct molecular mechanisms found within specific endotypes, can obscure the transcriptional contribution of their peers, which distorts the transcriptional landscape that defines SjD. While we initially had concerns that by stratifying in this way would abrogate the mitophagy signal present between clusters, we in fact find that the mitochondrial signals emerge in GSEA for cluster M_5_. Importantly, the influence of the ISGs also persists despite stratification by mitophagy signature, which highlights the shared pathological basis of IFN dysregulation in SjD. Yet by stratifying in this way, we were able to more fluidly observe that seemingly non-descript, yet discernable, breaks in heatmaps can conceal important transcriptional dynamics operating well below the towering heights of the IFN signatures that ultimately must be accounted for.

In the present study, all recruited participants were female to reflect the marked female bias observed in SjD. While it was not our original goal to ascertain biologic sex by GEP, these data provide an opportunity to study the relationship between mitophagy and biological sex within the context of complex autoimmunity as in SjD. The fact that *XIST* and *RPS4Y1* show up as DEGs in M_3_ results is intriguing given recent work in which GEP data derived from parotid glands of SjD patients was reanalyzed and showed not only these transcripts, but several additional correlated transcripts from the X and Y chromosomes, including *KDM5D*, *EIF1AY*, *USP9Y*, *TXLNGY*, *TAC1*, and *COMMD9*. The protein lysine demethylase 5D *(KDM5D)* is a male-specific, Y-linked H_3_K_4_ histone demethylase whose upregulation driven by KRAS-mediated activation of the STAT4 transcription factor^106^. Future research efforts will involve confirming the inferred biological sex directly and segregating these individuals to better delineate the effect of biological sex on mitophagic processes in SjD.

Our work highlights the importance of altered mitochondrial dynamics and compromised respiratory chain condition on fatigue in SjD. Based on our data, we postulate that autoimmune-related mitochondrial damage associated with the onset of a pro-oxidant environment is implicated in SjD pathogenesis. Certainly, as with any study, there are limitations. Certainly, expansion of these studies to larger cohorts of SjD patients will be essential to confirm our findings and map the spectrum of possible mitochondrial metabolic phenotypes in SjD. Additionally, replication with independent sets of GEP datasets (microarray and/or RNA-seq) will be required but will also provide opportunities to identify the contributions of additional immune cell subsets, including B, NK, and dendritic cells (DCs). Further, there will be opportunities to distinguish between mitochondrial dysfunction in systemic, circulating immune cells and those that reside in the exocrine glands of SjD patients. Further, future work will establish *bona fide* clinical correlations that the authors hypothesize exist and constitute molecular endotypes of SjD attributable to transcriptional differences in mitophagic processes.

## Footnotes

### Deceased

KAK deceased on 02 May 2022.

## Acknowledgments

The authors would like to thank Dr. Lida Radfar, DDS, for her expertise and contributions to research conducted under the auspices of the Oklahoma Sjögren’s Syndrome Center of Research Translation. Across many years, Dr. Radfar helped us to evaluate and characterize SjD patients in addition to collecting vital samples that allow our research efforts to continue to this day. We would also like to thank our patients and study participants who so graciously assist us to advance SjD research with the shared hope that improved diagnostics and treatments will someday alleviate the tremendous burdens of this disease.

