## Supplemental Materials for "Mitochondrial Dysfunction and Fatigue in Sjögren’s Disease"

### Online Supplemental Figure S1

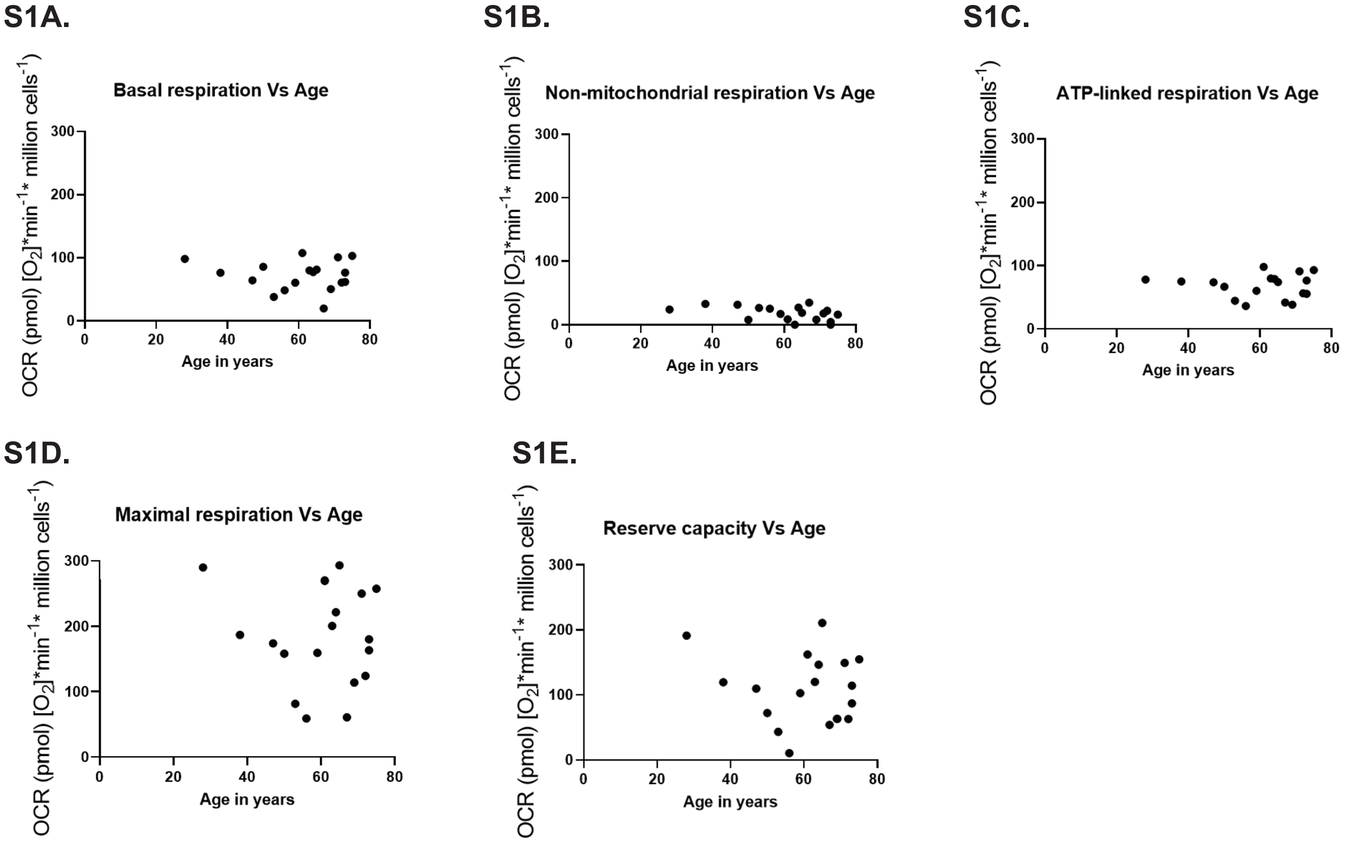


**Online Supplemental Figure S1:** Scatter plots of individual values of basal respiration (S1A) ,non-mitochondrial respiration (S1B), ATP-linked respiration (S1C), maximal respiration (S1D), and reserve capacity (S1E) of T cells from SjD subjects compared to their respective age. Oxygen consumption rate is given as pmol of oxygen consumed by one million T cells in one minute.

### Online Supplemental Figure S2

**S2A.**

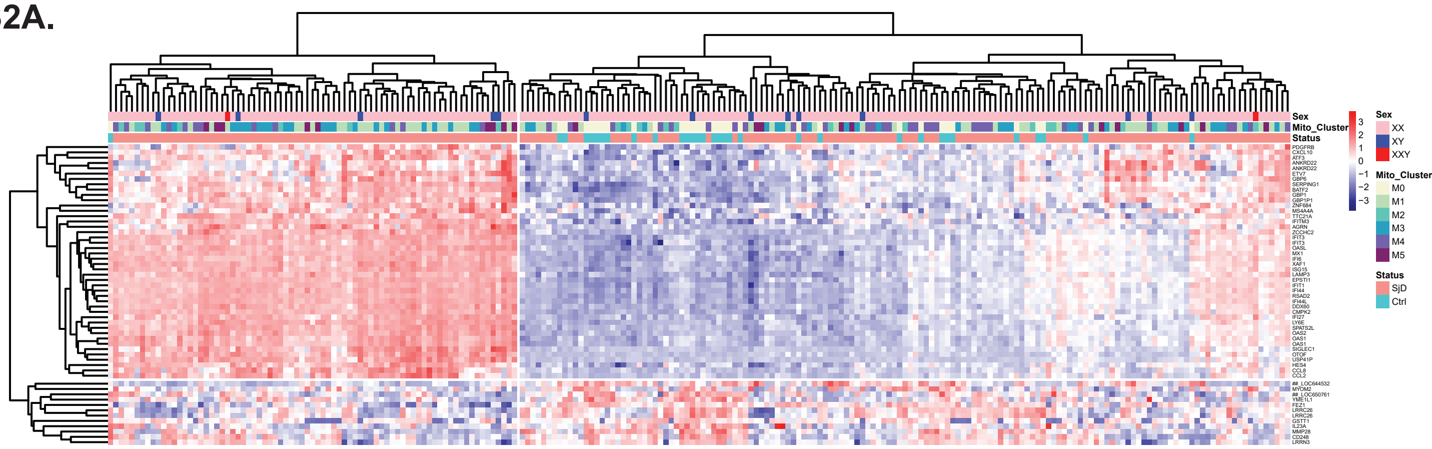


**S2B.**

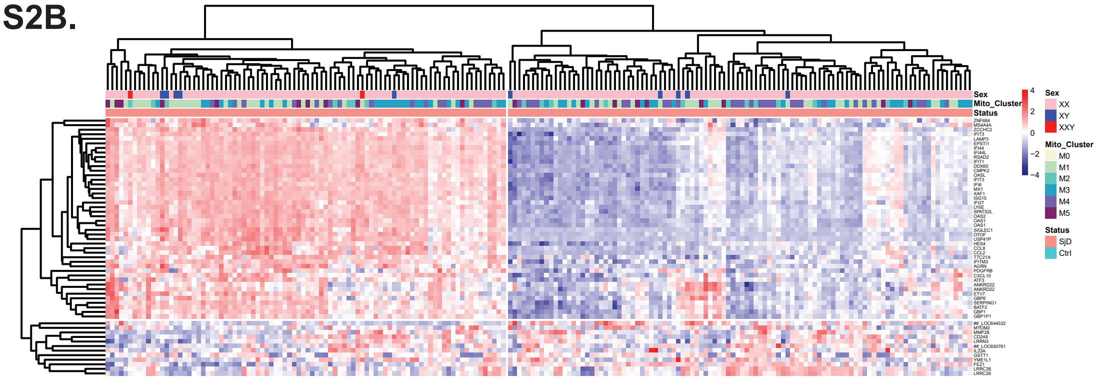


**S2C.**

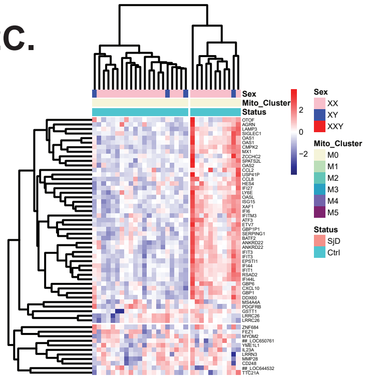


**Online Supplemental Figure S2:** Heatmaps showing differentially expressed transcripts meeting $p_{adj}$<0.05 and |$log_2$FC|>1 (n=56) from the comparison between all SjD and healthy subjects (overall $S_1$ analysis) for all subjects (S2A), SjD subjects alone (S2B), and healthy subjects alone (S2C).

### Online Supplemental Figure S3

S3A.

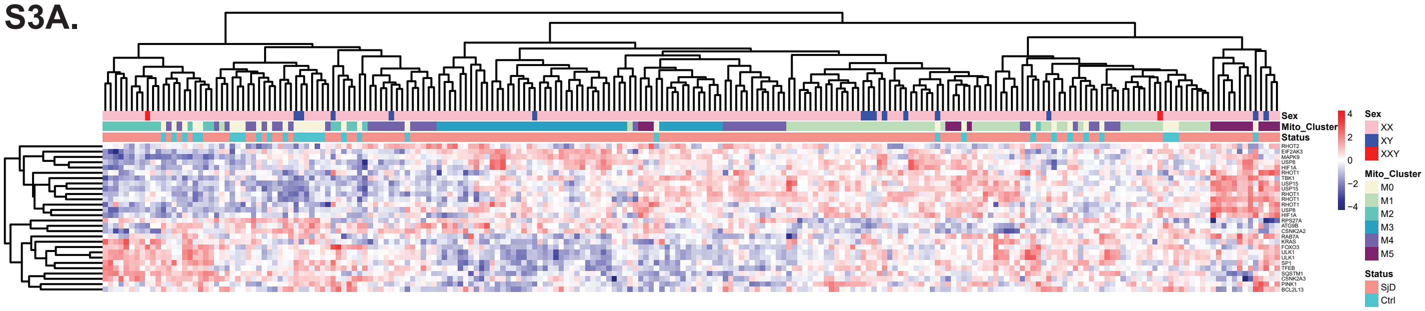


S3B.

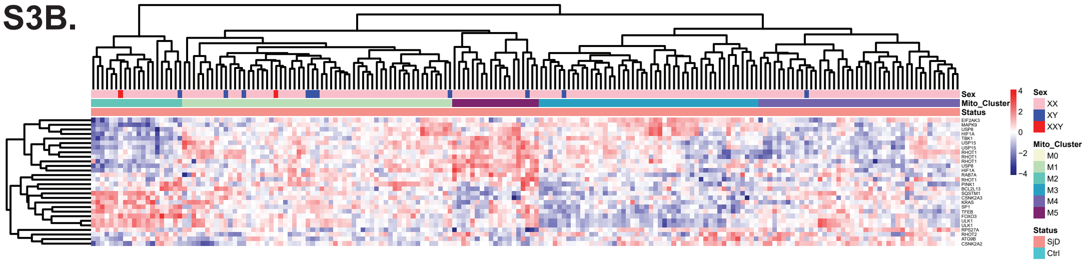


S3C.

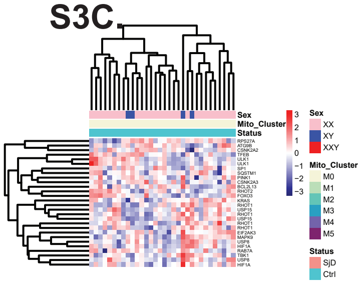


**Online Supplemental Figure S3:** Heatmaps showing extracted transcripts found within the KEGG mitophagy pathway and meeting $p_{adj}$<0.05 (n=28) when comparing SjD and healthy subjects (overall $S_1$ analysis) for all subjects (S3A), SjD subjects alone (S3B), and healthy subjects alone (S3C).

### Online Supplemental Figure S4

## S4A.

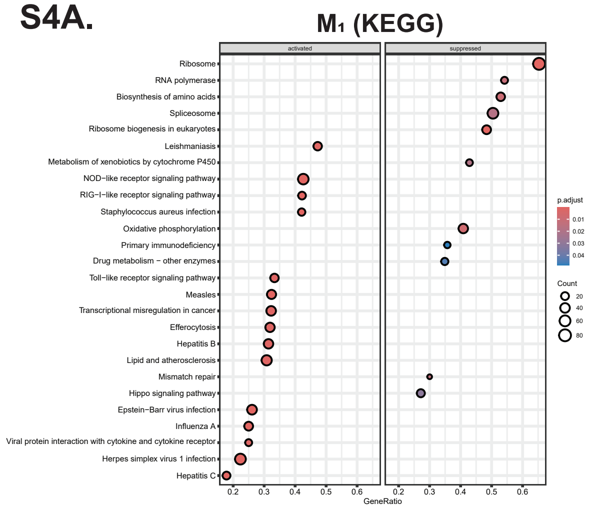


## S4B.

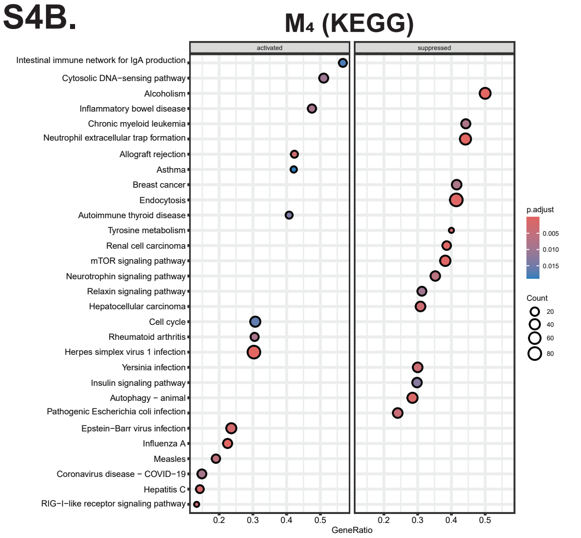


## S4C.

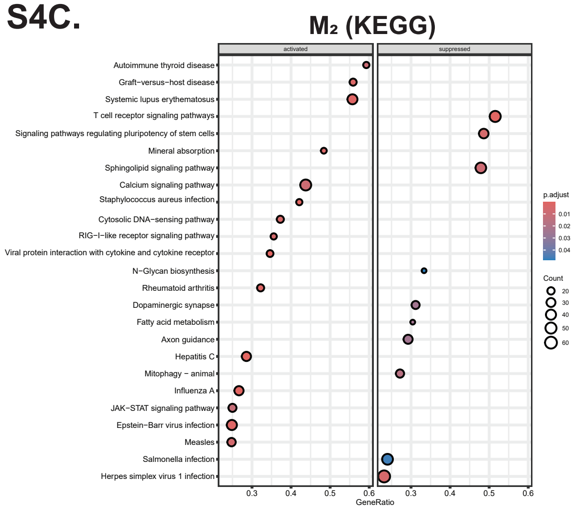


## S4D.

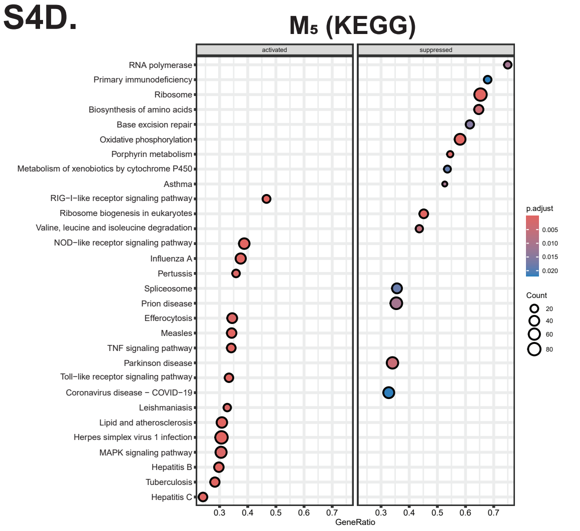


## S4E.

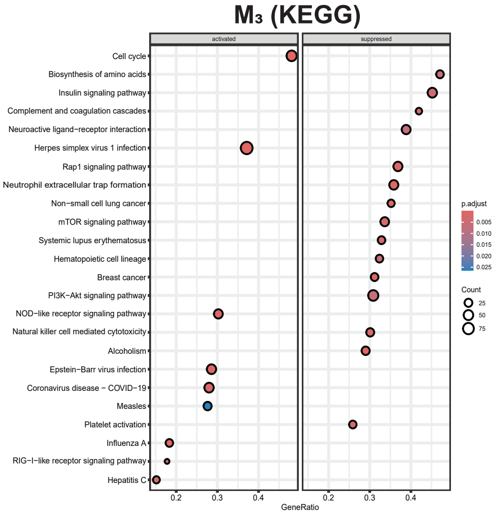


**Online Supplemental Figure S4:** Gene Set Enrichment Analysis (GSEA) results showing those significantly enriched KEGG pathways using gene expression data from clusters M$_1$-M$_5$, identified using $p_{adj}$<0.05.

### Online Supplemental Figure S5

**S5A.**

$M_1$ (GO)

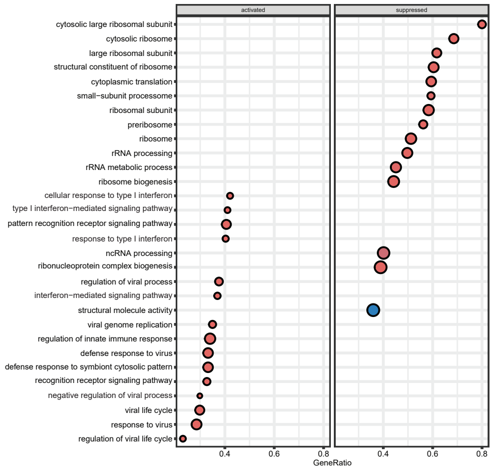


**S5B.**

$M_4$ (GO)

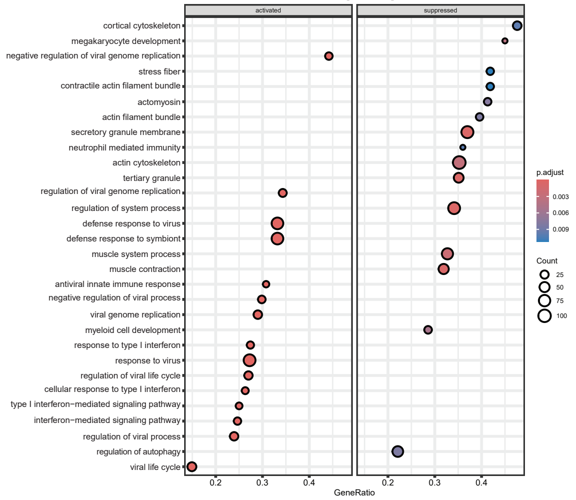


**S5C.**

$M_2$ (GO)

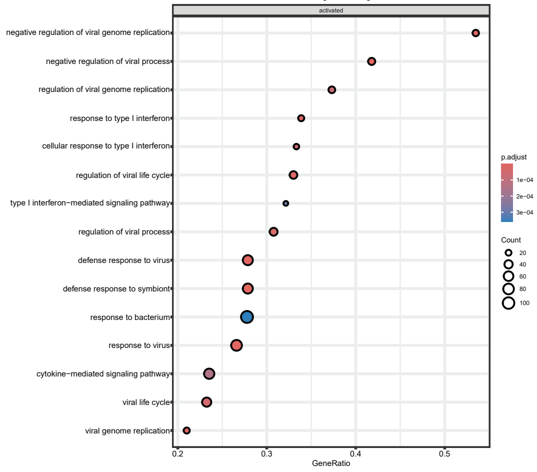


**S5D.**

$M_5$ (GO)

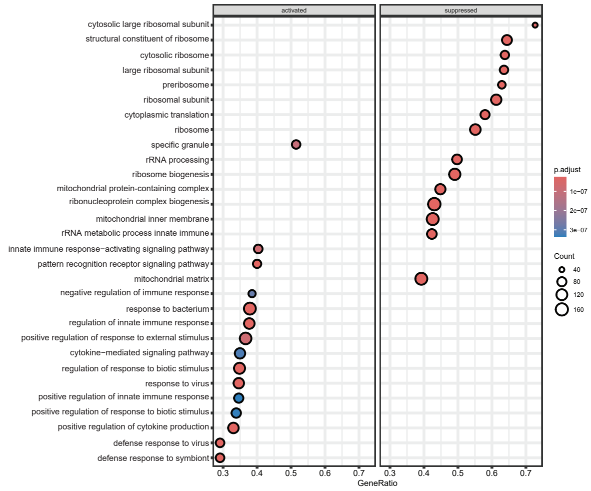


**S5E.**

$M_3$ (GO)

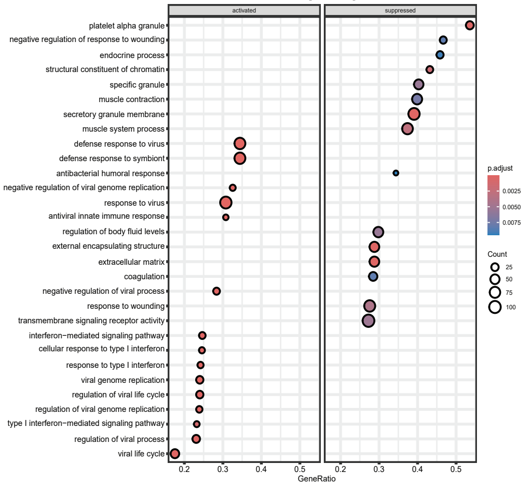


**Online Supplemental Figure S5:** Gene Set Enrichment Analysis (GSEA) results showing those significantly enriched GO pathways using gene expression data from clusters $M_1$-$M_5$, identified using $p_{adj}$<0.05.

| Age (Years) | Basal Respiration | Non-mitochondrial Respiration | ATP-linked Respiration | Maximal Respiration | Reserve Capacity |
|---|---|---|---|---|---|
| 28 | 98.43 | 24.1 | 78.19 | 289.76 | 191.13 |
| 38 | 76.37 | 32.91 | 75.08 | 187.09 | 119.73 |
| 47 | 64.28 | 31.58 | 73.79 | 174.18 | 109.9 |
| 50 | 85.98 | 7.95 | 66.86 | 158.4 | 72.26 |
| 53 | 38.15 | 26.73 | 44.61 | 81.62 | 43.47 |
| 56 | 48.61 | 25.28 | 36.45 | 59.12 | 10.51 |
| 59 | 60.69 | 17.19 | 60.43 | 159.61 | 102.94 |
| 61 | 107.6 | 8.7 | 98.15 | 269.9 | 162.4 |
| 63 | 80.15 | 0 | 79.81 | 200.86 | 120.17 |
| 64 | 77.32 | 27.11 | 79.02 | 221.84 | 146.52 |
| 65 | 81.37 | 18.78 | 74.15 | 293.2 | 210.85 |
| 67 | 19.78 | 34.94 | 41.89 | 60.96 | 54.09 |
| 69 | 50.57 | 8.16 | 38.4 | 114.05 | 63.48 |
| 72 | 60.96 | 22.07 | 56.38 | 124.43 | 63.2 |
| 73 | 76.53 | 0 | 76.53 | 163.66 | 87.13 |
| 73 | 61.75 | 4.33 | 55.56 | 180.45 | 114.37 |
| 75 | 103.09 | 16.07 | 93.03 | 257.85 | 154.76 |

**Online Supplemental Table S1:** Individual values of basal respiration, non-mitochondrial respiration, ATP-linked respiration, maximal respiration, and reserve capacity of T cells from SjD subjects compared with respective ages. Oxygen consumption rate is given as pmol of oxygen consumed by one million T cells in one minute.

### Online Supplemental Table S2: Overlapping significantly enriched KEGG pathways by GSEA showing directional change in the overall $S_1$ and $M_1$ through $M_5$ analyses.

| | | $M_1$ through $M_5$ | | | | | $S_1$ | | $M_1$ | | $M_2$ | | $M_3$ | | $M_4$ | | $M_5$ | |
|---|---|---|---|---|---|---|---|---|---|---|---|---|---|---|---|---|---|---|
| ID | Description | NES (Max) | NES (Max) Cluster | NES (Min) | NES (Min) Cluster | *Min Padj* | NES | *Padj* | NES | *Padj* | NES | *Padj* | NES | *Padj* | NES | *Padj* | NES | *Padj* |
| hsa05168 | Herpes simplex virus 1 infection | 2.076 | M3 | -1.442 | M2 | 1.356E-10 | 1.803 | 3.432E-06 | 1.749 | 1.359E-05 | -1.442 | 4.450E-03 | 2.076 | 1.356E-10 | 1.635 | 8.718E-05 | 1.849 | 1.910E-07 |
| hsa04613 | Neutrophil extracellular trap formation | 1.595 | N5 | -2.158 | M3 | 5.179E-08 | - | - | 1.578 | 4.390E-03 | - | - | -2.158 | 5.179E-08 | -2.065 | 2.664E-07 | 1.595 | 1.942E-03 |
| hsa05171 | Coronavirus disease - COVID-19 | 1.544 | M3 | -1.352 | M5 | 2.075E-04 | 1.715 | 2.075E-04 | 1.430 | 2.053E-02 | 1.391 | 1.367E-02 | 1.544 | 1.629E-03 | 1.405 | 8.971E-03 | -1.352 | 2.185E-02 |
| hsa05322 | Systemic lupus erythematosus | 1.750 | M1 | -1.613 | M3 | 1.163E-03 | - | - | 1.740 | 4.146E-03 | 1.750 | 1.163E-03 | -1.613 | 3.648E-03 | - | - | - | - |
| hsa00860 | Porphyrin metabolism | 1.617 | M2 | -1.843 | M5 | 1.545E-03 | - | - | - | - | 1.617 | 1.772E-02 | -1.589 | 2.530E-02 | -1.540 | 2.453E-02 | -1.843 | 1.545E-03 |
| hsa04610 | Complement and coagulation cascades | 1.751 | M5 | -1.800 | M3 | 2.045E-03 | - | - | - | - | - | - | -1.800 | 2.045E-03 | - | - | 1.751 | 3.082E-03 |
| hsa05135 | Yersinia infection | 1.429 | M5 | -1.541 | M5 | 3.748E-03 | - | - | - | - | - | - | - | - | -1.541 | 3.748E-03 | 1.429 | 1.755E-02 |
| hsa04080 | Neuroactive ligand-receptor interaction | 1.336 | M2 | -1.501 | M3 | 7.085E-03 | - | - | - | - | 1.336 | 3.641E-02 | -1.501 | 7.085E-03 | - | - | - | - |
| hsa04020 | Calcium signaling pathway | 1.439 | M2 | -1.280 | M3 | 8.105E-03 | - | - | - | - | 1.439 | 8.105E-03 | -1.280 | 4.100E-02 | - | - | - | - |
| hsa05310 | Asthma | 1.684 | M4 | -1.741 | M5 | 1.099E-02 | - | - | - | - | - | - | - | - | 1.684 | 1.891E-02 | -1.741 | 1.099E-02 |
| hsa04060 | Cytokine-cytokine receptor interaction | 1.436 | M2 | -1.322 | M3 | 1.388E-02 | - | - | 1.357 | 4.239E-02 | 1.436 | 1.388E-02 | -1.322 | 3.249E-02 | - | - | - | - |
| hsa05132 | Salmonella infection | 1.336 | M5 | -1.375 | M4 | 1.541E-02 | - | - | - | - | -1.284 | 4.648E-02 | - | - | -1.375 | 1.541E-02 | 1.336 | 2.572E-02 |

### Online Supplemental Table S3: Overlapping significantly enriched GO pathways by GSEA showing directional change in the overall $S_1$ and $M_1$ through $M_5$ analyses.

| | | $M_1$ through $M_5$ | | | | | $S_1$ | | $M_1$ | | $M_2$ | | $M_3$ | | $M_4$ | | $M_5$ | |
|---|---|---|---|---|---|---|---|---|---|---|---|---|---|---|---|---|---|---|
| ID | Description | NES (Max) | NES (Max) Cluster | NES (Min) | NES (Min) Cluster | Min Padj | NES | Padj | NES | Padj | NES | Padj | NES | Padj | NES | Padj | NES | Padj |
| GO:0022613 | ribonucleoprotein complex biogenesis | 1.442 | M3 | -2.118 | M5 | 1.489E-08 | - | - | -2.085 | 1.489E-08 | - | - | 1.442 | 3.416E-02 | - | - | -2.118 | 2.531E-08 |
| GO:0042254 | ribosome biogenesis | 1.497 | M3 | -2.340 | M5 | 1.489E-08 | | | -2.259 | 1.489E-08 | | | 1.497 | 3.021E-02 | 1.467 | 4.544E-02 | -2.340 | 2.531E-08 |
| GO:0042581 | specific granule | 2.236 | M5 | -1.819 | M3 | 9.930E-08 | - | - | 2.099 | 7.389E-06 | - | - | -1.819 | 5.469E-03 | -1.690 | 2.701E-02 | 2.236 | 9.930E-08 |
| GO:0034470 | ncRNA processing | 1.527 | M4 | -1.860 | M1 | 4.502E-07 | - | - | -1.860 | 4.502E-07 | - | - | - | - | 1.527 | 1.477E-02 | -1.852 | 9.860E-07 |
| GO:0030667 | secretory granule membrane | 1.850 | M5 | -1.864 | M3 | 8.439E-05 | - | - | 1.708 | 2.382E-03 | - | - | -1.864 | 8.439E-05 | -1.819 | 5.282E-04 | 1.850 | 1.019E-04 |
| GO:0070820 | tertiary granule | 1.961 | M5 | -1.951 | M4 | 1.194E-04 | - | - | 1.587 | 3.203E-02 | - | - | -1.771 | 1.193E-02 | -1.951 | 5.422E-04 | 1.961 | 1.194E-04 |
| GO:0140053 | mitochondrial gene expression | 1.705 | M4 | -1.889 | M5 | 1.907E-04 | - | - | -1.468 | 4.746E-02 | - | - | - | - | 1.705 | 1.707E-02 | -1.889 | 1.907E-04 |
| GO:0005766 | primary lysosome | 1.853 | M1 | -1.793 | M4 | 8.065E-04 | - | - | 1.853 | 8.065E-04 | - | - | -1.665 | 3.035E-02 | -1.793 | 1.119E-02 | 1.563 | 3.681E-02 |
| GO:0042582 | azurophil granule | 1.853 | M1 | -1.793 | M4 | 8.065E-04 | - | - | 1.853 | 8.065E-04 | - | - | -1.665 | 3.035E-02 | -1.793 | 1.119E-02 | 1.563 | 3.681E-02 |
| GO:0009611 | response to wounding | 1.538 | M1 | -1.601 | M3 | 4.072E-03 | - | - | 1.538 | 9.994E-03 | - | - | -1.601 | 4.072E-03 | - | - | 1.415 | 3.019E-02 |
| GO:0006959 | humoral immune response | 1.685 | M1 | -1.591 | M3 | 6.205E-03 | - | - | 1.685 | 6.205E-03 | - | - | -1.591 | 3.657E-02 | - | - | - | - |
| GO:0034774 | secretory granule lumen | 1.514 | M1 | -1.625 | M4 | 9.485E-03 | - | - | 1.514 | 2.399E-02 | - | - | -1.589 | 9.485E-03 | -1.625 | 1.310E-02 | - | - |
| GO:0019731 | antibacterial humoral response | 2.000 | M2 | -2.074 | M3 | 9.485E-03 | - | - | - | - | 2.000 | 4.184E-02 | -2.074 | 9.485E-03 | - | - | - | - |
| GO:0002446 | neutrophil mediated immunity | 1.903 | M5 | -2.049 | M3 | 9.911E-03 | - | - | - | - | - | - | -2.049 | 1.839E-02 | -2.033 | 9.911E-03 | 1.903 | 1.457E-02 |
| GO:0042060 | wound healing | 1.556 | M1 | -1.512 | M3 | 1.035E-02 | - | - | 1.556 | 1.035E-02 | - | - | -1.512 | 2.743E-02 | - | - | 1.436 | 3.760E-02 |
| GO:0060205 | cytoplasmic vesicle lumen | 1.517 | M1 | -1.593 | M4 | 1.306E-02 | - | - | 1.517 | 1.404E-02 | - | - | -1.584 | 1.306E-02 | -1.593 | 2.589E-02 | - | - |
| GO:0031983 | vesicle lumen | 1.517 | M1 | -1.593 | M3 | 1.306E-02 | - | - | 1.517 | 1.404E-02 | - | - | -1.584 | 1.306E-02 | -1.593 | 2.589E-02 | - | - |
| GO:0019730 | antimicrobial humoral response | 1.777 | M1 | -1.916 | M3 | 1.385E-02 | - | - | 1.777 | 1.737E-02 | - | - | -1.916 | 1.385E-02 | - | - | - | - |
| GO:0010562 | positive regulation of phosphorus metabolic process | 1.362 | M5 | -1.421 | M4 | 3.176E-02 | - | - | - | - | - | - | - | - | -1.421 | 4.583E-02 | 1.362 | 3.176E-02 |
| GO:0045937 | positive regulation of phosphate metabolic process | 1.362 | M5 | -1.421 | M4 | 3.176E-02 | - | - | - | - | - | - | - | - | -1.421 | 4.583E-02 | 1.362 | 3.176E-02 |
| GO:0071496 | cellular response to external stimulus | 1.443 | M1 | -1.546 | M4 | 3.983E-02 | - | - | 1.443 | 3.983E-02 | - | - | - | - | -1.546 | 4.302E-02 | 1.422 | 4.010E-02 |
| GO:0101002 | ficolin-1-rich granule | 1.499 | M5 | -1.590 | M4 | 4.406E-02 | - | - | - | - | - | - | - | - | -1.590 | 4.406E-02 | 1.499 | 4.834E-02 |

### Online Supplemental Table S4: All transcripts showing differential expression in at least 1 comparison of overall $S_1$ and $M_1$ through $M_5$ analyses sorted by delta FC as a measure of inter-cluster differences (M1 through M5).

| | | | S1 | | | M1 through M5 | | | | | M1 | | | | M2 | | | | M3 | | | | M4 | | | | M5 | | | |
|---|---|---|---|---|---|---|---|---|---|---|---|---|---|---|---|---|---|---|---|---|---|---|---|---|---|---|---|---|---|---|
| ID | Gene_Symbol | Cytoband | FC | $P_{val}$ | $P_{adj}$ | Max $P_{adj}$ | Min $P_{adj}$ | ΔFC | Max FC | Min FC | FC | FC Rank | $P_{val}$ | $P_{adj}$ | FC | FC Rank | $P_{val}$ | $P_{adj}$ | FC | FC Rank | $P_{val}$ | $P_{adj}$ | FC | FC Rank | $P_{val}$ | $P_{adj}$ | FC | FC Rank | $P_{val}$ | $P_{adj}$ |
| ILMN_1708983 | DNAI7 | 12p12.1b | -0.19 | 3.21E-01 | 4.83E-01 | 9.98E-01 | 1.11E-12 | 3.00 | 2.39 | -0.61 | -0.25 | 385 | 2.16E-01 | 3.89E-01 | 2.39 | 3 | 1.47E-16 | 1.11E-12 | -0.61 | 436 | 3.24E-03 | 8.73E-03 | 0.00 | 301 | 9.96E-01 | 9.98E-01 | -0.61 | 398 | 3.25E-02 | 7.20E-02 |
| ILMN_2318568 | HCFC1R1 | 16p13.3d | -0.77 | 3.44E-04 | 3.69E-03 | 7.07E-01 | 5.31E-09 | 2.89 | 1.07 | -1.82 | -0.75 | 521 | 4.31E-04 | 4.05E-03 | 1.07 | 63 | 1.55E-04 | 1.33E-03 | -1.40 | 559 | 2.26E-10 | 5.31E-09 | -0.15 | 384 | 4.90E-01 | 7.07E-01 | -1.82 | 560 | 2.61E-09 | 6.50E-08 |
| ILMN_1723436 | PFKFB2 | 1q32.1h-q32.2a | 0.53 | 5.03E-03 | 2.56E-02 | 6.72E-01 | 1.23E-13 | 2.79 | 2.41 | -0.38 | 0.81 | 87 | 3.77E-05 | 6.08E-04 | -0.38 | 406 | 1.32E-01 | 2.48E-01 | 0.28 | 236 | 1.45E-01 | 2.16E-01 | 0.16 | 205 | 4.43E-01 | 6.72E-01 | 2.41 | 2 | 1.39E-16 | 1.23E-13 |
| ILMN_1881081 | H3C6 | 0 | 0.10 | 6.75E-01 | 7.91E-01 | 6.79E-01 | 2.33E-05 | 2.71 | 1.79 | -0.92 | 0.17 | 302 | 5.10E-01 | 6.79E-01 | -0.92 | 497 | 8.79E-03 | 3.12E-02 | 0.27 | 241 | 3.05E-01 | 3.98E-01 | -0.44 | 474 | 1.15E-01 | 3.11E-01 | 1.79 | 14 | 2.65E-06 | 2.33E-05 |
| ILMN_1723433 | FAM72B | 1p12a | -0.90 | 6.20E-03 | 2.97E-02 | 7.02E-01 | 4.97E-07 | 2.71 | -0.28 | -2.98 | -1.01 | 547 | 7.00E-03 | 3.43E-02 | -0.28 | 377 | 5.69E-01 | 6.99E-01 | -0.90 | 485 | 1.60E-02 | 3.44E-02 | -0.28 | 430 | 4.81E-01 | 7.02E-01 | -2.98 | 565 | 2.84E-08 | 4.97E-07 |
| ILMN_1681728 | ##_LOC643505 | 7q36.1b | -0.79 | 8.80E-05 | 1.45E-03 | 5.31E-01 | 3.84E-09 | 2.67 | 0.88 | -1.78 | -0.75 | 522 | 2.57E-04 | 2.70E-03 | 0.88 | 91 | 1.13E-03 | 6.26E-03 | -1.37 | 558 | 1.55E-10 | 3.84E-09 | -0.23 | 420 | 2.90E-01 | 5.31E-01 | -1.78 | 558 | 1.69E-09 | 4.56E-08 |
| ILMN_1707925 | ABHD12B | 14q22.1c | -0.58 | 4.34E-03 | 2.29E-02 | 4.73E-01 | 1.26E-07 | 2.63 | 1.39 | -1.24 | -0.46 | 452 | 2.68E-02 | 9.13E-02 | 1.39 | 19 | 6.91E-07 | 1.90E-05 | -1.24 | 542 | 9.13E-09 | 1.26E-07 | -0.26 | 427 | 2.35E-01 | 4.73E-01 | -0.94 | 423 | 1.29E-03 | 4.79E-03 |
| ILMN_1657619 | DNAJB14 | 4q23b | 0.33 | 2.24E-02 | 7.51E-02 | 9.33E-01 | 9.82E-10 | 2.56 | 1.13 | -1.42 | 0.42 | 208 | 3.88E-03 | 2.21E-02 | -1.42 | 554 | 1.12E-12 | 9.82E-10 | 0.71 | 140 | 1.50E-06 | 1.09E-05 | -0.03 | 320 | 8.54E-01 | 9.33E-01 | 1.13 | 137 | 4.43E-08 | 7.19E-07 |
| ILMN_1752668 | DAAM2 | 6p21.2a | -0.38 | 1.61E-01 | 3.01E-01 | 6.08E-01 | 1.31E-03 | 2.51 | 1.42 | -1.10 | -0.24 | 382 | 4.29E-01 | 6.08E-01 | 0.35 | 155 | 3.77E-01 | 5.29E-01 | -1.10 | 524 | 3.65E-04 | 1.31E-03 | -0.36 | 461 | 2.55E-01 | 4.95E-01 | 1.42 | 53 | 9.50E-04 | 3.70E-03 |
| ILMN_1651652 | RTN3 | 11q13.1a | 0.35 | 5.68E-02 | 1.46E-01 | 6.59E-01 | 4.82E-11 | 2.45 | 2.04 | -0.41 | 0.61 | 127 | 1.45E-03 | 1.05E-02 | -0.41 | 412 | 9.96E-02 | 2.00E-01 | 0.26 | 244 | 1.67E-01 | 2.43E-01 | -0.16 | 389 | 4.27E-01 | 6.59E-01 | 2.04 | 6 | 3.74E-13 | 4.82E-11 |
| ILMN_1723139 | GPD2 | 2q24.1c | 1.00 | 3.28E-09 | 9.51E-07 | 1.39E-01 | 5.11E-14 | 2.41 | 1.88 | -0.53 | 1.21 | 37 | 1.59E-13 | 3.43E-10 | -0.53 | 440 | 9.57E-03 | 3.33E-02 | 1.37 | 25 | 2.64E-16 | 5.11E-14 | 0.36 | 121 | 2.90E-02 | 1.39E-01 | 1.88 | 11 | 8.53E-16 | 4.94E-13 |
| ILMN_1700202 | TMEM135 | 11q14.2a-q14.2b | 0.51 | 1.79E-03 | 1.21E-02 | 5.67E-01 | 7.59E-10 | 2.36 | 1.12 | -1.25 | 0.47 | 187 | 2.98E-03 | 1.82E-02 | -1.25 | 544 | 4.83E-09 | 4.60E-07 | 1.10 | 55 | 2.39E-11 | 7.59E-10 | 0.16 | 202 | 3.28E-01 | 5.67E-01 | 1.12 | 147 | 6.11E-07 | 6.67E-06 |
| ILMN_1781060 | SYN2 | 3p25.2a | 0.14 | 5.33E-01 | 6.80E-01 | 7.39E-01 | 3.86E-05 | 2.35 | 1.58 | -0.77 | 0.48 | 176 | 4.57E-02 | 1.33E-01 | -0.77 | 483 | 1.55E-02 | 4.81E-02 | -0.11 | 326 | 6.63E-01 | 7.39E-01 | -0.35 | 457 | 1.69E-01 | 3.90E-01 | 1.58 | 34 | 4.73E-06 | 3.86E-05 |
| ILMN_1691143 | RAB18 | 10p12.1a | 0.39 | 9.08E-03 | 3.88E-02 | 8.82E-01 | 1.91E-07 | 2.34 | 1.15 | -1.19 | 0.51 | 161 | 4.21E-04 | 3.98E-03 | -1.19 | 537 | 1.56E-09 | 1.91E-07 | 0.78 | 127 | 1.52E-07 | 1.48E-06 | -0.05 | 332 | 7.55E-01 | 8.82E-01 | 1.15 | 130 | 2.92E-08 | 5.10E-07 |
| ILMN_1781077 | CEP120 | 5q23.2b | 0.36 | 4.61E-02 | 1.26E-01 | 5.92E-01 | 2.02E-06 | 2.29 | 0.87 | -1.43 | 0.31 | 251 | 1.04E-01 | 2.38E-01 | -1.43 | 556 | 3.48E-08 | 2.02E-06 | 0.83 | 112 | 2.00E-05 | 1.03E-04 | 0.19 | 187 | 3.53E-01 | 5.92E-01 | 0.87 | 248 | 1.22E-03 | 4.58E-03 |
| ILMN_1872419 | ##_HS.544637 | 0 | 0.45 | 2.89E-03 | 1.72E-02 | 7.07E-01 | 1.59E-07 | 2.26 | 1.10 | -1.15 | 0.43 | 203 | 4.80E-03 | 2.58E-02 | -1.15 | 534 | 2.79E-08 | 1.70E-06 | 0.90 | 102 | 1.19E-08 | 1.59E-07 | 0.11 | 228 | 4.90E-01 | 7.07E-01 | 1.10 | 151 | 4.75E-07 | 5.42E-06 |
| ILMN_1748123 | KLHL14 | 18q12.1e | 0.67 | 2.34E-03 | 1.48E-02 | 1.63E-01 | 1.32E-07 | 2.25 | 1.39 | -0.87 | 0.46 | 188 | 4.68E-02 | 1.36E-01 | -0.87 | 492 | 4.84E-03 | 1.98E-02 | 1.39 | 23 | 9.64E-09 | 1.32E-07 | 0.51 | 87 | 3.87E-02 | 1.63E-01 | 0.56 | 287 | 8.51E-02 | 1.57E-01 |
| ILMN_1741281 | RNF175 | 4q31.3d | -0.45 | 2.00E-02 | 6.90E-02 | 2.59E-01 | 1.11E-05 | 2.23 | 1.45 | -0.79 | -0.46 | 454 | 2.78E-02 | 9.37E-02 | 1.45 | 14 | 3.30E-07 | 1.11E-05 | -0.79 | 469 | 2.20E-04 | 8.46E-04 | -0.38 | 466 | 8.48E-02 | 2.59E-01 | -0.74 | 407 | 1.24E-02 | 3.22E-02 |
| ILMN_1774028 | MTFR1 | 8q13.1a | 0.59 | 5.31E-04 | 5.11E-03 | 5.59E-01 | 2.60E-09 | 2.21 | 1.19 | -1.01 | 0.56 | 145 | 9.15E-04 | 7.37E-03 | -1.01 | 505 | 7.57E-06 | 1.23E-04 | 1.14 | 47 | 9.74E-11 | 2.60E-09 | 0.18 | 193 | 3.18E-01 | 5.59E-01 | 1.19 | 115 | 8.10E-07 | 8.49E-06 |
| ILMN_2076658 | MRPL1 | 4q21.1c | 0.31 | 9.20E-02 | 2.05E-01 | 8.06E-01 | 7.06E-07 | 2.21 | 1.04 | -1.17 | 0.08 | 318 | 6.81E-01 | 8.06E-01 | -1.17 | 535 | 3.32E-06 | 6.53E-05 | 1.04 | 76 | 6.54E-08 | 7.06E-07 | 0.18 | 189 | 3.46E-01 | 5.85E-01 | 0.24 | 320 | 3.60E-01 | 4.83E-01 |
| ILMN_2270100 | CEP85L | 6q22.31a | 0.85 | 5.89E-05 | 1.08E-03 | 4.83E-01 | 2.67E-09 | 2.19 | 1.53 | -0.66 | 0.81 | 90 | 3.00E-04 | 3.05E-03 | -0.66 | 467 | 2.29E-02 | 6.49E-02 | 1.50 | 15 | 1.01E-10 | 2.67E-09 | 0.27 | 147 | 2.45E-01 | 4.83E-01 | 1.53 | 38 | 1.22E-06 | 1.20E-05 |
| ILMN_1808768 | ROCK1 | 18q11.1c-q11.1d | 0.18 | 2.28E-01 | 3.82E-01 | 7.90E-01 | 5.58E-07 | 2.19 | 0.93 | -1.26 | 0.36 | 235 | 2.37E-02 | 8.29E-02 | -1.26 | 546 | 6.07E-09 | 5.58E-07 | 0.32 | 221 | 4.34E-02 | 7.94E-02 | -0.09 | 354 | 6.07E-01 | 7.90E-01 | 0.93 | 242 | 3.79E-05 | 2.29E-04 |
| ILMN_1651316 | CD69 | 12p13.31a | 0.68 | 2.08E-04 | 2.63E-03 | 2.38E-01 | 7.23E-11 | 2.13 | 1.40 | -0.74 | 0.46 | 189 | 1.34E-02 | 5.51E-02 | -0.74 | 478 | 2.76E-03 | 1.27E-02 | 1.40 | 21 | 1.55E-12 | 7.23E-11 | 0.35 | 124 | 7.32E-02 | 2.38E-01 | 1.23 | 105 | 3.96E-06 | 3.30E-05 |
| ILMN_1669700 | KPNA5 | 6q22.2a | 0.46 | 5.60E-03 | 2.76E-02 | 3.87E-01 | 8.96E-10 | 2.13 | 1.16 | -0.96 | 0.29 | 263 | 8.14E-02 | 2.01E-01 | -0.96 | 498 | 1.50E-05 | 2.13E-04 | 1.16 | 43 | 2.88E-11 | 8.96E-10 | 0.31 | 132 | 7.38E-02 | 2.39E-01 | 0.26 | 317 | 2.69E-01 | 3.87E-01 |
| ILMN_2198270 | FAM209A | 20q13.31a | 0.32 | 9.74E-03 | 4.07E-02 | 7.73E-01 | 2.23E-10 | 2.11 | 1.30 | -0.82 | 0.40 | 214 | 1.47E-03 | 1.06E-02 | -0.82 | 489 | 1.31E-06 | 3.18E-05 | 0.40 | 202 | 1.45E-03 | 4.35E-03 | 0.07 | 252 | 5.82E-01 | 7.73E-01 | 1.30 | 76 | 2.48E-12 | 2.23E-10 |
| ILMN_2207291 | IFNG | 12q15a | 0.86 | 1.35E-04 | 1.94E-03 | 1.40E-01 | 9.63E-08 | 2.11 | 1.48 | -0.63 | 0.82 | 86 | 1.01E-03 | 7.90E-03 | -0.63 | 460 | 5.27E-02 | 1.23E-01 | 1.48 | 16 | 6.66E-09 | 9.63E-08 | 0.57 | 78 | 2.98E-02 | 1.40E-01 | 0.76 | 260 | 2.75E-02 | 6.28E-02 |
| ILMN_1804350 | ##_LOC644852 | 1q44e | -0.97 | 4.34E-04 | 4.39E-03 | 2.31E-01 | 1.19E-05 | 2.09 | 0.61 | -1.47 | -0.56 | 480 | 6.32E-02 | 1.69E-01 | 0.61 | 121 | 1.21E-01 | 2.31E-01 | -1.47 | 561 | 1.66E-06 | 1.19E-05 | -1.38 | 564 | 1.94E-05 | 2.12E-03 | -1.26 | 529 | 2.77E-03 | 9.13E-03 |
| ILMN_2249282 | TIPRL | 1q24.2a | 0.61 | 3.80E-04 | 3.97E-03 | 5.75E-01 | 1.26E-08 | 2.07 | 1.61 | -0.47 | 0.52 | 157 | 3.00E-03 | 1.83E-02 | -0.47 | 420 | 4.35E-02 | 1.07E-01 | 1.12 | 52 | 1.05E-09 | 1.99E-08 | 0.18 | 192 | 3.37E-01 | 5.75E-01 | 1.61 | 31 | 3.62E-10 | 1.26E-08 |
| ILMN_1774685 | IL24 | 1q32.1h | -0.32 | 2.67E-02 | 8.53E-02 | 9.46E-01 | 5.39E-08 | 2.06 | 1.32 | -0.74 | -0.44 | 442 | 4.34E-03 | 2.40E-02 | 1.32 | 24 | 2.90E-10 | 5.39E-08 | -0.62 | 438 | 6.48E-05 | 2.90E-04 | 0.02 | 288 | 8.78E-01 | 9.46E-01 | -0.74 | 408 | 6.04E-04 | 2.50E-03 |
| ILMN_1763603 | FAM177A1 | 14q13.2a | 0.12 | 3.80E-01 | 5.41E-01 | 6.45E-01 | 5.50E-07 | 2.06 | 1.17 | -0.89 | 0.21 | 294 | 1.52E-01 | 3.09E-01 | -0.89 | 495 | 6.49E-06 | 1.09E-04 | 0.25 | 246 | 8.92E-02 | 1.44E-01 | -0.13 | 371 | 4.09E-01 | 6.45E-01 | 1.17 | 120 | 3.21E-08 | 5.50E-07 |
| ILMN_1812281 | ARG1 | 6q23.2a | -0.39 | 1.60E-01 | 3.00E-01 | 8.56E-01 | 5.90E-03 | 2.06 | 1.07 | -0.99 | 0.10 | 314 | 7.60E-01 | 8.56E-01 | -0.67 | 469 | 1.07E-01 | 2.10E-01 | -0.99 | 497 | 2.06E-03 | 5.90E-03 | -0.72 | 524 | 3.12E-02 | 1.44E-01 | 1.07 | 170 | 1.57E-02 | 3.94E-02 |
| ILMN_2226271 | INSL3 | 19p13.11d | 0.35 | 5.76E-02 | 1.47E-01 | 9.72E-01 | 1.19E-05 | 2.05 | 1.42 | -0.64 | 0.48 | 178 | 1.96E-02 | 7.29E-02 | -0.64 | 462 | 1.81E-02 | 5.43E-02 | 0.34 | 217 | 1.01E-01 | 1.60E-01 | 0.02 | 292 | 9.37E-01 | 9.72E-01 | 1.42 | 52 | 1.20E-06 | 1.19E-05 |
| ILMN_2092041 | CYTIP | 2q24.1d | 0.10 | 4.56E-01 | 6.14E-01 | 4.31E-01 | 1.43E-08 | 2.04 | 0.83 | -1.21 | 0.18 | 300 | 1.70E-01 | 3.32E-01 | -1.21 | 540 | 5.33E-11 | 1.43E-08 | 0.29 | 229 | 2.85E-02 | 5.61E-02 | -0.18 | 395 | 2.02E-01 | 4.31E-01 | 0.83 | 253 | 1.13E-05 | 8.16E-05 |
| ILMN_1766195 | ZNF558 | 19p13.2d | 0.41 | 9.83E-04 | 7.93E-03 | 7.23E-02 | 1.06E-10 | 2.02 | 0.87 | -1.15 | 0.27 | 268 | 1.93E-02 | 7.23E-02 | -1.15 | 532 | 1.37E-12 | 9.82E-10 | 0.87 | 107 | 2.47E-12 | 1.06E-10 | 0.34 | 125 | 6.76E-03 | 5.66E-02 | 0.69 | 270 | 3.04E-05 | 1.91E-04 |
| ILMN_1740083 | ORC4 | 2q23.1a | 0.37 | 8.25E-03 | 3.62E-02 | 8.22E-01 | 4.11E-06 | 2.02 | 1.00 | -1.02 | 0.39 | 222 | 6.56E-03 | 3.26E-02 | -1.02 | 506 | 1.23E-07 | 5.34E-06 | 0.73 | 133 | 4.87E-07 | 4.11E-06 | 0.07 | 257 | 6.55E-01 | 8.22E-01 | 1.00 | 235 | 8.29E-07 | 8.66E-06 |
| ILMN_1662846 | GPR160 | 3q26.2b | 0.16 | 2.44E-01 | 4.00E-01 | 5.36E-01 | 9.39E-06 | 2.01 | 1.02 | -0.99 | 0.29 | 262 | 4.53E-02 | 1.33E-01 | -0.99 | 500 | 4.09E-07 | 1.31E-05 | 0.32 | 219 | 2.62E-02 | 5.22E-02 | -0.16 | 390 | 2.95E-01 | 5.36E-01 | 1.02 | 216 | 9.10E-07 | 9.39E-06 |
| ILMN_1837219 | ##_HS.163426 | 0 | -0.16 | 3.48E-01 | 5.12E-01 | 1.38E-01 | 1.43E-04 | 2.01 | 0.94 | -1.07 | -0.39 | 426 | 2.66E-02 | 9.07E-02 | 0.94 | 89 | 8.11E-05 | 7.99E-04 | -0.31 | 371 | 8.41E-02 | 1.38E-01 | 0.44 | 103 | 1.87E-02 | 1.06E-01 | -1.07 | 479 | 2.18E-05 | 1.43E-04 |
| ILMN_1777564 | MAD2L1 | 4q27a | 0.58 | 2.31E-04 | 2.82E-03 | 1.11E-01 | 6.36E-12 | 1.99 | 1.25 | -0.75 | 0.35 | 236 | 2.41E-02 | 8.42E-02 | -0.75 | 480 | 3.38E-04 | 2.43E-03 | 1.25 | 33 | 8.28E-14 | 6.36E-12 | 0.39 | 116 | 2.01E-02 | 1.11E-01 | 0.93 | 241 | 2.91E-05 | 1.84E-04 |
| ILMN_1727567 | OLIG2 | 21q22.11c | -0.20 | 4.65E-01 | 6.22E-01 | 9.24E-01 | 5.84E-03 | 1.98 | 0.57 | -1.41 | -0.09 | 347 | 7.73E-01 | 8.64E-01 | 0.57 | 124 | 1.74E-01 | 3.03E-01 | -0.38 | 385 | 2.38E-01 | 3.25E-01 | 0.07 | 255 | 8.36E-01 | 9.24E-01 | -1.41 | 541 | 1.63E-03 | 5.84E-03 |
| ILMN_2209299 | ZNF333 | 19p13.12b | 0.60 | 4.65E-05 | 9.09E-04 | 6.63E-01 | 1.02E-12 | 1.97 | 1.75 | -0.22 | 0.68 | 110 | 5.45E-06 | 1.33E-04 | -0.22 | 353 | 2.53E-01 | 3.99E-01 | 0.85 | 110 | 2.56E-08 | 3.12E-07 | 0.12 | 220 | 4.32E-01 | 6.63E-01 | 1.75 | 17 | 2.32E-15 | 1.02E-12 |
| ILMN_1773427 | KANK1 | 9p24.3b | 0.77 | 5.18E-04 | 5.00E-03 | 9.16E-02 | 4.48E-06 | 1.97 | 1.26 | -0.71 | 0.54 | 149 | 2.69E-02 | 9.16E-02 | -0.71 | 473 | 2.86E-02 | 7.73E-02 | 1.26 | 31 | 5.37E-07 | 4.48E-06 | 0.70 | 63 | 7.22E-03 | 5.91E-02 | 1.18 | 118 | 6.05E-04 | 2.51E-03 |
| ILMN_1698164 | ##_LOC641693 | 0 | 0.18 | 2.74E-01 | 4.34E-01 | 8.44E-01 | 1.65E-04 | 1.96 | 1.13 | -0.83 | 0.36 | 234 | 5.47E-02 | 1.52E-01 | -0.83 | 490 | 1.03E-03 | 5.81E-03 | 0.10 | 270 | 5.97E-01 | 6.81E-01 | -0.08 | 348 | 6.94E-01 | 8.44E-01 | 1.13 | 136 | 2.57E-05 | 1.65E-04 |
| ILMN_1674696 | OLAH | 10p13c | 0.35 | 5.88E-02 | 1.49E-01 | 8.59E-01 | 7.60E-09 | 1.96 | 1.91 | -0.05 | 0.55 | 148 | 8.11E-03 | 3.84E-02 | 0.48 | 134 | 7.58E-02 | 1.63E-01 | -0.05 | 314 | 8.10E-01 | 8.59E-01 | 0.09 | 237 | 6.63E-01 | 8.27E-01 | 1.91 | 9 | 2.00E-10 | 7.60E-09 |
| ILMN_1853344 | ##_HS.543470 | 0 | -0.28 | 6.88E-02 | 1.67E-01 | 7.86E-01 | 3.47E-06 | 1.96 | 1.15 | -0.80 | -0.29 | 399 | 6.99E-02 | 1.81E-01 | 1.15 | 46 | 6.74E-08 | 3.47E-06 | -0.80 | 473 | 7.03E-07 | 5.66E-06 | 0.09 | 241 | 6.00E-01 | 7.86E-01 | -0.55 | 393 | 1.28E-02 | 3.32E-02 |
| ILMN_1788095 | SPRYD3 | 12q13.13e | -0.78 | 1.43E-08 | 2.83E-06 | 1.60E-02 | 6.27E-18 | 1.95 | 0.63 | -1.32 | -0.71 | 515 | 1.99E-08 | 1.96E-06 | 0.63 | 119 | 1.18E-04 | 1.07E-03 | -1.29 | 547 | 2.92E-21 | 6.27E-18 | -0.44 | 475 | 7.24E-04 | 1.60E-02 | -1.32 | 532 | 4.44E-13 | 5.22E-11 |
| ILMN_1659843 | DNAJC3 | 13q32.1b | -0.07 | 6.84E-01 | 7.98E-01 | 6.84E-01 | 5.24E-04 | 1.94 | 0.90 | -1.04 | 0.12 | 312 | 5.16E-01 | 6.84E-01 | -1.04 | 511 | 4.74E-05 | 5.24E-04 | -0.12 | 329 | 5.42E-01 | 6.32E-01 | -0.32 | 442 | 1.09E-01 | 3.02E-01 | 0.90 | 245 | 8.96E-04 | 3.52E-03 |
| ILMN_2204545 | ST3GAL4 | 11q24.2c | -0.54 | 1.78E-03 | 1.21E-02 | 3.02E-01 | 7.78E-07 | 1.94 | 0.45 | -1.50 | -0.28 | 395 | 1.46E-01 | 3.02E-01 | -1.50 | 558 | 9.35E-09 | 7.78E-07 | -0.64 | 444 | 8.95E-04 | 2.86E-03 | -0.94 | 549 | 5.17E-06 | 9.72E-04 | 0.45 | 297 | 9.47E-02 | 1.71E-01 |
| ILMN_1694325 | NFIX | 19p13.13c-p13.13b | -0.07 | 7.72E-01 | 8.61E-01 | 9.66E-01 | 9.67E-04 | 1.94 | 0.39 | -1.55 | 0.29 | 264 | 3.35E-01 | 5.21E-01 | -1.55 | 561 | 1.04E-04 | 9.67E-04 | -0.02 | 308 | 9.52E-01 | 9.66E-01 | -0.60 | 503 | 5.77E-02 | 2.07E-01 | 0.39 | 304 | 3.49E-01 | 4.73E-01 |
| ILMN_2049303 | ECI1 | 16p13.3d | -0.13 | 2.55E-01 | 4.13E-01 | 3.65E-01 | 1.83E-09 | 1.93 | 1.13 | -0.79 | -0.21 | 374 | 7.41E-02 | 1.88E-01 | 1.13 | 50 | 3.03E-12 | 1.83E-09 | -0.38 | 386 | 1.46E-03 | 4.37E-03 | 0.18 | 195 | 1.51E-01 | 3.65E-01 | -0.79 | 412 | 2.12E-06 | 1.93E-05 |
| ILMN_1653553 | TEDC1 | 14q32.33d | -0.24 | 4.17E-02 | 1.17E-01 | 7.80E-01 | 2.70E-09 | 1.92 | 0.80 | -1.12 | -0.23 | 379 | 4.59E-02 | 1.34E-01 | 0.80 | 96 | 4.12E-07 | 1.31E-05 | -0.53 | 419 | 1.11E-05 | 6.26E-05 | 0.07 | 258 | 5.92E-01 | 7.80E-01 | -1.12 | 499 | 5.78E-11 | 2.70E-09 |
| ILMN_1796138 | BMX | Xp22.2 | -0.38 | 4.61E-02 | 1.26E-01 | 8.56E-01 | 6.58E-08 | 1.92 | 0.72 | -1.21 | 0.13 | 310 | 4.93E-01 | 6.65E-01 | -0.07 | 304 | 7.74E-01 | 8.56E-01 | -1.21 | 537 | 4.29E-09 | 6.58E-08 | -0.50 | 484 | 1.63E-02 | 9.68E-02 | 0.72 | 266 | 9.52E-03 | 2.59E-02 |
| ILMN_1651490 | CYP26A1 | 10q23.33a | -0.41 | 1.33E-02 | 5.12E-02 | 7.86E-01 | 2.63E-08 | 1.92 | 0.83 | -1.08 | -0.08 | 343 | 6.52E-01 | 7.86E-01 | 0.83 | 94 | 2.78E-04 | 2.09E-03 | -1.08 | 519 | 1.46E-09 | 2.63E-08 | -0.31 | 434 | 8.88E-02 | 2.65E-01 | -0.41 | 384 | 8.77E-02 | 1.61E-01 |
| ILMN_1791006 | AHI1 | 6q23.3a | -0.16 | 3.59E-01 | 5.21E-01 | 4.88E-01 | 8.33E-07 | 1.91 | 0.51 | -1.41 | -0.36 | 423 | 4.32E-02 | 1.28E-01 | -1.41 | 552 | 1.02E-08 | 8.33E-07 | 0.51 | 179 | 5.40E-03 | 1.36E-02 | -0.22 | 418 | 2.49E-01 | 4.88E-01 | -0.49 | 388 | 5.12E-02 | 1.04E-01 |
| ILMN_1807423 | IGF2BP3 | 7p15.3b | 0.66 | 1.24E-04 | 1.83E-03 | 8.24E-01 | 4.82E-06 | 1.91 | 1.27 | -0.64 | 0.94 | 70 | 2.00E-07 | 1.08E-05 | -0.64 | 461 | 6.58E-03 | 2.50E-02 | 0.91 | 100 | 5.83E-07 | 4.82E-06 | 0.08 | 246 | 6.58E-01 | 8.24E-01 | 1.27 | 85 | 4.95E-07 | 5.60E-06 |
| ILMN_1705297 | MYBPH | 1q32.1e | -0.56 | 1.23E-02 | 4.84E-02 | 5.89E-01 | 1.20E-05 | 1.89 | 0.70 | -1.20 | -0.41 | 429 | 9.42E-02 | 2.23E-01 | 0.70 | 108 | 2.97E-02 | 7.95E-02 | -1.20 | 536 | 1.67E-06 | 1.20E-05 | -0.37 | 463 | 1.51E-01 | 3.64E-01 | -0.24 | 367 | 4.71E-01 | 5.89E-01 |
| ILMN_1744968 | KCNAB1 | 3q25.31b | -0.33 | 7.13E-02 | 1.71E-01 | 9.37E-01 | 3.82E-04 | 1.89 | 1.16 | -0.73 | -0.32 | 407 | 1.20E-01 | 2.63E-01 | 1.16 | 45 | 3.16E-05 | 3.82E-04 | -0.63 | 443 | 2.63E-03 | 7.29E-03 | -0.04 | 327 | 8.60E-01 | 9.37E-01 | -0.73 | 406 | 1.23E-02 | 3.21E-02 |
| ILMN_2052891 | PKD2 | 4q22.1b | 0.62 | 7.12E-08 | 7.82E-06 | 7.26E-03 | 1.75E-14 | 1.89 | 1.13 | -0.76 | 0.64 | 121 | 3.42E-09 | 5.01E-07 | -0.76 | 481 | 9.70E-08 | 4.51E-06 | 0.94 | 96 | 6.96E-17 | 1.75E-14 | 0.42 | 108 | 1.87E-04 | 7.26E-03 | 1.13 | 141 | 2.58E-13 | 3.72E-11 |
| ILMN_1788203 | HEY1 | 8q21.13a | -0.13 | 6.26E-01 | 7.54E-01 | 9.94E-01 | 1.25E-02 | 1.88 | 1.24 | -0.64 | 0.00 | 337 | 9.89E-01 | 9.94E-01 | -0.64 | 464 | 1.14E-01 | 2.21E-01 | -0.30 | 366 | 3.34E-01 | 4.28E-01 | -0.40 | 469 | 2.15E-01 | 4.49E-01 | 1.24 | 97 | 4.02E-03 | 1.25E-02 |
| ILMN_1777286 | RPL13P12 | 17p11.2h | -0.03 | 8.50E-01 | 9.12E-01 | 6.60E-01 | 3.38E-06 | 1.88 | 0.62 | -1.26 | -0.18 | 364 | 3.02E-01 | 4.85E-01 | -1.26 | 545 | 6.52E-08 | 3.38E-06 | 0.62 | 160 | 3.63E-04 | 1.31E-03 | -0.14 | 379 | 4.28E-01 | 6.60E-01 | -0.77 | 411 | 1.47E-03 | 5.37E-03 |
| ILMN_1738116 | TMEM119 | 12q24.11a | 0.44 | 5.87E-02 | 1.49E-01 | 9.62E-01 | 1.35E-05 | 1.88 | 1.83 | -0.05 | 0.59 | 133 | 2.57E-02 | 8.87E-02 | -0.05 | 294 | 8.92E-01 | 9.35E-01 | 0.28 | 233 | 283E-01 | 3.75E-01 | -0.03 | 321 | 9.13E-01 | 9.62E-01 | 1.83 | 12 | 1.39E-06 | 1.35E-05 |
| ILMN_2327812 | IL5RA | 3p26.3a | -0.26 | 2.51E-01 | 4.08E-01 | 9.64E-01 | 1.01E-02 | 1.88 | 0.76 | -1.12 | -0.28 | 393 | 3.02E-01 | 4.85E-01 | 0.76 | 100 | 3.24E-02 | 8.48E-02 | -0.38 | 387 | 1.58E-01 | 2.32E-01 | 0.03 | 281 | 9.19E-01 | 9.64E-01 | -1.12 | 498 | 3.12E-03 | 1.01E-02 |
| ILMN_1696270 | PLAG1 | 8q12.1b | -0.14 | 4.43E-01 | 6.03E-01 | 9.69E-01 | 1.17E-06 | 1.88 | 0.40 | -1.47 | -0.41 | 430 | 3.29E-02 | 1.06E-01 | -1.47 | 557 | 1.66E-08 | 1.17E-06 | 0.40 | 201 | 3.54E-02 | 6.72E-02 | -0.02 | 315 | 9.31E-01 | 9.69E-01 | -0.25 | 369 | 3.40E-01 | 4.63E-01 |

### Online Supplemental Table S4 (Continued)

| | | | S1 | | | M1 through M5 | | | | | M1 | | | | M2 | | | | M3 | | | | M4 | | | | M5 | | | |
|---|---|---|---|---|---|---|---|---|---|---|---|---|---|---|---|---|---|---|---|---|---|---|---|---|---|---|---|---|---|---|
| ID | Gene_Symbol | Cytoband | FC | $P_{val}$ | $P_{adj}$ | Max $P_{adj}$ | Min $P_{adj}$ | ΔFC | Max FC | Min FC | FC | FC Rank | $P_{val}$ | $P_{adj}$ | FC | FC Rank | $P_{val}$ | $P_{adj}$ | FC | FC Rank | $P_{val}$ | $P_{adj}$ | FC | FC Rank | $P_{val}$ | $P_{adj}$ | FC | FC Rank | $P_{val}$ | $P_{adj}$ |
| ILMN_1801387 | YEATS4 | 12q15c | 0.26 | 3.25E-02 | 9.83E-02 | 6.11E-01 | 4.55E-10 | 1.87 | 0.80 | -1.07 | 0.10 | 313 | 3.78E-01 | 5.65E-01 | -1.07 | 518 | 5.00E-12 | 2.43E-09 | 0.80 | 121 | 1.31E-11 | 4.55E-10 | 0.11 | 231 | 3.73E-01 | 6.11E-01 | 0.49 | 293 | 1.84E-03 | 6.47E-03 |
| ILMN_1730176 | ITGAX | 16p11.2c | 0.27 | 1.14E-01 | 2.38E-01 | 9.00E-01 | 1.26E-08 | 1.87 | 1.62 | -0.25 | 0.50 | 167 | 4.83E-03 | 2.59E-02 | 0.53 | 128 | 2.49E-02 | 6.94E-02 | -0.25 | 357 | 1.60E-01 | 2.34E-01 | 0.05 | 268 | 7.88E-01 | 9.00E-01 | 1.62 | 29 | 3.63E-10 | 1.26E-08 |
| ILMN_2352633 | ARHGAP24 | 4q21.23b-q21.3a | 0.38 | 8.62E-03 | 3.74E-02 | 5.89E-01 | 1.27E-07 | 1.86 | 1.36 | -0.50 | 0.44 | 196 | 6.34E-03 | 3.18E-02 | -0.50 | 433 | 1.89E-02 | 5.63E-02 | 0.47 | 183 | 3.93E-03 | 1.03E-02 | 0.16 | 203 | 3.51E-01 | 5.89E-01 | 1.36 | 59 | 5.79E-09 | 1.27E-07 |
| ILMN_1696749 | LMNA | 1q22c | -0.35 | 6.52E-03 | 3.06E-02 | 4.43E-01 | 1.37E-07 | 1.86 | 0.73 | -1.14 | -0.28 | 394 | 4.03E-02 | 1.22E-01 | 0.73 | 106 | 5.32E-05 | 5.73E-04 | -0.61 | 435 | 1.13E-05 | 6.32E-05 | -0.18 | 394 | 2.11E-01 | 4.43E-01 | -1.14 | 502 | 6.37E-09 | 1.37E-07 |
| ILMN_2067607 | TMEM106B | 7p21.3a | 0.28 | 1.06E-01 | 2.26E-01 | 7.63E-01 | 6.25E-05 | 1.86 | 0.81 | -1.06 | 0.21 | 297 | 2.48E-01 | 4.27E-01 | -1.06 | 513 | 1.22E-05 | 1.79E-04 | 0.81 | 117 | 1.11E-05 | 6.25E-05 | 0.11 | 229 | 5.66E-01 | 7.63E-01 | 0.30 | 312 | 2.35E-01 | 3.50E-01 |
| ILMN_1810752 | C4BPA | 1q32.2a | -0.69 | 6.99E-02 | 1.69E-01 | 8.86E-01 | 3.06E-02 | 1.86 | 0.63 | -1.23 | -1.23 | 556 | 6.04E-03 | 3.06E-02 | -0.88 | 494 | 1.36E-01 | 2.53E-01 | -0.82 | 478 | 6.75E-02 | 1.15E-01 | -0.14 | 378 | 7.62E-01 | 8.86E-01 | 0.63 | 277 | 3.11E-01 | 4.33E-01 |
| ILMN_1863592 | ##_HS.336370 | 0 | 0.13 | 2.94E-01 | 4.55E-01 | 3.33E-01 | 2.31E-07 | 1.86 | 1.06 | -0.80 | 0.23 | 288 | 7.85E-02 | 1.96E-01 | -0.80 | 487 | 3.14E-06 | 6.26E-05 | 0.29 | 232 | 2.52E-02 | 5.06E-02 | -0.20 | 411 | 1.30E-01 | 3.33E-01 | 1.06 | 186 | 1.17E-08 | 2.31E-07 |
| ILMN_1696584 | ORM1 | 9q32d | -0.32 | 3.13E-01 | 4.75E-01 | 9.72E-01 | 7.32E-04 | 1.85 | 0.51 | -1.34 | 0.09 | 315 | 7.97E-01 | 8.79E-01 | -0.10 | 315 | 8.38E-01 | 9.01E-01 | -1.34 | 554 | 1.86E-04 | 7.32E-04 | 0.03 | 280 | 9.37E-01 | 9.72E-01 | 0.51 | 292 | 3.05E-01 | 4.27E-01 |
| ILMN_1654389 | ##_FCGR1A | 1q21.2a | 0.70 | 8.28E-04 | 7.00E-03 | 6.46E-01 | 6.61E-06 | 1.85 | 1.63 | -0.22 | 1.08 | 51 | 3.22E-06 | 8.63E-05 | -0.22 | 350 | 4.66E-01 | 6.13E-01 | 0.66 | 149 | 4.03E-03 | 1.06E-02 | 0.20 | 178 | 4.11E-01 | 6.46E-01 | 1.63 | 27 | 6.05E-07 | 6.61E-06 |
| ILMN_1705984 | HNMT | 2q22.1b | 0.64 | 1.11E-04 | 1.71E-03 | 5.29E-01 | 2.82E-07 | 1.85 | 1.34 | -0.51 | 0.71 | 108 | 4.71E-05 | 7.18E-04 | -0.51 | 434 | 2.36E-02 | 6.65E-02 | 0.99 | 90 | 2.27E-08 | 2.82E-07 | 0.19 | 182 | 2.89E-01 | 5.29E-01 | 1.34 | 66 | 6.59E-08 | 1.01E-06 |
| ILMN_2345908 | DDX11 | 12p11.21b | 0.26 | 3.14E-01 | 4.76E-01 | 8.75E-01 | 2.14E-02 | 1.84 | 0.75 | -1.09 | 0.08 | 317 | 7.90E-01 | 8.75E-01 | -1.09 | 521 | 5.37E-03 | 2.14E-02 | 0.75 | 130 | 1.17E-02 | 2.63E-02 | 0.24 | 166 | 4.36E-01 | 6.67E-01 | 0.18 | 324 | 6.55E-01 | 7.51E-01 |
| ILMN_1670572 | IDO2 | 8p11.21c | 0.32 | 9.12E-02 | 2.03E-01 | 6.50E-01 | 1.02E-04 | 1.83 | 1.34 | -0.49 | 0.37 | 232 | 8.96E-02 | 2.15E-01 | -0.49 | 431 | 8.65E-02 | 1.80E-01 | 0.31 | 223 | 1.50E-01 | 2.23E-01 | 0.19 | 186 | 4.15E-01 | 6.50E-01 | 1.34 | 63 | 1.47E-05 | 1.02E-04 |
| ILMN_1752755 | VWF | 12p13.31e | -0.54 | 1.75E-02 | 6.27E-02 | 6.41E-01 | 5.51E-05 | 1.83 | 0.70 | -1.14 | -0.18 | 366 | 4.66E-01 | 6.41E-01 | 0.70 | 109 | 3.62E-02 | 9.27E-02 | -1.14 | 528 | 9.63E-06 | 5.51E-05 | -0.56 | 493 | 3.68E-02 | 1.59E-01 | -0.59 | 397 | 9.33E-02 | 1.69E-01 |
| ILMN_1785615 | SUMO1P1 | 20q13.2c | 0.30 | 3.69E-02 | 1.08E-01 | 9.01E-01 | 8.63E-10 | 1.83 | 1.50 | -0.33 | 0.48 | 175 | 1.50E-03 | 1.07E-02 | -0.33 | 391 | 1.02E-01 | 2.03E-01 | 0.14 | 264 | 3.62E-01 | 4.57E-01 | 0.04 | 278 | 7.92E-01 | 9.01E-01 | 1.50 | 44 | 1.41E-11 | 8.63E-10 |
| ILMN_1659770 | KCNJ15 | 21q22.13b | 0.55 | 1.12E-03 | 8.65E-03 | 9.99E-01 | 3.99E-10 | 1.82 | 1.82 | 0.00 | 0.78 | 92 | 2.03E-05 | 3.76E-04 | 0.00 | 280 | 9.99E-01 | 9.99E-01 | 0.28 | 235 | 1.16E-01 | 1.80E-01 | 0.25 | 164 | 1.93E-01 | 4.20E-01 | 1.82 | 13 | 5.30E-12 | 3.99E-10 |
| ILMN_2100287 | CFAP58 | 10q25.1a | 0.62 | 1.43E-05 | 3.83E-04 | 4.97E-01 | 3.64E-11 | 1.82 | 1.64 | -0.19 | 0.85 | 80 | 4.85E-08 | 3.86E-06 | -0.19 | 343 | 3.46E-01 | 4.97E-01 | 0.42 | 197 | 5.47E-03 | 1.37E-02 | 0.40 | 113 | 1.21E-02 | 8.09E-02 | 1.64 | 26 | 2.49E-13 | 3.64E-11 |
| ILMN_1695413 | ZNF429 | 19p12d | 0.14 | 5.14E-01 | 6.64E-01 | 8.16E-01 | 1.65E-03 | 1.82 | 1.20 | -0.62 | 0.40 | 218 | 9.75E-02 | 2.28E-01 | -0.62 | 459 | 4.93E-02 | 1.17E-01 | 0.07 | 278 | 7.58E-01 | 8.16E-01 | -0.23 | 421 | 3.65E-01 | 6.03E-01 | 1.20 | 112 | 3.70E-04 | 1.65E-03 |
| ILMN_2225608 | FAM135A | 6q13a | 0.26 | 4.69E-02 | 1.27E-01 | 8.41E-01 | 1.47E-07 | 1.82 | 0.70 | -1.12 | 0.30 | 257 | 2.44E-02 | 8.49E-02 | -1.12 | 527 | 1.10E-09 | 1.47E-07 | 0.70 | 142 | 3.12E-07 | 2.76E-06 | 0.06 | 266 | 6.88E-01 | 8.41E-01 | 0.25 | 318 | 1.85E-01 | 2.91E-01 |
| ILMN_1797301 | INKA2 | 1p13.2d | -0.63 | 2.84E-03 | 1.70E-02 | 2.57E-01 | 1.73E-03 | 1.82 | 0.69 | -1.13 | -0.37 | 425 | 1.16E-01 | 2.57E-01 | -1.13 | 529 | 3.00E-04 | 2.22E-03 | -0.76 | 464 | 1.37E-03 | 4.13E-03 | -1.10 | 555 | 1.44E-05 | 1.73E-03 | 0.69 | 271 | 3.71E-02 | 8.03E-02 |
| ILMN_1778478 | CCDC28B | 1p35.1b | -0.47 | 9.28E-04 | 7.59E-03 | 8.95E-01 | 1.06E-09 | 1.82 | 0.33 | -1.49 | -0.57 | 483 | 2.09E-04 | 2.31E-03 | 0.33 | 158 | 9.44E-02 | 1.92E-01 | -0.62 | 440 | 4.96E-05 | 2.29E-04 | -0.04 | 330 | 7.78E-01 | 8.95E-01 | -1.49 | 546 | 1.88E-11 | 1.06E-09 |
| ILMN_1725244 | HAT1 | 2q31.1d | 0.22 | 1.44E-01 | 2.79E-01 | 9.64E-01 | 7.64E-06 | 1.81 | 0.73 | -1.09 | 0.15 | 306 | 3.18E-01 | 5.02E-01 | -1.09 | 520 | 1.94E-07 | 7.64E-06 | 0.73 | 135 | 4.17E-06 | 2.67E-05 | -0.02 | 314 | 9.19E-01 | 9.64E-01 | 0.53 | 290 | 1.38E-02 | 3.52E-02 |
| ILMN_1695157 | CA4 | 17q23.1a | -0.51 | 1.31E-03 | 9.71E-03 | 6.86E-01 | 1.34E-13 | 1.81 | 0.50 | -1.31 | -0.19 | 368 | 2.14E-01 | 3.88E-01 | 0.50 | 132 | 1.12E-02 | 3.76E-02 | -1.31 | 548 | 8.46E-16 | 1.34E-13 | -0.44 | 473 | 5.78E-03 | 5.17E-02 | 0.12 | 331 | 5.79E-01 | 6.86E-01 |
| ILMN_1684563 | SPIN4 | Xq11.1b | 0.64 | 1.43E-04 | 2.03E-03 | 1.53E-01 | 1.38E-07 | 1.81 | 1.07 | -0.74 | 0.68 | 111 | 1.82E-04 | 2.07E-03 | -0.74 | 479 | 1.75E-03 | 8.95E-03 | 1.07 | 67 | 1.01E-08 | 1.38E-07 | 0.40 | 114 | 3.45E-02 | 1.53E-01 | 0.93 | 243 | 2.52E-04 | 1.19E-03 |
| ILMN_1680192 | APOBEC3A | 22q13.1c | 0.98 | 7.84E-07 | 4.51E-05 | 3.92E-01 | 7.29E-09 | 1.81 | 1.48 | -0.33 | 0.98 | 64 | 8.83E-06 | 1.94E-04 | -0.33 | 392 | 2.47E-01 | 3.92E-01 | 1.42 | 20 | 3.25E-10 | 7.29E-09 | 0.63 | 72 | 5.95E-03 | 5.28E-02 | 1.48 | 45 | 1.65E-06 | 1.56E-05 |
| ILMN_1758093 | TMEM260 | 14q23.1a | 0.30 | 2.46E-02 | 8.03E-02 | 7.52E-01 | 1.63E-06 | 1.80 | 1.11 | -0.70 | 0.33 | 241 | 2.25E-02 | 8.01E-02 | -0.70 | 472 | 3.18E-04 | 2.32E-03 | 0.46 | 186 | 1.78E-03 | 5.19E-03 | 0.09 | 238 | 5.51E-01 | 7.52E-01 | 1.11 | 149 | 1.17E-07 | 1.63E-06 |
| ILMN_1651950 | TPST1 | 7q11.21e | -0.21 | 2.78E-01 | 4.39E-01 | 8.34E-01 | 1.53E-03 | 1.80 | 1.05 | -0.75 | 0.07 | 319 | 7.27E-01 | 8.34E-01 | 0.27 | 180 | 3.23E-01 | 4.74E-01 | -0.75 | 462 | 4.38E-04 | 1.53E-03 | -0.36 | 460 | 9.97E-02 | 2.86E-01 | 1.05 | 188 | 3.87E-04 | 1.71E-03 |
| ILMN_1655766 | ANKRD34B | 5q14.1e | 0.35 | 5.79E-02 | 1.48E-01 | 8.87E-01 | 6.71E-05 | 1.79 | 1.31 | -0.49 | 0.57 | 141 | 5.92E-03 | 3.02E-02 | -0.49 | 428 | 7.41E-02 | 1.60E-01 | 0.28 | 239 | 1.83E-01 | 2.62E-01 | 0.06 | 259 | 7.66E-01 | 8.87E-01 | 1.31 | 75 | 9.04E-06 | 6.71E-05 |
| ILMN_2388090 | MAPK14 | 6p21.31b | 0.00 | 9.88E-01 | 9.94E-01 | 4.39E-01 | 1.41E-07 | 1.79 | 1.06 | -0.73 | 0.21 | 295 | 9.73E-02 | 2.28E-01 | -0.73 | 477 | 1.75E-05 | 2.43E-04 | -0.12 | 330 | 3.44E-01 | 4.39E-01 | -0.32 | 441 | 1.58E-02 | 9.47E-02 | 1.06 | 178 | 6.57E-09 | 1.41E-07 |
| ILMN_1730032 | BOK | 2q37.3g | -0.41 | 5.32E-03 | 2.67E-02 | 4.04E-01 | 3.52E-10 | 1.79 | 0.76 | -1.04 | -0.21 | 373 | 1.50E-01 | 3.07E-01 | 0.76 | 101 | 9.00E-05 | 8.66E-04 | -1.04 | 511 | 9.76E-12 | 3.52E-10 | -0.20 | 410 | 1.80E-01 | 4.04E-01 | -0.54 | 392 | 8.23E-03 | 2.29E-02 |
| ILMN_1723048 | GJC2 | 1q42.13c | -0.95 | 1.90E-08 | 3.40E-06 | 5.71E-01 | 3.74E-14 | 1.79 | 0.18 | -1.61 | -0.76 | 525 | 7.65E-06 | 1.74E-04 | 0.18 | 211 | 4.20E-01 | 5.71E-01 | -1.49 | 562 | 1.79E-16 | 3.74E-14 | -0.70 | 518 | 9.64E-05 | 5.20E-03 | -1.61 | 555 | 3.91E-11 | 1.93E-09 |
| ILMN_1734867 | NR2C1 | 12q22d | 0.35 | 4.16E-03 | 2.23E-02 | 2.02E-01 | 4.99E-08 | 1.79 | 0.78 | -1.01 | 0.22 | 290 | 8.16E-02 | 2.02E-01 | -1.01 | 504 | 5.34E-09 | 5.00E-07 | 0.78 | 124 | 3.12E-09 | 4.99E-08 | 0.32 | 131 | 1.82E-02 | 1.04E-01 | 0.46 | 296 | 1.03E-02 | 2.78E-02 |
| ILMN_1785646 | PMP22 | 17p12a | -0.02 | 9.29E-01 | 9.61E-01 | 9.85E-01 | 3.10E-02 | 1.79 | 0.79 | -1.00 | 0.01 | 333 | 9.73E-01 | 9.85E-01 | 0.79 | 98 | 3.56E-02 | 9.15E-02 | -0.30 | 369 | 2.92E-01 | 3.84E-01 | 0.50 | 88 | 9.74E-02 | 2.81E-01 | -1.00 | 434 | 1.18E-02 | 3.10E-02 |
| ILMN_1735910 | VMO1 | 17p13.2c | 0.06 | 7.42E-01 | 8.40E-01 | 9.17E-01 | 5.34E-04 | 1.79 | 1.18 | -0.61 | 0.04 | 324 | 8.55E-01 | 9.17E-01 | 1.18 | 41 | 4.88E-05 | 5.34E-04 | -0.23 | 353 | 2.78E-01 | 3.70E-01 | 0.45 | 97 | 4.80E-02 | 1.86E-01 | -0.61 | 399 | 4.31E-02 | 9.08E-02 |
| ILMN_1766264 | PI16 | 6p21.2c | -0.29 | 1.44E-01 | 2.80E-01 | 4.21E-01 | 1.50E-03 | 1.78 | 0.64 | -1.14 | -0.52 | 468 | 2.11E-02 | 7.71E-02 | 0.64 | 118 | 3.08E-02 | 8.15E-02 | -0.53 | 422 | 1.82E-02 | 3.85E-02 | 0.31 | 135 | 1.93E-01 | 4.21E-01 | -1.14 | 504 | 3.29E-04 | 1.50E-03 |
| ILMN_1737089 | CAPN5 | 11q13.5c | -0.78 | 1.61E-06 | 7.79E-05 | 4.83E-01 | 2.29E-08 | 1.77 | 0.22 | -1.55 | -0.70 | 511 | 7.54E-05 | 1.06E-03 | 0.22 | 193 | 3.32E-01 | 4.83E-01 | -1.08 | 518 | 2.41E-09 | 4.02E-08 | -0.56 | 495 | 2.20E-03 | 2.92E-02 | -1.55 | 552 | 7.37E-10 | 2.29E-08 |
| ILMN_1715065 | ZNF580 | 19q13.42c | 0.22 | 1.43E-01 | 2.78E-01 | 9.70E-01 | 2.94E-06 | 1.77 | 1.22 | -0.55 | 0.05 | 322 | 7.56E-01 | 8.53E-01 | -0.55 | 444 | 1.14E-02 | 3.81E-02 | 0.53 | 172 | 1.48E-03 | 4.43E-03 | -0.01 | 312 | 9.34E-01 | 9.70E-01 | 1.22 | 106 | 2.35E-07 | 2.94E-06 |
| ILMN_1663390 | CDC20 | 1p34.2a | 0.32 | 1.76E-01 | 3.20E-01 | 4.80E-01 | 1.61E-03 | 1.77 | 1.33 | -0.44 | 0.28 | 266 | 2.96E-01 | 4.80E-01 | 1.33 | 23 | 2.00E-04 | 1.61E-03 | 0.28 | 237 | 2.98E-01 | 3.90E-01 | 0.37 | 119 | 1.96E-01 | 4.23E-01 | -0.44 | 385 | 2.43E-01 | 3.59E-01 |
| ILMN_1757129 | TMEM88 | 17p13.1d | -0.18 | 3.00E-01 | 4.61E-01 | 8.48E-01 | 2.16E-04 | 1.76 | 1.13 | -0.63 | 0.06 | 320 | 7.48E-01 | 8.48E-01 | -0.50 | 432 | 5.06E-02 | 1.20E-01 | -0.63 | 442 | 1.21E-03 | 3.72E-03 | -0.37 | 464 | 6.92E-02 | 2.31E-01 | 1.13 | 140 | 3.54E-05 | 2.16E-04 |
| ILMN_2350357 | ZNF254 | 19p12b | 0.07 | 6.47E-01 | 7.70E-01 | 8.73E-01 | 5.01E-05 | 1.76 | 0.73 | -1.03 | 0.04 | 323 | 7.88E-01 | 8.73E-01 | -1.03 | 508 | 2.34E-06 | 5.01E-05 | 0.31 | 226 | 5.93E-02 | 1.03E-01 | -0.11 | 362 | 5.27E-01 | 7.36E-01 | 0.73 | 263 | 1.34E-03 | 4.96E-03 |
| ILMN_2113938 | TOR1AIP2 | 1q25.2c | 0.35 | 6.91E-03 | 3.20E-02 | 7.02E-01 | 9.58E-08 | 1.75 | 1.15 | -0.60 | 0.47 | 184 | 5.59E-04 | 4.97E-03 | -0.60 | 455 | 7.71E-04 | 4.63E-03 | 0.43 | 195 | 1.62E-03 | 4.79E-03 | 0.10 | 234 | 4.81E-01 | 7.02E-01 | 1.15 | 131 | 4.14E-09 | 9.58E-08 |
| ILMN_1795499 | MAML3 | 4q31.1d-q31.1e | 0.15 | 3.64E-01 | 5.27E-01 | 9.99E-01 | 5.21E-04 | 1.75 | 1.04 | -0.71 | 0.38 | 225 | 4.27E-02 | 1.27E-01 | -0.71 | 474 | 4.49E-03 | 1.86E-02 | 0.00 | 298 | 9.99E-01 | 9.99E-01 | -0.05 | 336 | 7.85E-01 | 8.98E-01 | 1.04 | 197 | 9.73E-05 | 5.21E-04 |
| ILMN_1739751 | SLC26A11 | 17q25.3d-q25.3e | -0.39 | 8.87E-03 | 3.82E-02 | 5.12E-01 | 3.36E-10 | 1.73 | 0.19 | -1.55 | -0.46 | 451 | 2.81E-03 | 1.75E-02 | 0.18 | 208 | 3.60E-01 | 5.12E-01 | -0.68 | 453 | 1.23E-05 | 6.83E-05 | 0.19 | 184 | 2.38E-01 | 4.75E-01 | -1.55 | 551 | 4.29E-12 | 3.36E-10 |
| ILMN_1702970 | NLRP6 | 11p15.5d | 0.21 | 1.47E-01 | 2.83E-01 | 9.17E-01 | 9.54E-10 | 1.73 | 1.58 | -0.15 | 0.25 | 280 | 1.11E-01 | 2.50E-01 | -0.15 | 331 | 4.62E-01 | 6.09E-01 | 0.02 | 291 | 8.86E-01 | 9.17E-01 | 0.04 | 277 | 8.00E-01 | 9.05E-01 | 1.58 | 33 | 1.61E-11 | 9.54E-10 |
| ILMN_1880342 | YME1L1 | 0 | -0.48 | 1.39E-03 | 1.02E-02 | 6.68E-01 | 7.47E-10 | 1.73 | 0.13 | -1.60 | -0.51 | 462 | 1.55E-03 | 1.10E-02 | 0.13 | 232 | 5.32E-01 | 6.68E-01 | -0.58 | 429 | 3.99E-04 | 1.42E-03 | -0.13 | 374 | 4.27E-01 | 6.59E-01 | -1.60 | 554 | 1.15E-11 | 7.47E-10 |
| ILMN_1861376 | PLXDC2 | 0 | 0.55 | 6.99E-04 | 6.22E-03 | 6.95E-01 | 6.47E-08 | 1.73 | 1.45 | -0.28 | 0.77 | 94 | 7.52E-06 | 1.72E-04 | -0.28 | 378 | 2.06E-01 | 3.44E-01 | 0.78 | 125 | 5.72E-06 | 3.51E-05 | -0.13 | 370 | 4.72E-01 | 6.95E-01 | 1.45 | 47 | 2.59E-09 | 6.47E-08 |
| ILMN_1737195 | CENPK | 5q12.3b | 0.08 | 7.46E-01 | 8.43E-01 | 9.74E-01 | 1.35E-02 | 1.73 | 0.65 | -1.08 | 0.02 | 331 | 9.51E-01 | 9.74E-01 | -1.08 | 519 | 2.97E-03 | 1.35E-02 | 0.65 | 152 | 1.71E-02 | 3.64E-02 | -0.14 | 377 | 6.24E-01 | 8.00E-01 | 0.07 | 337 | 8.49E-01 | 8.98E-01 |
| ILMN_2375879 | VEGFA | 6p21.1c | -0.27 | 6.76E-02 | 1.65E-01 | 9.04E-01 | 1.91E-06 | 1.73 | 0.71 | -1.02 | 0.03 | 325 | 8.35E-01 | 9.04E-01 | 0.71 | 107 | 3.86E-04 | 2.69E-03 | -0.80 | 474 | 2.04E-07 | 1.91E-06 | -0.06 | 342 | 6.95E-01 | 8.45E-01 | -1.02 | 448 | 1.94E-06 | 1.79E-05 |
| ILMN_1727618 | NDUFAF6 | 8q22.1c | 0.63 | 2.06E-07 | 1.69E-05 | 1.40E-03 | 5.97E-15 | 1.73 | 1.07 | -0.65 | 0.50 | 168 | 2.17E-05 | 3.97E-04 | -0.65 | 466 | 2.71E-05 | 3.42E-04 | 1.07 | 63 | 1.74E-17 | 5.97E-15 | 0.55 | 81 | 1.06E-05 | 1.40E-03 | 0.72 | 264 | 1.31E-05 | 9.28E-05 |
| ILMN_1866504 | ##_HS.446301 | 0 | 0.34 | 5.14E-02 | 1.36E-01 | 9.37E-01 | 6.26E-05 | 1.72 | 1.24 | -0.48 | 0.31 | 253 | 1.18E-01 | 2.59E-01 | -0.48 | 427 | 6.03E-02 | 1.37E-01 | 0.64 | 153 | 1.11E-03 | 3.44E-03 | -0.04 | 325 | 8.61E-01 | 9.37E-01 | 1.24 | 100 | 8.33E-06 | 6.26E-05 |
| ILMN_1752117 | PIP4P2 | 8q21.3e | 0.22 | 5.59E-02 | 1.44E-01 | 5.25E-01 | 5.21E-11 | 1.72 | 1.23 | -0.49 | 0.42 | 210 | 3.41E-04 | 3.38E-03 | -0.49 | 430 | 1.32E-03 | 7.08E-03 | 0.20 | 253 | 7.54E-02 | 1.26E-01 | -0.13 | 373 | 2.83E-01 | 5.25E-01 | 1.23 | 102 | 4.36E-13 | 5.21E-11 |
| ILMN_1779043 | CACNG6 | 19q13.42a | -0.38 | 1.38E-01 | 2.71E-01 | 8.97E-01 | 2.06E-03 | 1.72 | 0.27 | -1.45 | -0.30 | 400 | 3.08E-01 | 4.91E-01 | 0.27 | 182 | 4.87E-01 | 6.30E-01 | -0.53 | 420 | 7.47E-02 | 1.25E-01 | -0.09 | 352 | 7.82E-01 | 8.97E-01 | -1.45 | 543 | 4.79E-04 | 2.06E-03 |
| ILMN_2402817 | ZBTB16 | 11q23.2a | -0.58 | 3.04E-03 | 1.78E-02 | 4.99E-01 | 1.30E-05 | 1.72 | 0.65 | -1.06 | -0.62 | 499 | 4.33E-03 | 2.39E-02 | 0.65 | 115 | 2.24E-02 | 6.38E-02 | -1.06 | 515 | 1.82E-06 | 1.30E-05 | -0.35 | 455 | 1.27E-01 | 3.28E-01 | -0.27 | 371 | 3.76E-01 | 4.99E-01 |
| ILMN_1789095 | BMPR2 | 2q33.1g | 0.59 | 2.59E-06 | 1.09E-04 | 2.36E-01 | 4.87E-13 | 1.72 | 1.07 | -0.65 | 0.60 | 131 | 9.08E-07 | 3.27E-05 | -0.65 | 465 | 5.19E-05 | 5.60E-04 | 1.01 | 87 | 3.88E-15 | 4.87E-13 | 0.23 | 169 | 7.20E-02 | 2.36E-01 | 1.07 | 172 | 6.87E-10 | 2.15E-08 |
| ILMN_1781672 | GAB1 | 4q31.21c | 0.09 | 5.05E-01 | 6.57E-01 | 9.58E-01 | 8.18E-06 | 1.71 | 1.02 | -0.69 | 0.40 | 217 | 5.87E-03 | 3.00E-02 | -0.69 | 470 | 3.63E-04 | 2.58E-03 | -0.01 | 304 | 9.40E-01 | 9.58E-01 | -0.30 | 432 | 4.78E-02 | 1.85E-01 | 1.02 | 214 | 7.77E-07 | 8.18E-06 |
| ILMN_1664047 | CACNA1E | 1q25.3b | 0.41 | 6.21E-02 | 1.55E-01 | 6.74E-01 | 1.11E-04 | 1.71 | 1.51 | -0.20 | 0.84 | 82 | 7.34E-04 | 6.17E-03 | 0.22 | 195 | 4.96E-01 | 6.38E-01 | -0.20 | 344 | 4.19E-01 | 5.14E-01 | 0.20 | 179 | 4.46E-01 | 6.74E-01 | 1.51 | 42 | 1.62E-05 | 1.11E-04 |
| ILMN_1863569 | RSF1 | 0 | -0.34 | 2.14E-02 | 7.26E-02 | 5.32E-01 | 1.16E-04 | 1.70 | 1.01 | -0.69 | -0.32 | 405 | 5.69E-02 | 1.56E-01 | 1.01 | 77 | 7.08E-06 | 1.16E-04 | -0.60 | 434 | 3.69E-04 | 1.32E-03 | -0.19 | 398 | 2.91E-01 | 5.32E-01 | -0.69 | 403 | 3.35E-03 | 1.07E-02 |
| ILMN_1735075 | PGBD2 | 1q44f | 0.32 | 1.21E-02 | 4.78E-02 | 1.54E-01 | 1.90E-07 | 1.70 | 0.64 | -1.06 | 0.29 | 261 | 2.39E-02 | 8.36E-02 | -1.06 | 516 | 1.54E-09 | 1.90E-07 | 0.60 | 162 | 4.31E-06 | 2.74E-05 | 0.29 | 142 | 3.48E-02 | 1.54E-01 | 0.64 | 276 | 4.43E-04 | 1.93E-03 |
| ILMN_1712718 | BPNT1 | 1q41d | 0.00 | 9.94E-01 | 9.96E-01 | 9.07E-01 | 1.80E-05 | 1.70 | 0.36 | -1.34 | -0.04 | 340 | 8.39E-01 | 9.07E-01 | -1.34 | 550 | 6.36E-07 | 1.80E-05 | 0.36 | 212 | 7.24E-02 | 1.21E-01 | -0.13 | 372 | 5.41E-01 | 7.46E-01 | 0.17 | 327 | 5.46E-01 | 6.56E-01 |
| ILMN_1798224 | KDM5C | Xp11.22a | 0.52 | 1.75E-03 | 1.20E-02 | 6.52E-01 | 3.77E-06 | 1.70 | 1.34 | -0.36 | 0.50 | 166 | 5.93E-03 | 3.02E-02 | -0.36 | 402 | 1.32E-01 | 2.47E-01 | 0.74 | 131 | 6.04E-05 | 2.72E-04 | 0.16 | 206 | 4.18E-01 | 6.52E-01 | 1.34 | 65 | 3.15E-07 | 3.77E-06 |
| ILMN_1829989 | ZEB1-AS1 | 0 | -0.01 | 9.32E-01 | 9.63E-01 | 8.70E-01 | 8.86E-10 | 1.70 | 0.69 | -1.00 | 0.03 | 326 | 7.82E-01 | 8.70E-01 | -1.00 | 501 | 8.87E-13 | 8.86E-10 | 0.08 | 277 | 4.45E-01 | 5.40E-01 | -0.12 | 363 | 2.78E-01 | 5.19E-01 | 0.69 | 268 | 1.40E-06 | 1.35E-05 |
| ILMN_1687958 | SLC25A22 | 11p15.5c | -0.27 | 2.66E-02 | 8.50E-02 | 7.52E-01 | 1.65E-07 | 1.69 | 0.68 | -1.01 | -0.26 | 387 | 3.47E-02 | 1.10E-01 | 0.68 | 111 | 2.73E-05 | 3.43E-04 | -0.58 | 430 | 2.72E-06 | 1.82E-05 | 0.08 | 250 | 5.51E-01 | 7.52E-01 | -1.01 | 440 | 7.96E-09 | 1.65E-07 |
| ILMN_2067852 | SLC30A1 | 1q32.3a | 0.54 | 2.71E-05 | 6.22E-04 | 7.37E-01 | 1.21E-08 | 1.69 | 1.21 | -0.48 | 0.73 | 103 | 8.74E-08 | 6.06E-06 | -0.48 | 425 | 6.53E-03 | 2.49E-02 | 0.72 | 136 | 1.40E-07 | 1.37E-06 | 0.09 | 240 | 5.29E-01 | 7.37E-01 | 1.21 | 109 | 3.45E-10 | 1.21E-08 |

### Online Supplemental Table S4 (Continued)

| ID | Gene_Symbol | Cytoband | S1 | | | M1 through M5 | | | | | M1 | | | | M2 | | | | M3 | | | | M4 | | | | M5 | | | |
|---|---|---|---|---|---|---|---|---|---|---|---|---|---|---|---|---|---|---|---|---|---|---|---|---|---|---|---|---|---|---|
| | | | FC | $P_{val}$ | $P_{adj}$ | Max $P_{adj}$ | Min $P_{adj}$ | ΔFC | Max FC | Min FC | FC | FC Rank | $P_{val}$ | $P_{adj}$ | FC | FC Rank | $P_{val}$ | $P_{adj}$ | FC | FC Rank | $P_{val}$ | $P_{adj}$ | FC | FC Rank | $P_{val}$ | $P_{adj}$ | FC | FC Rank | $P_{val}$ | $P_{adj}$ |
| ILMN_2408415 | RPL9 | 4p14b | 0.42 | 1.89E-02 | 6.64E-02 | 6.11E-01 | 2.11E-07 | 1.69 | 1.10 | -0.59 | 0.15 | 309 | 4.32E-01 | 6.11E-01 | -0.59 | 453 | 1.82E-02 | 5.46E-02 | 1.10 | 53 | 1.64E-08 | 2.11E-07 | 0.27 | 148 | 1.74E-01 | 3.96E-01 | 0.32 | 310 | 2.16E-01 | 3.29E-01 |
| ILMN_1756878 | SLC39A9 | 14q24.1e | 0.22 | 1.23E-01 | 2.51E-01 | 9.03E-01 | 8.78E-06 | 1.69 | 1.09 | -0.60 | 0.22 | 292 | 1.59E-01 | 3.19E-01 | -0.60 | 456 | 3.14E-03 | 1.41E-02 | 0.41 | 200 | 7.96E-03 | 1.90E-02 | -0.04 | 328 | 7.95E-01 | 9.03E-01 | 1.09 | 164 | 8.41E-07 | 8.78E-06 |
| ILMN_1789781 | PIM3 | 22q13.33a | 0.06 | 7.17E-01 | 8.22E-01 | 6.72E-01 | 3.48E-04 | 1.69 | 0.57 | -1.12 | 0.24 | 287 | 2.27E-01 | 4.02E-01 | -1.12 | 528 | 2.77E-05 | 3.48E-04 | 0.14 | 263 | 4.71E-01 | 5.66E-01 | -0.16 | 391 | 4.43E-01 | 6.72E-01 | 0.57 | 284 | 4.21E-02 | 8.90E-02 |
| ILMN_1666453 | STK3 | 8q22.2a | 0.33 | 2.25E-03 | 1.43E-02 | 7.59E-01 | 4.82E-11 | 1.69 | 1.09 | -0.59 | 0.50 | 170 | 1.63E-06 | 5.13E-05 | -0.59 | 454 | 1.36E-05 | 1.97E-04 | 0.46 | 187 | 1.09E-05 | 6.14E-05 | -0.06 | 343 | 5.62E-01 | 7.59E-01 | 1.09 | 155 | 3.84E-13 | 4.82E-11 |
| ILMN_1694127 | ##_LOC650761 | 0 | -1.26 | 3.37E-09 | 9.57E-07 | 3.46E-01 | 2.24E-08 | 1.68 | -0.38 | -2.06 | -1.53 | 561 | 1.84E-10 | 5.42E-08 | -0.38 | 405 | 2.08E-01 | 3.46E-01 | -1.35 | 556 | 1.36E-08 | 1.79E-07 | -0.69 | 517 | 4.81E-03 | 4.64E-02 | -2.06 | 563 | 7.19E-10 | 2.24E-08 |
| ILMN_1688423 | FCER1A | 1q23.2a | -0.46 | 2.37E-02 | 7.83E-02 | 9.65E-01 | 1.64E-05 | 1.68 | 0.03 | -1.65 | -0.54 | 474 | 2.60E-02 | 8.94E-02 | -0.44 | 417 | 1.67E-01 | 2.95E-01 | -0.40 | 391 | 102E-01 | 1.61E-01 | 0.03 | 287 | 9.21E-01 | 9.65E-01 | -1.65 | 556 | 1.75E-06 | 1.64E-05 |
| ILMN_1763433 | TRIM9 | 14q22.1c | -0.43 | 2.30E-02 | 7.66E-02 | 2.25E-01 | 8.27E-04 | 1.68 | 0.53 | -1.15 | -0.36 | 422 | 9.59E-02 | 2.25E-01 | -1.15 | 531 | 8.48E-05 | 8.27E-04 | -0.54 | 424 | 1.34E-02 | 2.95E-02 | -0.60 | 502 | 1.00E-02 | 7.16E-02 | 0.53 | 289 | 8.03E-02 | 1.49E-01 |
| ILMN_2171295 | CDK14 | 7q21.13c | 0.32 | 1.95E-02 | 6.79E-02 | 8.58E-01 | 1.18E-07 | 1.67 | 1.26 | -0.41 | 0.48 | 177 | 1.41E-03 | 1.02E-02 | -0.41 | 413 | 3.56E-02 | 9.16E-02 | 0.29 | 230 | 5.05E-02 | 9.03E-02 | 0.06 | 265 | 7.16E-01 | 8.58E-01 | 1.26 | 88 | 5.31E-09 | 1.18E-07 |
| ILMN_1655913 | NUCB2 | 11p15.1d | 0.11 | 3.88E-01 | 5.50E-01 | 8.28E-01 | 1.41E-07 | 1.67 | 0.64 | -1.03 | -0.05 | 341 | 6.85E-01 | 8.08E-01 | -1.03 | 509 | 1.02E-09 | 1.41E-07 | 0.64 | 154 | 3.54E-07 | 3.09E-06 | -0.06 | 338 | 6.65E-01 | 8.28E-01 | 0.29 | 313 | 8.64E-02 | 1.59E-01 |
| ILMN_1704139 | DHRSX | Xp22.33d-p22.33c,Yp11.31c-p11.31 | 0.36 | 7.67E-03 | 3.45E-02 | 9.73E-01 | 3.91E-06 | 1.67 | 1.10 | -0.57 | 0.54 | 150 | 3.58E-04 | 3.50E-03 | -0.57 | 450 | 3.83E-03 | 1.64E-02 | 0.44 | 191 | 3.39E-03 | 9.10E-03 | 0.01 | 296 | 9.41E-01 | 9.73E-01 | 1.10 | 152 | 3.30E-07 | 3.91E-06 |
| ILMN_1728742 | FAXDC2 | 5q33.2b | -0.59 | 1.58E-03 | 1.12E-02 | 4.93E-01 | 2.38E-07 | 1.67 | 0.51 | -1.16 | -0.20 | 371 | 3.09E-01 | 4.93E-01 | 0.51 | 130 | 5.19E-02 | 1.22E-01 | -1.16 | 531 | 1.88E-08 | 2.38E-07 | -0.66 | 513 | 1.71E-03 | 2.53E-02 | -1.02 | 449 | 2.90E-04 | 1.34E-03 |
| ILMN_1792323 | HDC | 15q21.2a | -0.75 | 3.90E-02 | 1.12E-01 | 9.95E-01 | 1.07E-02 | 1.67 | 0.10 | -1.56 | -0.79 | 530 | 6.45E-02 | 1.71E-01 | 0.01 | 276 | 9.89E-01 | 9.95E-01 | -1.24 | 541 | 4.11E-03 | 1.07E-02 | 0.10 | 232 | 8.18E-01 | 9.15E-01 | -1.56 | 553 | 9.17E-03 | 2.51E-02 |
| ILMN_1779979 | SLC37A3 | 7q34c | 0.12 | 3.56E-01 | 5.19E-01 | 8.87E-01 | 2.66E-07 | 1.66 | 1.22 | -0.43 | 0.24 | 283 | 9.99E-02 | 2.32E-01 | -0.43 | 416 | 2.72E-02 | 7.44E-02 | -0.08 | 321 | 5.86E-01 | 6.72E-01 | 0.05 | 271 | 7.64E-01 | 8.87E-01 | 1.22 | 107 | 1.39E-08 | 2.66E-07 |
| ILMN_2136455 | EOGT | 3p14.1b | 0.36 | 1.06E-02 | 4.33E-02 | 8.46E-01 | 3.19E-06 | 1.66 | 1.09 | -0.57 | 0.26 | 276 | 9.08E-02 | 2.17E-01 | -0.57 | 447 | 5.27E-03 | 2.11E-02 | 0.80 | 118 | 3.68E-07 | 3.19E-06 | 0.06 | 260 | 6.96E-01 | 8.46E-01 | 1.09 | 163 | 7.97E-07 | 8.36E-06 |
| ILMN_2339006 | VWA8 | 13q14.11b-q14.11c | 0.32 | 5.52E-03 | 2.73E-02 | 3.94E-01 | 8.65E-08 | 1.65 | 1.08 | -0.57 | 0.16 | 305 | 2.20E-01 | 3.94E-01 | -0.57 | 448 | 7.23E-04 | 4.39E-03 | 0.58 | 166 | 8.25E-06 | 4.83E-05 | 0.26 | 153 | 5.10E-02 | 1.92E-01 | 1.08 | 167 | 3.66E-09 | 8.65E-08 |
| ILMN_1728645 | ##_LOC649095 | 22q11.23c | -0.21 | 2.04E-01 | 3.54E-01 | 8.49E-01 | 9.45E-07 | 1.65 | 0.23 | -1.42 | -0.45 | 449 | 1.36E-02 | 5.57E-02 | -1.42 | 553 | 1.20E-08 | 9.45E-07 | 0.23 | 248 | 2.03E-01 | 2.85E-01 | -0.19 | 397 | 3.33E-01 | 5.72E-01 | -0.07 | 351 | 7.82E-01 | 8.49E-01 |
| ILMN_2137084 | LIN9 | 1q42.12c-q42.12d | 0.53 | 1.47E-04 | 2.07E-03 | 1.87E-01 | 8.22E-14 | 1.64 | 1.19 | -0.45 | 0.24 | 285 | 7.34E-02 | 1.87E-01 | -0.45 | 419 | 1.22E-02 | 4.01E-02 | 1.19 | 41 | 4.90E-16 | 8.22E-14 | 0.36 | 122 | 1.31E-02 | 8.50E-02 | 0.82 | 254 | 2.29E-05 | 1.50E-04 |
| ILMN_1654875 | CLC | 19q13.2b | -0.12 | 5.53E-01 | 6.98E-01 | 9.48E-01 | 4.73E-04 | 1.64 | 0.31 | -1.32 | -0.19 | 369 | 4.16E-01 | 5.97E-01 | 0.31 | 163 | 3.18E-01 | 4.70E-01 | -0.02 | 309 | 9.27E-01 | 9.48E-01 | 0.20 | 176 | 4.20E-01 | 6.53E-01 | -1.32 | 535 | 8.71E-05 | 4.73E-04 |
| ILMN_1689172 | ATXN1L | 16q22.3a | -0.57 | 2.05E-03 | 1.35E-02 | 4.57E-01 | 3.58E-05 | 1.64 | 0.49 | -1.15 | -0.22 | 378 | 2.75E-01 | 4.57E-01 | 0.49 | 133 | 6.70E-02 | 1.48E-01 | -0.93 | 489 | 5.86E-06 | 3.58E-05 | -0.74 | 528 | 5.34E-04 | 1.33E-02 | -1.15 | 508 | 5.92E-05 | 3.36E-04 |
| ILMN_1776653 | SCML1 | Xp22.13d | 0.17 | 3.10E-01 | 4.72E-01 | 9.47E-01 | 4.44E-04 | 1.63 | 0.59 | -1.04 | 0.02 | 329 | 9.04E-01 | 9.47E-01 | -1.04 | 510 | 3.83E-05 | 4.44E-04 | 0.59 | 165 | 1.94E-03 | 5.62E-03 | 0.22 | 171 | 2.64E-01 | 5.04E-01 | 0.04 | 343 | 8.90E-01 | 9.27E-01 |
| ILMN_2394210 | SLC26A8 | 6p21.31b | 0.54 | 1.46E-02 | 5.48E-02 | 9.62E-01 | 4.13E-05 | 1.63 | 1.62 | -0.02 | 0.96 | 68 | 1.35E-04 | 1.65E-03 | 0.15 | 223 | 6.48E-01 | 7.64E-01 | -0.02 | 306 | 9.47E-01 | 9.62E-01 | 0.27 | 149 | 3.10E-01 | 5.50E-01 | 1.62 | 30 | 5.12E-06 | 4.13E-05 |
| ILMN_2332368 | VPS13B | 8q22.2a-q22.2b | 0.40 | 2.20E-03 | 1.41E-02 | 7.82E-01 | 5.32E-10 | 1.63 | 1.41 | -0.23 | 0.44 | 202 | 1.86E-03 | 1.27E-02 | -0.23 | 355 | 2.17E-01 | 3.58E-01 | 0.56 | 168 | 9.01E-05 | 3.87E-04 | 0.08 | 248 | 5.95E-01 | 7.82E-01 | 1.41 | 54 | 7.49E-12 | 5.32E-10 |
| ILMN_1791568 | CEP57 | 11q21c | 0.51 | 3.46E-05 | 7.43E-04 | 1.68E-01 | 8.54E-15 | 1.63 | 1.06 | -0.57 | 0.43 | 205 | 2.31E-04 | 2.49E-03 | -0.57 | 449 | 1.94E-04 | 1.58E-03 | 1.06 | 69 | 2.72E-17 | 8.54E-15 | 0.25 | 160 | 4.03E-02 | 1.68E-01 | 0.58 | 282 | 3.58E-04 | 1.61E-03 |
| ILMN_1810962 | PTPRK | 6q22.33b-q22.33c | 0.53 | 6.02E-03 | 2.91E-02 | 5.06E-01 | 4.61E-06 | 1.63 | 1.08 | -0.54 | 0.26 | 275 | 2.14E-01 | 3.88E-01 | -0.54 | 443 | 5.07E-02 | 1.20E-01 | 1.08 | 61 | 5.54E-07 | 4.61E-06 | 0.52 | 86 | 1.93E-02 | 1.08E-01 | 0.26 | 316 | 3.82E-01 | 5.06E-01 |
| ILMN_2132599 | ANKRD22 | 10q23.31b | 1.09 | 5.79E-07 | 3.60E-05 | 5.55E-01 | 1.75E-07 | 1.63 | 1.89 | 0.26 | 1.53 | 18 | 8.81E-10 | 1.75E-07 | 0.26 | 184 | 4.04E-01 | 5.55E-01 | 1.09 | 60 | 8.59E-06 | 4.99E-05 | 0.45 | 99 | 7.40E-02 | 2.39E-01 | 1.89 | 10 | 4.49E-08 | 7.24E-07 |
| ILMN_1796335 | LPCAT2 | 16q12.2c | 0.30 | 2.34E-01 | 3.89E-01 | 6.95E-01 | 1.01E-02 | 1.62 | 1.18 | -0.45 | 0.84 | 81 | 3.63E-03 | 2.11E-02 | 1.18 | 40 | 2.04E-03 | 1.01E-02 | -0.45 | 404 | 1.20E-01 | 1.85E-01 | 0.22 | 172 | 4.71E-01 | 6.95E-01 | 0.36 | 307 | 3.74E-01 | 4.97E-01 |
| ILMN_1887305 | ##_HS.549784 | 0 | -0.09 | 5.86E-01 | 7.24E-01 | 4.55E-01 | 3.91E-05 | 1.62 | 1.13 | -0.49 | 0.19 | 299 | 2.74E-01 | 4.55E-01 | -0.41 | 411 | 7.48E-02 | 1.61E-01 | -0.33 | 376 | 6.12E-02 | 1.06E-01 | -0.49 | 479 | 8.34E-03 | 6.41E-02 | 1.13 | 139 | 4.82E-06 | 3.91E-05 |
| ILMN_2223805 | CEP41 | 7q32.2b | -0.39 | 3.09E-03 | 1.80E-02 | 9.89E-01 | 1.83E-12 | 1.62 | 0.00 | -1.62 | -0.58 | 485 | 5.06E-05 | 7.61E-04 | -1.62 | 563 | 4.27E-16 | 1.83E-12 | 0.00 | 302 | 9.84E-01 | 9.89E-01 | -0.31 | 433 | 3.99E-02 | 1.67E-01 | -0.52 | 390 | 8.38E-03 | 2.32E-02 |
| ILMN_2371055 | EFNA1 | 1q22a | -0.41 | 3.78E-02 | 1.09E-01 | 7.96E-01 | 4.31E-05 | 1.62 | 0.19 | -1.42 | -0.74 | 520 | 9.70E-04 | 7.69E-03 | -1.42 | 555 | 1.93E-06 | 4.31E-05 | -0.17 | 342 | 4.33E-01 | 5.28E-01 | -0.12 | 365 | 6.16E-01 | 7.96E-01 | 0.19 | 323 | 5.35E-01 | 6.47E-01 |
| ILMN_1661466 | SPINK8 | 0 | 0.15 | 2.65E-01 | 4.25E-01 | 9.91E-01 | 4.28E-08 | 1.61 | 1.32 | -0.29 | 0.30 | 254 | 4.29E-02 | 1.28E-01 | -0.29 | 383 | 1.36E-01 | 2.53E-01 | -0.07 | 316 | 654E-01 | 7.30E-01 | 0.00 | 302 | 9.79E-01 | 9.91E-01 | 1.32 | 70 | 1.57E-09 | 4.28E-08 |
| ILMN_1767960 | NSUN7 | 4p14a | 0.19 | 1.63E-01 | 3.04E-01 | 9.86E-01 | 1.90E-10 | 1.61 | 1.46 | -0.15 | 0.34 | 239 | 1.48E-02 | 5.90E-02 | -0.07 | 300 | 7.21E-01 | 8.18E-01 | 0.00 | 303 | 979E-01 | 9.86E-01 | -0.15 | 383 | 3.06E-01 | 5.46E-01 | 1.46 | 46 | 2.06E-12 | 1.90E-10 |
| ILMN_1695853 | CLK4 | 5q35.3c | 0.38 | 1.94E-04 | 2.51E-03 | 3.22E-01 | 2.86E-11 | 1.61 | 1.09 | -0.52 | 0.37 | 230 | 2.19E-04 | 2.39E-03 | -0.52 | 435 | 9.29E-05 | 8.90E-04 | 0.63 | 159 | 1.71E-09 | 3.04E-08 | 0.16 | 201 | 1.23E-01 | 3.22E-01 | 1.09 | 159 | 1.81E-13 | 2.86E-11 |
| ILMN_1802053 | ZNF91 | 19p12b | 0.07 | 5.72E-01 | 7.13E-01 | 7.47E-01 | 3.44E-07 | 1.61 | 0.55 | -1.05 | -0.08 | 345 | 5.45E-01 | 7.07E-01 | -1.05 | 512 | 3.33E-09 | 3.44E-07 | 0.55 | 169 | 3.16E-05 | 1.54E-04 | -0.08 | 350 | 5.42E-01 | 7.47E-01 | 0.17 | 326 | 3.46E-01 | 4.69E-01 |
| ILMN_1711408 | ANXA4 | 2p14a | 0.63 | 8.51E-06 | 2.66E-04 | 4.12E-01 | 4.73E-11 | 1.60 | 1.08 | -0.53 | 0.77 | 93 | 1.41E-07 | 8.54E-06 | -0.53 | 438 | 5.08E-03 | 2.05E-02 | 1.08 | 62 | 9.26E-13 | 4.73E-11 | 0.20 | 177 | 1.86E-01 | 4.12E-01 | 0.67 | 272 | 8.56E-04 | 3.39E-03 |
| ILMN_1713764 | ##_LOC440928 | 0 | 0.38 | 2.11E-03 | 1.37E-02 | 7.63E-01 | 3.39E-09 | 1.60 | 1.25 | -0.36 | 0.50 | 165 | 1.43E-04 | 1.73E-03 | -0.36 | 399 | 3.85E-02 | 9.72E-02 | 0.45 | 190 | 7.67E-04 | 2.51E-03 | 0.08 | 247 | 5.68E-01 | 7.63E-01 | 1.25 | 95 | 7.71E-11 | 3.39E-09 |
| ILMN_1785170 | ARMCX2 | Xq22.1d | 0.07 | 6.10E-01 | 7.43E-01 | 7.01E-01 | 1.02E-05 | 1.60 | 0.54 | -1.06 | -0.22 | 376 | 1.56E-01 | 3.15E-01 | -1.06 | 514 | 2.90E-07 | 1.02E-05 | 0.54 | 170 | 4.45E-04 | 1.55E-03 | 0.11 | 226 | 4.80E-01 | 7.01E-01 | 0.29 | 314 | 1.78E-01 | 2.82E-01 |
| ILMN_1663640 | MAOA | Xp11.3c | 0.27 | 2.33E-01 | 3.88E-01 | 9.87E-01 | 2.42E-04 | 1.60 | 1.56 | -0.04 | 0.45 | 193 | 9.27E-02 | 2.20E-01 | 0.22 | 191 | 5.27E-01 | 6.64E-01 | -0.04 | 313 | 8.88E-01 | 9.18E-01 | -0.01 | 305 | 9.69E-01 | 9.87E-01 | 1.56 | 36 | 4.03E-05 | 2.42E-04 |
| ILMN_1766346 | ZNF512B | 20q13.33e | -0.21 | 1.41E-01 | 2.75E-01 | 2.57E-01 | 1.53E-05 | 1.60 | 0.54 | -1.06 | -0.44 | 441 | 4.95E-03 | 2.64E-02 | 0.54 | 127 | 8.00E-03 | 2.91E-02 | -0.30 | 368 | 5.35E-02 | 9.48E-02 | 0.28 | 143 | 8.37E-02 | 2.57E-01 | -1.06 | 470 | 1.61E-06 | 1.53E-05 |
| ILMN_1651354 | SPP1 | 4q22.1b | -0.69 | 1.51E-02 | 5.62E-02 | 7.53E-01 | 2.88E-03 | 1.60 | 0.20 | -1.40 | -0.31 | 402 | 3.47E-01 | 5.34E-01 | -1.40 | 551 | 1.31E-03 | 7.02E-03 | -1.10 | 525 | 9.01E-04 | 2.88E-03 | -0.70 | 520 | 4.23E-02 | 1.72E-01 | 0.20 | 322 | 6.57E-01 | 7.53E-01 |
| ILMN_1653612 | KCTD9 | 8p21.2c | 0.43 | 7.20E-04 | 6.35E-03 | 8.15E-01 | 3.10E-09 | 1.60 | 1.01 | -0.58 | 0.39 | 220 | 2.12E-03 | 1.41E-02 | -0.58 | 452 | 5.19E-04 | 3.40E-03 | 0.85 | 108 | 1.20E-10 | 3.10E-09 | 0.06 | 263 | 6.45E-01 | 8.15E-01 | 1.01 | 221 | 2.91E-08 | 5.09E-07 |
| ILMN_1689378 | NOCT | 4q31.1b | 0.09 | 5.68E-01 | 7.09E-01 | 9.28E-01 | 6.97E-08 | 1.59 | 1.50 | -0.09 | 0.08 | 316 | 6.50E-01 | 7.85E-01 | 1.50 | 12 | 4.21E-10 | 6.97E-08 | -0.09 | 322 | 6.17E-01 | 6.99E-01 | 0.09 | 244 | 6.41E-01 | 8.12E-01 | 0.03 | 345 | 8.91E-01 | 9.28E-01 |
| ILMN_1792931 | LYSMD1 | 1q21.2d | -0.07 | 5.41E-01 | 6.87E-01 | 9.21E-01 | 5.47E-12 | 1.59 | 1.35 | -0.24 | -0.13 | 357 | 2.61E-01 | 4.42E-01 | 1.35 | 22 | 1.82E-15 | 5.47E-12 | -0.22 | 349 | 6.50E-02 | 1.11E-01 | -0.03 | 318 | 8.31E-01 | 9.21E-01 | -0.24 | 366 | 1.56E-01 | 2.54E-01 |
| ILMN_1742872 | UBA2 | 19q13.11c | -0.30 | 5.60E-02 | 1.44E-01 | 9.89E-01 | 2.74E-07 | 1.58 | 0.16 | -1.42 | -0.34 | 414 | 5.12E-02 | 1.45E-01 | 0.16 | 219 | 4.79E-01 | 6.23E-01 | -0.30 | 367 | 8.58E-02 | 1.40E-01 | -0.01 | 303 | 9.73E-01 | 9.89E-01 | -1.42 | 542 | 1.44E-08 | 2.74E-07 |
| ILMN_1682081 | RNF19B | 1p35.1b | 0.07 | 5.78E-01 | 7.18E-01 | 3.12E-01 | 1.27E-06 | 1.58 | 0.56 | -1.02 | 0.30 | 259 | 2.48E-02 | 8.60E-02 | -1.02 | 507 | 1.85E-08 | 1.27E-06 | 0.16 | 260 | 2.27E-01 | 3.12E-01 | -0.32 | 439 | 2.22E-02 | 1.17E-01 | 0.56 | 286 | 2.64E-03 | 8.75E-03 |
| ILMN_1753648 | TRIM51 | 11q11c | -0.49 | 5.50E-02 | 1.42E-01 | 9.70E-01 | 5.23E-03 | 1.58 | 0.23 | -1.35 | -0.46 | 453 | 1.24E-01 | 2.69E-01 | 0.23 | 188 | 5.64E-01 | 6.95E-01 | -0.81 | 477 | 7.37E-03 | 1.77E-02 | 0.03 | 285 | 9.34E-01 | 9.70E-01 | -1.35 | 539 | 1.43E-03 | 5.23E-03 |
| ILMN_1690783 | TREML1 | 6p21.1g | -0.74 | 5.92E-04 | 5.52E-03 | 4.84E-01 | 2.24E-06 | 1.57 | 0.30 | -1.27 | -0.33 | 410 | 1.61E-01 | 3.21E-01 | 0.30 | 165 | 3.33E-01 | 4.84E-01 | -1.27 | 543 | 2.44E-07 | 2.24E-06 | -0.84 | 539 | 9.41E-04 | 1.82E-02 | -0.97 | 426 | 3.89E-03 | 1.21E-02 |
| ILMN_1752226 | P2RY11 | 19p13.2c | -0.44 | 2.95E-04 | 3.31E-03 | 9.75E-01 | 7.53E-10 | 1.57 | 0.36 | -1.21 | -0.62 | 498 | 6.09E-07 | 2.40E-05 | 0.36 | 153 | 2.54E-02 | 7.04E-02 | -0.61 | 437 | 1.11E-06 | 8.44E-06 | 0.01 | 298 | 9.45E-01 | 9.75E-01 | -1.21 | 520 | 1.17E-11 | 7.53E-10 |
| ILMN_1684982 | PDK4 | 7q21.3b | 0.79 | 8.39E-05 | 1.40E-03 | 7.22E-01 | 1.21E-04 | 1.56 | 1.40 | -0.16 | 0.75 | 99 | 1.27E-03 | 9.45E-03 | -0.16 | 336 | 5.95E-01 | 7.22E-01 | 0.98 | 92 | 2.78E-05 | 1.38E-04 | 0.76 | 54 | 1.98E-03 | 2.75E-02 | 1.40 | 55 | 1.79E-05 | 1.21E-04 |
| ILMN_2228044 | TBC1D23 | 3q12.1c-q12.2a | 0.43 | 4.37E-04 | 4.41E-03 | 9.69E-01 | 1.19E-08 | 1.56 | 1.02 | -0.54 | 0.51 | 163 | 3.61E-05 | 5.92E-04 | -0.54 | 442 | 8.05E-04 | 4.79E-03 | 0.78 | 123 | 5.65E-10 | 1.19E-08 | 0.01 | 297 | 9.32E-01 | 9.69E-01 | 1.02 | 215 | 5.83E-09 | 1.27E-07 |
| ILMN_1798458 | KAZN | 1p36.21b-p36.21a | -0.03 | 9.01E-01 | 9.45E-01 | 9.26E-01 | 6.43E-03 | 1.56 | 1.11 | -0.45 | 0.33 | 242 | 1.92E-01 | 3.61E-01 | -0.05 | 296 | 8.78E-01 | 9.26E-01 | -0.45 | 403 | 7.90E-02 | 1.31E-01 | -0.41 | 470 | 1.22E-01 | 3.22E-01 | 1.11 | 148 | 1.83E-03 | 6.43E-03 |
| ILMN_1678037 | HIRIP3 | 16p11.2d | -0.32 | 4.97E-03 | 2.53E-02 | 5.12E-01 | 1.36E-08 | 1.55 | 0.51 | -1.05 | -0.47 | 458 | 5.92E-05 | 8.71E-04 | 0.51 | 131 | 9.27E-04 | 5.38E-03 | -0.52 | 418 | 1.04E-05 | 5.88E-05 | 0.13 | 218 | 2.72E-01 | 5.12E-01 | -1.05 | 464 | 3.97E-10 | 1.36E-08 |
| ILMN_1852022 | PLIN4 | 0 | 0.05 | 7.37E-01 | 8.37E-01 | 9.13E-01 | 3.09E-06 | 1.55 | 1.07 | -0.48 | 0.29 | 260 | 4.44E-02 | 1.31E-01 | -0.05 | 293 | 8.08E-01 | 8.80E-01 | -0.48 | 409 | 1.07E-03 | 3.34E-03 | -0.04 | 326 | 8.14E-01 | 9.13E-01 | 1.07 | 171 | 2.50E-07 | 3.09E-06 |
| ILMN_1731785 | ORM2 | 9q32d | 0.13 | 5.95E-01 | 7.31E-01 | 8.22E-01 | 2.09E-02 | 1.55 | 1.07 | -0.48 | 0.48 | 179 | 9.21E-02 | 2.19E-01 | 0.13 | 234 | 7.28E-01 | 8.22E-01 | -0.48 | 410 | 9.19E-02 | 1.48E-01 | 0.17 | 200 | 5.76E-01 | 7.69E-01 | 1.07 | 177 | 7.42E-03 | 2.09E-02 |
| ILMN_1706935 | CCDC136 | 7q32.1a | -0.21 | 9.18E-02 | 2.04E-01 | 5.41E-01 | 3.34E-07 | 1.54 | 0.47 | -1.08 | -0.34 | 417 | 9.51E-03 | 4.30E-02 | 0.47 | 137 | 7.59E-03 | 2.79E-02 | -0.28 | 361 | 3.36E-02 | 6.44E-02 | 0.14 | 213 | 3.00E-01 | 5.41E-01 | -1.08 | 480 | 1.81E-08 | 3.34E-07 |
| ILMN_1673299 | ZNF80 | 3q13.31a | 0.07 | 6.21E-01 | 7.50E-01 | 6.18E-01 | 8.00E-06 | 1.54 | 1.09 | -0.45 | -0.16 | 362 | 3.01E-01 | 4.84E-01 | 1.09 | 56 | 2.09E-07 | 8.00E-06 | 0.10 | 271 | 5.27E-01 | 6.18E-01 | 0.26 | 156 | 1.20E-01 | 3.19E-01 | -0.45 | 386 | 3.93E-02 | 8.42E-02 |
| ILMN_1723815 | NPEPPS | 0 | -0.22 | 1.78E-01 | 3.22E-01 | 8.68E-01 | 1.18E-04 | 1.54 | 0.43 | -1.11 | 0.05 | 321 | 7.79E-01 | 8.68E-01 | -1.11 | 526 | 7.29E-06 | 1.18E-04 | -0.28 | 362 | 1.26E-01 | 1.92E-01 | -0.50 | 485 | 1.01E-02 | 7.22E-02 | 0.43 | 300 | 9.46E-02 | 1.71E-01 |
| ILMN_1851133 | ##_HS.152655 | 0 | -0.17 | 1.75E-01 | 3.19E-01 | 8.30E-01 | 1.33E-06 | 1.54 | 1.07 | -0.48 | -0.22 | 377 | 1.21E-01 | 2.64E-01 | 1.07 | 62 | 1.97E-08 | 1.33E-06 | -0.48 | 408 | 7.51E-04 | 2.46E-03 | 0.06 | 261 | 6.69E-01 | 8.30E-01 | -0.23 | 364 | 2.43E-01 | 3.59E-01 |
| ILMN_1652490 | MANSC1 | 12p13.2a | -0.52 | 3.71E-04 | 3.92E-03 | 6.86E-01 | 7.17E-13 | 1.54 | 0.36 | -1.18 | -0.13 | 356 | 3.54E-01 | 5.42E-01 | -0.11 | 321 | 5.53E-01 | 6.86E-01 | -1.18 | 534 | 6.14E-15 | 7.17E-13 | -0.64 | 507 | 2.35E-05 | 2.46E-03 | 0.36 | 306 | 6.80E-02 | 1.31E-01 |
| ILMN_1810628 | PRUNE2 | 9q21.13c | 0.53 | 8.00E-02 | 1.85E-01 | 9.05E-01 | 1.24E-02 | 1.54 | 1.45 | -0.09 | 0.25 | 281 | 4.83E-01 | 6.56E-01 | -0.09 | 313 | 8.46E-01 | 9.05E-01 | 0.71 | 138 | 4.65E-02 | 8.42E-02 | 0.79 | 49 | 3.62E-02 | 1.57E-01 | 1.45 | 48 | 4.00E-03 | 1.24E-02 |
| ILMN_2050911 | SLC22A4 | 5q31.1b | 0.12 | 2.97E-01 | 4.58E-01 | 8.22E-01 | 3.35E-08 | 1.53 | 1.04 | -0.49 | 0.27 | 269 | 2.00E-02 | 7.39E-02 | -0.49 | 429 | 1.69E-03 | 8.66E-03 | 0.03 | 289 | 7.66E-01 | 8.22E-01 | -0.16 | 387 | 2.09E-01 | 4.41E-01 | 1.04 | 199 | 1.18E-09 | 3.35E-08 |
| ILMN_1785732 | TNFAIP6 | 2q23.3b | 0.73 | 1.70E-06 | 8.06E-05 | 7.12E-01 | 5.62E-10 | 1.53 | 1.64 | 0.12 | 0.82 | 85 | 1.04E-06 | 3.59E-05 | 0.12 | 239 | 5.84E-01 | 7.12E-01 | 0.78 | 126 | 3.41E-06 | 2.23E-05 | 0.42 | 109 | 1.64E-02 | 9.73E-02 | 1.64 | 24 | 8.02E-12 | 5.62E-10 |
| ILMN_1761820 | EDARADD | 1q43a | -0.35 | 9.65E-03 | 4.05E-02 | 5.58E-01 | 2.16E-07 | 1.53 | 0.36 | -1.16 | -0.54 | 473 | 1.59E-04 | 1.87E-03 | 0.36 | 151 | 5.02E-02 | 1.19E-01 | -0.54 | 423 | 1.46E-04 | 5.92E-04 | 0.15 | 210 | 3.17E-01 | 5.58E-01 | -1.16 | 514 | 1.08E-08 | 2.16E-07 |

### Online Supplemental Table S4 (Continued)

| | | | S1 | | | M1 through M5 | | | | | M1 | | | | M2 | | | | M3 | | | | M4 | | | | M5 | | | |
|---|---|---|---|---|---|---|---|---|---|---|---|---|---|---|---|---|---|---|---|---|---|---|---|---|---|---|---|---|---|---|
| ID | Gene_Symbol | Cytoband | FC | $P_{val}$ | $P_{adj}$ | Max $P_{adj}$ | Min $P_{adj}$ | ΔFC | Max FC | Min FC | FC | FC Rank | $P_{val}$ | $P_{adj}$ | FC | FC Rank | $P_{val}$ | $P_{adj}$ | FC | FC Rank | $P_{val}$ | $P_{adj}$ | FC | FC Rank | $P_{val}$ | $P_{adj}$ | FC | FC Rank | $P_{val}$ | $P_{adj}$ |
| ILMN_1676895 | BTNL8 | 0 | -0.44 | 4.92E-03 | 2.51E-02 | 5.15E-01 | 2.84E-09 | 1.52 | 0.44 | -1.09 | -0.16 | 361 | 3.30E-01 | 5.15E-01 | 0.30 | 167 | 1.60E-01 | 2.86E-01 | -1.09 | 520 | 1.09E-10 | 2.84E-09 | -0.52 | 488 | 2.47E-03 | 3.09E-02 | 0.44 | 299 | 5.04E-02 | 1.03E-01 |
| ILMN_2047618 | KCNE1 | 21q22.12a | 0.33 | 1.80E-02 | 6.41E-02 | 8.93E-01 | 5.88E-09 | 1.52 | 1.44 | -0.08 | 0.58 | 140 | 2.13E-04 | 2.34E-03 | -0.08 | 309 | 6.97E-01 | 8.00E-01 | 0.13 | 266 | 3.90E-01 | 4.86E-01 | 0.05 | 272 | 7.76E-01 | 8.93E-01 | 1.44 | 49 | 1.48E-10 | 5.88E-09 |
| ILMN_1751020 | PACSIN1 | 6p21.31e | -0.50 | 3.72E-04 | 3.92E-03 | 5.92E-01 | 1.16E-05 | 1.51 | 0.48 | -1.04 | -0.59 | 489 | 1.06E-04 | 1.38E-03 | 0.48 | 135 | 1.67E-02 | 5.09E-02 | -0.75 | 461 | 1.61E-06 | 1.16E-05 | -0.15 | 382 | 3.53E-01 | 5.92E-01 | -1.04 | 462 | 1.73E-06 | 1.63E-05 |
| ILMN_2179873 | PHC3 | 3q26.2b | 0.63 | 4.57E-07 | 3.09E-05 | 2.22E-01 | 3.40E-10 | 1.51 | 1.24 | -0.27 | 0.58 | 139 | 1.24E-05 | 2.49E-04 | -0.27 | 373 | 1.14E-01 | 2.22E-01 | 0.94 | 98 | 9.37E-12 | 3.40E-10 | 0.40 | 112 | 3.59E-03 | 3.88E-02 | 1.24 | 99 | 7.33E-11 | 3.30E-09 |
| ILMN_2205032 | MAGEE1 | Xq13.3c | -0.02 | 8.73E-01 | 9.26E-01 | 9.69E-01 | 5.89E-07 | 1.50 | 0.39 | -1.11 | -0.18 | 365 | 1.98E-01 | 3.68E-01 | -1.11 | 525 | 6.61E-09 | 5.89E-07 | 0.39 | 205 | 5.20E-03 | 1.31E-02 | -0.01 | 310 | 9.31E-01 | 9.69E-01 | -0.07 | 350 | 7.28E-01 | 8.09E-01 |
| ILMN_1751161 | COL7A1 | 3p21.31e | 0.20 | 8.26E-02 | 1.89E-01 | 7.16E-01 | 4.82E-11 | 1.50 | 1.32 | -0.19 | 0.22 | 289 | 6.86E-02 | 1.78E-01 | -0.19 | 341 | 2.48E-01 | 3.93E-01 | 0.07 | 279 | 5.58E-01 | 6.46E-01 | 0.09 | 242 | 5.02E-01 | 7.16E-01 | 1.32 | 71 | 3.86E-13 | 4.82E-11 |
| ILMN_1816342 | ##_HS.534061 | 0 | -0.44 | 2.37E-03 | 1.49E-02 | 2.74E-01 | 2.67E-05 | 1.50 | 0.42 | -1.08 | -0.25 | 384 | 1.28E-01 | 2.74E-01 | 0.42 | 144 | 5.09E-02 | 1.20E-01 | -0.68 | 454 | 3.73E-05 | 1.79E-04 | -0.45 | 477 | 8.33E-03 | 6.41E-02 | -1.08 | 484 | 3.09E-06 | 2.67E-05 |
| ILMN_1656818 | GPR141 | 7p14.1e | 0.04 | 8.00E-01 | 8.81E-01 | 7.86E-01 | 2.46E-05 | 1.50 | 1.04 | -0.46 | 0.34 | 240 | 2.98E-02 | 9.90E-02 | 0.21 | 198 | 3.02E-01 | 4.52E-01 | -0.46 | 405 | 3.09E-03 | 8.37E-03 | -0.09 | 353 | 6.00E-01 | 7.86E-01 | 1.04 | 201 | 2.81E-06 | 2.46E-05 |
| ILMN_2386967 | VPS54 | 2p15a-p14c | 0.41 | 6.36E-04 | 5.81E-03 | 5.41E-01 | 1.81E-09 | 1.50 | 1.21 | -0.29 | 0.35 | 237 | 5.09E-03 | 2.69E-02 | -0.29 | 379 | 8.21E-02 | 1.73E-01 | 0.68 | 144 | 1.20E-07 | 1.20E-06 | 0.14 | 217 | 3.00E-01 | 5.41E-01 | 1.21 | 110 | 3.54E-11 | 1.81E-09 |
| ILMN_2089977 | FKBP9P1 | 7p11.2b | 0.02 | 9.17E-01 | 9.54E-01 | 6.11E-01 | 1.04E-03 | 1.50 | 1.01 | -0.48 | 0.33 | 244 | 8.96E-02 | 2.15E-01 | 0.47 | 136 | 6.23E-02 | 1.40E-01 | -0.48 | 412 | 1.30E-02 | 2.87E-02 | -0.18 | 396 | 3.73E-01 | 6.11E-01 | 1.01 | 223 | 2.16E-04 | 1.04E-03 |
| ILMN_2319000 | MATK | 19p13.3e | -0.65 | 2.60E-07 | 2.01E-05 | 4.82E-01 | 1.86E-10 | 1.49 | 0.16 | -1.33 | -0.74 | 519 | 2.19E-08 | 2.11E-06 | 0.16 | 217 | 3.30E-01 | 4.82E-01 | -0.92 | 486 | 9.01E-12 | 3.29E-10 | -0.21 | 415 | 1.15E-01 | 3.11E-01 | -1.33 | 536 | 1.98E-12 | 1.86E-10 |
| ILMN_1898771 | ##_HS.494932 | 0 | -0.36 | 2.87E-02 | 8.99E-02 | 9.97E-01 | 2.37E-07 | 1.48 | 0.00 | -1.48 | -0.61 | 495 | 7.01E-04 | 5.93E-03 | 0.00 | 282 | 9.94E-01 | 9.97E-01 | -0.14 | 335 | 4.51E-01 | 5.45E-01 | -0.03 | 323 | 8.67E-01 | 9.40E-01 | -1.48 | 544 | 1.21E-08 | 2.37E-07 |
| ILMN_1728898 | SPINK8 | 3p21.31f | 0.15 | 2.39E-01 | 3.95E-01 | 9.70E-01 | 5.74E-08 | 1.48 | 1.25 | -0.23 | 0.31 | 249 | 2.94E-02 | 9.78E-02 | -0.23 | 358 | 2.23E-01 | 3.64E-01 | -0.05 | 315 | 7.10E-01 | 7.77E-01 | -0.01 | 309 | 9.34E-01 | 9.70E-01 | 1.25 | 96 | 2.23E-09 | 5.74E-08 |
| ILMN_1705364 | BAG6 | 6p21.33a | -0.43 | 1.09E-04 | 1.69E-03 | 1.70E-01 | 3.84E-11 | 1.48 | 0.46 | -1.02 | -0.20 | 370 | 6.37E-02 | 1.70E-01 | 0.46 | 138 | 1.19E-03 | 6.52E-03 | -0.81 | 476 | 7.17E-13 | 3.84E-11 | -0.39 | 467 | 5.87E-04 | 1.41E-02 | -1.02 | 446 | 7.58E-11 | 3.35E-09 |
| ILMN_1760688 | SAMD14 | 17q21.33a | -0.73 | 1.10E-03 | 8.57E-03 | 6.86E-01 | 5.55E-06 | 1.47 | 0.19 | -1.28 | -0.45 | 450 | 6.94E-02 | 1.80E-01 | 0.19 | 203 | 5.53E-01 | 6.86E-01 | -1.28 | 544 | 6.87E-07 | 5.55E-06 | -0.68 | 515 | 1.03E-02 | 7.29E-02 | -0.80 | 413 | 2.29E-02 | 5.39E-02 |
| ILMN_1899760 | ##_HS.438905 | 0 | 0.38 | 2.30E-03 | 1.46E-02 | 2.63E-01 | 2.35E-06 | 1.47 | 1.07 | -0.40 | 0.44 | 200 | 2.22E-03 | 1.46E-02 | -0.40 | 410 | 3.20E-02 | 8.41E-02 | 0.39 | 206 | 6.19E-03 | 1.53E-02 | 0.26 | 155 | 8.74E-02 | 2.63E-01 | 1.07 | 175 | 1.80E-07 | 2.35E-06 |
| ILMN_1758057 | TOR1AIP2 | 1q25.2c | 0.31 | 4.58E-03 | 2.38E-02 | 2.15E-01 | 1.59E-08 | 1.47 | 1.07 | -0.40 | 0.32 | 247 | 6.53E-03 | 3.25E-02 | -0.40 | 408 | 1.05E-02 | 3.58E-02 | 0.32 | 222 | 7.10E-03 | 1.71E-02 | 0.23 | 168 | 6.18E-02 | 2.15E-01 | 1.07 | 176 | 4.76E-10 | 1.59E-08 |
| ILMN_1673119 | AFF1 | 4q21.3e-q22.1a | 0.52 | 7.74E-04 | 6.67E-03 | 9.50E-01 | 1.25E-07 | 1.47 | 1.44 | -0.02 | 0.65 | 117 | 1.66E-04 | 1.93E-03 | -0.02 | 288 | 9.15E-01 | 9.50E-01 | 0.36 | 211 | 3.44E-02 | 6.55E-02 | 0.42 | 107 | 2.08E-02 | 1.13E-01 | 1.44 | 50 | 5.68E-09 | 1.25E-07 |
| ILMN_1744006 | GFOD2 | 16q22.1b | -0.50 | 1.00E-05 | 2.99E-04 | 3.91E-02 | 7.51E-17 | 1.46 | 0.44 | -1.03 | -0.30 | 401 | 3.48E-03 | 2.05E-02 | 0.44 | 139 | 1.22E-03 | 6.66E-03 | -1.03 | 505 | 7.62E-20 | 7.51E-17 | -0.32 | 437 | 3.66E-03 | 3.91E-02 | -0.76 | 410 | 2.23E-07 | 2.81E-06 |
| ILMN_1812915 | TNFRSF10B | 8p21.3a | -0.62 | 2.03E-06 | 9.21E-05 | 2.52E-01 | 3.15E-15 | 1.46 | 0.25 | -1.21 | -0.34 | 411 | 9.58E-03 | 4.32E-02 | 0.25 | 187 | 1.35E-01 | 2.52E-01 | -1.21 | 538 | 7.95E-18 | 3.15E-15 | -0.65 | 510 | 3.18E-06 | 8.40E-04 | -0.28 | 373 | 1.25E-01 | 2.13E-01 |
| ILMN_1803479 | ##_LOC650546 | 0 | 0.28 | 1.90E-02 | 6.66E-02 | 9.14E-01 | 6.68E-09 | 1.46 | 1.18 | -0.28 | 0.48 | 180 | 2.14E-04 | 2.35E-03 | -0.28 | 376 | 9.72E-02 | 1.97E-01 | 0.22 | 250 | 7.98E-02 | 1.32E-01 | -0.03 | 322 | 8.14E-01 | 9.14E-01 | 1.18 | 117 | 1.73E-10 | 6.68E-09 |
| ILMN_1805985 | SLF1 | 5q15b | 0.50 | 9.63E-07 | 5.22E-05 | 8.25E-02 | 2.68E-11 | 1.46 | 1.14 | -0.32 | 0.55 | 147 | 2.04E-07 | 1.09E-05 | -0.32 | 390 | 1.80E-02 | 5.42E-02 | 0.67 | 146 | 6.15E-10 | 1.28E-08 | 0.27 | 146 | 1.25E-02 | 8.25E-02 | 1.14 | 135 | 1.61E-13 | 2.68E-11 |
| ILMN_1669323 | BACE2 | 21q22.3a | -0.27 | 1.26E-01 | 2.54E-01 | 6.56E-01 | 1.26E-03 | 1.46 | 0.43 | -1.03 | -0.32 | 404 | 1.13E-01 | 2.52E-01 | 0.43 | 141 | 1.06E-01 | 2.10E-01 | -0.47 | 407 | 1.89E-02 | 3.97E-02 | 0.17 | 198 | 4.23E-01 | 6.56E-01 | -1.03 | 457 | 2.70E-04 | 1.26E-03 |
| ILMN_1816603 | SLC8A1 | 0 | 0.19 | 2.67E-01 | 4.26E-01 | 9.78E-01 | 5.32E-04 | 1.45 | 1.08 | -0.37 | 0.53 | 153 | 7.01E-03 | 3.43E-02 | -0.37 | 403 | 1.55E-01 | 2.79E-01 | -0.09 | 323 | 6.44E-01 | 7.21E-01 | -0.01 | 308 | 9.52E-01 | 9.78E-01 | 1.08 | 166 | 9.96E-05 | 5.32E-04 |
| ILMN_1803838 | CNFN | 19q13.2c | -0.35 | 7.69E-03 | 3.45E-02 | 8.17E-01 | 1.02E-06 | 1.45 | 0.35 | -1.10 | -0.42 | 436 | 3.12E-03 | 1.89E-02 | 0.35 | 154 | 5.67E-02 | 1.30E-01 | -0.47 | 406 | 9.91E-04 | 3.12E-03 | -0.07 | 344 | 6.48E-01 | 8.17E-01 | -1.10 | 490 | 6.68E-08 | 1.02E-06 |
| ILMN_1688865 | PPP1R9B | 17q21.33a | 0.27 | 8.26E-02 | 1.89E-01 | 8.57E-01 | 5.31E-06 | 1.44 | 1.25 | -0.19 | 0.41 | 212 | 1.87E-02 | 7.07E-02 | -0.06 | 298 | 7.76E-01 | 8.57E-01 | 0.30 | 227 | 8.18E-02 | 1.34E-01 | -0.19 | 402 | 2.92E-01 | 5.33E-01 | 1.25 | 92 | 4.63E-07 | 5.31E-06 |
| ILMN_1668928 | MLKL | 16q22.3c | 0.26 | 2.03E-01 | 3.53E-01 | 9.95E-01 | 3.43E-04 | 1.44 | 1.35 | -0.10 | 0.51 | 162 | 3.19E-02 | 1.04E-01 | 0.27 | 178 | 3.80E-01 | 5.32E-01 | 0.00 | 296 | 9.92E-01 | 9.95E-01 | -0.10 | 359 | 7.03E-01 | 8.51E-01 | 1.35 | 61 | 6.07E-05 | 3.43E-04 |
| ILMN_1721008 | DUT | 15q21.1d | -0.56 | 8.32E-07 | 4.67E-05 | 5.93E-01 | 3.64E-11 | 1.44 | 0.12 | -1.32 | -0.61 | 493 | 1.02E-06 | 3.59E-05 | 0.12 | 238 | 4.44E-01 | 5.93E-01 | -0.59 | 432 | 2.10E-06 | 1.46E-05 | -0.34 | 451 | 8.32E-03 | 6.41E-02 | -1.32 | 533 | 2.49E-13 | 3.64E-11 |
| ILMN_1747759 | WSB1 | 17q11.1b | 0.57 | 8.71E-06 | 2.69E-04 | 6.87E-01 | 3.12E-09 | 1.43 | 1.32 | -0.11 | 0.75 | 98 | 1.53E-07 | 9.03E-06 | -0.11 | 320 | 5.55E-01 | 6.87E-01 | 0.58 | 167 | 4.43E-05 | 2.08E-04 | 0.30 | 139 | 4.14E-02 | 1.70E-01 | 1.32 | 68 | 6.83E-11 | 3.12E-09 |
| ILMN_1667408 | EPB41 | 1p35.3a | -0.53 | 2.13E-03 | 1.38E-02 | 4.92E-01 | 2.75E-07 | 1.43 | 0.37 | -1.06 | -0.19 | 367 | 3.09E-01 | 4.92E-01 | 0.37 | 149 | 1.24E-01 | 2.36E-01 | -1.06 | 514 | 2.21E-08 | 2.75E-07 | -0.49 | 480 | 1.07E-02 | 7.48E-02 | -0.85 | 416 | 1.04E-03 | 3.99E-03 |
| ILMN_2363586 | SDCBP | 8q12.1c | 0.21 | 2.58E-02 | 8.30E-02 | 4.60E-01 | 1.62E-11 | 1.43 | 1.03 | -0.39 | 0.30 | 255 | 1.21E-03 | 9.10E-03 | -0.39 | 407 | 1.50E-03 | 7.88E-03 | 0.29 | 231 | 2.04E-03 | 5.84E-03 | -0.12 | 366 | 2.23E-01 | 4.60E-01 | 1.03 | 203 | 8.79E-14 | 1.62E-11 |
| ILMN_2256295 | GNLY | 2p11.2f | -0.47 | 3.80E-02 | 1.10E-01 | 7.43E-01 | 7.05E-03 | 1.43 | 0.28 | -1.14 | -0.55 | 477 | 3.82E-02 | 1.18E-01 | 0.28 | 173 | 4.16E-01 | 5.66E-01 | -0.59 | 433 | 2.47E-02 | 4.98E-02 | -0.17 | 393 | 5.36E-01 | 7.43E-01 | -1.14 | 506 | 2.04E-03 | 7.05E-03 |
| ILMN_1769849 | HORMAD1 | 1q21.2c | 0.25 | 1.96E-02 | 6.82E-02 | 7.62E-01 | 1.75E-10 | 1.43 | 1.17 | -0.25 | 0.35 | 238 | 2.06E-03 | 1.38E-02 | -0.25 | 366 | 8.70E-02 | 1.80E-01 | 0.18 | 259 | 1.03E-01 | 1.62E-01 | 0.07 | 256 | 5.66E-01 | 7.62E-01 | 1.17 | 119 | 1.82E-12 | 1.75E-10 |
| ILMN_1658071 | ATP1B1 | 1q24.2b | 0.66 | 5.64E-05 | 1.06E-03 | 4.62E-01 | 1.10E-09 | 1.43 | 1.20 | -0.23 | 0.58 | 136 | 7.79E-04 | 6.47E-03 | -0.23 | 357 | 3.11E-01 | 4.62E-01 | 1.20 | 38 | 3.67E-11 | 1.10E-09 | 0.32 | 130 | 8.01E-02 | 2.51E-01 | 0.78 | 257 | 1.33E-03 | 4.94E-03 |
| ILMN_1673521 | KISS1R | 19p13.3i | -0.83 | 1.46E-05 | 3.89E-04 | 8.11E-01 | 1.65E-08 | 1.42 | 0.10 | -1.32 | -0.87 | 537 | 3.47E-05 | 5.73E-04 | 0.10 | 245 | 7.11E-01 | 8.11E-01 | -1.32 | 552 | 8.35E-10 | 1.65E-08 | -0.26 | 426 | 2.33E-01 | 4.71E-01 | -1.22 | 521 | 3.39E-05 | 2.09E-04 |
| ILMN_1769734 | NT5C3A | 7p14.3c-p14.3b | 0.46 | 1.08E-05 | 3.17E-04 | 6.00E-01 | 1.28E-10 | 1.42 | 1.06 | -0.36 | 0.52 | 158 | 5.33E-07 | 2.21E-05 | -0.36 | 400 | 7.28E-03 | 2.70E-02 | 0.74 | 132 | 5.77E-12 | 2.27E-10 | 0.10 | 235 | 3.61E-01 | 6.00E-01 | 1.06 | 182 | 1.21E-12 | 1.28E-10 |
| ILMN_1882093 | ##_HS.30972 | 0 | 0.23 | 4.39E-02 | 1.22E-01 | 9.83E-01 | 1.41E-06 | 1.42 | 1.00 | -0.42 | 0.30 | 256 | 2.11E-02 | 7.69E-02 | -0.42 | 414 | 1.57E-02 | 4.86E-02 | 0.25 | 245 | 5.22E-02 | 9.28E-02 | 0.01 | 299 | 9.62E-01 | 9.83E-01 | 1.00 | 232 | 9.89E-08 | 1.41E-06 |
| ILMN_1749403 | TSPAN33 | 7q32.1a | -0.45 | 6.09E-04 | 5.64E-03 | 3.67E-01 | 3.21E-07 | 1.41 | 0.32 | -1.09 | -0.32 | 406 | 2.37E-02 | 8.29E-02 | 0.32 | 160 | 8.07E-02 | 1.71E-01 | -0.81 | 475 | 2.64E-08 | 3.21E-07 | -0.21 | 414 | 1.54E-01 | 3.67E-01 | -1.09 | 486 | 7.64E-08 | 1.14E-06 |
| ILMN_2066060 | HLA-DRB6 | 6p21.32b | 0.81 | 1.99E-02 | 6.88E-02 | 4.68E-01 | 3.33E-03 | 1.41 | 1.84 | 0.42 | 0.42 | 207 | 2.86E-01 | 4.68E-01 | 1.84 | 4 | 5.07E-04 | 3.33E-03 | 0.76 | 128 | 5.53E-02 | 9.74E-02 | 1.25 | 11 | 3.04E-03 | 3.48E-02 | 0.57 | 285 | 3.07E-01 | 4.29E-01 |
| ILMN_2215355 | LOC400499 | 16p13.13b | 0.38 | 4.51E-02 | 1.24E-01 | 9.14E-01 | 4.93E-05 | 1.41 | 1.36 | -0.05 | 0.55 | 146 | 9.00E-03 | 4.14E-02 | 0.55 | 126 | 4.70E-02 | 1.13E-01 | 0.21 | 252 | 3.22E-01 | 4.16E-01 | -0.05 | 334 | 8.15E-01 | 9.14E-01 | 1.36 | 60 | 6.30E-06 | 4.93E-05 |
| ILMN_1746317 | NUP58 | 13q12.13a | 0.61 | 1.07E-04 | 1.66E-03 | 7.29E-01 | 1.58E-05 | 1.41 | 1.14 | -0.27 | 0.85 | 79 | 6.40E-07 | 2.49E-05 | -0.27 | 374 | 2.14E-01 | 3.54E-01 | 0.81 | 116 | 2.30E-06 | 1.58E-05 | 0.11 | 225 | 5.18E-01 | 7.29E-01 | 1.14 | 134 | 2.12E-06 | 1.93E-05 |
| ILMN_1735712 | KRT1 | 12q13.13d | -0.28 | 3.16E-01 | 4.78E-01 | 7.82E-01 | 3.67E-02 | 1.41 | 0.30 | -1.11 | 0.15 | 308 | 6.46E-01 | 7.82E-01 | 0.30 | 166 | 4.76E-01 | 6.21E-01 | -0.72 | 459 | 2.63E-02 | 5.25E-02 | -0.35 | 458 | 3.03E-01 | 5.43E-01 | -1.11 | 493 | 1.45E-02 | 3.67E-02 |
| ILMN_2209671 | DCDC1 | 11p14.1a-p13f | -0.51 | 5.23E-05 | 9.96E-04 | 9.64E-01 | 1.23E-07 | 1.41 | 0.31 | -1.09 | -0.76 | 523 | 1.49E-08 | 1.56E-06 | 0.31 | 162 | 6.58E-02 | 1.46E-01 | -0.67 | 451 | 4.96E-07 | 4.18E-06 | -0.01 | 311 | 9.18E-01 | 9.64E-01 | -1.09 | 488 | 5.53E-09 | 1.23E-07 |
| ILMN_1798588 | HLTF | 3q24f | 0.02 | 9.14E-01 | 9.52E-01 | 6.89E-01 | 1.44E-05 | 1.41 | 0.34 | -1.07 | -0.10 | 348 | 5.22E-01 | 6.89E-01 | -1.07 | 517 | 4.67E-07 | 1.44E-05 | 0.34 | 216 | 3.19E-02 | 6.16E-02 | 0.17 | 199 | 3.09E-01 | 5.49E-01 | -0.16 | 357 | 4.70E-01 | 5.88E-01 |
| ILMN_2271875 | ALAS2 | Xp11.21a | -0.54 | 1.92E-02 | 6.71E-02 | 9.62E-01 | 9.29E-04 | 1.40 | 0.28 | -1.13 | 0.02 | 330 | 9.30E-01 | 9.62E-01 | 0.28 | 175 | 4.08E-01 | 5.59E-01 | -0.95 | 491 | 2.46E-04 | 9.29E-04 | -0.89 | 543 | 1.15E-03 | 2.02E-02 | -1.13 | 501 | 1.85E-03 | 6.51E-03 |
| ILMN_1671478 | CKB | 14q32.33a | 0.34 | 6.89E-02 | 1.67E-01 | 4.09E-01 | 2.99E-03 | 1.40 | 1.02 | -0.38 | 0.32 | 246 | 1.38E-01 | 2.89E-01 | 1.02 | 75 | 4.42E-04 | 2.99E-03 | 0.43 | 194 | 5.05E-02 | 9.02E-02 | 0.31 | 136 | 1.84E-01 | 4.09E-01 | -0.38 | 382 | 2.14E-01 | 3.25E-01 |
| ILMN_1661461 | LINC00639 | 14q21.1b | 0.18 | 2.84E-01 | 4.45E-01 | 9.64E-01 | 3.49E-05 | 1.40 | 1.23 | -0.17 | 0.37 | 233 | 5.10E-02 | 1.44E-01 | -0.17 | 340 | 4.86E-01 | 6.29E-01 | 0.01 | 293 | 9.49E-01 | 9.64E-01 | -0.05 | 337 | 7.82E-01 | 8.97E-01 | 1.23 | 103 | 4.22E-06 | 3.49E-05 |
| ILMN_2112256 | TNFRSF4 | 1p36.33b | 0.06 | 6.13E-01 | 7.45E-01 | 7.28E-01 | 4.38E-07 | 1.40 | 1.06 | -0.34 | -0.07 | 342 | 5.73E-01 | 7.28E-01 | 1.06 | 67 | 4.51E-09 | 4.38E-07 | -0.11 | 327 | 3.97E-01 | 4.94E-01 | 0.33 | 126 | 1.86E-02 | 1.05E-01 | -0.34 | 378 | 6.34E-02 | 1.24E-01 |
| ILMN_1792356 | DPYSL4 | 10q26.3e | 0.27 | 3.59E-01 | 5.21E-01 | 9.90E-01 | 2.03E-02 | 1.40 | 1.30 | -0.10 | 0.01 | 334 | 9.81E-01 | 9.90E-01 | 1.30 | 28 | 5.04E-03 | 2.03E-02 | 0.13 | 267 | 7.15E-01 | 7.81E-01 | 0.72 | 58 | 5.04E-02 | 1.91E-01 | -0.10 | 354 | 8.39E-01 | 8.91E-01 |
| ILMN_1743939 | C5orf24 | 5q31.1e | 0.64 | 1.84E-05 | 4.67E-04 | 9.92E-01 | 2.70E-07 | 1.40 | 1.39 | 0.00 | 0.64 | 122 | 1.96E-04 | 2.21E-03 | 0.00 | 283 | 9.86E-01 | 9.92E-01 | 0.73 | 134 | 2.38E-05 | 1.20E-04 | 0.49 | 90 | 6.81E-03 | 5.70E-02 | 1.39 | 56 | 1.42E-08 | 2.70E-07 |
| ILMN_2172318 | TAFA2 | 12q14.1d | 0.14 | 3.73E-01 | 5.35E-01 | 8.16E-01 | 3.65E-04 | 1.39 | 1.06 | -0.33 | 0.27 | 270 | 1.50E-01 | 3.07E-01 | -0.33 | 394 | 1.77E-01 | 3.07E-01 | 0.19 | 258 | 3.18E-01 | 4.11E-01 | -0.09 | 356 | 6.46E-01 | 8.16E-01 | 1.06 | 184 | 6.52E-05 | 3.65E-04 |
| ILMN_1797372 | DIPK2A | 3q24a | 0.47 | 4.91E-06 | 1.76E-04 | 5.09E-01 | 2.82E-11 | 1.39 | 1.03 | -0.36 | 0.53 | 154 | 1.64E-07 | 9.50E-06 | -0.36 | 401 | 5.78E-03 | 2.27E-02 | 0.76 | 129 | 4.93E-13 | 2.82E-11 | 0.12 | 224 | 2.68E-01 | 5.09E-01 | 1.03 | 206 | 1.41E-12 | 1.44E-10 |
| ILMN_2273447 | NPRL3 | 16p13.3f | -0.69 | 1.61E-02 | 5.90E-02 | 9.23E-01 | 2.93E-03 | 1.39 | -0.07 | -1.46 | -0.10 | 349 | 7.50E-01 | 8.50E-01 | -0.07 | 303 | 8.72E-01 | 9.23E-01 | -0.94 | 490 | 4.52E-03 | 1.17E-02 | -1.46 | 565 | 3.31E-05 | 2.93E-03 | -0.81 | 414 | 7.70E-02 | 1.45E-01 |
| ILMN_2374352 | DBNDD1 | 16q24.3b | -0.49 | 2.10E-03 | 1.37E-02 | 9.47E-01 | 1.33E-06 | 1.39 | 0.03 | -1.36 | -0.73 | 518 | 4.47E-05 | 6.92E-04 | -0.22 | 348 | 3.53E-01 | 5.04E-01 | -0.49 | 413 | 6.54E-03 | 1.60E-02 | 0.03 | 282 | 8.81E-01 | 9.47E-01 | -1.36 | 540 | 9.12E-08 | 1.33E-06 |
| ILMN_1678859 | ##_LOC649210 | 0 | -0.42 | 1.01E-01 | 2.19E-01 | 5.48E-01 | 3.48E-02 | 1.39 | 0.34 | -1.05 | -0.40 | 427 | 1.90E-01 | 3.58E-01 | 0.34 | 156 | 3.97E-01 | 5.48E-01 | -0.50 | 417 | 9.77E-02 | 1.56E-01 | -0.38 | 465 | 2.36E-01 | 4.73E-01 | -1.05 | 466 | 1.36E-02 | 3.48E-02 |
| ILMN_1763207 | BATF3 | 1q32.3c | 0.33 | 2.11E-02 | 7.17E-02 | 2.96E-01 | 3.20E-05 | 1.39 | 1.08 | -0.30 | 0.33 | 245 | 4.92E-02 | 1.41E-01 | 1.08 | 59 | 1.32E-06 | 3.20E-05 | 0.31 | 225 | 6.53E-02 | 1.12E-01 | 0.41 | 111 | 2.04E-02 | 1.11E-01 | -0.30 | 374 | 1.89E-01 | 2.96E-01 |
| ILMN_2189406 | ARPIN/ARPIN-AP3S2 | 15q26.1b | 0.25 | 1.76E-01 | 3.20E-01 | 9.59E-01 | 1.81E-05 | 1.39 | 1.40 | 0.02 | 0.38 | 228 | 6.94E-02 | 1.80E-01 | 1.40 | 18 | 6.42E-07 | 1.81E-05 | 0.02 | 292 | 9.42E-01 | 9.59E-01 | 0.07 | 251 | 7.37E-01 | 8.72E-01 | 0.17 | 325 | 5.53E-01 | 6.63E-01 |
| ILMN_2206141 | BAGE5 | 0 | 0.30 | 6.97E-02 | 1.68E-01 | 4.68E-01 | 1.01E-04 | 1.38 | 1.13 | -0.25 | 0.53 | 156 | 4.25E-03 | 2.36E-02 | -0.24 | 362 | 3.17E-01 | 4.68E-01 | 0.40 | 203 | 2.90E-02 | 5.68E-02 | -0.25 | 425 | 1.93E-01 | 4.21E-01 | 1.13 | 142 | 1.45E-05 | 1.01E-04 |
| ILMN_1651752 | TASL | Xp21.2a | 0.72 | 1.36E-05 | 3.73E-04 | 6.64E-01 | 1.04E-07 | 1.38 | 1.13 | -0.26 | 0.92 | 72 | 2.98E-07 | 1.44E-05 | -0.26 | 370 | 2.68E-01 | 4.15E-01 | 1.05 | 71 | 7.25E-09 | 1.04E-07 | 0.14 | 212 | 4.33E-01 | 6.64E-01 | 1.13 | 146 | 6.65E-06 | 5.16E-05 |
| ILMN_1756676 | PHF19 | 9q33.2a | -0.20 | 2.68E-02 | 8.55E-02 | 7.61E-01 | 4.95E-11 | 1.38 | 0.38 | -1.00 | -0.15 | 359 | 1.07E-01 | 2.44E-01 | 0.38 | 148 | 2.28E-03 | 1.10E-02 | -0.33 | 379 | 4.07E-04 | 1.44E-03 | -0.06 | 339 | 5.65E-01 | 7.61E-01 | -1.00 | 433 | 4.01E-13 | 4.95E-11 |
| ILMN_2396287 | RFX2 | 19p13.3b | -0.83 | 1.94E-05 | 4.82E-04 | 8.19E-01 | 1.76E-07 | 1.38 | 0.10 | -1.28 | -0.69 | 508 | 1.70E-03 | 1.18E-02 | -0.58 | 451 | 4.24E-02 | 1.05E-01 | -1.28 | 545 | 1.33E-08 | 1.76E-07 | -0.87 | 542 | 1.76E-04 | 6.99E-03 | 0.10 | 335 | 7.43E-01 | 8.19E-01 |
| ILMN_1728197 | CLDN5 | 22q11.21c | -0.58 | 4.22E-03 | 2.25E-02 | 6.01E-01 | 1.26E-04 | 1.38 | 0.23 | -1.15 | -0.27 | 389 | 2.44E-01 | 4.21E-01 | 0.23 | 190 | 4.53E-01 | 6.01E-01 | -0.98 | 496 | 2.50E-05 | 1.26E-04 | -0.56 | 494 | 2.08E-02 | 1.13E-01 | -1.15 | 509 | 3.69E-04 | 1.64E-03 |

### Online Supplemental Table S4 (Continued)

| | | | S1 | | | M1 through M5 | | | | | M1 | | | | M2 | | | | M3 | | | | M4 | | | | M5 | | | |
|---|---|---|---|---|---|---|---|---|---|---|---|---|---|---|---|---|---|---|---|---|---|---|---|---|---|---|---|---|---|---|
| ID | Gene_Symbol | Cytoband | FC | $P_{val}$ | $P_{adj}$ | Max $P_{adj}$ | Min $P_{adj}$ | ΔFC | Max FC | Min FC | FC | FC Rank | $P_{val}$ | $P_{adj}$ | FC | FC Rank | $P_{val}$ | $P_{adj}$ | FC | FC Rank | $P_{val}$ | $P_{adj}$ | FC | FC Rank | $P_{val}$ | $P_{adj}$ | FC | FC Rank | $P_{val}$ | $P_{adj}$ |
| ILMN_1690561 | GZMM | 19p13.3j | -0.38 | 4.47E-02 | 1.24E-01 | 8.26E-01 | 1.76E-04 | 1.37 | 0.06 | -1.30 | -0.45 | 448 | 4.22E-02 | 1.26E-01 | -1.30 | 549 | 1.19E-05 | 1.76E-04 | 0.06 | 280 | 7.70E-01 | 8.26E-01 | -0.59 | 499 | 1.22E-02 | 8.14E-02 | -0.59 | 396 | 5.84E-02 | 1.16E-01 |
| ILMN_1786036 | GPATCH2 | 1q41b | 0.57 | 6.26E-06 | 2.13E-04 | 1.13E-01 | 9.79E-12 | 1.37 | 1.03 | -0.34 | 0.38 | 229 | 4.22E-03 | 2.35E-02 | -0.34 | 396 | 4.71E-02 | 1.13E-01 | 1.03 | 83 | 1.41E-13 | 9.79E-12 | 0.45 | 100 | 1.17E-03 | 2.03E-02 | 0.65 | 273 | 4.36E-04 | 1.90E-03 |
| ILMN_1906626 | ##_HS.562110 | 0 | 0.27 | 4.69E-02 | 1.27E-01 | 9.92E-01 | 4.10E-08 | 1.37 | 1.29 | -0.08 | 0.51 | 159 | 5.25E-04 | 4.73E-03 | -0.08 | 306 | 6.93E-01 | 7.97E-01 | 0.00 | 297 | 9.89E-01 | 9.92E-01 | 0.02 | 291 | 9.04E-01 | 9.58E-01 | 1.29 | 78 | 1.49E-09 | 4.10E-08 |
| ILMN_1738468 | KEL | 7q34f | -0.64 | 6.05E-03 | 2.92E-02 | 7.45E-01 | 1.17E-03 | 1.36 | 0.17 | -1.19 | -0.15 | 360 | 5.64E-01 | 7.22E-01 | 0.17 | 214 | 6.25E-01 | 7.45E-01 | -0.96 | 492 | 3.19E-04 | 1.17E-03 | -1.19 | 561 | 2.40E-05 | 2.46E-03 | -0.94 | 421 | 1.13E-02 | 3.00E-02 |
| ILMN_1663484 | HYPK | 15q15.3b | 0.30 | 2.02E-02 | 6.96E-02 | 6.71E-01 | 4.33E-06 | 1.36 | 1.01 | -0.36 | 0.16 | 304 | 2.55E-01 | 4.35E-01 | -0.36 | 398 | 5.02E-02 | 1.19E-01 | 0.60 | 163 | 1.85E-05 | 9.65E-05 | 0.11 | 227 | 4.42E-01 | 6.71E-01 | 1.01 | 230 | 3.69E-07 | 4.33E-06 |
| ILMN_1742230 | BAZ1A | 14q13.2a | 0.38 | 7.67E-07 | 4.43E-05 | 2.53E-01 | 1.33E-18 | 1.36 | 1.06 | -0.29 | 0.44 | 198 | 1.22E-09 | 2.25E-07 | -0.29 | 382 | 1.58E-03 | 8.23E-03 | 0.52 | 173 | 1.66E-12 | 7.63E-11 | 0.13 | 219 | 8.11E-02 | 2.53E-01 | 1.06 | 181 | 1.77E-22 | 1.33E-18 |
| ILMN_1714445 | SLC6A9 | 1p34.1f | -0.69 | 1.95E-03 | 1.30E-02 | 9.05E-01 | 3.15E-06 | 1.36 | 0.06 | -1.29 | -0.23 | 380 | 3.40E-01 | 5.26E-01 | 0.06 | 255 | 8.45E-01 | 9.05E-01 | -1.29 | 546 | 3.63E-07 | 3.15E-06 | -0.83 | 536 | 1.66E-03 | 2.50E-02 | -0.97 | 425 | 5.34E-03 | 1.59E-02 |
| ILMN_1860008 | LOC102724323 | 0 | 0.22 | 1.10E-01 | 2.32E-01 | 7.30E-01 | 1.80E-06 | 1.35 | 1.13 | -0.22 | 0.46 | 190 | 2.08E-03 | 1.39E-02 | 0.33 | 159 | 9.35E-02 | 1.91E-01 | -0.22 | 351 | 1.32E-01 | 2.00E-01 | 0.10 | 233 | 5.19E-01 | 7.30E-01 | 1.13 | 144 | 1.32E-07 | 1.80E-06 |
| ILMN_1767805 | ANKRD20A11P | 21q11.2b-q11.2c | -0.14 | 5.47E-01 | 6.92E-01 | 7.93E-01 | 3.27E-02 | 1.35 | 1.00 | -0.35 | -0.28 | 397 | 3.33E-01 | 5.20E-01 | 1.00 | 82 | 9.37E-03 | 3.27E-02 | -0.11 | 328 | 6.99E-01 | 7.68E-01 | -0.16 | 386 | 6.13E-01 | 7.93E-01 | -0.35 | 379 | 3.89E-01 | 5.12E-01 |
| ILMN_1878885 | IL23A | 0 | -1.03 | 2.52E-07 | 1.97E-05 | 2.34E-01 | 6.90E-07 | 1.35 | -0.47 | -1.82 | -1.25 | 557 | 1.21E-07 | 7.62E-06 | -0.47 | 422 | 1.22E-01 | 2.34E-01 | -0.88 | 483 | 1.72E-04 | 6.82E-04 | -0.79 | 534 | 1.27E-03 | 2.15E-02 | -1.82 | 559 | 4.23E-08 | 6.90E-07 |
| ILMN_2163051 | HARBI1 | 11p11.2c | 0.43 | 1.16E-03 | 8.85E-03 | 7.69E-01 | 9.63E-06 | 1.35 | 1.03 | -0.32 | 0.50 | 169 | 7.50E-04 | 6.26E-03 | -0.32 | 389 | 9.45E-02 | 1.93E-01 | 0.63 | 156 | 2.26E-05 | 1.15E-04 | 0.09 | 243 | 5.76E-01 | 7.69E-01 | 1.03 | 208 | 9.40E-07 | 9.63E-06 |
| ILMN_1670353 | RAD51AP1 | 12p13.32a | 0.54 | 2.56E-04 | 3.02E-03 | 3.91E-01 | 7.02E-11 | 1.35 | 1.12 | -0.23 | 0.39 | 219 | 8.77E-03 | 4.06E-02 | -0.23 | 356 | 2.46E-01 | 3.91E-01 | 1.12 | 50 | 1.48E-12 | 7.02E-11 | 0.28 | 145 | 8.14E-02 | 2.53E-01 | 0.44 | 298 | 3.58E-02 | 7.80E-02 |
| ILMN_2166831 | RPS4X | Xq13.1e | -0.32 | 5.19E-04 | 5.01E-03 | 7.28E-01 | 1.14E-12 | 1.35 | 0.28 | -1.07 | -0.52 | 464 | 3.09E-08 | 2.77E-06 | 0.28 | 176 | 2.04E-02 | 5.97E-02 | -0.32 | 374 | 4.07E-04 | 1.44E-03 | 0.06 | 262 | 5.17E-01 | 7.28E-01 | -1.07 | 477 | 2.80E-15 | 1.14E-12 |
| ILMN_1759023 | WFS1 | 4p16.1f | -0.37 | 4.91E-03 | 2.51E-02 | 5.71E-01 | 8.46E-07 | 1.35 | 0.28 | -1.06 | -0.61 | 491 | 1.09E-05 | 2.26E-04 | 0.28 | 172 | 1.13E-01 | 2.20E-01 | -0.50 | 416 | 2.89E-04 | 1.07E-03 | 0.14 | 216 | 3.32E-01 | 5.71E-01 | -1.06 | 472 | 5.42E-08 | 8.46E-07 |
| ILMN_1763745 | CCNJL | 5q33.3d | -0.67 | 2.45E-06 | 1.05E-04 | 6.67E-01 | 1.14E-12 | 1.35 | 0.12 | -1.23 | -0.42 | 434 | 4.92E-03 | 2.62E-02 | -0.32 | 387 | 1.01E-01 | 2.03E-01 | -1.23 | 540 | 1.06E-14 | 1.14E-12 | -0.72 | 523 | 6.86E-06 | 1.09E-03 | 0.12 | 329 | 5.58E-01 | 6.67E-01 |
| ILMN_1685122 | COL9A2 | 1p34.2d | 0.43 | 3.54E-03 | 1.98E-02 | 7.67E-01 | 5.04E-06 | 1.35 | 1.24 | -0.10 | 0.47 | 183 | 6.39E-03 | 3.20E-02 | -0.10 | 318 | 6.53E-01 | 7.67E-01 | 0.45 | 188 | 9.71E-03 | 2.24E-02 | 0.25 | 161 | 1.68E-01 | 3.89E-01 | 1.24 | 98 | 4.36E-07 | 5.04E-06 |
| ILMN_1903312 | ##_HS.170946 | 0 | -0.18 | 1.69E-01 | 3.12E-01 | 8.22E-01 | 1.35E-05 | 1.35 | 1.01 | -0.33 | -0.27 | 391 | 6.62E-02 | 1.74E-01 | 1.01 | 76 | 4.26E-07 | 1.35E-05 | -0.33 | 378 | 2.55E-02 | 5.12E-02 | -0.20 | 407 | 1.97E-01 | 4.25E-01 | -0.07 | 349 | 7.48E-01 | 8.22E-01 |
| ILMN_1675124 | DDX17 | 22q13.1b | 0.64 | 2.96E-07 | 2.25E-05 | 6.04E-01 | 5.22E-11 | 1.35 | 1.22 | -0.13 | 0.60 | 129 | 4.28E-06 | 1.10E-04 | -0.13 | 323 | 4.57E-01 | 6.04E-01 | 0.97 | 95 | 1.05E-12 | 5.22E-11 | 0.31 | 134 | 2.14E-02 | 1.15E-01 | 1.22 | 108 | 7.66E-11 | 3.37E-09 |
| ILMN_1763561 | OSBPL6 | 2q31.2b | -0.52 | 1.19E-05 | 3.40E-04 | 1.25E-01 | 4.89E-09 | 1.34 | 0.32 | -1.02 | -0.33 | 408 | 9.79E-03 | 4.40E-02 | 0.32 | 161 | 5.36E-02 | 1.25E-01 | -0.84 | 480 | 2.07E-10 | 4.89E-09 | -0.50 | 483 | 2.23E-04 | 8.16E-03 | -1.02 | 451 | 2.04E-08 | 3.71E-07 |
| ILMN_1808979 | CLEC4D | 12p13.31b | 0.34 | 1.05E-01 | 2.24E-01 | 8.53E-01 | 4.44E-03 | 1.34 | 1.10 | -0.24 | 0.51 | 160 | 3.25E-02 | 1.05E-01 | -0.24 | 363 | 4.37E-01 | 5.87E-01 | 0.49 | 180 | 4.10E-02 | 7.58E-02 | -0.09 | 358 | 7.08E-01 | 8.53E-01 | 1.10 | 153 | 1.18E-03 | 4.44E-03 |
| ILMN_1669317 | C5AR2 | 19q13.32b-q13.32c | -0.69 | 6.07E-07 | 3.75E-05 | 9.49E-01 | 6.03E-16 | 1.34 | 0.02 | -1.32 | -0.42 | 432 | 2.52E-03 | 1.61E-02 | 0.02 | 268 | 9.14E-01 | 9.49E-01 | -1.32 | 551 | 1.13E-18 | 6.03E-16 | -0.57 | 496 | 9.10E-05 | 4.98E-03 | -0.32 | 376 | 8.95E-02 | 1.63E-01 |
| ILMN_1722206 | MAF | 16q23.1e | 0.63 | 7.43E-06 | 2.40E-04 | 2.37E-01 | 3.11E-09 | 1.34 | 1.03 | -0.31 | 0.49 | 174 | 1.44E-03 | 1.04E-02 | -0.31 | 386 | 1.25E-01 | 2.37E-01 | 1.03 | 82 | 1.21E-10 | 3.11E-09 | 0.53 | 84 | 1.15E-03 | 2.02E-02 | 0.78 | 256 | 2.95E-04 | 1.37E-03 |
| ILMN_1772189 | ABCD1 | Xq28f | 0.34 | 2.03E-02 | 6.99E-02 | 7.13E-01 | 2.50E-05 | 1.33 | 1.09 | -0.25 | 0.47 | 186 | 4.31E-03 | 2.39E-02 | -0.25 | 364 | 2.52E-01 | 3.97E-01 | 0.46 | 185 | 5.00E-03 | 1.27E-02 | -0.12 | 364 | 4.98E-01 | 7.13E-01 | 1.09 | 161 | 2.87E-06 | 2.50E-05 |
| ILMN_1667444 | ##_LOC644532 | 0 | -1.16 | 4.26E-10 | 2.38E-07 | 5.95E-01 | 4.48E-11 | 1.33 | -0.20 | -1.53 | -1.10 | 551 | 1.21E-07 | 7.62E-06 | -1.20 | 538 | 9.40E-06 | 1.46E-04 | -1.53 | 564 | 8.71E-13 | 4.48E-11 | -1.13 | 557 | 2.26E-07 | 2.55E-04 | -0.20 | 361 | 4.79E-01 | 5.95E-01 |
| ILMN_1760315 | VWCE | 11q12.2b | -0.75 | 1.77E-04 | 2.35E-03 | 7.65E-01 | 2.61E-05 | 1.33 | 0.13 | -1.20 | -0.42 | 433 | 5.65E-02 | 1.56E-01 | 0.13 | 233 | 6.50E-01 | 7.65E-01 | -1.03 | 509 | 4.07E-06 | 2.61E-05 | -1.05 | 554 | 8.66E-06 | 1.26E-03 | -1.20 | 519 | 1.05E-04 | 5.58E-04 |
| ILMN_1785762 | RHOT1 | 17q11.2d | 0.56 | 6.85E-07 | 4.03E-05 | 9.40E-01 | 1.45E-11 | 1.33 | 1.31 | -0.02 | 0.67 | 113 | 3.11E-08 | 2.77E-06 | -0.02 | 286 | 9.01E-01 | 9.40E-01 | 0.66 | 151 | 6.70E-08 | 7.18E-07 | 0.26 | 152 | 3.48E-02 | 1.53E-01 | 1.31 | 73 | 7.72E-14 | 1.45E-11 |
| ILMN_1792396 | GOLGA6L6** | 15q11.2c | 0.28 | 3.93E-02 | 1.12E-01 | 7.16E-01 | 2.88E-07 | 1.33 | 1.26 | -0.07 | 0.38 | 226 | 1.34E-02 | 5.51E-02 | 0.29 | 169 | 1.52E-01 | 2.75E-01 | -0.07 | 318 | 6.38E-01 | 7.16E-01 | 0.33 | 127 | 4.39E-02 | 1.76E-01 | 1.26 | 89 | 1.53E-08 | 2.88E-07 |
| ILMN_1774596 | BSCL2 | 11q12.3a | -0.53 | 1.00E-09 | 3.78E-07 | 1.61E-01 | 3.34E-16 | 1.33 | 0.19 | -1.14 | -0.52 | 465 | 1.01E-09 | 1.96E-07 | 0.19 | 207 | 7.47E-02 | 1.61E-01 | -0.73 | 460 | 1.12E-16 | 2.55E-14 | -0.31 | 436 | 3.48E-04 | 1.04E-02 | -1.14 | 503 | 8.87E-20 | 3.34E-16 |
| ILMN_2120022 | ARL5B | 10p12.33b | 0.53 | 2.29E-05 | 5.46E-04 | 8.43E-01 | 2.99E-08 | 1.33 | 1.04 | -0.29 | 0.72 | 106 | 3.13E-08 | 2.78E-06 | -0.29 | 381 | 7.97E-02 | 1.69E-01 | 0.79 | 122 | 1.68E-09 | 2.99E-08 | 0.05 | 267 | 6.91E-01 | 8.43E-01 | 1.04 | 202 | 1.14E-08 | 2.27E-07 |
| ILMN_2378708 | TAF6 | 7q22.1c | -0.12 | 4.54E-01 | 6.12E-01 | 8.13E-01 | 2.77E-05 | 1.33 | 0.11 | -1.22 | -0.09 | 346 | 6.32E-01 | 7.71E-01 | -1.22 | 542 | 1.09E-06 | 2.77E-05 | 0.11 | 269 | 5.61E-01 | 6.49E-01 | -0.32 | 443 | 9.64E-02 | 2.80E-01 | 0.09 | 336 | 7.33E-01 | 8.13E-01 |
| ILMN_1822671 | YME1L1 | 0 | -1.80 | 3.70E-08 | 5.41E-06 | 1.94E-02 | 1.06E-05 | 1.32 | -1.31 | -2.63 | -1.93 | 565 | 5.78E-07 | 2.33E-05 | -2.01 | 565 | 6.73E-05 | 6.88E-04 | -1.79 | 565 | 3.48E-06 | 2.27E-05 | -1.31 | 563 | 1.07E-03 | 1.94E-02 | -2.63 | 564 | 1.05E-06 | 1.06E-05 |
| ILMN_2047112 | DPCD | 10q24.32a | -0.71 | 2.84E-04 | 3.23E-03 | 6.55E-01 | 2.33E-06 | 1.32 | 0.18 | -1.14 | -0.34 | 418 | 1.07E-01 | 2.43E-01 | 0.18 | 209 | 5.17E-01 | 6.55E-01 | -1.14 | 527 | 2.56E-07 | 2.33E-06 | -0.82 | 535 | 3.38E-04 | 1.02E-02 | -1.11 | 495 | 2.43E-04 | 1.15E-03 |
| ILMN_1776157 | SEPTIN4 | 17q22d | 0.15 | 4.32E-01 | 5.93E-01 | 9.95E-01 | 3.06E-03 | 1.32 | 1.00 | -0.32 | 0.44 | 199 | 4.04E-02 | 1.23E-01 | 1.00 | 80 | 4.56E-04 | 3.06E-03 | -0.32 | 372 | 1.43E-01 | 2.14E-01 | 0.08 | 245 | 7.08E-01 | 8.53E-01 | 0.00 | 347 | 9.91E-01 | 9.95E-01 |
| ILMN_1755843 | SLC26A8 | 6p21.31b | 0.45 | 3.34E-03 | 1.90E-02 | 8.95E-01 | 6.10E-07 | 1.32 | 1.37 | 0.05 | 0.73 | 102 | 2.71E-05 | 4.71E-04 | 0.05 | 257 | 8.30E-01 | 8.95E-01 | 0.10 | 272 | 5.69E-01 | 6.57E-01 | 0.21 | 175 | 2.44E-01 | 4.83E-01 | 1.37 | 58 | 3.63E-08 | 6.10E-07 |
| ILMN_1766551 | CPA3 | 3q24f | -0.76 | 1.56E-02 | 5.78E-02 | 7.86E-01 | 1.21E-02 | 1.32 | -0.21 | -1.52 | -0.72 | 517 | 5.34E-02 | 1.50E-01 | -0.53 | 441 | 2.80E-01 | 4.29E-01 | -1.00 | 498 | 8.41E-03 | 1.98E-02 | -0.21 | 412 | 6.00E-01 | 7.86E-01 | -1.52 | 547 | 3.87E-03 | 1.21E-02 |
| ILMN_1816105 | ##_HS.372928 | 0 | 0.04 | 7.48E-01 | 8.45E-01 | 9.69E-01 | 1.91E-08 | 1.32 | 1.12 | -0.19 | -0.01 | 338 | 9.41E-01 | 9.69E-01 | 1.12 | 54 | 7.49E-11 | 1.91E-08 | -0.19 | 343 | 1.32E-01 | 2.00E-01 | 0.19 | 185 | 1.54E-01 | 3.68E-01 | -0.19 | 359 | 2.73E-01 | 3.93E-01 |
| ILMN_1712755 | LRRC41 | 1p33d | -0.28 | 4.56E-03 | 2.38E-02 | 9.26E-01 | 6.40E-11 | 1.32 | 0.22 | -1.10 | -0.27 | 392 | 8.14E-03 | 3.84E-02 | 0.22 | 194 | 1.04E-01 | 2.06E-01 | -0.48 | 411 | 4.63E-06 | 2.91E-05 | 0.02 | 289 | 8.41E-01 | 9.26E-01 | -1.10 | 489 | 5.52E-13 | 6.40E-11 |
| ILMN_1699085 | ##_C8ORF16 | 8p23.1b | 0.53 | 4.64E-03 | 2.41E-02 | 5.25E-01 | 1.15E-05 | 1.31 | 1.04 | -0.28 | 0.39 | 221 | 6.59E-02 | 1.74E-01 | -0.28 | 375 | 3.20E-01 | 4.72E-01 | 1.04 | 78 | 1.59E-06 | 1.15E-05 | 0.24 | 167 | 2.84E-01 | 5.25E-01 | 0.77 | 259 | 9.15E-03 | 2.50E-02 |
| ILMN_1722025 | CPEB4 | 5q35.2a-q35.2b | 0.31 | 6.43E-02 | 1.59E-01 | 5.23E-01 | 3.18E-04 | 1.31 | 1.06 | -0.25 | 0.65 | 119 | 5.15E-04 | 4.67E-03 | -0.22 | 349 | 3.71E-01 | 5.23E-01 | 0.28 | 234 | 1.26E-01 | 1.93E-01 | -0.25 | 423 | 2.02E-01 | 4.32E-01 | 1.06 | 185 | 5.54E-05 | 3.18E-04 |
| ILMN_1724668 | REPS2 | Xp22.2 | -0.51 | 8.30E-04 | 7.01E-03 | 8.97E-01 | 4.98E-11 | 1.31 | 0.14 | -1.17 | -0.11 | 351 | 4.91E-01 | 6.63E-01 | 0.14 | 229 | 5.01E-01 | 6.42E-01 | -1.17 | 532 | 9.95E-13 | 4.98E-11 | -0.59 | 500 | 3.50E-04 | 1.04E-02 | 0.04 | 342 | 8.47E-01 | 8.97E-01 |
| ILMN_1653466 | HES4 | 1p36.33b | 1.15 | 1.26E-05 | 3.54E-04 | 4.03E-01 | 2.22E-04 | 1.31 | 1.76 | 0.46 | 1.26 | 31 | 4.88E-05 | 7.40E-04 | 1.76 | 6 | 1.58E-05 | 2.22E-04 | 1.10 | 57 | 3.88E-04 | 1.38E-03 | 1.14 | 16 | 4.76E-04 | 1.24E-02 | 0.46 | 295 | 2.83E-01 | 4.03E-01 |
| ILMN_1758087 | TAOK1 | 17q11.2b | 0.42 | 2.92E-02 | 9.10E-02 | 6.06E-01 | 2.94E-03 | 1.31 | 1.01 | -0.30 | 0.59 | 132 | 5.65E-03 | 2.92E-02 | -0.30 | 384 | 2.90E-01 | 4.39E-01 | 0.72 | 137 | 9.22E-04 | 2.94E-03 | -0.20 | 409 | 3.68E-01 | 6.06E-01 | 1.01 | 227 | 8.25E-04 | 3.28E-03 |
| ILMN_1751393 | ZNF684 | 1p34.2c | 1.01 | 9.39E-09 | 2.08E-06 | 5.16E-01 | 4.18E-11 | 1.30 | 1.52 | 0.22 | 1.08 | 53 | 1.35E-08 | 1.44E-06 | 0.22 | 196 | 3.64E-01 | 5.16E-01 | 1.39 | 22 | 8.02E-13 | 4.18E-11 | 0.56 | 80 | 4.35E-03 | 4.32E-02 | 1.52 | 39 | 9.76E-09 | 1.99E-07 |
| ILMN_2069632 | GTSF1 | 12q13.2a | -0.25 | 1.35E-01 | 2.66E-01 | 6.32E-01 | 3.42E-04 | 1.30 | 0.19 | -1.11 | -0.14 | 358 | 4.55E-01 | 6.32E-01 | 0.19 | 205 | 4.49E-01 | 5.97E-01 | -0.26 | 358 | 1.79E-01 | 2.57E-01 | -0.20 | 404 | 3.41E-01 | 5.80E-01 | -1.11 | 494 | 6.03E-05 | 3.42E-04 |
| ILMN_1684227 | GPR146 | 7p22.3b | -0.37 | 7.13E-02 | 1.71E-01 | 9.53E-01 | 7.00E-03 | 1.30 | 0.29 | -1.01 | 0.02 | 328 | 9.16E-01 | 9.53E-01 | 0.29 | 168 | 3.34E-01 | 4.85E-01 | -0.65 | 445 | 5.27E-03 | 1.33E-02 | -0.60 | 504 | 1.39E-02 | 8.83E-02 | -1.01 | 435 | 2.02E-03 | 7.00E-03 |
| ILMN_1766363 | FLT3 | 13q12.2b | 0.04 | 8.39E-01 | 9.06E-01 | 8.97E-01 | 1.78E-03 | 1.30 | 1.06 | -0.23 | 0.16 | 303 | 4.40E-01 | 6.18E-01 | 0.10 | 244 | 7.12E-01 | 8.12E-01 | -0.23 | 354 | 2.70E-01 | 3.60E-01 | -0.06 | 341 | 7.83E-01 | 8.97E-01 | 1.06 | 179 | 4.03E-04 | 1.78E-03 |
| ILMN_1854031 | ##_HS.214235 | 0 | 0.57 | 6.91E-04 | 6.17E-03 | 9.17E-01 | 5.36E-06 | 1.30 | 1.34 | 0.04 | 0.75 | 100 | 7.27E-05 | 1.03E-03 | 0.66 | 114 | 7.33E-03 | 2.72E-02 | 0.60 | 164 | 1.41E-03 | 4.23E-03 | 0.04 | 274 | 8.20E-01 | 9.17E-01 | 1.34 | 64 | 4.68E-07 | 5.36E-06 |
| ILMN_1879482 | ##_HS.560732 | 0 | 0.05 | 7.21E-01 | 8.24E-01 | 9.33E-01 | 5.74E-06 | 1.30 | 1.17 | -0.12 | 0.02 | 327 | 8.79E-01 | 9.33E-01 | 1.17 | 44 | 1.37E-07 | 5.74E-06 | -0.12 | 332 | 4.50E-01 | 5.44E-01 | 0.04 | 275 | 8.00E-01 | 9.05E-01 | 0.24 | 319 | 2.94E-01 | 4.16E-01 |
| ILMN_1695414 | ASF1B | 19p13.12c | -0.80 | 7.00E-08 | 7.81E-06 | 8.30E-01 | 1.82E-14 | 1.29 | -0.07 | -1.36 | -0.44 | 443 | 3.50E-03 | 2.06E-02 | -0.07 | 299 | 7.39E-01 | 8.30E-01 | -1.36 | 557 | 7.60E-17 | 1.82E-14 | -0.71 | 521 | 1.10E-05 | 1.41E-03 | -1.14 | 505 | 1.28E-07 | 1.76E-06 |
| ILMN_1807300 | PKD2 | 4q22.1b | 0.63 | 1.76E-09 | 6.03E-07 | 3.69E-01 | 3.55E-13 | 1.29 | 1.13 | -0.17 | 0.64 | 123 | 4.65E-09 | 5.93E-07 | -0.17 | 338 | 2.28E-01 | 3.69E-01 | 0.89 | 103 | 2.64E-15 | 3.55E-13 | 0.41 | 110 | 2.50E-04 | 8.61E-03 | 1.13 | 145 | 4.32E-13 | 5.20E-11 |
| ILMN_1790771 | ##_LOC650795 | 0 | -0.28 | 3.59E-02 | 1.06E-01 | 3.71E-01 | 1.43E-06 | 1.29 | 0.22 | -1.07 | -0.62 | 496 | 1.51E-05 | 2.92E-04 | 0.22 | 192 | 2.29E-01 | 3.71E-01 | -0.22 | 350 | 1.12E-01 | 1.74E-01 | 0.21 | 174 | 1.46E-01 | 3.58E-01 | -1.07 | 478 | 1.01E-07 | 1.43E-06 |
| ILMN_1732049 | DPM2 | 9q34.11a | -0.53 | 1.06E-03 | 8.36E-03 | 4.00E-01 | 5.49E-05 | 1.29 | 0.28 | -1.01 | -0.21 | 375 | 2.25E-01 | 4.00E-01 | 0.28 | 174 | 2.31E-01 | 3.73E-01 | -0.80 | 472 | 9.59E-06 | 5.49E-05 | -0.77 | 531 | 4.87E-05 | 3.52E-03 | -1.01 | 442 | 5.45E-05 | 3.14E-04 |
| ILMN_1862198 | ##_HS.495112 | 0 | -0.54 | 6.19E-04 | 5.69E-03 | 9.16E-01 | 8.35E-06 | 1.29 | 0.04 | -1.25 | -0.83 | 534 | 3.84E-06 | 1.00E-04 | 0.04 | 259 | 8.61E-01 | 9.16E-01 | -0.40 | 393 | 2.33E-02 | 4.74E-02 | -0.19 | 399 | 3.15E-01 | 5.56E-01 | -1.25 | 527 | 7.95E-07 | 8.35E-06 |
| ILMN_1755974 | ALDOC | 17q11.2a | -0.51 | 2.66E-09 | 8.27E-07 | 8.51E-02 | 7.51E-17 | 1.29 | 0.27 | -1.03 | -0.55 | 476 | 1.34E-11 | 8.77E-09 | 0.27 | 183 | 8.82E-03 | 3.13E-02 | -0.77 | 466 | 7.98E-20 | 7.51E-17 | -0.20 | 408 | 1.31E-02 | 8.51E-02 | -1.03 | 454 | 2.15E-18 | 3.59E-15 |
| ILMN_1715963 | FBXO7 | 22q12.3a | -0.58 | 1.10E-04 | 1.70E-03 | 6.76E-01 | 1.55E-06 | 1.29 | 0.13 | -1.16 | -0.25 | 386 | 1.17E-01 | 2.59E-01 | 0.13 | 235 | 5.42E-01 | 6.76E-01 | -0.87 | 482 | 1.61E-07 | 1.55E-06 | -0.71 | 522 | 3.95E-05 | 3.20E-03 | -1.16 | 513 | 5.34E-07 | 5.95E-06 |
| ILMN_2375484 | CPEB2 | 4p15.33a | 0.38 | 3.20E-03 | 1.84E-02 | 6.01E-01 | 1.42E-07 | 1.29 | 1.15 | -0.14 | 0.59 | 134 | 2.67E-05 | 4.66E-04 | -0.14 | 327 | 4.54E-01 | 6.01E-01 | 0.28 | 240 | 4.60E-02 | 8.33E-02 | 0.14 | 215 | 3.36E-01 | 5.74E-01 | 1.15 | 129 | 6.63E-09 | 1.42E-07 |
| ILMN_1662340 | ZNF358 | 19p13.2e | -0.45 | 2.03E-06 | 9.21E-05 | 3.24E-01 | 2.17E-12 | 1.29 | 0.17 | -1.12 | -0.44 | 445 | 6.18E-06 | 1.47E-04 | 0.17 | 216 | 1.91E-01 | 3.24E-01 | -0.66 | 450 | 5.88E-11 | 1.66E-09 | -0.19 | 400 | 6.44E-02 | 2.21E-01 | -1.12 | 500 | 6.64E-15 | 2.17E-12 |
| ILMN_1680772 | ##_LOC642083 | 0 | -0.73 | 2.00E-03 | 1.32E-02 | 6.28E-01 | 5.34E-04 | 1.29 | -0.26 | -1.55 | -0.65 | 504 | 2.11E-02 | 7.69E-02 | -0.26 | 371 | 4.85E-01 | 6.28E-01 | -0.98 | 495 | 5.64E-04 | 1.91E-03 | -0.51 | 487 | 8.24E-02 | 2.55E-01 | -1.55 | 550 | 1.00E-04 | 5.34E-04 |
| ILMN_2121316 | GORAB | 1q24.2c | 0.52 | 4.59E-06 | 1.70E-04 | 1.74E-01 | 1.35E-14 | 1.29 | 1.03 | -0.26 | 0.45 | 192 | 7.56E-05 | 1.06E-03 | -0.26 | 372 | 8.26E-02 | 1.74E-01 | 1.03 | 81 | 5.02E-17 | 1.35E-14 | 0.25 | 165 | 4.03E-02 | 1.68E-01 | 0.54 | 288 | 7.48E-04 | 3.02E-03 |
| ILMN_1724299 | ANKRD20A11P | 21q11.2b-q11.2c | -0.03 | 8.93E-01 | 9.40E-01 | 9.34E-01 | 5.07E-03 | 1.29 | 1.09 | -0.19 | -0.13 | 355 | 6.05E-01 | 7.53E-01 | 1.09 | 57 | 8.65E-04 | 5.07E-03 | -0.09 | 324 | 7.05E-01 | 7.73E-01 | 0.05 | 270 | 8.55E-01 | 9.34E-01 | -0.19 | 360 | 5.76E-01 | 6.83E-01 |
| ILMN_2114720 | SLPI | 20q13.12b | -0.40 | 5.46E-02 | 1.42E-01 | 9.18E-01 | 1.61E-04 | 1.28 | 0.27 | -1.01 | -0.11 | 353 | 6.46E-01 | 7.82E-01 | 0.27 | 177 | 3.83E-01 | 5.34E-01 | -1.01 | 501 | 3.32E-05 | 1.61E-04 | -0.42 | 472 | 9.90E-02 | 2.85E-01 | 0.05 | 339 | 8.79E-01 | 9.18E-01 |
| ILMN_1683178 | JAK2 | 9p24.1c | 0.52 | 4.98E-08 | 6.36E-06 | 1.70E-01 | 9.14E-13 | 1.28 | 1.06 | -0.22 | 0.65 | 118 | 4.07E-12 | 3.83E-09 | -0.22 | 352 | 6.24E-02 | 1.40E-01 | 0.67 | 147 | 1.29E-12 | 6.17E-11 | 0.19 | 181 | 4.13E-02 | 1.70E-01 | 1.06 | 183 | 1.94E-15 | 9.14E-13 |

### Online Supplemental Table S4 (Continued)

| | | | S1 | | | M1 through M5 | | | | | M1 | | | | M2 | | | | M3 | | | | M4 | | | | M5 | | | |
|---|---|---|---|---|---|---|---|---|---|---|---|---|---|---|---|---|---|---|---|---|---|---|---|---|---|---|---|---|---|---|
| ID | Gene_Symbol | Cytoband | FC | $P_{val}$ | $P_{adj}$ | Max $P_{adj}$ | Min $P_{adj}$ | ΔFC | Max FC | Min FC | FC | FC Rank | $P_{val}$ | $P_{adj}$ | FC | FC Rank | $P_{val}$ | $P_{adj}$ | FC | FC Rank | $P_{val}$ | $P_{adj}$ | FC | FC Rank | $P_{val}$ | $P_{adj}$ | FC | FC Rank | $P_{val}$ | $P_{adj}$ |
| ILMN_1808238 | RBPMS2 | 15q22.31b | -0.98 | 2.25E-07 | 1.82E-05 | 6.95E-01 | 1.22E-09 | 1.28 | -0.16 | -1.44 | -0.81 | 532 | 1.02E-04 | 1.34E-03 | -0.16 | 333 | 5.64E-01 | 6.95E-01 | -1.44 | 560 | 4.13E-11 | 1.22E-09 | -0.86 | 540 | 1.01E-04 | 5.35E-03 | -1.15 | 510 | 8.52E-05 | 4.64E-04 |
| ILMN_1774211 | TMPPE | 3p22.3c | 0.08 | 5.81E-01 | 7.20E-01 | 9.97E-01 | 2.02E-05 | 1.28 | 1.08 | -0.20 | 0.26 | 273 | 1.01E-01 | 2.34E-01 | 0.00 | 278 | 9.95E-01 | 9.97E-01 | -0.16 | 340 | 3.18E-01 | 4.11E-01 | -0.20 | 405 | 2.45E-01 | 4.83E-01 | 1.08 | 168 | 2.24E-06 | 2.02E-05 |
| ILMN_2404850 | RPL14 | 3p22.1c | -0.24 | 1.93E-01 | 3.41E-01 | 7.12E-01 | 1.06E-03 | 1.28 | 0.17 | -1.10 | -0.56 | 481 | 7.93E-03 | 3.76E-02 | 0.15 | 221 | 5.85E-01 | 7.12E-01 | -0.13 | 334 | 5.42E-01 | 6.32E-01 | 0.17 | 196 | 4.35E-01 | 6.66E-01 | -1.10 | 492 | 2.21E-04 | 1.06E-03 |
| ILMN_1804601 | ##_LOC649923 | 0 | -0.61 | 4.77E-02 | 1.29E-01 | 7.69E-01 | 1.85E-02 | 1.28 | 0.21 | -1.07 | -1.07 | 550 | 3.02E-03 | 1.85E-02 | 0.21 | 199 | 6.55E-01 | 7.69E-01 | -0.22 | 346 | 5.41E-01 | 6.31E-01 | 0.65 | 508 | 8.70E-02 | 2.62E-01 | -0.62 | 400 | 2.16E-01 | 3.28E-01 |
| ILMN_1806448 | NTNG2 | 9q34.13b | 0.69 | 3.75E-05 | 7.80E-04 | 9.87E-02 | 3.35E-08 | 1.28 | 1.63 | 0.35 | 0.91 | 75 | 1.54E-06 | 4.87E-05 | 0.75 | 103 | 2.15E-03 | 1.05E-02 | 0.35 | 213 | 5.62E-02 | 9.87E-02 | 0.48 | 92 | 1.49E-02 | 9.19E-02 | 1.63 | 28 | 1.17E-09 | 3.35E-08 |
| ILMN_1690101 | FAIM | 3q22.3c | -0.20 | 1.57E-01 | 2.97E-01 | 3.89E-01 | 2.91E-05 | 1.28 | 0.27 | -1.01 | -0.35 | 421 | 2.15E-02 | 7.78E-02 | -1.01 | 502 | 1.15E-06 | 2.91E-05 | 0.27 | 243 | 7.90E-02 | 1.31E-01 | -0.22 | 419 | 1.68E-01 | 3.89E-01 | -0.38 | 381 | 7.67E-02 | 1.44E-01 |
| ILMN_1783702 | MORC3 | 21q22.12b | 0.35 | 6.08E-04 | 5.63E-03 | 6.31E-01 | 2.71E-09 | 1.27 | 1.04 | -0.23 | 0.44 | 197 | 6.03E-05 | 8.85E-04 | -0.23 | 359 | 1.02E-01 | 2.04E-01 | 0.45 | 189 | 5.25E-05 | 2.41E-04 | 0.10 | 236 | 3.95E-01 | 6.31E-01 | 1.04 | 196 | 5.81E-11 | 2.71E-09 |
| ILMN_1657932 | MUC6 | 0 | -0.91 | 4.12E-04 | 4.21E-03 | 9.05E-01 | 3.91E-05 | 1.27 | -0.07 | -1.35 | -0.59 | 488 | 4.42E-02 | 1.30E-01 | -0.07 | 305 | 8.45E-01 | 9.05E-01 | -1.35 | 555 | 6.49E-06 | 3.91E-05 | -1.27 | 562 | 4.81E-05 | 3.50E-03 | -0.71 | 405 | 8.20E-02 | 1.52E-01 |
| ILMN_1786847 | TGM3 | 20p13d | -0.76 | 2.68E-04 | 3.10E-03 | 5.64E-01 | 3.38E-09 | 1.27 | -0.24 | -1.51 | -0.33 | 409 | 1.43E-01 | 2.97E-01 | -0.24 | 361 | 4.14E-01 | 5.64E-01 | -1.51 | 563 | 1.34E-10 | 3.38E-09 | -0.73 | 525 | 2.35E-03 | 3.04E-02 | -0.49 | 387 | 1.22E-01 | 2.09E-01 |
| ILMN_1704629 | SLC1A7 | 1p32.3c | 0.29 | 1.29E-01 | 2.59E-01 | 5.07E-01 | 4.28E-03 | 1.27 | 1.00 | -0.27 | 0.28 | 267 | 2.14E-01 | 3.87E-01 | 1.00 | 84 | 6.98E-04 | 4.28E-03 | 0.20 | 254 | 3.63E-01 | 4.58E-01 | 0.45 | 98 | 5.37E-02 | 1.99E-01 | -0.27 | 372 | 3.83E-01 | 5.07E-01 |
| ILMN_1857136 | ##_HS.567124 | 0 | 0.27 | 6.32E-02 | 1.57E-01 | 9.60E-01 | 5.90E-07 | 1.27 | 1.28 | 0.01 | 0.42 | 209 | 9.92E-03 | 4.44E-02 | 0.19 | 202 | 3.57E-01 | 5.08E-01 | 0.01 | 294 | 9.43E-01 | 9.60E-01 | 0.08 | 249 | 6.48E-01 | 8.17E-01 | 1.28 | 83 | 3.50E-08 | 5.90E-07 |
| ILMN_1676555 | IFT56 | 7q34b | 0.28 | 1.20E-01 | 2.47E-01 | 9.69E-01 | 1.11E-03 | 1.27 | 1.07 | -0.20 | 0.47 | 182 | 2.17E-02 | 7.82E-02 | -0.20 | 345 | 4.60E-01 | 6.07E-01 | -0.01 | 305 | 955E-01 | 9.69E-01 | 0.23 | 170 | 2.97E-01 | 5.37E-01 | 1.07 | 173 | 2.34E-04 | 1.11E-03 |
| ILMN_1801254 | ##_LOC648984 | 0 | 0.28 | 2.99E-02 | 9.24E-02 | 8.87E-01 | 3.73E-06 | 1.27 | 1.05 | -0.22 | 0.47 | 185 | 1.14E-03 | 8.67E-03 | -0.22 | 351 | 2.44E-01 | 3.89E-01 | 0.19 | 257 | 1.80E-01 | 2.58E-01 | -0.04 | 331 | 7.66E-01 | 8.87E-01 | 1.05 | 191 | 3.11E-07 | 3.73E-06 |
| ILMN_1700248 | WDR86 | 7q36.1d | -0.55 | 3.20E-03 | 1.84E-02 | 9.48E-01 | 2.05E-04 | 1.27 | 0.03 | -1.24 | -0.77 | 528 | 2.73E-04 | 2.83E-03 | 0.03 | 261 | 9.11E-01 | 9.48E-01 | -0.69 | 455 | 1.16E-03 | 3.59E-03 | -0.14 | 376 | 5.29E-01 | 7.37E-01 | -1.24 | 526 | 3.31E-05 | 2.05E-04 |
| ILMN_1652417 | ZNG1E | 0 | 0.66 | 4.46E-06 | 1.67E-04 | 7.37E-01 | 2.53E-06 | 1.27 | 1.16 | -0.10 | 0.76 | 95 | 2.15E-06 | 6.27E-05 | -0.10 | 319 | 6.14E-01 | 7.37E-01 | 0.83 | 111 | 2.82E-07 | 2.53E-06 | 0.37 | 118 | 2.59E-02 | 1.30E-01 | 1.16 | 127 | 2.78E-07 | 3.37E-06 |
| ILMN_1681603 | UBE2A | Xq24c | 0.24 | 6.09E-02 | 1.53E-01 | 9.69E-01 | 3.76E-06 | 1.27 | 1.03 | -0.24 | 0.30 | 258 | 3.18E-02 | 1.04E-01 | -0.24 | 360 | 1.96E-01 | 3.31E-01 | 0.27 | 242 | 5.18E-02 | 9.23E-02 | 0.01 | 295 | 9.31E-01 | 9.69E-01 | 1.03 | 211 | 3.14E-07 | 3.76E-06 |
| ILMN_1791955 | RASGRP4 | 19q13.2a | 0.12 | 3.73E-01 | 5.35E-01 | 6.19E-01 | 1.03E-06 | 1.27 | 1.13 | -0.13 | 0.26 | 272 | 7.13E-02 | 1.84E-01 | 0.14 | 228 | 4.73E-01 | 6.19E-01 | -0.10 | 325 | 4.89E-01 | 5.83E-01 | -0.13 | 375 | 3.79E-01 | 6.17E-01 | 1.13 | 143 | 6.74E-08 | 1.03E-06 |
| ILMN_1665557 | USP15 | 12q14.1d | 0.34 | 1.01E-04 | 1.59E-03 | 6.64E-01 | 2.09E-12 | 1.26 | 1.01 | -0.25 | 0.45 | 194 | 5.28E-07 | 2.20E-05 | -0.25 | 365 | 2.88E-02 | 7.77E-02 | 0.42 | 198 | 2.54E-06 | 1.72E-05 | 0.07 | 253 | 4.33E-01 | 6.64E-01 | 1.01 | 222 | 5.96E-15 | 2.09E-12 |
| ILMN_2302757 | FCGBP | 19q13.2b | -0.76 | 3.54E-05 | 7.56E-04 | 4.51E-01 | 3.73E-06 | 1.26 | -0.27 | -1.53 | -0.99 | 544 | 3.59E-06 | 9.46E-05 | -0.53 | 439 | 5.43E-02 | 1.26E-01 | -0.83 | 479 | 8.67E-05 | 3.75E-04 | -0.27 | 429 | 2.16E-01 | 4.51E-01 | -1.53 | 549 | 3.11E-07 | 3.73E-06 |
| ILMN_1668984 | SPNS3 | 17p13.2c | -0.36 | 2.87E-02 | 8.98E-02 | 9.41E-01 | 3.98E-05 | 1.26 | 0.03 | -1.23 | -0.40 | 428 | 3.27E-02 | 1.06E-01 | 0.03 | 262 | 9.02E-01 | 9.41E-01 | -0.27 | 359 | 1.55E-01 | 2.29E-01 | -0.21 | 413 | 2.88E-01 | 5.29E-01 | -1.23 | 522 | 4.90E-06 | 3.98E-05 |
| ILMN_1679280 | PPIAP66 | 2q31.1f | -0.24 | 6.69E-02 | 1.64E-01 | 9.42E-01 | 8.65E-08 | 1.26 | 0.02 | -1.23 | -0.34 | 412 | 2.00E-02 | 7.39E-02 | 0.02 | 266 | 9.04E-01 | 9.42E-01 | -0.07 | 317 | 6.35E-01 | 7.14E-01 | -0.05 | 333 | 7.56E-01 | 8.83E-01 | -1.23 | 525 | 3.66E-09 | 8.65E-08 |
| ILMN_1772387 | TLR2 | 4q31.3d | 0.16 | 1.96E-01 | 3.44E-01 | 9.50E-01 | 4.05E-07 | 1.25 | 1.04 | -0.21 | 0.41 | 211 | 1.60E-03 | 1.13E-02 | 0.02 | 269 | 9.16E-01 | 9.50E-01 | -0.04 | 312 | 7.79E-01 | 8.34E-01 | -0.21 | 416 | 1.15E-01 | 3.11E-01 | 1.04 | 198 | 2.24E-08 | 4.05E-07 |
| ILMN_2094875 | ABCB1 | 7q21.12a-q21.12b | -0.21 | 1.67E-01 | 3.09E-01 | 9.48E-01 | 1.09E-05 | 1.25 | 0.04 | -1.21 | -0.42 | 437 | 1.57E-02 | 6.15E-02 | -1.21 | 541 | 3.18E-07 | 1.09E-05 | -0.02 | 310 | 8.96E-01 | 9.25E-01 | 0.03 | 283 | 8.83E-01 | 9.48E-01 | 0.04 | 341 | 8.58E-01 | 9.04E-01 |
| ILMN_1781700 | IL18R1 | 2q12.1a | 0.18 | 6.35E-02 | 1.57E-01 | 7.88E-01 | 5.35E-09 | 1.25 | 1.05 | -0.20 | 0.29 | 265 | 1.07E-02 | 4.67E-02 | -0.20 | 346 | 1.72E-01 | 3.01E-01 | 0.04 | 287 | 7.23E-01 | 7.88E-01 | 0.09 | 239 | 4.43E-01 | 6.72E-01 | 1.05 | 190 | 1.33E-10 | 5.35E-09 |
| ILMN_2112915 | DRD4 | 11p15.5d | -0.12 | 4.17E-01 | 5.78E-01 | 8.00E-01 | 3.46E-05 | 1.25 | 0.15 | -1.09 | -0.12 | 354 | 4.91E-01 | 6.63E-01 | -1.09 | 522 | 1.46E-06 | 3.46E-05 | 0.15 | 262 | 3.60E-01 | 4.55E-01 | -0.27 | 428 | 1.31E-01 | 3.35E-01 | -0.09 | 353 | 7.16E-01 | 8.00E-01 |
| ILMN_1770454 | AGRN | 1p36.33b | 1.01 | 1.93E-06 | 8.88E-05 | 5.57E-01 | 7.38E-06 | 1.25 | 1.51 | 0.26 | 1.12 | 47 | 4.36E-06 | 1.11E-04 | 0.26 | 185 | 4.06E-01 | 5.57E-01 | 1.21 | 36 | 9.54E-07 | 7.38E-06 | 0.59 | 77 | 2.00E-02 | 1.10E-01 | 1.51 | 43 | 1.01E-05 | 7.37E-05 |
| ILMN_1781966 | OSBP2 | 22q12.2b-q12.2c | -0.41 | 7.14E-02 | 1.71E-01 | 9.96E-01 | 6.50E-03 | 1.25 | 0.16 | -1.08 | 0.00 | 336 | 9.92E-01 | 9.96E-01 | 0.16 | 218 | 6.30E-01 | 7.49E-01 | -0.79 | 471 | 2.30E-03 | 6.50E-03 | -0.54 | 491 | 4.59E-02 | 1.81E-01 | -1.08 | 483 | 2.86E-03 | 9.36E-03 |
| ILMN_1681721 | OASL | 12q24.31a | 0.70 | 2.26E-02 | 7.56E-02 | 5.99E-01 | 4.77E-03 | 1.24 | 1.59 | 0.35 | 1.10 | 49 | 2.29E-03 | 1.50E-02 | 1.59 | 9 | 8.01E-04 | 4.77E-03 | 0.35 | 214 | 3.31E-01 | 4.25E-01 | 0.49 | 91 | 1.96E-01 | 4.24E-01 | 0.35 | 308 | 4.83E-01 | 5.99E-01 |
| ILMN_1667830 | ANKRD13D | 11q13.1f | 0.31 | 3.95E-02 | 1.13E-01 | 8.32E-01 | 1.53E-06 | 1.24 | 1.29 | 0.05 | 0.26 | 277 | 1.24E-01 | 2.69E-01 | 0.27 | 181 | 2.21E-01 | 3.62E-01 | 0.05 | 284 | 7.78E-01 | 8.32E-01 | 0.44 | 102 | 1.28E-02 | 8.39E-02 | 1.29 | 80 | 1.09E-07 | 1.53E-06 |
| ILMN_1815184 | ASPM | 1q31.3c | 0.59 | 2.13E-04 | 2.67E-03 | 6.22E-01 | 2.24E-08 | 1.23 | 1.07 | -0.16 | 0.57 | 142 | 8.10E-04 | 6.68E-03 | -0.16 | 334 | 4.77E-01 | 6.22E-01 | 1.07 | 64 | 1.20E-09 | 2.24E-08 | 0.18 | 190 | 3.06E-01 | 5.46E-01 | 0.74 | 262 | 1.73E-03 | 6.14E-03 |
| ILMN_1789599 | MICOS10-NBL1/NBL1 | 1p36.13a | -0.40 | 3.93E-02 | 1.13E-01 | 8.85E-01 | 2.17E-04 | 1.23 | -0.07 | -1.30 | -0.66 | 506 | 3.57E-03 | 2.08E-02 | -1.30 | 548 | 1.54E-05 | 2.17E-04 | -0.16 | 341 | 4.62E-01 | 5.57E-01 | -0.07 | 347 | 7.60E-01 | 8.85E-01 | -0.57 | 395 | 6.76E-02 | 1.30E-01 |
| ILMN_2166457 | HPGD | 4q34.1d | -0.40 | 1.66E-02 | 6.05E-02 | 7.52E-01 | 2.90E-04 | 1.23 | 0.12 | -1.10 | -0.45 | 446 | 2.20E-02 | 7.92E-02 | -1.10 | 523 | 2.21E-05 | 2.90E-04 | -0.33 | 375 | 9.49E-02 | 1.52E-01 | -0.35 | 453 | 9.26E-02 | 2.73E-01 | 0.12 | 330 | 6.56E-01 | 7.52E-01 |
| ILMN_1798557 | FAM153A/FAM153B | 5q35.2d | -0.15 | 4.16E-01 | 5.78E-01 | 7.82E-01 | 1.91E-03 | 1.22 | 0.22 | -1.01 | -0.51 | 463 | 1.31E-02 | 5.42E-02 | -1.01 | 503 | 2.47E-04 | 1.91E-03 | 0.22 | 251 | 2.94E-01 | 3.87E-01 | 0.15 | 207 | 4.78E-01 | 6.99E-01 | -0.11 | 355 | 6.94E-01 | 7.82E-01 |
| ILMN_1716468 | NUDT16L2P | 3q22.1b | 0.39 | 5.39E-04 | 5.17E-03 | 9.32E-01 | 3.35E-10 | 1.22 | 1.20 | -0.02 | 0.63 | 124 | 1.90E-07 | 1.04E-05 | 0.13 | 231 | 3.93E-01 | 5.44E-01 | 0.32 | 220 | 6.30E-03 | 1.55E-02 | -0.02 | 317 | 8.52E-01 | 9.32E-01 | 1.20 | 114 | 4.14E-12 | 3.35E-10 |
| ILMN_1794863 | CAMK2N1 | 1p36.12b | -0.58 | 1.94E-03 | 1.29E-02 | 3.25E-01 | 4.11E-06 | 1.22 | -0.35 | -1.57 | -0.72 | 516 | 1.00E-03 | 7.89E-03 | -1.57 | 562 | 8.46E-08 | 4.11E-06 | -0.44 | 401 | 4.07E-02 | 7.54E-02 | -0.35 | 456 | 1.25E-01 | 3.25E-01 | -0.88 | 417 | 3.96E-03 | 1.23E-02 |
| ILMN_2352380 | TRPM2 | 21q22.3d | 0.18 | 2.27E-01 | 3.81E-01 | 6.34E-01 | 3.15E-05 | 1.22 | 1.07 | -0.15 | 0.32 | 248 | 5.40E-02 | 1.50E-01 | 1.07 | 61 | 1.29E-06 | 3.15E-05 | -0.15 | 338 | 3.73E-01 | 4.69E-01 | 0.25 | 162 | 1.52E-01 | 3.65E-01 | -0.15 | 356 | 5.21E-01 | 6.34E-01 |
| ILMN_1785345 | GPR84 | 12q13.13f | 0.37 | 2.40E-02 | 7.90E-02 | 9.78E-01 | 4.73E-05 | 1.22 | 1.21 | -0.01 | 0.61 | 126 | 1.13E-03 | 8.56E-03 | 0.07 | 253 | 7.89E-01 | 8.67E-01 | 0.20 | 255 | 2.80E-01 | 3.72E-01 | -0.01 | 307 | 9.51E-01 | 9.78E-01 | 1.21 | 111 | 6.01E-06 | 4.73E-05 |
| ILMN_1759772 | LRRC56 | 11p15.5d | -0.42 | 2.31E-03 | 1.46E-02 | 8.98E-01 | 1.30E-06 | 1.21 | 0.04 | -1.17 | -0.66 | 507 | 1.65E-05 | 3.15E-04 | -0.43 | 415 | 3.22E-02 | 8.45E-02 | -0.37 | 384 | 1.49E-02 | 3.23E-02 | 0.04 | 276 | 7.84E-01 | 8.98E-01 | -1.17 | 515 | 8.87E-08 | 1.30E-06 |
| ILMN_1732404 | C2orf15 | 2q11.2c | -0.27 | 5.38E-02 | 1.40E-01 | 8.40E-01 | 4.88E-06 | 1.21 | 0.06 | -1.14 | -0.45 | 447 | 5.08E-03 | 2.69E-02 | -1.14 | 530 | 1.11E-07 | 4.88E-06 | 0.05 | 286 | 7.73E-01 | 8.28E-01 | -0.25 | 422 | 1.40E-01 | 3.49E-01 | 0.06 | 338 | 7.71E-01 | 8.40E-01 |
| ILMN_1668228 | ##_LOC136143 | 7q31.31d | -0.42 | 5.28E-06 | 1.84E-04 | 6.53E-01 | 3.44E-12 | 1.21 | 0.14 | -1.07 | -0.56 | 482 | 4.41E-09 | 5.81E-07 | 0.14 | 227 | 2.52E-01 | 3.98E-01 | -0.49 | 415 | 2.27E-07 | 2.10E-06 | -0.08 | 349 | 4.19E-01 | 6.53E-01 | -1.07 | 474 | 1.14E-14 | 3.44E-12 |
| ILMN_2388800 | PLPP3 | 1p32.2c | 0.34 | 9.27E-03 | 3.94E-02 | 5.85E-01 | 7.18E-06 | 1.21 | 1.05 | -0.15 | 0.40 | 216 | 7.09E-03 | 3.45E-02 | -0.15 | 330 | 4.35E-01 | 5.85E-01 | 0.29 | 228 | 4.81E-02 | 8.66E-02 | 0.15 | 208 | 3.29E-01 | 5.68E-01 | 1.05 | 187 | 6.68E-07 | 7.18E-06 |
| ILMN_1790781 | DHRS13 | 17q11.2a | -0.48 | 8.98E-04 | 7.42E-03 | 4.85E-01 | 8.58E-09 | 1.20 | 0.20 | -1.01 | -0.27 | 390 | 8.01E-02 | 1.99E-01 | 0.20 | 201 | 3.34E-01 | 4.85E-01 | -1.01 | 502 | 3.91E-10 | 8.58E-09 | -0.34 | 449 | 3.96E-02 | 1.66E-01 | -0.26 | 370 | 2.34E-01 | 3.49E-01 |
| ILMN_1774890 | LAS1L | Xq11.1c | -0.20 | 7.41E-02 | 1.76E-01 | 5.28E-01 | 8.68E-08 | 1.20 | 0.18 | -1.03 | -0.37 | 424 | 2.33E-03 | 1.52E-02 | 0.14 | 226 | 3.76E-01 | 5.28E-01 | -0.20 | 345 | 9.41E-02 | 1.51E-01 | 0.18 | 194 | 1.60E-01 | 3.77E-01 | -1.03 | 453 | 3.69E-09 | 8.68E-08 |
| ILMN_2276952 | TSC22D3 | Xq22.3b | -0.58 | 1.32E-05 | 3.66E-04 | 5.04E-01 | 1.00E-10 | 1.20 | 0.17 | -1.03 | -0.41 | 431 | 2.99E-03 | 1.83E-02 | 0.17 | 213 | 3.53E-01 | 5.04E-01 | -1.03 | 507 | 2.30E-12 | 1.00E-10 | -0.51 | 486 | 5.55E-04 | 1.36E-02 | -0.21 | 363 | 2.73E-01 | 3.93E-01 |
| ILMN_1904578 | TENM1 | 0 | 0.04 | 8.03E-01 | 8.83E-01 | 9.77E-01 | 2.39E-03 | 1.19 | 1.03 | -0.16 | 0.02 | 332 | 9.43E-01 | 9.69E-01 | -0.16 | 337 | 5.53E-01 | 6.86E-01 | 0.01 | 295 | 9.67E-01 | 9.77E-01 | -0.02 | 316 | 9.23E-01 | 9.65E-01 | 1.03 | 209 | 5.72E-04 | 2.39E-03 |
| ILMN_1658798 | MEGF9 | 9q33.2a | 0.25 | 2.12E-02 | 7.20E-02 | 9.83E-01 | 3.27E-10 | 1.19 | 1.16 | -0.03 | 0.49 | 173 | 2.05E-05 | 3.78E-04 | 0.10 | 247 | 5.22E-01 | 6.59E-01 | -0.03 | 311 | 7.75E-01 | 8.29E-01 | 0.01 | 300 | 9.61E-01 | 9.83E-01 | 1.16 | 128 | 3.98E-12 | 3.27E-10 |
| ILMN_1699809 | CAPNS2 | 16q12.2c | 0.37 | 2.99E-03 | 1.76E-02 | 9.39E-01 | 8.18E-08 | 1.19 | 1.17 | -0.02 | 0.53 | 155 | 1.17E-04 | 1.48E-03 | -0.02 | 287 | 9.00E-01 | 9.39E-01 | 0.23 | 249 | 9.43E-02 | 1.51E-01 | 0.18 | 191 | 2.07E-01 | 4.39E-01 | 1.17 | 124 | 3.42E-09 | 8.18E-08 |
| ILMN_1657373 | P3H2 | 3q28b-q28c | -0.51 | 9.69E-06 | 2.93E-04 | 6.04E-01 | 4.16E-07 | 1.19 | 0.13 | -1.06 | -0.55 | 475 | 4.15E-05 | 6.55E-04 | 0.13 | 236 | 4.56E-01 | 6.04E-01 | -0.54 | 425 | 4.67E-05 | 2.18E-04 | -0.41 | 471 | 2.94E-03 | 3.42E-02 | -1.06 | 469 | 2.31E-08 | 4.16E-07 |
| ILMN_1796560 | PFKFB2 | 1q32.1h-q32.2a | 0.15 | 2.17E-01 | 3.70E-01 | 9.30E-01 | 2.60E-07 | 1.18 | 1.17 | -0.02 | 0.22 | 291 | 1.26E-01 | 2.71E-01 | 0.19 | 206 | 3.03E-01 | 4.53E-01 | -0.02 | 307 | 9.03E-01 | 9.30E-01 | 0.05 | 269 | 7.47E-01 | 8.78E-01 | 1.17 | 121 | 1.36E-08 | 2.60E-07 |
| ILMN_1716733 | MYOM2 | 8p23.3a | -1.07 | 3.37E-03 | 1.91E-02 | 2.76E-01 | 4.03E-03 | 1.18 | -0.81 | -1.99 | -0.89 | 538 | 3.99E-02 | 1.21E-01 | -0.81 | 488 | 1.53E-01 | 2.76E-01 | -1.32 | 550 | 2.44E-03 | 6.83E-03 | -0.91 | 546 | 4.67E-02 | 1.83E-01 | -1.99 | 562 | 1.05E-03 | 4.03E-03 |
| ILMN_2380163 | PTPRF | 1p34.2a | -0.47 | 3.95E-04 | 4.09E-03 | 7.16E-01 | 3.83E-11 | 1.18 | 0.18 | -1.00 | -0.08 | 344 | 5.56E-01 | 7.16E-01 | 0.18 | 210 | 2.97E-01 | 4.47E-01 | -1.00 | 499 | 7.06E-13 | 3.83E-11 | -0.59 | 501 | 2.79E-05 | 2.67E-03 | -0.32 | 375 | 8.23E-02 | 1.52E-01 |
| ILMN_1798804 | SRPK1 | 6p21.31b | 0.14 | 1.14E-01 | 2.38E-01 | 9.83E-01 | 3.62E-13 | 1.18 | 1.05 | -0.13 | 0.25 | 279 | 3.17E-03 | 1.91E-02 | -0.13 | 325 | 2.42E-01 | 3.86E-01 | 0.00 | 301 | 9.76E-01 | 9.83E-01 | -0.06 | 340 | 5.05E-01 | 7.18E-01 | 1.05 | 193 | 5.05E-16 | 3.62E-13 |
| ILMN_1730964 | CLEC18A/CLEC18C | 16q22.1f | 0.17 | 2.17E-01 | 3.71E-01 | 7.86E-01 | 6.37E-06 | 1.18 | 1.09 | -0.08 | 0.21 | 293 | 1.63E-01 | 3.24E-01 | 0.19 | 204 | 3.30E-01 | 4.82E-01 | 0.09 | 274 | 5.60E-01 | 6.48E-01 | -0.08 | 351 | 6.00E-01 | 7.86E-01 | 1.09 | 156 | 5.78E-07 | 6.37E-06 |
| ILMN_2384056 | GPER1 | 7p22.3b | 0.10 | 6.92E-01 | 8.04E-01 | 7.96E-01 | 3.18E-02 | 1.18 | 1.02 | -0.16 | 0.21 | 296 | 4.67E-01 | 6.42E-01 | 0.15 | 222 | 6.91E-01 | 7.96E-01 | -0.14 | 336 | 6.19E-01 | 7.01E-01 | -0.16 | 388 | 6.03E-01 | 7.88E-01 | 1.02 | 217 | 1.22E-02 | 3.18E-02 |
| ILMN_1785252 | SLC26A6 | 3p21.31e | 0.17 | 9.43E-02 | 2.08E-01 | 9.97E-01 | 2.57E-09 | 1.17 | 1.02 | -0.15 | 0.15 | 307 | 1.60E-01 | 3.20E-01 | 1.02 | 74 | 5.46E-12 | 2.57E-09 | -0.15 | 337 | 1.73E-01 | 2.50E-01 | 0.42 | 106 | 2.24E-04 | 8.16E-03 | 0.00 | 346 | 9.96E-01 | 9.97E-01 |
| ILMN_1654692 | RALGPS2 | 1q25.2b | 0.69 | 1.53E-04 | 2.12E-03 | 8.54E-01 | 3.29E-06 | 1.17 | 1.09 | -0.08 | 0.62 | 125 | 3.10E-03 | 1.88E-02 | -0.08 | 310 | 7.70E-01 | 8.54E-01 | 1.09 | 59 | 3.81E-07 | 3.29E-06 | 0.46 | 96 | 3.74E-02 | 1.60E-01 | 0.87 | 247 | 3.09E-03 | 9.98E-03 |
| ILMN_1823172 | IL23A | 0 | -0.47 | 1.37E-03 | 1.01E-02 | 8.41E-01 | 1.75E-06 | 1.17 | -0.07 | -1.23 | -0.78 | 529 | 2.69E-06 | 7.48E-05 | -0.23 | 354 | 2.91E-01 | 4.40E-01 | -0.34 | 380 | 3.82E-02 | 7.15E-02 | -0.07 | 345 | 6.88E-01 | 8.41E-01 | -1.23 | 524 | 1.28E-07 | 1.75E-06 |
| ILMN_1788059 | PCGF5 | 10q23.32a | 0.73 | 2.34E-07 | 1.87E-05 | 9.19E-01 | 9.63E-10 | 1.17 | 1.13 | -0.03 | 0.89 | 77 | 4.23E-09 | 5.73E-07 | -0.03 | 292 | 8.66E-01 | 9.19E-01 | 1.02 | 86 | 3.13E-11 | 9.63E-10 | 0.28 | 144 | 7.02E-02 | 2.32E-01 | 1.13 | 138 | 7.25E-08 | 1.10E-06 |
| ILMN_1701914 | CD274 | 9p24.1c | 0.56 | 7.37E-04 | 6.46E-03 | 8.13E-01 | 2.76E-05 | 1.16 | 1.25 | 0.09 | 0.75 | 97 | 8.98E-05 | 1.21E-03 | 0.09 | 248 | 7.14E-01 | 8.13E-01 | 0.48 | 181 | 1.09E-02 | 2.47E-02 | 0.29 | 140 | 1.47E-01 | 3.58E-01 | 1.25 | 90 | 3.23E-06 | 2.76E-05 |
| ILMN_1808846 | TBKBP1 | 17q21.32b | 0.12 | 3.99E-01 | 5.61E-01 | 9.84E-01 | 1.87E-04 | 1.16 | 1.01 | -0.16 | 0.31 | 250 | 6.71E-02 | 1.75E-01 | 0.01 | 275 | 9.72E-01 | 9.84E-01 | -0.16 | 339 | 3.55E-01 | 4.51E-01 | -0.09 | 355 | 6.25E-01 | 8.01E-01 | 1.01 | 228 | 2.96E-05 | 1.87E-04 |
| ILMN_1686097 | TOP2A | 17q21.2a | 0.99 | 2.47E-07 | 1.94E-05 | 4.55E-01 | 1.83E-09 | 1.16 | 1.45 | 0.29 | 1.02 | 60 | 2.36E-06 | 6.70E-05 | 0.29 | 171 | 3.04E-01 | 4.55E-01 | 1.45 | 18 | 6.59E-11 | 1.83E-09 | 0.68 | 66 | 2.53E-03 | 3.13E-02 | 0.61 | 278 | 3.97E-02 | 8.49E-02 |
| ILMN_1661577 | MAFA | 8q24.3f | 0.09 | 5.98E-01 | 7.33E-01 | 9.95E-01 | 1.38E-03 | 1.16 | 1.04 | -0.12 | 0.00 | 335 | 9.90E-01 | 9.95E-01 | 1.04 | 73 | 1.62E-04 | 1.38E-03 | -0.12 | 331 | 5.54E-01 | 6.43E-01 | 0.25 | 159 | 2.50E-01 | 4.90E-01 | 0.11 | 332 | 6.93E-01 | 7.81E-01 |

### Online Supplemental Table S4 (Continued)

| | | | S1 | | | M1 through M5 | | | | | M1 | | | | M2 | | | | M3 | | | | M4 | | | | M5 | | | |
|---|---|---|---|---|---|---|---|---|---|---|---|---|---|---|---|---|---|---|---|---|---|---|---|---|---|---|---|---|---|---|
| ID | Gene_Symbol | Cytoband | FC | $P_{val}$ | $P_{adj}$ | Max $P_{adj}$ | Min $P_{adj}$ | ΔFC | Max FC | Min FC | FC | FC Rank | $P_{val}$ | $P_{adj}$ | FC | FC Rank | $P_{val}$ | $P_{adj}$ | FC | FC Rank | $P_{val}$ | $P_{adj}$ | FC | FC Rank | $P_{val}$ | $P_{adj}$ | FC | FC Rank | $P_{val}$ | $P_{adj}$ |
| ILMN_1880930 | ##_HS.543027 | 0 | 0.35 | 1.52E-02 | 5.66E-02 | 8.93E-01 | 2.74E-06 | 1.16 | 1.19 | 0.03 | 0.68 | 112 | 2.88E-05 | 4.96E-04 | 0.08 | 252 | 7.14E-01 | 8.13E-01 | 0.03 | 290 | 8.54E-01 | 8.93E-01 | 0.12 | 223 | 4.89E-01 | 7.06E-01 | 1.19 | 116 | 2.16E-07 | 2.74E-06 |
| ILMN_1806473 | BEX5 | Xq22.1d | -0.47 | 2.11E-03 | 1.37E-02 | 9.27E-01 | 7.16E-06 | 1.15 | -0.03 | -1.19 | -0.81 | 531 | 2.19E-06 | 6.33E-05 | -0.08 | 311 | 7.13E-01 | 8.12E-01 | -0.32 | 373 | 5.87E-02 | 1.02E-01 | -0.03 | 324 | 8.43E-01 | 9.27E-01 | -1.19 | 518 | 6.65E-07 | 7.16E-06 |
| ILMN_1724822 | LINC00999 | 10p11.21a | -0.53 | 6.20E-05 | 1.13E-03 | 6.14E-01 | 4.87E-10 | 1.15 | 0.13 | -1.01 | -0.35 | 419 | 1.55E-02 | 6.10E-02 | -0.33 | 393 | 7.98E-02 | 1.70E-01 | -1.01 | 503 | 1.42E-11 | 4.87E-10 | -0.54 | 490 | 3.75E-04 | 1.09E-02 | 0.13 | 328 | 4.99E-01 | 6.14E-01 |
| ILMN_1680424 | CTSG | 14q12a | 0.50 | 9.84E-02 | 2.14E-01 | 9.89E-01 | 9.55E-03 | 1.15 | 1.13 | -0.01 | 1.13 | 45 | 1.29E-03 | 9.55E-03 | 0.59 | 123 | 1.98E-01 | 3.34E-01 | 0.08 | 276 | 8.22E-01 | 8.68E-01 | 0.01 | 306 | 9.74E-01 | 9.89E-01 | 0.40 | 301 | 4.09E-01 | 5.31E-01 |
| ILMN_2058782 | IFI27 | 14q32.13a | 2.85 | 1.38E-06 | 7.01E-05 | 4.69E-02 | 6.85E-05 | 1.15 | 3.21 | 2.07 | 3.21 | 1 | 5.17E-06 | 1.27E-04 | 2.93 | 1 | 1.40E-03 | 7.44E-03 | 3.09 | 1 | 1.24E-05 | 6.85E-05 | 2.07 | 2 | 4.84E-03 | 4.65E-02 | 2.27 | 5 | 1.94E-02 | 4.69E-02 |
| ILMN_1757702 | TPT1P9 | 0 | -0.22 | 1.30E-01 | 2.60E-01 | 8.34E-01 | 1.63E-04 | 1.14 | 0.14 | -1.00 | -0.34 | 415 | 4.30E-02 | 1.28E-01 | 0.14 | 225 | 5.12E-01 | 6.50E-01 | 0.05 | 285 | 7.80E-01 | 8.34E-01 | -0.16 | 392 | 3.51E-01 | 5.90E-01 | -1.00 | 430 | 2.53E-05 | 1.63E-04 |
| ILMN_1663437 | NECAB2 | 16q23.3b | -0.50 | 5.09E-03 | 2.58E-02 | 9.34E-01 | 1.75E-06 | 1.14 | 0.10 | -1.04 | -0.11 | 352 | 5.72E-01 | 7.28E-01 | 0.03 | 260 | 8.91E-01 | 9.34E-01 | -1.04 | 512 | 1.85E-07 | 1.75E-06 | -0.77 | 530 | 2.11E-04 | 7.89E-03 | 0.10 | 334 | 7.03E-01 | 7.90E-01 |
| ILMN_1656310 | IDO1 | 8p11.22a-p11.21c | 0.84 | 2.41E-04 | 2.90E-03 | 8.79E-01 | 3.12E-03 | 1.14 | 1.05 | -0.08 | 0.98 | 63 | 3.08E-04 | 3.12E-03 | 1.05 | 69 | 3.22E-03 | 1.44E-02 | 0.85 | 109 | 1.84E-03 | 5.34E-03 | 0.95 | 32 | 9.13E-04 | 1.79E-02 | -0.08 | 352 | 8.23E-01 | 8.79E-01 |
| ILMN_1742824 | SPATA13 | 13q12.12b | 0.62 | 4.13E-04 | 4.23E-03 | 9.55E-01 | 9.01E-07 | 1.13 | 1.09 | -0.05 | 0.69 | 109 | 3.09E-04 | 3.12E-03 | -0.05 | 295 | 8.42E-01 | 9.03E-01 | 1.05 | 73 | 8.68E-08 | 9.01E-07 | 0.03 | 286 | 8.96E-01 | 9.55E-01 | 1.09 | 165 | 5.62E-05 | 3.22E-04 |
| ILMN_2056074 | RGPD4** | 2p11.2d | 0.53 | 2.08E-02 | 7.10E-02 | 9.60E-01 | 6.01E-03 | 1.13 | 1.16 | 0.03 | 0.19 | 298 | 4.68E-01 | 6.43E-01 | 0.03 | 264 | 9.33E-01 | 9.60E-01 | 0.83 | 113 | 2.11E-03 | 6.01E-03 | 0.67 | 69 | 1.80E-02 | 1.03E-01 | 1.16 | 125 | 1.91E-03 | 6.69E-03 |
| ILMN_1717063 | FBXO9 | 6p12.1d | -0.60 | 4.43E-05 | 8.85E-04 | 7.28E-01 | 1.58E-07 | 1.13 | 0.11 | -1.02 | -0.24 | 381 | 1.29E-01 | 2.76E-01 | 0.11 | 241 | 6.02E-01 | 7.28E-01 | -0.93 | 488 | 1.18E-08 | 1.58E-07 | -0.79 | 533 | 3.02E-06 | 8.28E-04 | -1.02 | 452 | 4.69E-06 | 3.83E-05 |
| ILMN_1779061 | ST6GALNAC3 | 1p31.1g-p31.1f | 0.22 | 4.16E-02 | 1.17E-01 | 9.06E-01 | 1.78E-08 | 1.13 | 1.09 | -0.03 | 0.18 | 301 | 1.34E-01 | 2.83E-01 | -0.03 | 291 | 8.47E-01 | 9.06E-01 | 0.24 | 247 | 4.47E-02 | 8.14E-02 | 0.07 | 254 | 5.79E-01 | 7.71E-01 | 1.09 | 154 | 5.49E-10 | 1.78E-08 |
| ILMN_1714650 | RASGRP4 | 19q13.2a | 0.12 | 3.02E-01 | 4.63E-01 | 7.97E-01 | 1.07E-06 | 1.13 | 1.00 | -0.12 | 0.33 | 243 | 1.11E-02 | 4.82E-02 | -0.07 | 301 | 6.92E-01 | 7.97E-01 | -0.12 | 333 | 3.36E-01 | 4.30E-01 | -0.11 | 361 | 4.29E-01 | 6.61E-01 | 1.00 | 234 | 7.01E-08 | 1.07E-06 |
| ILMN_1776939 | MS4A1 | 11q12.2a | 0.71 | 1.55E-03 | 1.11E-02 | 9.49E-01 | 4.91E-05 | 1.12 | 1.15 | 0.03 | 0.40 | 215 | 1.13E-01 | 2.52E-01 | 0.09 | 249 | 7.89E-01 | 8.67E-01 | 1.15 | 45 | 8.42E-06 | 4.91E-05 | 0.96 | 31 | 4.04E-04 | 1.13E-02 | 0.03 | 344 | 9.24E-01 | 9.49E-01 |
| ILMN_1788936 | KREMEN1 | 22q12.1c | 0.18 | 1.91E-01 | 3.39E-01 | 9.00E-01 | 1.37E-05 | 1.12 | 1.05 | -0.07 | 0.26 | 278 | 9.26E-02 | 2.20E-01 | -0.07 | 302 | 7.29E-01 | 8.23E-01 | 0.09 | 275 | 5.73E-01 | 6.61E-01 | -0.04 | 329 | 7.88E-01 | 9.00E-01 | 1.05 | 192 | 1.42E-06 | 1.37E-05 |
| ILMN_2325506 | BCAS4 | 20q13.13f | -0.34 | 2.39E-03 | 1.49E-02 | 9.62E-01 | 1.13E-07 | 1.11 | 0.11 | -1.01 | -0.61 | 492 | 6.12E-07 | 2.41E-05 | 0.11 | 242 | 4.97E-01 | 6.39E-01 | -0.24 | 355 | 4.18E-02 | 7.69E-02 | 0.01 | 294 | 9.14E-01 | 9.62E-01 | -1.01 | 437 | 5.04E-09 | 1.13E-07 |
| ILMN_1822495 | ##_HS.544127 | 0 | 0.40 | 7.41E-04 | 6.47E-03 | 4.17E-01 | 1.90E-09 | 1.11 | 1.31 | 0.20 | 0.43 | 206 | 1.73E-03 | 1.20E-02 | 0.20 | 200 | 2.69E-01 | 4.17E-01 | 0.38 | 210 | 5.64E-03 | 1.41E-02 | 0.25 | 163 | 8.25E-02 | 2.55E-01 | 1.31 | 72 | 3.82E-11 | 1.90E-09 |
| ILMN_1803824 | ZDHHC9 | Xq25h | -0.46 | 2.20E-06 | 9.67E-05 | 5.89E-01 | 1.93E-10 | 1.11 | 0.10 | -1.01 | -0.51 | 460 | 3.75E-07 | 1.73E-05 | 0.10 | 246 | 4.39E-01 | 5.89E-01 | -0.62 | 441 | 8.76E-10 | 1.71E-08 | -0.15 | 385 | 1.34E-01 | 3.39E-01 | -1.01 | 439 | 2.11E-12 | 1.93E-10 |
| ILMN_1799848 | ANKRD22 | 10q23.31b | 1.05 | 1.48E-06 | 7.36E-05 | 1.41E-01 | 5.89E-07 | 1.11 | 1.67 | 0.56 | 1.48 | 22 | 4.57E-09 | 5.89E-07 | 1.01 | 78 | 1.82E-03 | 9.21E-03 | 0.82 | 114 | 8.36E-04 | 2.70E-03 | 0.56 | 79 | 3.03E-02 | 1.41E-01 | 1.67 | 23 | 1.64E-06 | 1.55E-05 |
| ILMN_1726981 | VEGFB | 11q13.1a | -0.40 | 6.81E-07 | 4.02E-05 | 6.26E-01 | 1.01E-18 | 1.10 | -0.07 | -1.17 | -0.53 | 469 | 3.83E-11 | 1.80E-08 | -0.15 | 329 | 1.43E-01 | 2.63E-01 | -0.41 | 395 | 1.59E-07 | 1.54E-06 | -0.07 | 346 | 3.89E-01 | 6.26E-01 | -1.17 | 517 | 6.68E-23 | 1.01E-18 |
| ILMN_1756953 | GBP6 | 1p22.2c | 1.21 | 1.46E-06 | 7.29E-05 | 1.68E-01 | 4.55E-07 | 1.10 | 1.75 | 0.65 | 1.73 | 11 | 2.87E-09 | 4.55E-07 | 0.65 | 117 | 7.91E-02 | 1.68E-01 | 1.07 | 66 | 1.72E-04 | 6.81E-04 | 0.68 | 67 | 2.24E-02 | 1.18E-01 | 1.75 | 18 | 1.17E-05 | 8.40E-05 |
| ILMN_2130180 | RPL13P5 | 12p13.31d | -0.35 | 3.49E-04 | 3.73E-03 | 6.75E-01 | 1.32E-09 | 1.10 | 0.08 | -1.02 | -0.53 | 470 | 5.53E-07 | 2.26E-05 | 0.08 | 250 | 5.40E-01 | 6.75E-01 | -0.22 | 348 | 3.28E-02 | 6.30E-02 | -0.12 | 368 | 2.62E-01 | 5.02E-01 | -1.02 | 445 | 2.47E-11 | 1.32E-09 |
| ILMN_1700896 | SAP30 | 4q34.1c | 0.54 | 4.10E-06 | 1.56E-04 | 8.63E-01 | 1.06E-08 | 1.10 | 1.14 | 0.04 | 0.66 | 114 | 2.26E-07 | 1.18E-05 | 0.04 | 258 | 7.83E-01 | 8.63E-01 | 0.67 | 145 | 1.53E-07 | 1.48E-06 | 0.14 | 211 | 2.68E-01 | 5.08E-01 | 1.14 | 132 | 2.94E-10 | 1.06E-08 |
| ILMN_1701882 | FAM72B | 0 | 0.74 | 1.25E-07 | 1.17E-05 | 5.90E-01 | 1.35E-07 | 1.10 | 1.25 | 0.15 | 0.81 | 88 | 2.65E-07 | 1.32E-05 | 0.15 | 220 | 4.40E-01 | 5.90E-01 | 0.91 | 99 | 9.92E-09 | 1.35E-07 | 0.44 | 104 | 7.25E-03 | 5.91E-02 | 1.25 | 93 | 1.51E-08 | 2.85E-07 |
| ILMN_2256050 | SERPINA1 | 14q32.13a | -0.58 | 8.26E-04 | 6.98E-03 | 9.76E-01 | 1.78E-07 | 1.10 | -0.01 | -1.11 | -0.28 | 398 | 1.30E-01 | 2.76E-01 | -0.13 | 326 | 5.92E-01 | 7.19E-01 | -1.11 | 526 | 1.36E-08 | 1.78E-07 | -0.62 | 505 | 2.07E-03 | 2.81E-02 | -0.01 | 348 | 9.63E-01 | 9.76E-01 |
| ILMN_1887011 | IL23A | 0 | -0.63 | 6.50E-04 | 5.91E-03 | 5.04E-01 | 4.72E-05 | 1.09 | -0.25 | -1.35 | -0.93 | 539 | 1.30E-05 | 2.57E-04 | -0.25 | 368 | 3.53E-01 | 5.04E-01 | -0.44 | 402 | 3.42E-02 | 6.53E-02 | -0.33 | 447 | 1.30E-01 | 3.34E-01 | -1.35 | 538 | 5.99E-06 | 4.72E-05 |
| ILMN_1692398 | CNTNAP1 | 17q21.31a | -0.35 | 2.71E-02 | 8.61E-02 | 9.64E-01 | 2.60E-04 | 1.09 | 0.06 | -1.03 | -0.65 | 505 | 3.00E-04 | 3.06E-03 | 0.02 | 270 | 9.39E-01 | 9.64E-01 | -0.29 | 364 | 1.07E-01 | 1.68E-01 | 0.06 | 264 | 7.54E-01 | 8.82E-01 | -1.03 | 460 | 4.40E-05 | 2.60E-04 |
| ILMN_2198239 | HGD | 0 | -0.81 | 5.53E-06 | 1.91E-04 | 7.52E-01 | 1.09E-07 | 1.09 | -0.13 | -1.22 | -0.63 | 503 | 2.12E-03 | 1.41E-02 | -0.13 | 324 | 6.33E-01 | 7.52E-01 | -1.22 | 539 | 7.74E-09 | 1.09E-07 | -0.69 | 516 | 1.48E-03 | 2.35E-02 | -0.91 | 418 | 1.46E-03 | 5.34E-03 |
| ILMN_1808405 | HLA-DQA1 | 6p21.32b | -0.41 | 3.87E-03 | 2.11E-02 | 6.31E-01 | 1.63E-06 | 1.09 | -0.15 | -1.23 | -0.43 | 439 | 8.13E-03 | 3.84E-02 | -0.17 | 339 | 4.23E-01 | 5.73E-01 | -0.44 | 398 | 7.46E-03 | 1.79E-02 | -0.15 | 380 | 3.95E-01 | 6.31E-01 | -1.23 | 523 | 1.18E-07 | 1.63E-06 |
| ILMN_1733270 | CD163 | 12p13.31c | 0.50 | 2.12E-04 | 2.67E-03 | 4.55E-01 | 1.27E-07 | 1.08 | 1.28 | 0.20 | 0.58 | 137 | 1.52E-04 | 1.82E-03 | 0.34 | 157 | 9.15E-02 | 1.88E-01 | 0.47 | 182 | 2.09E-03 | 5.96E-03 | 0.20 | 180 | 2.19E-01 | 4.55E-01 | 1.28 | 84 | 5.78E-09 | 1.27E-07 |
| ILMN_1656292 | RPL21P131 | 19q13.11b | -0.23 | 9.64E-02 | 2.11E-01 | 9.87E-01 | 1.96E-05 | 1.08 | 0.02 | -1.07 | -0.34 | 416 | 2.91E-02 | 9.71E-02 | 0.02 | 271 | 9.38E-01 | 9.63E-01 | -0.08 | 320 | 627E-01 | 7.07E-01 | -0.01 | 304 | 9.70E-01 | 9.87E-01 | -1.07 | 473 | 2.16E-06 | 1.96E-05 |
| ILMN_1808059 | BCAS4 | 20q13.13f | -0.36 | 8.99E-04 | 7.42E-03 | 9.53E-01 | 3.02E-09 | 1.08 | -0.02 | -1.10 | -0.61 | 494 | 2.17E-07 | 1.14E-05 | -0.20 | 344 | 1.89E-01 | 3.22E-01 | -0.22 | 347 | 5.49E-02 | 9.68E-02 | -0.02 | 313 | 8.92E-01 | 9.53E-01 | -1.10 | 491 | 6.56E-11 | 3.02E-09 |
| ILMN_2413537 | HEMGN | 9q22.33b | -0.48 | 2.43E-02 | 7.97E-02 | 9.71E-01 | 2.88E-03 | 1.08 | -0.02 | -1.10 | -0.02 | 339 | 9.47E-01 | 9.71E-01 | -0.16 | 335 | 6.24E-01 | 7.44E-01 | -0.57 | 428 | 1.99E-02 | 4.15E-02 | -1.10 | 556 | 3.12E-05 | 2.88E-03 | -0.63 | 401 | 6.87E-02 | 1.32E-01 |
| ILMN_1730054 | GSTT1 | 22q11.23b | -1.03 | 4.26E-03 | 2.27E-02 | 6.93E-01 | 7.90E-03 | 1.08 | -0.32 | -1.40 | -1.40 | 559 | 1.00E-03 | 7.90E-03 | -0.32 | 388 | 5.62E-01 | 6.93E-01 | -1.02 | 504 | 1.65E-02 | 3.53E-02 | -0.66 | 511 | 1.42E-01 | 3.51E-01 | -1.16 | 512 | 5.01E-02 | 1.02E-01 |
| ILMN_1756469 | GAMT | 19p13.3i | -0.50 | 5.37E-07 | 3.42E-05 | 6.75E-01 | 1.16E-12 | 1.08 | -0.08 | -1.16 | -0.70 | 513 | 1.09E-11 | 7.82E-09 | -0.08 | 308 | 5.41E-01 | 6.75E-01 | -0.53 | 421 | 1.51E-07 | 1.47E-06 | -0.10 | 360 | 3.49E-01 | 5.87E-01 | -1.16 | 511 | 2.92E-15 | 1.16E-12 |
| ILMN_1773650 | LRRN3 | 7q31.1b | -1.15 | 6.39E-05 | 1.16E-03 | 1.95E-01 | 2.17E-05 | 1.08 | -0.67 | -1.75 | -1.68 | 562 | 5.14E-07 | 2.17E-05 | -1.75 | 564 | 6.04E-05 | 6.36E-04 | -0.78 | 468 | 1.70E-02 | 3.64E-02 | -0.67 | 514 | 5.20E-02 | 1.95E-01 | -1.15 | 507 | 1.24E-02 | 3.23E-02 |
| ILMN_1788439 | TNRC18 | 7p22.1c | 0.18 | 1.96E-01 | 3.45E-01 | 7.20E-01 | 8.57E-05 | 1.08 | 1.00 | -0.07 | 0.26 | 274 | 1.05E-01 | 2.39E-01 | 0.17 | 212 | 4.13E-01 | 5.64E-01 | -0.07 | 319 | 6.42E-01 | 7.20E-01 | 0.12 | 222 | 4.92E-01 | 7.08E-01 | 1.00 | 231 | 1.20E-05 | 8.57E-05 |
| ILMN_1680777 | MAP2K3 | 17p11.2b | -0.55 | 7.84E-03 | 3.50E-02 | 7.86E-01 | 7.54E-04 | 1.08 | -0.10 | -1.18 | -0.10 | 350 | 6.51E-01 | 7.86E-01 | -0.25 | 367 | 4.06E-01 | 5.57E-01 | -0.65 | 447 | 5.34E-03 | 1.35E-02 | -1.18 | 560 | 2.57E-06 | 7.54E-04 | -0.53 | 391 | 1.03E-01 | 1.83E-01 |
| ILMN_1821352 | ##_HS.564504 | 0 | -0.47 | 1.98E-03 | 1.31E-02 | 9.66E-01 | 2.26E-05 | 1.08 | 0.02 | -1.06 | -0.84 | 535 | 5.51E-07 | 2.26E-05 | -0.08 | 307 | 7.16E-01 | 8.14E-01 | -0.35 | 381 | 3.32E-02 | 6.36E-02 | 0.02 | 293 | 9.24E-01 | 9.66E-01 | -1.06 | 471 | 5.69E-06 | 4.52E-05 |
| ILMN_1655525 | FLCN | 17p11.2h | -0.71 | 8.37E-04 | 7.04E-03 | 9.85E-01 | 8.09E-05 | 1.08 | 0.01 | -1.07 | -0.35 | 420 | 1.47E-01 | 3.02E-01 | 0.01 | 273 | 9.72E-01 | 9.85E-01 | -1.07 | 516 | 1.51E-05 | 8.09E-05 | -0.99 | 551 | 1.34E-04 | 6.10E-03 | -0.92 | 420 | 6.87E-03 | 1.96E-02 |
| ILMN_1768575 | SFTPD | 10q22.3f | 0.40 | 5.65E-02 | 1.45E-01 | 9.24E-01 | 3.40E-03 | 1.08 | 1.12 | 0.05 | 0.24 | 286 | 3.17E-01 | 5.02E-01 | 1.12 | 53 | 5.19E-04 | 3.40E-03 | 0.20 | 256 | 4.11E-01 | 5.07E-01 | 0.71 | 61 | 6.18E-03 | 5.40E-02 | 0.05 | 340 | 8.87E-01 | 9.24E-01 |
| ILMN_1865736 | SAMD4A-AS1 | 0 | -0.73 | 5.83E-05 | 1.08E-03 | 9.98E-01 | 5.17E-06 | 1.07 | 0.00 | -1.07 | -0.55 | 478 | 8.63E-03 | 4.02E-02 | 0.00 | 281 | 9.97E-01 | 9.98E-01 | -1.07 | 517 | 6.34E-07 | 5.17E-06 | -0.66 | 512 | 2.96E-03 | 3.43E-02 | -1.03 | 455 | 4.74E-04 | 2.04E-03 |
| ILMN_1685538 | RPL4P5 | 0 | -0.35 | 1.42E-04 | 2.02E-03 | 8.84E-01 | 5.20E-11 | 1.06 | 0.03 | -1.03 | -0.54 | 472 | 6.75E-08 | 4.94E-06 | 0.03 | 263 | 8.14E-01 | 8.84E-01 | -0.27 | 360 | 5.22E-03 | 1.32E-02 | -0.05 | 335 | 6.05E-01 | 7.89E-01 | -1.03 | 459 | 4.28E-13 | 5.20E-11 |
| ILMN_2131022 | LINC01002 | 19p13.3j | -0.62 | 8.08E-05 | 1.36E-03 | 7.92E-01 | 4.10E-09 | 1.06 | -0.09 | -1.15 | -0.46 | 457 | 7.32E-03 | 3.54E-02 | -0.09 | 314 | 6.85E-01 | 7.92E-01 | -1.15 | 530 | 1.67E-10 | 4.10E-09 | -0.55 | 492 | 2.75E-03 | 3.31E-02 | -0.20 | 362 | 3.94E-01 | 5.17E-01 |
| ILMN_1658247 | OAS1 | 12q24.13b | 0.90 | 4.89E-03 | 2.50E-02 | 5.70E-01 | 1.56E-02 | 1.06 | 1.46 | 0.40 | 1.16 | 42 | 2.41E-03 | 1.56E-02 | 1.46 | 13 | 3.83E-03 | 1.64E-02 | 0.66 | 150 | 8.45E-02 | 1.38E-01 | 0.80 | 48 | 4.66E-02 | 1.83E-01 | 0.40 | 302 | 4.50E-01 | 5.70E-01 |
| ILMN_1821975 | IL23A | 0 | -0.39 | 4.74E-03 | 2.45E-02 | 9.60E-01 | 2.55E-05 | 1.06 | 0.02 | -1.04 | -0.62 | 500 | 7.72E-05 | 1.08E-03 | -0.29 | 380 | 1.60E-01 | 2.86E-01 | -0.30 | 365 | 5.70E-02 | 9.98E-02 | 0.02 | 290 | 9.08E-01 | 9.60E-01 | -1.04 | 463 | 2.94E-06 | 2.55E-05 |
| ILMN_1732296 | ID3 | 1p36.12a | -0.44 | 2.47E-04 | 2.94E-03 | 9.91E-01 | 3.74E-07 | 1.05 | 0.00 | -1.05 | -0.69 | 510 | 1.85E-07 | 1.04E-05 | 0.00 | 277 | 9.83E-01 | 9.91E-01 | -0.40 | 392 | 2.08E-03 | 5.96E-03 | -0.03 | 319 | 8.40E-01 | 9.26E-01 | -1.05 | 465 | 2.05E-08 | 3.74E-07 |
| ILMN_2072598 | LOC401357 | 7p11.2a | -0.54 | 5.89E-07 | 3.65E-05 | 9.10E-01 | 1.19E-15 | 1.05 | 0.03 | -1.03 | -0.43 | 438 | 9.68E-05 | 1.29E-03 | 0.03 | 265 | 8.53E-01 | 9.10E-01 | -1.03 | 506 | 2.53E-18 | 1.19E-15 | -0.35 | 454 | 2.39E-03 | 3.06E-02 | -0.33 | 377 | 3.05E-02 | 6.85E-02 |
| ILMN_1722158 | CASP5 | 11q22.3b | 0.24 | 1.34E-01 | 2.65E-01 | 9.96E-01 | 3.14E-04 | 1.05 | 1.05 | 0.00 | 0.40 | 213 | 2.78E-02 | 9.39E-02 | 0.06 | 254 | 7.88E-01 | 8.67E-01 | 0.00 | 300 | 9.95E-01 | 9.96E-01 | 0.11 | 230 | 5.82E-01 | 7.73E-01 | 1.05 | 189 | 5.47E-05 | 3.14E-04 |
| ILMN_1726153 | CYTOR | 2p11.2d | 0.40 | 2.78E-03 | 1.67E-02 | 9.82E-01 | 1.49E-05 | 1.05 | 1.04 | -0.01 | 0.57 | 144 | 2.28E-04 | 2.47E-03 | -0.01 | 284 | 9.68E-01 | 9.82E-01 | 0.39 | 207 | 1.11E-02 | 2.51E-02 | 0.04 | 279 | 8.22E-01 | 9.17E-01 | 1.04 | 195 | 1.56E-06 | 1.49E-05 |
| ILMN_2257015 | AGER | 6p21.32b | 0.30 | 1.55E-02 | 5.74E-02 | 8.36E-01 | 6.71E-07 | 1.05 | 1.09 | 0.04 | 0.38 | 223 | 5.36E-03 | 2.79E-02 | 0.42 | 143 | 2.10E-02 | 6.09E-02 | 0.04 | 288 | 7.83E-01 | 8.36E-01 | 0.19 | 183 | 1.88E-01 | 4.13E-01 | 1.09 | 162 | 4.09E-08 | 6.71E-07 |
| ILMN_1732534 | CHMP5 | 9p13.3e | 0.69 | 4.80E-07 | 3.16E-05 | 3.03E-01 | 1.11E-08 | 1.05 | 1.31 | 0.26 | 0.73 | 104 | 1.03E-06 | 3.59E-05 | 0.26 | 186 | 1.74E-01 | 3.03E-01 | 0.94 | 97 | 5.22E-10 | 1.11E-08 | 0.36 | 120 | 1.78E-02 | 1.03E-01 | 1.31 | 74 | 5.98E-10 | 1.91E-08 |
| ILMN_1760062 | IFI44 | 1p31.1e | 1.78 | 3.61E-09 | 1.01E-06 | 7.14E-02 | 1.04E-07 | 1.05 | 2.06 | 1.01 | 1.86 | 7 | 1.33E-07 | 8.19E-06 | 1.01 | 79 | 2.58E-02 | 7.14E-02 | 2.06 | 5 | 7.23E-09 | 1.04E-07 | 1.50 | 7 | 4.58E-05 | 3.44E-03 | 2.00 | 7 | 3.90E-05 | 2.34E-04 |
| ILMN_1661174 | RPL21P97 | 11p15.4a | -0.18 | 1.83E-01 | 3.28E-01 | 9.98E-01 | 3.27E-05 | 1.04 | 0.03 | -1.02 | -0.28 | 396 | 6.95E-02 | 1.80E-01 | -0.09 | 312 | 6.72E-01 | 7.82E-01 | 000 | 299 | 9.98E-01 | 9.98E-01 | 0.03 | 284 | 8.71E-01 | 9.43E-01 | -1.02 | 444 | 3.92E-06 | 3.27E-05 |
| ILMN_1818577 | CPLX2 | 0 | 0.51 | 8.54E-02 | 1.94E-01 | 7.69E-01 | 3.83E-02 | 1.04 | 1.17 | 0.14 | 0.31 | 252 | 3.80E-01 | 5.67E-01 | 1.17 | 43 | 1.15E-02 | 3.83E-02 | 0.14 | 265 | 6.99E-01 | 7.69E-01 | 1.00 | 27 | 6.97E-03 | 5.76E-02 | 0.30 | 311 | 5.40E-01 | 6.52E-01 |
| ILMN_2390586 | SP100 | 2q37.1a | 0.64 | 9.16E-10 | 3.63E-07 | 5.23E-01 | 4.37E-11 | 1.04 | 1.16 | 0.13 | 0.75 | 96 | 2.85E-11 | 1.38E-08 | 0.13 | 237 | 3.71E-01 | 5.23E-01 | 0.80 | 119 | 1.82E-12 | 8.24E-11 | 0.26 | 150 | 2.02E-02 | 1.11E-01 | 1.16 | 126 | 3.14E-13 | 4.37E-11 |
| ILMN_1654890 | LENG8 | 19q13.42a | 0.70 | 2.37E-07 | 1.88E-05 | 9.72E-01 | 4.85E-09 | 1.04 | 1.02 | -0.01 | 0.72 | 105 | 1.70E-06 | 5.27E-05 | -0.01 | 285 | 9.51E-01 | 9.72E-01 | 0.98 | 94 | 2.04E-10 | 4.85E-09 | 0.36 | 123 | 2.20E-02 | 1.17E-01 | 1.02 | 213 | 1.09E-06 | 1.09E-05 |
| ILMN_1726589 | CD248 | 11q13.1e | -1.47 | 9.02E-08 | 9.22E-06 | 6.04E-02 | 4.94E-07 | 1.04 | -0.89 | -1.93 | -1.93 | 564 | 3.35E-09 | 4.94E-07 | -1.51 | 559 | 3.11E-04 | 2.28E-03 | -1.34 | 553 | 2.87E-05 | 1.42E-04 | -0.89 | 544 | 7.52E-03 | 6.04E-02 | -1.84 | 561 | 3.79E-05 | 2.29E-04 |
| ILMN_2322498 | RORA | 15q22.2a-q22.2b | 0.54 | 2.34E-04 | 2.84E-03 | 9.33E-01 | 3.35E-08 | 1.03 | 1.00 | -0.03 | 0.45 | 195 | 5.58E-03 | 2.88E-02 | -0.03 | 290 | 8.90E-01 | 9.33E-01 | 1.00 | 89 | 1.93E-09 | 3.35E-08 | 0.26 | 154 | 1.26E-01 | 3.26E-01 | 0.57 | 283 | 1.15E-02 | 3.03E-02 |
| ILMN_1724708 | RAB33A | Xq25h | -0.40 | 2.50E-05 | 5.84E-04 | 8.97E-01 | 1.91E-10 | 1.03 | -0.03 | -1.06 | -0.51 | 461 | 1.04E-06 | 3.60E-05 | -0.03 | 289 | 8.33E-01 | 8.97E-01 | -0.41 | 394 | 8.31E-05 | 3.61E-04 | -0.13 | 369 | 2.43E-01 | 4.81E-01 | -1.06 | 468 | 2.08E-12 | 1.91E-10 |
| ILMN_1683844 | HSD17B13 | 4q22.1a | 0.39 | 4.78E-04 | 4.71E-03 | 8.07E-01 | 5.56E-08 | 1.03 | 1.09 | 0.06 | 0.47 | 181 | 1.89E-04 | 2.13E-03 | 0.06 | 256 | 7.06E-01 | 8.07E-01 | 0.31 | 224 | 1.40E-02 | 3.08E-02 | 0.26 | 151 | 4.65E-02 | 1.83E-01 | 1.09 | 157 | 2.14E-09 | 5.56E-08 |
| ILMN_1753347 | DEFA4 | 8p23.1f | 0.58 | 4.50E-02 | 1.24E-01 | 9.92E-01 | 1.61E-02 | 1.03 | 1.03 | 0.01 | 1.03 | 58 | 2.52E-03 | 1.61E-02 | 0.01 | 274 | 9.85E-01 | 9.92E-01 | 0.32 | 218 | 3.40E-01 | 4.35E-01 | 012 | 221 | 7.35E-01 | 8.71E-01 | 0.92 | 244 | 5.29E-02 | 1.07E-01 |

### Online Supplemental Table S4 (Continued)

| | | | S1 | | | M1 through M5 | | | | | M1 | | | | M2 | | | | M3 | | | | M4 | | | | M5 | | | |
|---|---|---|---|---|---|---|---|---|---|---|---|---|---|---|---|---|---|---|---|---|---|---|---|---|---|---|---|---|---|---|
| ID | Gene_Symbol | Cytoband | FC | $P_{val}$ | $P_{adj}$ | Max $P_{adj}$ | Min $P_{adj}$ | ΔFC | Max FC | Min FC | FC | FC Rank | $P_{val}$ | $P_{adj}$ | FC | FC Rank | $P_{val}$ | $P_{adj}$ | FC | FC Rank | $P_{val}$ | $P_{adj}$ | FC | FC Rank | $P_{val}$ | $P_{adj}$ | FC | FC Rank | $P_{val}$ | $P_{adj}$ |
| ILMN_2354191 | CD8B | 2p11.2e | -0.64 | 8.12E-04 | 6.90E-03 | 4.20E-01 | 2.12E-04 | 1.02 | -0.31 | -1.33 | -0.76 | 524 | 8.72E-04 | 7.07E-03 | -0.56 | 446 | 5.77E-02 | 1.32E-01 | -0.65 | 448 | 4.03E-03 | 1.06E-02 | -0.31 | 435 | 1.92E-01 | 4.20E-01 | -1.33 | 537 | 3.46E-05 | 2.12E-04 |
| ILMN_1762284 | ASPRV1 | 2p14a | 0.36 | 2.51E-03 | 1.55E-02 | 7.17E-01 | 1.74E-07 | 1.02 | 1.08 | 0.06 | 0.60 | 130 | 5.05E-06 | 1.25E-04 | 0.17 | 215 | 3.24E-01 | 4.75E-01 | 0.06 | 281 | 6.39E-01 | 7.17E-01 | 0.14 | 214 | 3.02E-01 | 5.42E-01 | 1.08 | 169 | 8.42E-09 | 1.74E-07 |
| ILMN_1691339 | CLEC1A | 12p13.2c | 0.36 | 6.45E-03 | 3.04E-02 | 9.97E-01 | 2.14E-05 | 1.01 | 1.01 | 0.00 | 0.59 | 135 | 1.14E-04 | 1.45E-03 | 0.00 | 279 | 9.95E-01 | 9.97E-01 | 0.10 | 273 | 5.16E-01 | 6.07E-01 | 0.17 | 197 | 2.82E-01 | 5.23E-01 | 1.01 | 224 | 2.41E-06 | 2.14E-05 |
| ILMN_1767322 | EDAR | 2q13a | -0.71 | 6.46E-04 | 5.89E-03 | 6.68E-01 | 2.79E-05 | 1.01 | -0.19 | -1.20 | -1.20 | 555 | 7.46E-07 | 2.79E-05 | -0.92 | 496 | 3.50E-03 | 1.54E-02 | -0.41 | 396 | 8.19E-02 | 1.35E-01 | -0.19 | 403 | 4.38E-01 | 6.68E-01 | -1.17 | 516 | 4.80E-04 | 2.07E-03 |
| ILMN_2048591 | LRRN3 | 7q31.1b | -0.93 | 7.91E-04 | 6.79E-03 | 2.42E-01 | 1.77E-04 | 1.00 | -0.44 | -1.44 | -1.44 | 560 | 7.89E-06 | 1.77E-04 | -1.23 | 543 | 3.41E-03 | 1.51E-02 | -0.44 | 399 | 1.66E-01 | 2.42E-01 | -0.70 | 519 | 3.66E-02 | 1.58E-01 | -0.94 | 422 | 3.39E-02 | 7.45E-02 |
| ILMN_1904208 | NT5DC4 | 0 | 0.24 | 8.56E-03 | 3.72E-02 | 9.49E-01 | 1.82E-10 | 1.00 | 1.02 | 0.01 | 0.27 | 271 | 6.56E-03 | 3.26E-02 | 0.01 | 272 | 9.15E-01 | 9.49E-01 | 0.12 | 268 | 2.12E-01 | 2.95E-01 | 0.15 | 209 | 1.41E-01 | 3.50E-01 | 1.02 | 218 | 1.90E-12 | 1.82E-10 |
| ILMN_1782938 | SLC16A10 | 6q21h | -0.64 | 8.61E-05 | 1.42E-03 | 1.90E-01 | 1.09E-05 | 1.00 | -0.29 | -1.28 | -0.96 | 540 | 4.26E-07 | 1.89E-05 | -1.28 | 547 | 3.17E-07 | 1.09E-05 | -0.29 | 363 | 1.24E-01 | 1.90E-01 | -0.47 | 478 | 1.69E-02 | 9.89E-02 | -0.70 | 404 | 7.40E-03 | 2.09E-02 |
| ILMN_1689786 | ASMTL | Xp22.33d,Yp11.32a | -0.44 | 1.68E-07 | 1.49E-05 | 5.25E-01 | 2.60E-14 | 0.99 | -0.10 | -1.09 | -0.44 | 444 | 3.91E-07 | 1.78E-05 | -0.10 | 317 | 3.73E-01 | 5.25E-01 | -0.57 | 427 | 1.80E-10 | 4.36E-09 | -0.19 | 401 | 3.59E-02 | 1.56E-01 | -1.09 | 487 | 2.59E-17 | 2.60E-14 |
| ILMN_1755910 | ##_LOC648366 | 0 | 0.52 | 3.09E-06 | 1.27E-04 | 9.38E-01 | 5.43E-07 | 0.99 | 1.01 | 0.02 | 0.65 | 116 | 5.09E-07 | 2.17E-05 | 0.02 | 267 | 8.98E-01 | 9.38E-01 | 0.51 | 178 | 7.82E-05 | 3.42E-04 | 0.31 | 133 | 1.94E-02 | 1.08E-01 | 1.01 | 225 | 3.16E-08 | 5.43E-07 |
| ILMN_1712719 | MAP7 | 6q23.3b | -0.65 | 9.29E-05 | 1.50E-03 | 6.58E-01 | 5.41E-08 | 0.98 | -0.16 | -1.14 | -0.42 | 435 | 2.43E-02 | 8.46E-02 | -0.16 | 332 | 5.20E-01 | 6.58E-01 | -1.14 | 529 | 3.41E-09 | 5.41E-08 | -0.58 | 497 | 3.33E-03 | 3.68E-02 | -0.23 | 365 | 3.70E-01 | 4.93E-01 |
| ILMN_1748601 | CD8B | 2p11.2e | -0.62 | 9.26E-04 | 7.58E-03 | 3.60E-01 | 1.89E-04 | 0.98 | -0.34 | -1.32 | -0.71 | 514 | 1.55E-03 | 1.10E-02 | -0.56 | 445 | 5.71E-02 | 1.31E-01 | -0.62 | 439 | 5.65E-03 | 1.41E-02 | -0.34 | 450 | 1.48E-01 | 3.60E-01 | -1.32 | 534 | 3.01E-05 | 1.89E-04 |
| ILMN_1672486 | TCF7L2 | 10q25.2b-q25.3a | 0.59 | 1.10E-04 | 1.70E-03 | 5.51E-01 | 2.22E-05 | 0.98 | 1.17 | 0.19 | 0.73 | 101 | 3.05E-05 | 5.20E-04 | 0.29 | 170 | 2.09E-01 | 3.48E-01 | 0.63 | 158 | 3.52E-04 | 1.27E-03 | 0.19 | 188 | 3.10E-01 | 5.51E-01 | 1.17 | 123 | 2.51E-06 | 2.22E-05 |
| ILMN_1808837 | RPL7AP62 | 0 | -0.42 | 7.92E-06 | 2.51E-04 | 7.98E-01 | 2.53E-10 | 0.98 | -0.05 | -1.03 | -0.59 | 487 | 1.47E-08 | 1.55E-06 | -0.05 | 297 | 6.94E-01 | 7.98E-01 | -0.36 | 382 | 4.24E-04 | 1.49E-03 | -0.15 | 381 | 1.61E-01 | 3.79E-01 | -1.03 | 456 | 2.87E-12 | 2.53E-10 |
| ILMN_1676580 | H4C4 | 6p22.1d | 0.26 | 7.28E-02 | 1.74E-01 | 9.01E-01 | 9.50E-05 | 0.96 | 1.01 | 0.05 | 0.43 | 204 | 8.31E-03 | 3.91E-02 | 0.14 | 224 | 4.98E-01 | 6.39E-01 | 0.05 | 283 | 7.45E-01 | 8.05E-01 | 0.05 | 273 | 7.90E-01 | 9.01E-01 | 1.01 | 229 | 1.35E-05 | 9.50E-05 |
| ILMN_1765165 | VPS13B | 8q22.2a-q22.2b | 0.45 | 8.62E-04 | 7.19E-03 | 7.41E-01 | 2.43E-05 | 0.96 | 1.06 | 0.10 | 0.54 | 152 | 7.50E-04 | 6.26E-03 | 0.10 | 243 | 6.20E-01 | 7.41E-01 | 0.43 | 192 | 6.82E-03 | 1.66E-02 | 0.30 | 138 | 7.40E-02 | 2.39E-01 | 1.06 | 180 | 2.78E-06 | 2.43E-05 |
| ILMN_2138589 | MERTK | 2q13c-q13d | 0.82 | 8.72E-07 | 4.84E-05 | 3.65E-01 | 3.39E-05 | 0.96 | 1.27 | 0.31 | 0.90 | 76 | 5.01E-06 | 1.25E-04 | 0.31 | 164 | 2.24E-01 | 3.65E-01 | 0.88 | 105 | 7.58E-06 | 4.47E-05 | 0.67 | 68 | 1.14E-03 | 2.02E-02 | 1.27 | 87 | 4.09E-06 | 3.39E-05 |
| ILMN_1787441 | CCDC17 | 1p34.1b | 0.37 | 3.15E-04 | 3.46E-03 | 6.98E-01 | 2.53E-08 | 0.96 | 1.01 | 0.06 | 0.54 | 151 | 3.17E-06 | 8.54E-05 | 0.42 | 142 | 4.75E-03 | 1.95E-02 | 0.06 | 282 | 6.16E-01 | 6.98E-01 | 0.33 | 128 | 6.64E-03 | 5.62E-02 | 1.01 | 220 | 8.31E-10 | 2.53E-08 |
| ILMN_2393693 | LRRC37A4P | 17q21.31d | 0.63 | 1.62E-03 | 1.14E-02 | 8.24E-01 | 4.04E-03 | 0.96 | 1.06 | 0.10 | 0.57 | 143 | 1.53E-02 | 6.06E-02 | 1.06 | 65 | 6.49E-04 | 4.04E-03 | 0.63 | 157 | 7.44E-03 | 1.78E-02 | 0.78 | 50 | 1.61E-03 | 2.47E-02 | 0.10 | 333 | 7.50E-01 | 8.24E-01 |
| ILMN_1791759 | CXCL10 | 4q21.1a | 1.60 | 1.89E-09 | 6.20E-07 | 2.36E-02 | 1.26E-09 | 0.95 | 2.04 | 1.09 | 1.74 | 9 | 1.18E-08 | 1.30E-06 | 1.19 | 34 | 2.24E-03 | 1.09E-02 | 2.04 | 6 | 4.29E-11 | 1.26E-09 | 1.16 | 14 | 2.27E-04 | 8.17E-03 | 1.09 | 160 | 8.55E-03 | 2.36E-02 |
| ILMN_1685339 | TPM1 | 15q22.2b | 0.32 | 6.03E-03 | 2.92E-02 | 7.71E-01 | 1.65E-06 | 0.95 | 1.03 | 0.08 | 0.25 | 282 | 6.46E-02 | 1.71E-01 | 0.08 | 251 | 6.57E-01 | 7.71E-01 | 0.38 | 208 | 4.82E-03 | 1.23E-02 | 0.22 | 173 | 1.27E-01 | 3.28E-01 | 1.03 | 210 | 1.19E-07 | 1.65E-06 |
| ILMN_1792538 | CD7 | 17q25.3g | -0.47 | 4.53E-08 | 6.09E-06 | 4.15E-01 | 2.97E-13 | 0.95 | -0.12 | -1.07 | -0.63 | 502 | 9.20E-12 | 6.93E-09 | -0.21 | 347 | 6.73E-02 | 1.49E-01 | -0.49 | 414 | 5.38E-08 | 5.95E-07 | -0.12 | 367 | 1.89E-01 | 4.15E-01 | -1.07 | 476 | 3.74E-16 | 2.97E-13 |
| ILMN_1695812 | KRT72 | 12q13.13d | 0.42 | 1.07E-01 | 2.27E-01 | 8.05E-01 | 3.18E-02 | 0.94 | 1.06 | 0.13 | 0.13 | 311 | 6.81E-01 | 8.05E-01 | 1.06 | 64 | 9.02E-03 | 3.18E-02 | 0.38 | 209 | 2.19E-01 | 3.03E-01 | 0.68 | 65 | 3.60E-02 | 1.56E-01 | 0.36 | 305 | 3.99E-01 | 5.22E-01 |
| ILMN_1703324 | PDSS1 | 10p12.1b | 0.45 | 1.75E-05 | 4.50E-04 | 4.06E-01 | 1.46E-09 | 0.93 | 1.09 | 0.16 | 0.44 | 201 | 1.07E-04 | 1.38E-03 | 0.23 | 189 | 1.24E-01 | 2.37E-01 | 0.61 | 161 | 1.20E-07 | 1.20E-06 | 0.16 | 204 | 1.82E-01 | 4.06E-01 | 1.09 | 158 | 2.76E-11 | 1.46E-09 |
| ILMN_1701603 | ALPL | 1p36.12b | -0.69 | 7.56E-07 | 4.38E-05 | 3.47E-01 | 1.11E-11 | 0.93 | -0.25 | -1.18 | -0.59 | 486 | 1.24E-04 | 1.55E-03 | -0.38 | 404 | 5.60E-02 | 1.29E-01 | -1.18 | 535 | 1.66E-13 | 1.11E-11 | -0.50 | 481 | 1.95E-03 | 2.74E-02 | -0.25 | 368 | 2.33E-01 | 3.47E-01 |
| ILMN_1782487 | GBP1P1 | 0 | 1.32 | 4.81E-11 | 5.17E-08 | 6.23E-02 | 2.43E-10 | 0.93 | 1.58 | 0.65 | 1.56 | 17 | 6.53E-12 | 5.46E-09 | 0.65 | 116 | 2.17E-02 | 6.23E-02 | 1.56 | 13 | 6.25E-12 | 2.43E-10 | 0.75 | 55 | 1.10E-03 | 1.99E-02 | 1.58 | 32 | 3.29E-07 | 3.91E-06 |
| ILMN_1708004 | GTSCR1 | 18q22.2b | 0.87 | 1.30E-04 | 1.89E-03 | 2.89E-01 | 8.79E-05 | 0.92 | 1.31 | 0.38 | 0.38 | 224 | 1.38E-01 | 2.89E-01 | 0.79 | 97 | 2.10E-02 | 6.09E-02 | 1.14 | 48 | 1.65E-05 | 8.79E-05 | 1.31 | 9 | 3.16E-06 | 8.40E-04 | 0.60 | 280 | 1.00E-01 | 1.78E-01 |
| ILMN_1703430 | YJU2 | 19p13.3d | -0.39 | 2.79E-03 | 1.68E-02 | 7.47E-01 | 1.43E-05 | 0.92 | -0.09 | -1.01 | -0.34 | 413 | 2.24E-02 | 8.00E-02 | -0.10 | 316 | 6.12E-01 | 7.35E-01 | -0.56 | 426 | 2.03E-04 | 7.89E-04 | -0.09 | 357 | 5.42E-01 | 7.47E-01 | -1.01 | 441 | 1.49E-06 | 1.43E-05 |
| ILMN_1657871 | RSAD2 | 2p25.2a | 2.26 | 1.13E-08 | 2.38E-06 | 3.13E-02 | 1.22E-06 | 0.92 | 2.49 | 1.57 | 2.49 | 3 | 9.98E-08 | 6.48E-06 | 1.57 | 10 | 8.82E-03 | 3.13E-02 | 2.48 | 3 | 1.22E-07 | 1.22E-06 | 1.84 | 4 | 1.47E-04 | 6.35E-03 | 2.41 | 3 | 1.82E-04 | 8.98E-04 |
| ILMN_1742001 | CD160 | 1q21.1b | -0.40 | 1.61E-02 | 5.89E-02 | 5.64E-01 | 2.49E-04 | 0.91 | -0.20 | -1.11 | -0.52 | 466 | 7.17E-03 | 3.48E-02 | -0.47 | 424 | 6.02E-02 | 1.37E-01 | -0.24 | 356 | 2.01E-01 | 2.83E-01 | -0.20 | 406 | 3.24E-01 | 5.64E-01 | -1.11 | 497 | 4.20E-05 | 2.49E-04 |
| ILMN_1771482 | ELAPOR1 | 1p13.3b | -0.81 | 7.82E-04 | 6.73E-03 | 8.26E-01 | 9.73E-04 | 0.91 | -0.12 | -1.03 | -0.76 | 526 | 6.47E-03 | 3.23E-02 | -0.12 | 322 | 7.34E-01 | 8.26E-01 | -1.03 | 510 | 2.59E-04 | 9.73E-04 | -0.84 | 537 | 4.70E-03 | 4.56E-02 | -0.56 | 394 | 1.48E-01 | 2.43E-01 |
| ILMN_1661516 | POLR2J2/POLR2J3 | 7q22.1f | -0.63 | 8.02E-05 | 1.35E-03 | 5.59E-01 | 3.36E-08 | 0.91 | -0.19 | -1.10 | -0.31 | 403 | 7.89E-02 | 1.97E-01 | -0.30 | 385 | 1.92E-01 | 3.25E-01 | -1.10 | 522 | 1.94E-09 | 3.36E-08 | -0.74 | 527 | 7.59E-05 | 4.42E-03 | -0.19 | 358 | 4.39E-01 | 5.59E-01 |
| ILMN_1783142 | RPS4Y1 | Yp11.31a | -0.80 | 4.89E-02 | 1.31E-01 | 7.52E-01 | 4.60E-02 | 0.89 | -0.25 | -1.14 | -0.25 | 383 | 6.04E-01 | 7.52E-01 | -0.61 | 458 | 3.30E-01 | 4.82E-01 | -1.09 | 521 | 2.25E-02 | 4.60E-02 | -1.14 | 558 | 2.33E-02 | 1.21E-01 | -0.98 | 428 | 1.40E-01 | 2.34E-01 |
| ILMN_1775016 | MPZL2 | 11q23.3d | 0.45 | 9.27E-04 | 7.58E-03 | 7.03E-01 | 4.89E-05 | 0.89 | 1.01 | 0.12 | 0.60 | 128 | 1.47E-04 | 1.77E-03 | 0.12 | 240 | 5.74E-01 | 7.03E-01 | 0.40 | 204 | 1.13E-02 | 2.55E-02 | 0.25 | 158 | 1.29E-01 | 3.31E-01 | 1.01 | 226 | 6.23E-06 | 4.89E-05 |
| ILMN_1775357 | SNX29 | 16p13.13a | 0.63 | 8.44E-05 | 1.40E-03 | 6.95E-01 | 4.70E-07 | 0.89 | 1.02 | 0.14 | 0.46 | 191 | 1.11E-02 | 4.83E-02 | 0.14 | 230 | 5.64E-01 | 6.95E-01 | 1.02 | 85 | 4.11E-08 | 4.70E-07 | 0.53 | 83 | 5.65E-03 | 5.11E-02 | 0.65 | 274 | 1.07E-02 | 2.86E-02 |
| ILMN_1781373 | IFIH1 | 2q24.2d | 0.93 | 1.40E-10 | 1.27E-07 | 1.55E-01 | 6.28E-11 | 0.88 | 1.25 | 0.37 | 1.05 | 56 | 8.20E-11 | 2.94E-08 | 0.37 | 150 | 7.12E-02 | 1.55E-01 | 1.16 | 44 | 1.31E-12 | 6.28E-11 | 0.53 | 82 | 1.23E-03 | 2.11E-02 | 1.25 | 91 | 2.02E-08 | 3.69E-07 |
| ILMN_1709925 | GPR155 | 2q31.1g | 0.38 | 3.61E-03 | 2.01E-02 | 3.65E-01 | 7.21E-06 | 0.87 | 1.03 | 0.16 | 0.50 | 171 | 6.76E-04 | 5.77E-03 | 0.55 | 125 | 3.92E-03 | 1.67E-02 | 0.16 | 261 | 2.74E-01 | 3.65E-01 | 0.29 | 141 | 6.01E-02 | 2.11E-01 | 1.03 | 204 | 6.71E-07 | 7.21E-06 |
| ILMN_1716062 | MYL4 | 17q21.32a | -0.58 | 6.92E-03 | 3.20E-02 | 6.66E-01 | 4.53E-03 | 0.87 | -0.17 | -1.04 | -0.17 | 363 | 4.94E-01 | 6.66E-01 | -0.26 | 369 | 4.30E-01 | 5.80E-01 | -0.77 | 467 | 1.82E-03 | 5.31E-03 | -1.04 | 553 | 7.88E-05 | 4.53E-03 | -0.66 | 402 | 5.69E-02 | 1.13E-01 |
| ILMN_2249018 | LRRC26 | 9q34.3e | -1.23 | 2.98E-09 | 8.80E-07 | 2.47E-02 | 1.40E-06 | 0.87 | -0.87 | -1.74 | -1.31 | 558 | 1.25E-07 | 7.81E-06 | -0.87 | 493 | 6.46E-03 | 2.47E-02 | -1.31 | 549 | 1.43E-07 | 1.40E-06 | -0.92 | 547 | 3.45E-04 | 1.03E-02 | -1.74 | 557 | 4.74E-07 | 5.41E-06 |
| ILMN_1860003 | ##_HS.347034 | 0 | 0.52 | 2.56E-03 | 1.57E-02 | 3.31E-01 | 1.53E-04 | 0.86 | 1.18 | 0.32 | 0.65 | 120 | 1.23E-03 | 9.18E-03 | 1.18 | 38 | 1.00E-05 | 1.53E-04 | 0.35 | 215 | 8.27E-02 | 1.36E-01 | 0.32 | 129 | 1.28E-01 | 3.31E-01 | 0.70 | 267 | 1.18E-02 | 3.10E-02 |
| ILMN_1815190 | METTL1 | 12q14.1a | -0.43 | 2.07E-04 | 2.63E-03 | 5.52E-01 | 2.02E-06 | 0.86 | -0.15 | -1.00 | -0.55 | 479 | 4.05E-05 | 6.43E-04 | -0.15 | 328 | 4.01E-01 | 5.52E-01 | -0.39 | 390 | 3.20E-03 | 8.64E-03 | -0.22 | 417 | 1.21E-01 | 3.21E-01 | -1.00 | 431 | 1.51E-07 | 2.02E-06 |
| ILMN_2116827 | RGPD4** | 0 | 0.59 | 7.79E-03 | 3.48E-02 | 5.26E-01 | 3.88E-03 | 0.85 | 1.10 | 0.24 | 0.24 | 284 | 3.39E-01 | 5.26E-01 | 1.08 | 60 | 1.52E-03 | 7.98E-03 | 0.51 | 175 | 4.56E-02 | 8.28E-02 | 1.10 | 19 | 6.19E-05 | 3.88E-03 | 0.69 | 269 | 5.23E-02 | 1.06E-01 |
| ILMN_1764030 | CCL23 | 17q12b | 0.99 | 9.92E-04 | 7.96E-03 | 6.29E-01 | 1.49E-02 | 0.85 | 1.18 | 0.33 | 1.05 | 55 | 3.43E-03 | 2.03E-02 | 1.14 | 49 | 1.63E-02 | 5.01E-02 | 0.99 | 91 | 6.02E-03 | 1.49E-02 | 1.18 | 13 | 2.01E-03 | 2.76E-02 | 0.33 | 309 | 5.15E-01 | 6.29E-01 |
| ILMN_1780444 | ARL3 | 10q24.32b | -0.51 | 1.01E-05 | 3.01E-04 | 1.04E-01 | 9.80E-09 | 0.85 | -0.36 | -1.21 | -0.53 | 471 | 6.09E-05 | 8.91E-04 | -1.21 | 539 | 3.06E-11 | 9.80E-09 | -0.44 | 400 | 9.04E-04 | 2.89E-03 | -0.50 | 482 | 3.96E-04 | 1.12E-02 | -0.36 | 380 | 5.11E-02 | 1.04E-01 |
| ILMN_1697460 | REEP6 | 19p13.3h | -0.88 | 1.82E-13 | 1.37E-09 | 1.65E-02 | 2.02E-11 | 0.84 | -0.45 | -1.29 | -1.06 | 548 | 2.68E-15 | 2.02E-11 | -0.79 | 486 | 2.61E-06 | 5.41E-05 | -0.96 | 493 | 3.88E-13 | 2.31E-11 | -0.45 | 476 | 7.78E-04 | 1.65E-02 | -1.29 | 531 | 2.49E-12 | 2.23E-10 |
| ILMN_1680757 | LRRC26 | 9q34.3e | -1.08 | 8.72E-08 | 9.06E-06 | 9.76E-02 | 7.80E-06 | 0.84 | -0.64 | -1.48 | -1.16 | 553 | 1.24E-06 | 4.13E-05 | -0.64 | 463 | 3.87E-02 | 9.76E-02 | -1.17 | 533 | 1.01E-06 | 7.80E-06 | -0.78 | 532 | 1.77E-03 | 2.58E-02 | -1.48 | 545 | 9.08E-06 | 6.74E-05 |
| ILMN_2399016 | MMP28 | 17q12b | -1.30 | 1.86E-09 | 6.20E-07 | 1.64E-02 | 1.11E-08 | 0.84 | -0.87 | -1.70 | -1.70 | 563 | 1.91E-11 | 1.11E-08 | -1.51 | 560 | 3.41E-06 | 6.65E-05 | -1.03 | 508 | 2.95E-05 | 1.46E-04 | -0.87 | 541 | 7.69E-04 | 1.64E-02 | -1.53 | 548 | 9.62E-06 | 7.09E-05 |
| ILMN_1805750 | IFITM3 | 11p15.5d | 1.20 | 1.61E-07 | 1.44E-05 | 3.85E-02 | 1.09E-05 | 0.83 | 1.67 | 0.84 | 1.40 | 25 | 2.06E-07 | 1.09E-05 | 0.88 | 92 | 1.16E-02 | 3.85E-02 | 1.23 | 34 | 4.85E-06 | 3.03E-05 | 0.84 | 44 | 2.64E-03 | 3.23E-02 | 1.67 | 22 | 8.18E-06 | 6.17E-05 |
| ILMN_1759075 | TNFRSF13B | 17p11.2h | -0.55 | 8.08E-03 | 3.57E-02 | 4.40E-01 | 9.14E-04 | 0.83 | -0.23 | -1.05 | -0.97 | 541 | 6.30E-05 | 9.14E-04 | -0.34 | 395 | 2.85E-01 | 4.34E-01 | -0.23 | 352 | 3.45E-01 | 4.40E-01 | -0.33 | 448 | 1.85E-01 | 4.11E-01 | -1.05 | 467 | 1.73E-03 | 6.15E-03 |
| ILMN_1710962 | TMEM97 | 17q11.2a | -0.49 | 8.19E-07 | 4.64E-05 | 1.36E-01 | 7.30E-10 | 0.83 | -0.25 | -1.08 | -0.62 | 497 | 3.10E-08 | 2.77E-06 | -0.48 | 426 | 8.40E-04 | 4.95E-03 | -0.39 | 388 | 4.30E-04 | 1.51E-03 | -0.25 | 424 | 2.82E-02 | 1.36E-01 | -1.08 | 481 | 1.11E-11 | 7.30E-10 |
| ILMN_1701114 | GBP1 | 1p22.2c | 1.14 | 8.16E-13 | 4.10E-09 | 1.15E-02 | 4.96E-11 | 0.83 | 1.51 | 0.68 | 1.30 | 29 | 5.99E-13 | 8.73E-10 | 0.68 | 110 | 2.40E-03 | 1.15E-02 | 1.29 | 27 | 9.88E-13 | 4.96E-11 | 0.77 | 51 | 2.29E-05 | 2.42E-03 | 1.51 | 40 | 9.87E-10 | 2.92E-08 |
| ILMN_1714402 | ZCCHC2 | 18q21.33a | 1.16 | 1.53E-06 | 7.54E-05 | 9.11E-02 | 2.17E-05 | 0.83 | 1.55 | 0.72 | 1.44 | 23 | 5.18E-07 | 2.17E-05 | 1.04 | 72 | 4.98E-03 | 2.02E-02 | 1.12 | 51 | 8.30E-05 | 3.60E-04 | 0.72 | 59 | 1.47E-02 | 9.11E-02 | 1.55 | 37 | 9.31E-05 | 5.02E-04 |
| ILMN_1708790 | CA6 | 1p36.23a | -0.51 | 3.27E-05 | 7.12E-04 | 4.62E-01 | 5.58E-06 | 0.82 | -0.19 | -1.01 | -0.52 | 467 | 2.50E-04 | 2.64E-03 | -0.19 | 342 | 3.11E-01 | 4.62E-01 | -0.59 | 431 | 3.63E-05 | 1.75E-04 | -0.32 | 440 | 2.91E-02 | 1.39E-01 | -1.01 | 438 | 4.93E-07 | 5.58E-06 |
| ILMN_1770387 | TMPRSS9 | 19p13.3g | -0.62 | 1.35E-02 | 5.17E-02 | 6.47E-01 | 8.45E-03 | 0.82 | -0.21 | -1.03 | -0.21 | 372 | 4.73E-01 | 6.47E-01 | -0.44 | 418 | 2.45E-01 | 3.90E-01 | -0.86 | 481 | 3.12E-03 | 8.45E-03 | -1.03 | 552 | 8.29E-04 | 1.70E-02 | -0.91 | 419 | 2.41E-02 | 5.61E-02 |
| ILMN_1766054 | ABCA1 | 9q31.1d | 0.39 | 9.62E-04 | 7.78E-03 | 3.72E-01 | 1.31E-06 | 0.81 | 1.02 | 0.21 | 0.49 | 172 | 2.68E-04 | 2.80E-03 | 0.21 | 197 | 2.29E-01 | 3.72E-01 | 0.28 | 238 | 3.71E-02 | 6.98E-02 | 0.25 | 157 | 7.02E-02 | 2.33E-01 | 1.02 | 212 | 9.02E-08 | 1.31E-06 |
| ILMN_1659985 | RSPH9 | 6p21.1c | 0.61 | 9.03E-04 | 7.44E-03 | 1.79E-01 | 2.46E-04 | 0.81 | 1.24 | 0.43 | 0.66 | 115 | 2.08E-03 | 1.39E-02 | 0.88 | 93 | 1.88E-03 | 9.43E-03 | 0.43 | 196 | 4.43E-02 | 8.09E-02 | 0.45 | 101 | 4.48E-02 | 1.79E-01 | 1.24 | 101 | 4.11E-05 | 2.46E-04 |
| ILMN_1700671 | ETV7 | 6p21.31a | 1.06 | 4.02E-06 | 1.54E-04 | 1.30E-01 | 8.55E-06 | 0.81 | 1.43 | 0.62 | 1.43 | 24 | 1.42E-07 | 8.55E-06 | 1.06 | 68 | 2.50E-03 | 1.18E-02 | 0.88 | 106 | 9.62E-04 | 3.04E-03 | 0.62 | 74 | 2.60E-02 | 1.30E-01 | 1.38 | 57 | 2.10E-04 | 1.01E-03 |
| ILMN_1783621 | CMPK2 | 2p25.2a | 1.96 | 1.07E-06 | 5.67E-05 | 2.80E-02 | 3.73E-05 | 0.80 | 2.32 | 1.53 | 2.32 | 4 | 1.10E-06 | 3.73E-05 | 1.65 | 8 | 7.61E-03 | 2.80E-02 | 1.97 | 7 | 3.32E-05 | 1.61E-04 | 1.53 | 5 | 2.05E-03 | 2.80E-02 | 1.96 | 8 | 2.85E-03 | 9.34E-03 |
| ILMN_1708110 | TMEM144 | 4q32.1d-q32.1e | 0.67 | 5.25E-06 | 1.84E-04 | 1.45E-01 | 6.02E-06 | 0.79 | 1.20 | 0.41 | 0.79 | 91 | 3.31E-06 | 8.82E-05 | 0.41 | 145 | 6.53E-02 | 1.45E-01 | 0.66 | 148 | 9.09E-05 | 3.90E-04 | 0.43 | 105 | 1.55E-02 | 9.39E-02 | 1.20 | 113 | 5.41E-07 | 6.02E-06 |
| ILMN_1695604 | DCBLD1 | 6q22.2b | 0.81 | 1.39E-07 | 1.28E-05 | 3.84E-01 | 1.05E-07 | 0.79 | 1.06 | 0.27 | 0.92 | 73 | 3.74E-07 | 1.73E-05 | 0.27 | 179 | 2.40E-01 | 3.84E-01 | 1.06 | 68 | 7.36E-09 | 1.05E-07 | 0.53 | 85 | 4.94E-03 | 4.70E-02 | 0.64 | 275 | 9.91E-03 | 2.68E-02 |
| ILMN_2402640 | DSC1 | 18q12.1d | -0.65 | 6.65E-03 | 3.11E-02 | 4.46E-01 | 1.13E-03 | 0.78 | -0.33 | -1.11 | -1.11 | 552 | 8.15E-05 | 1.13E-03 | -0.78 | 484 | 3.29E-02 | 8.58E-02 | -0.33 | 377 | 2.36E-01 | 3.23E-01 | -0.36 | 462 | 2.13E-01 | 4.46E-01 | -0.51 | 389 | 1.87E-01 | 2.92E-01 |
| ILMN_1693836 | ##_LOC653344 | 0 | -0.73 | 4.69E-06 | 1.73E-04 | 2.56E-01 | 6.43E-06 | 0.78 | -0.33 | -1.11 | -0.98 | 542 | 9.73E-08 | 6.43E-06 | -1.11 | 524 | 4.21E-06 | 7.75E-05 | -0.69 | 456 | 1.33E-04 | 5.43E-04 | -0.33 | 444 | 8.32E-02 | 2.56E-01 | -0.98 | 429 | 1.12E-04 | 5.86E-04 |
| ILMN_1686664 | MT2A | 16q13b | 0.66 | 4.38E-06 | 1.65E-04 | 3.64E-01 | 5.86E-05 | 0.78 | 1.04 | 0.27 | 0.81 | 89 | 1.97E-06 | 5.86E-05 | 1.04 | 71 | 2.93E-06 | 5.91E-05 | 0.52 | 174 | 1.89E-03 | 5.49E-03 | 0.60 | 75 | 7.03E-04 | 1.57E-02 | 0.27 | 315 | 2.48E-01 | 3.64E-01 |

### Online Supplemental Table S4 (Continued)

| | | | S1 | | | M1 through M5 | | | | | M1 | | | | M2 | | | | M3 | | | | M4 | | | | M5 | | | |
|---|---|---|---|---|---|---|---|---|---|---|---|---|---|---|---|---|---|---|---|---|---|---|---|---|---|---|---|---|---|---|
| ID | Gene_Symbol | Cytoband | FC | $P_{val}$ | $P_{adj}$ | Max $P_{adj}$ | Min $P_{adj}$ | ΔFC | Max FC | Min FC | FC | FC Rank | $P_{val}$ | $P_{adj}$ | FC | FC Rank | $P_{val}$ | $P_{adj}$ | FC | FC Rank | $P_{val}$ | $P_{adj}$ | FC | FC Rank | $P_{val}$ | $P_{adj}$ | FC | FC Rank | $P_{val}$ | $P_{adj}$ |
| ILMN_1877196 | LRCH3 | 0 | 0.53 | 7.27E-05 | 1.27E-03 | 3.96E-01 | 1.82E-05 | 0.78 | 1.00 | 0.23 | 0.38 | 227 | 1.12E-02 | 4.85E-02 | 1.00 | 81 | 6.48E-07 | 1.82E-05 | 0.47 | 184 | 1.83E-03 | 5.32E-03 | 0.85 | 42 | 1.70E-07 | 2.55E-04 | 0.23 | 321 | 2.76E-01 | 3.96E-01 |
| ILMN_1711514 | COCH | 14q12e | -0.64 | 3.73E-03 | 2.05E-02 | 2.02E-01 | 4.21E-03 | 0.78 | -0.39 | -1.17 | -0.82 | 533 | 1.68E-03 | 1.18E-02 | -1.17 | 536 | 6.86E-04 | 4.21E-03 | -0.39 | 389 | 1.29E-01 | 1.96E-01 | -0.52 | 489 | 5.55E-02 | 2.02E-01 | -0.84 | 415 | 2.02E-02 | 4.85E-02 |
| ILMN_1720048 | CCL2 | 17q12a | 1.19 | 3.94E-05 | 8.08E-04 | 3.12E-01 | 2.86E-04 | 0.77 | 1.38 | 0.60 | 1.31 | 28 | 1.34E-04 | 1.65E-03 | 1.27 | 30 | 4.60E-03 | 1.90E-02 | 1.38 | 24 | 6.39E-05 | 2.86E-04 | 0.95 | 33 | 7.97E-03 | 6.27E-02 | 0.60 | 279 | 2.03E-01 | 3.12E-01 |
| ILMN_1670305 | SERPING1 | 11q12.1a | 1.48 | 9.21E-08 | 9.27E-06 | 2.95E-02 | 3.08E-06 | 0.77 | 1.80 | 1.03 | 1.80 | 8 | 3.56E-08 | 3.08E-06 | 1.41 | 17 | 8.20E-04 | 4.87E-03 | 1.44 | 19 | 8.95E-06 | 5.17E-05 | 1.03 | 26 | 2.25E-03 | 2.95E-02 | 1.73 | 19 | 1.17E-04 | 6.12E-04 |
| ILMN_1674009 | TKTL1 | Xq28g | -0.61 | 1.26E-04 | 1.86E-03 | 3.23E-01 | 2.47E-04 | 0.77 | -0.30 | -1.07 | -0.62 | 501 | 7.48E-04 | 6.25E-03 | -0.40 | 409 | 9.71E-02 | 1.96E-01 | -0.75 | 463 | 6.30E-05 | 2.83E-04 | -0.30 | 431 | 1.23E-01 | 3.23E-01 | -1.07 | 475 | 4.14E-05 | 2.47E-04 |
| ILMN_2377829 | NANOS1 | 10q26.11c | 0.46 | 1.83E-02 | 6.48E-02 | 2.98E-01 | 2.18E-03 | 0.76 | 1.13 | 0.37 | 0.37 | 231 | 1.14E-01 | 2.55E-01 | 1.13 | 51 | 2.93E-04 | 2.18E-03 | 0.54 | 171 | 2.25E-02 | 4.61E-02 | 0.40 | 115 | 1.07E-01 | 2.98E-01 | 0.46 | 294 | 1.60E-01 | 2.59E-01 |
| ILMN_1793915 | MXI1 | 10q25.2a | -0.65 | 1.81E-03 | 1.22E-02 | 4.72E-01 | 5.61E-03 | 0.76 | -0.26 | -1.02 | -0.26 | 388 | 2.89E-01 | 4.72E-01 | -0.47 | 421 | 1.43E-01 | 2.64E-01 | -0.76 | 465 | 1.94E-03 | 5.61E-03 | -0.98 | 550 | 1.62E-04 | 6.64E-03 | -1.02 | 443 | 2.98E-03 | 9.70E-03 |
| ILMN_1674640 | CXCR6 | 3p21.31i | -0.81 | 2.98E-05 | 6.64E-04 | 3.39E-01 | 2.68E-04 | 0.76 | -0.36 | -1.11 | -1.00 | 546 | 1.44E-05 | 2.81E-04 | -0.77 | 482 | 9.96E-03 | 3.44E-02 | -0.92 | 487 | 5.93E-05 | 2.68E-04 | -0.36 | 459 | 1.34E-01 | 3.39E-01 | -1.11 | 496 | 4.86E-04 | 2.09E-03 |
| ILMN_1690241 | BATF2 | 11q13.1c | 1.31 | 1.82E-07 | 1.56E-05 | 3.36E-02 | 1.90E-06 | 0.75 | 1.65 | 0.90 | 1.65 | 14 | 1.91E-08 | 1.90E-06 | 1.56 | 11 | 4.08E-05 | 4.64E-04 | 1.17 | 42 | 5.61E-05 | 2.55E-04 | 0.90 | 38 | 2.85E-03 | 3.36E-02 | 1.57 | 35 | 1.01E-04 | 5.40E-04 |
| ILMN_2370336 | MS4A4A | 11q12.2a | 1.02 | 1.23E-08 | 2.50E-06 | 1.15E-01 | 2.05E-08 | 0.75 | 1.27 | 0.52 | 1.08 | 52 | 1.40E-07 | 8.54E-06 | 0.52 | 129 | 4.80E-02 | 1.15E-01 | 1.27 | 29 | 1.09E-09 | 2.05E-08 | 0.72 | 60 | 7.79E-04 | 1.65E-02 | 1.14 | 133 | 5.90E-05 | 3.35E-04 |
| ILMN_1671260 | WLS | 1p31.3a | -0.72 | 7.11E-06 | 2.34E-04 | 2.58E-01 | 9.15E-08 | 0.74 | -0.35 | -1.10 | -0.46 | 456 | 1.13E-02 | 4.87E-02 | -0.35 | 397 | 1.40E-01 | 2.58E-01 | -1.10 | 523 | 6.23E-09 | 9.15E-08 | -0.84 | 538 | 1.75E-05 | 2.01E-03 | -0.38 | 383 | 1.30E-01 | 2.20E-01 |
| ILMN_1707979 | CARD17P | 11q22.3b | 0.85 | 1.69E-05 | 4.36E-04 | 8.64E-02 | 6.35E-05 | 0.74 | 1.33 | 0.59 | 1.09 | 50 | 2.20E-06 | 6.35E-05 | 0.90 | 90 | 2.67E-03 | 1.24E-02 | 0.71 | 139 | 1.74E-03 | 5.10E-03 | 0.59 | 76 | 1.34E-02 | 8.64E-02 | 1.33 | 67 | 3.26E-05 | 2.03E-04 |
| ILMN_1710303 | ODAD4 | 17q21.2b | -0.80 | 7.26E-04 | 6.39E-03 | 3.24E-01 | 1.14E-03 | 0.74 | -0.43 | -1.17 | -0.43 | 440 | 1.17E-01 | 2.58E-01 | -0.47 | 423 | 1.90E-01 | 3.24E-01 | -1.00 | 500 | 3.11E-04 | 1.14E-03 | -1.17 | 559 | 7.45E-05 | 4.35E-03 | -0.97 | 427 | 1.17E-02 | 3.08E-02 |
| ILMN_1723944 | TARP | 7p14.1e | -0.46 | 3.68E-03 | 2.04E-02 | 2.70E-01 | 3.98E-04 | 0.73 | -0.30 | -1.04 | -0.57 | 484 | 2.23E-03 | 1.47E-02 | -0.69 | 471 | 4.80E-03 | 1.96E-02 | -0.30 | 370 | 1.02E-01 | 1.61E-01 | -0.33 | 446 | 9.10E-02 | 2.70E-01 | -1.04 | 461 | 7.20E-05 | 3.98E-04 |
| ILMN_1678833 | CCR1 | 3p21.31i | 0.60 | 3.14E-06 | 1.28E-04 | 1.90E-01 | 7.23E-06 | 0.73 | 1.03 | 0.30 | 0.71 | 107 | 1.65E-06 | 5.16E-05 | 0.36 | 152 | 5.92E-02 | 1.35E-01 | 0.64 | 155 | 1.38E-05 | 7.52E-05 | 0.30 | 137 | 5.01E-02 | 1.90E-01 | 1.03 | 207 | 6.74E-07 | 7.23E-06 |
| ILMN_2148785 | GBP1 | 1p22.2c | 0.95 | 2.23E-10 | 1.52E-07 | 1.10E-01 | 4.36E-09 | 0.71 | 1.15 | 0.43 | 1.15 | 44 | 2.60E-11 | 1.38E-08 | 0.43 | 140 | 4.50E-02 | 1.10E-01 | 1.10 | 56 | 1.81E-10 | 4.36E-09 | 0.66 | 70 | 1.62E-04 | 6.64E-03 | 0.87 | 249 | 1.98E-04 | 9.61E-04 |
| ILMN_1695485 | FAM225A | 0 | 0.94 | 2.50E-06 | 1.07E-04 | 1.97E-01 | 2.17E-05 | 0.71 | 1.17 | 0.47 | 1.17 | 40 | 5.16E-07 | 2.17E-05 | 0.78 | 99 | 9.23E-03 | 3.24E-02 | 1.04 | 77 | 8.00E-06 | 4.69E-05 | 0.47 | 93 | 5.29E-02 | 1.97E-01 | 0.99 | 236 | 2.04E-03 | 7.05E-03 |
| ILMN_1740426 | RASD1 | 17p11.2g | -0.62 | 1.25E-05 | 3.54E-04 | 2.15E-01 | 7.23E-05 | 0.70 | -0.32 | -1.02 | -0.60 | 490 | 2.48E-04 | 2.63E-03 | -0.72 | 476 | 7.97E-04 | 4.76E-03 | -0.71 | 458 | 1.86E-05 | 9.70E-05 | -0.32 | 438 | 6.19E-02 | 2.15E-01 | -1.02 | 450 | 9.84E-06 | 7.23E-05 |
| ILMN_2239772 | ABHD17AP1 | 1q21.1c | -0.46 | 8.98E-07 | 4.93E-05 | 3.23E-02 | 8.31E-10 | 0.69 | -0.33 | -1.02 | -0.46 | 455 | 9.70E-06 | 2.08E-04 | -0.53 | 436 | 1.22E-04 | 1.09E-03 | -0.44 | 397 | 3.03E-05 | 1.48E-04 | -0.33 | 445 | 2.65E-03 | 3.23E-02 | -1.02 | 447 | 1.35E-11 | 8.31E-10 |
| ILMN_1706502 | EIF2AK2 | 2p22.2b | 0.96 | 1.14E-09 | 4.09E-07 | 3.80E-02 | 2.98E-07 | 0.69 | 1.28 | 0.60 | 1.04 | 57 | 1.86E-08 | 1.86E-06 | 0.60 | 122 | 1.14E-02 | 3.80E-02 | 1.03 | 80 | 2.41E-08 | 2.98E-07 | 0.73 | 56 | 1.26E-04 | 5.91E-03 | 1.28 | 81 | 5.31E-07 | 5.93E-06 |
| ILMN_1740200 | ##_USP18 | 22q11.21b | 0.99 | 5.01E-05 | 9.61E-04 | 2.36E-01 | 1.32E-03 | 0.69 | 1.27 | 0.59 | 1.07 | 54 | 2.25E-04 | 2.44E-03 | 1.27 | 29 | 8.23E-04 | 4.88E-03 | 1.03 | 79 | 3.67E-04 | 1.32E-03 | 0.89 | 40 | 3.46E-03 | 3.77E-02 | 0.59 | 281 | 1.42E-01 | 2.36E-01 |
| ILMN_1735364 | ANKFY1 | 17p13.2c | 0.59 | 1.68E-05 | 4.34E-04 | 1.00E-02 | 1.53E-06 | 0.68 | 1.19 | 0.50 | 0.50 | 164 | 1.37E-03 | 1.00E-02 | 1.19 | 35 | 2.31E-08 | 1.53E-06 | 0.51 | 176 | 1.25E-03 | 3.83E-03 | 0.64 | 71 | 1.38E-04 | 6.18E-03 | 0.78 | 258 | 4.47E-04 | 1.95E-03 |
| ILMN_2215656 | MAGOH | 1p32.3c | -0.53 | 4.30E-04 | 4.36E-03 | 2.21E-01 | 2.99E-04 | 0.66 | -0.34 | -1.01 | -0.69 | 509 | 1.07E-04 | 1.38E-03 | -0.53 | 437 | 2.25E-02 | 6.41E-02 | -0.37 | 383 | 3.56E-02 | 6.75E-02 | -0.34 | 452 | 6.48E-02 | 2.21E-01 | -1.01 | 436 | 5.16E-05 | 2.99E-04 |
| ILMN_1783909 | COL6A2 | 21q22.3f | -0.67 | 2.40E-03 | 1.50E-02 | 3.58E-01 | 7.45E-03 | 0.66 | -0.40 | -1.06 | -0.86 | 536 | 9.30E-04 | 7.45E-03 | -1.06 | 515 | 2.02E-03 | 1.00E-02 | -0.67 | 452 | 9.65E-03 | 2.23E-02 | -0.40 | 468 | 1.47E-01 | 3.58E-01 | -0.75 | 409 | 3.94E-02 | 8.44E-02 |
| ILMN_2049766 | NFE2L3 | 7p15.2b | 0.83 | 1.31E-11 | 1.66E-08 | 7.65E-02 | 1.11E-11 | 0.66 | 1.05 | 0.39 | 0.89 | 78 | 1.79E-10 | 5.39E-08 | 0.39 | 147 | 2.82E-02 | 7.65E-02 | 1.05 | 74 | 1.66E-13 | 1.11E-11 | 0.69 | 64 | 1.43E-06 | 5.75E-04 | 0.52 | 291 | 5.18E-03 | 1.55E-02 |
| ILMN_1664543 | IFIT3 | 10q23.31b | 1.40 | 1.36E-05 | 3.72E-04 | 5.01E-02 | 1.91E-04 | 0.66 | 1.75 | 1.09 | 1.68 | 13 | 8.68E-06 | 1.91E-04 | 1.17 | 42 | 1.63E-02 | 5.01E-02 | 1.37 | 26 | 2.63E-04 | 9.86E-04 | 1.09 | 23 | 5.34E-03 | 4.96E-02 | 1.75 | 16 | 7.99E-04 | 3.19E-03 |
| ILMN_1669321 | MATK | 19p13.3e | -0.90 | 1.79E-06 | 8.39E-05 | 8.80E-02 | 6.68E-05 | 0.65 | -0.61 | -1.26 | -1.00 | 545 | 6.79E-06 | 1.58E-04 | -0.61 | 457 | 3.39E-02 | 8.80E-02 | -0.97 | 494 | 1.20E-05 | 6.68E-05 | -0.65 | 509 | 4.92E-03 | 4.70E-02 | -1.26 | 528 | 4.46E-05 | 2.63E-04 |
| ILMN_1709747 | EXOG | 3p22.2a | 0.79 | 4.24E-07 | 2.96E-05 | 1.92E-01 | 5.27E-06 | 0.65 | 1.05 | 0.40 | 0.99 | 62 | 7.32E-08 | 5.27E-06 | 1.05 | 70 | 1.25E-05 | 1.83E-04 | 0.70 | 143 | 1.12E-04 | 4.69E-04 | 0.62 | 73 | 1.02E-03 | 1.89E-02 | 0.40 | 303 | 1.09E-01 | 1.92E-01 |
| ILMN_1765332 | TIMM10 | 11q12.1a | 0.76 | 1.35E-04 | 1.94E-03 | 1.11E-01 | 1.38E-03 | 0.64 | 1.07 | 0.43 | 0.91 | 74 | 1.08E-04 | 1.38E-03 | 0.74 | 105 | 1.52E-02 | 4.75E-02 | 0.43 | 193 | 6.45E-02 | 1.11E-01 | 0.80 | 47 | 1.15E-03 | 2.02E-02 | 1.07 | 174 | 1.06E-03 | 4.06E-03 |
| ILMN_2170814 | LAMP3 | 3q27.1a | 1.75 | 3.13E-08 | 5.02E-06 | 2.47E-02 | 9.75E-06 | 0.64 | 1.96 | 1.32 | 1.96 | 6 | 1.86E-07 | 1.04E-05 | 1.32 | 25 | 6.46E-03 | 2.47E-02 | 1.81 | 8 | 1.31E-06 | 9.75E-06 | 1.51 | 6 | 1.13E-04 | 5.65E-03 | 1.78 | 15 | 5.68E-04 | 2.37E-03 |
| ILMN_1663347 | OTOF | 2p23.3b | 2.70 | 1.75E-06 | 8.23E-05 | 2.35E-02 | 1.08E-04 | 0.64 | 2.88 | 2.25 | 2.79 | 2 | 3.70E-05 | 6.01E-04 | 2.47 | 2 | 5.03E-03 | 2.03E-02 | 2.88 | 2 | 2.11E-05 | 1.08E-04 | 2.25 | 1 | 1.49E-03 | 2.35E-02 | 2.67 | 1 | 4.33E-03 | 1.33E-02 |
| ILMN_1690105 | STAT1 | 2q32.2b | 0.72 | 1.50E-10 | 1.27E-07 | 5.15E-02 | 6.74E-09 | 0.63 | 1.01 | 0.39 | 0.82 | 84 | 2.01E-10 | 5.83E-08 | 0.39 | 146 | 1.69E-02 | 5.15E-02 | 0.81 | 115 | 2.97E-10 | 6.74E-09 | 0.46 | 95 | 4.68E-04 | 1.23E-02 | 1.01 | 219 | 1.34E-08 | 2.58E-07 |
| ILMN_2371724 | CEACAM1 | 19q13.2c | 0.61 | 6.92E-08 | 7.78E-06 | 4.20E-02 | 7.16E-08 | 0.62 | 1.00 | 0.38 | 0.82 | 83 | 2.73E-10 | 7.16E-08 | 0.74 | 104 | 9.13E-06 | 1.42E-04 | 0.42 | 199 | 9.50E-04 | 3.01E-03 | 0.38 | 117 | 4.17E-03 | 4.20E-02 | 1.00 | 233 | 2.82E-08 | 4.95E-07 |
| ILMN_1723912 | IFI44L | 1p31.1e | 2.20 | 4.33E-08 | 5.93E-06 | 1.51E-02 | 3.48E-06 | 0.62 | 2.42 | 1.80 | 2.29 | 5 | 1.40E-06 | 4.50E-05 | 1.80 | 5 | 3.43E-03 | 1.51E-02 | 2.42 | 4 | 4.05E-07 | 3.48E-06 | 1.90 | 3 | 1.26E-04 | 5.91E-03 | 2.29 | 4 | 4.61E-04 | 2.00E-03 |
| ILMN_1695404 | LY6E | 8q24.3f | 1.46 | 5.27E-08 | 6.62E-06 | 1.78E-02 | 1.43E-06 | 0.58 | 1.68 | 1.10 | 1.51 | 19 | 1.90E-06 | 5.71E-05 | 1.18 | 37 | 4.02E-03 | 1.70E-02 | 1.68 | 11 | 1.46E-07 | 1.43E-06 | 1.10 | 20 | 8.95E-04 | 1.78E-02 | 1.64 | 25 | 1.86E-04 | 9.12E-04 |
| ILMN_2388547 | EPSTI1 | 13q14.11c-q14.11d | 1.57 | 2.07E-10 | 1.52E-07 | 3.86E-03 | 1.80E-08 | 0.58 | 1.78 | 1.19 | 1.69 | 12 | 4.47E-09 | 5.81E-07 | 1.31 | 26 | 3.87E-04 | 2.70E-03 | 1.78 | 9 | 9.31E-10 | 1.80E-08 | 1.19 | 12 | 6.13E-05 | 3.86E-03 | 1.71 | 21 | 1.52E-05 | 1.05E-04 |
| ILMN_2410826 | OAS1 | 12q24.13b | 1.01 | 4.84E-05 | 9.38E-04 | 1.01E-01 | 1.68E-03 | 0.58 | 1.38 | 0.81 | 1.13 | 46 | 1.37E-04 | 1.68E-03 | 1.38 | 20 | 3.92E-04 | 2.73E-03 | 0.98 | 93 | 9.50E-04 | 3.01E-03 | 0.84 | 43 | 6.69E-03 | 5.63E-02 | 0.81 | 255 | 4.93E-02 | 1.01E-01 |
| ILMN_1745242 | PLSCR1 | 3q24d | 0.72 | 1.21E-04 | 1.80E-03 | 1.76E-01 | 1.64E-04 | 0.57 | 1.03 | 0.46 | 0.99 | 61 | 7.06E-06 | 1.64E-04 | 0.94 | 88 | 1.10E-03 | 6.11E-03 | 0.51 | 177 | 1.93E-02 | 4.06E-02 | 0.46 | 94 | 4.36E-02 | 1.76E-01 | 1.03 | 205 | 7.10E-04 | 2.88E-03 |
| ILMN_1683678 | SPATS2L | 2q33.1e | 1.50 | 1.19E-07 | 1.15E-05 | 3.20E-02 | 3.72E-06 | 0.57 | 1.70 | 1.13 | 1.62 | 16 | 1.36E-06 | 4.42E-05 | 1.13 | 52 | 9.07E-03 | 3.20E-02 | 1.70 | 10 | 4.37E-07 | 3.72E-06 | 1.15 | 15 | 9.74E-04 | 1.86E-02 | 1.43 | 51 | 1.92E-03 | 6.72E-03 |
| ILMN_1675640 | OAS1 | 12q24.13b | 1.08 | 3.82E-05 | 7.91E-04 | 9.27E-02 | 1.54E-03 | 0.56 | 1.43 | 0.87 | 1.16 | 43 | 1.96E-04 | 2.20E-03 | 1.43 | 16 | 5.09E-04 | 3.35E-03 | 1.10 | 54 | 4.39E-04 | 1.54E-03 | 0.94 | 35 | 4.10E-03 | 4.17E-02 | 0.87 | 246 | 4.43E-02 | 9.27E-02 |
| ILMN_1683792 | LAP3 | 4p15.32b | 0.84 | 2.68E-08 | 4.38E-06 | 4.65E-02 | 2.14E-08 | 0.55 | 1.05 | 0.49 | 0.93 | 71 | 3.90E-08 | 3.32E-06 | 0.63 | 120 | 3.85E-03 | 1.65E-02 | 1.04 | 75 | 1.14E-09 | 2.14E-08 | 0.49 | 89 | 4.84E-03 | 4.65E-02 | 1.05 | 194 | 7.96E-06 | 6.02E-05 |
| ILMN_1707695 | IFIT1 | 10q23.31b | 1.54 | 4.54E-07 | 3.08E-05 | 3.91E-02 | 4.84E-05 | 0.55 | 1.73 | 1.18 | 1.73 | 10 | 1.84E-06 | 5.57E-05 | 1.18 | 39 | 1.18E-02 | 3.91E-02 | 1.61 | 12 | 8.27E-06 | 4.84E-05 | 1.28 | 10 | 6.87E-04 | 1.54E-02 | 1.72 | 20 | 5.89E-04 | 2.45E-03 |
| ILMN_1674811 | OASL | 12q24.31a | 1.13 | 1.44E-06 | 7.24E-05 | 4.56E-02 | 3.88E-05 | 0.54 | 1.35 | 0.82 | 1.35 | 27 | 1.15E-06 | 3.88E-05 | 1.18 | 36 | 1.05E-03 | 5.93E-03 | 1.07 | 65 | 1.08E-04 | 4.54E-04 | 0.82 | 45 | 4.69E-03 | 4.56E-02 | 1.27 | 86 | 9.26E-04 | 3.62E-03 |
| ILMN_1779071 | FEZ1 | 11q24.2b | -1.04 | 1.82E-05 | 4.63E-04 | 8.72E-02 | 6.58E-04 | 0.54 | -0.75 | -1.28 | -1.19 | 554 | 4.20E-05 | 6.58E-04 | -1.15 | 533 | 2.27E-03 | 1.10E-02 | -1.05 | 513 | 2.71E-04 | 1.01E-03 | -0.75 | 529 | 1.36E-02 | 8.72E-02 | -1.28 | 530 | 1.44E-03 | 5.27E-03 |
| ILMN_1795181 | DDX60 | 4q32.3e | 1.15 | 4.70E-09 | 1.20E-06 | 2.53E-02 | 4.33E-07 | 0.52 | 1.32 | 0.80 | 1.23 | 35 | 8.34E-08 | 5.82E-06 | 0.80 | 95 | 6.66E-03 | 2.53E-02 | 1.27 | 28 | 3.73E-08 | 4.33E-07 | 0.92 | 36 | 1.11E-04 | 5.60E-03 | 1.32 | 69 | 3.03E-05 | 1.90E-04 |
| ILMN_1775175 | RPL3L | 16p13.3e | -0.66 | 3.50E-03 | 1.97E-02 | 1.72E-01 | 1.99E-02 | 0.51 | -0.49 | -1.00 | -0.49 | 459 | 6.52E-02 | 1.72E-01 | -0.78 | 485 | 2.51E-02 | 6.97E-02 | -0.66 | 449 | 1.37E-02 | 3.02E-02 | -0.74 | 526 | 8.37E-03 | 6.42E-02 | -1.00 | 432 | 7.00E-03 | 1.99E-02 |
| ILMN_1739428 | IFIT2 | 10q23.31b | 0.93 | 3.34E-08 | 5.19E-06 | 3.08E-02 | 1.67E-06 | 0.50 | 1.17 | 0.66 | 1.11 | 48 | 1.66E-08 | 1.67E-06 | 0.66 | 113 | 8.62E-03 | 3.08E-02 | 0.91 | 101 | 3.33E-06 | 2.19E-05 | 0.73 | 57 | 3.62E-04 | 1.06E-02 | 1.17 | 122 | 1.68E-05 | 1.14E-04 |
| ILMN_1699695 | TNFRSF21 | 6p12.3d | -0.68 | 1.52E-04 | 2.11E-03 | 7.42E-02 | 1.83E-03 | 0.49 | -0.59 | -1.08 | -0.70 | 512 | 1.40E-03 | 1.02E-02 | -0.72 | 475 | 1.23E-02 | 4.03E-02 | -0.65 | 446 | 2.77E-03 | 7.63E-03 | -0.59 | 498 | 1.06E-02 | 7.42E-02 | -1.08 | 482 | 4.18E-04 | 1.83E-03 |
| ILMN_2239754 | IFIT3 | 10q23.31b | 0.98 | 1.45E-04 | 2.05E-03 | 9.03E-02 | 5.99E-04 | 0.48 | 1.25 | 0.77 | 1.25 | 32 | 3.67E-05 | 5.99E-04 | 1.08 | 58 | 6.08E-03 | 2.36E-02 | 0.80 | 120 | 7.80E-03 | 1.86E-02 | 0.77 | 52 | 1.45E-02 | 9.03E-02 | 1.25 | 94 | 2.93E-03 | 9.56E-03 |
| ILMN_1725320 | SIGLEC1 | 20p13b | 1.09 | 2.83E-05 | 6.41E-04 | 8.14E-02 | 9.07E-04 | 0.48 | 1.28 | 0.81 | 1.18 | 39 | 1.24E-04 | 1.55E-03 | 0.96 | 87 | 1.62E-02 | 4.98E-02 | 1.13 | 49 | 2.38E-04 | 9.07E-04 | 0.81 | 46 | 1.23E-02 | 8.14E-02 | 1.28 | 82 | 2.69E-03 | 8.88E-03 |
| ILMN_1708303 | CYP4F22 | 19p13.12a | 0.97 | 6.16E-05 | 1.13E-03 | 1.60E-01 | 3.52E-04 | 0.47 | 1.15 | 0.67 | 0.96 | 67 | 8.29E-04 | 6.82E-03 | 0.67 | 112 | 7.40E-02 | 1.60E-01 | 1.15 | 46 | 8.07E-05 | 3.52E-04 | 0.87 | 41 | 4.38E-03 | 4.34E-02 | 0.84 | 252 | 3.59E-02 | 7.82E-02 |
| ILMN_1742618 | XAF1 | 17p13.2a | 1.19 | 5.09E-08 | 6.44E-06 | 1.83E-02 | 2.21E-05 | 0.46 | 1.43 | 0.97 | 1.22 | 36 | 2.30E-06 | 6.55E-05 | 1.43 | 15 | 2.41E-05 | 3.10E-04 | 1.20 | 37 | 3.36E-06 | 2.21E-05 | 1.12 | 18 | 3.60E-05 | 3.05E-03 | 0.97 | 237 | 6.32E-03 | 1.83E-02 |
| ILMN_2054019 | ISG15 | 1p36.33b | 1.54 | 4.49E-07 | 3.08E-05 | 2.53E-02 | 9.70E-05 | 0.45 | 1.74 | 1.29 | 1.63 | 15 | 5.80E-06 | 1.40E-04 | 1.74 | 7 | 2.23E-04 | 1.76E-03 | 1.54 | 14 | 1.86E-05 | 9.70E-05 | 1.46 | 8 | 1.17E-04 | 5.67E-03 | 1.29 | 79 | 9.26E-03 | 2.53E-02 |
| ILMN_1658579 | CDK5RAP3 | 17q21.32b | 0.74 | 1.45E-04 | 2.05E-03 | 4.33E-02 | 2.86E-03 | 0.42 | 1.00 | 0.58 | 0.58 | 138 | 9.62E-03 | 4.33E-02 | 1.00 | 83 | 7.47E-04 | 4.51E-03 | 0.71 | 141 | 1.74E-03 | 5.10E-03 | 1.00 | 28 | 3.04E-05 | 2.86E-03 | 0.84 | 251 | 7.05E-03 | 2.01E-02 |
| ILMN_2386790 | KLRC3 | 12p13.2c | -0.75 | 1.49E-03 | 1.08E-02 | 1.55E-01 | 2.33E-02 | 0.40 | -0.63 | -1.03 | -0.76 | 527 | 6.55E-03 | 3.26E-02 | -0.66 | 468 | 7.09E-02 | 1.55E-01 | -0.70 | 457 | 1.25E-02 | 2.77E-02 | -0.63 | 506 | 3.18E-02 | 1.46E-01 | -1.03 | 458 | 8.41E-03 | 2.33E-02 |
| ILMN_2374865 | ATF3 | 1q32.3c | 1.20 | 5.48E-08 | 6.69E-06 | 9.33E-03 | 8.83E-06 | 0.40 | 1.38 | 0.98 | 1.36 | 26 | 1.48E-07 | 8.83E-06 | 1.38 | 21 | 4.65E-05 | 5.16E-04 | 1.19 | 40 | 3.97E-06 | 2.55E-05 | 0.98 | 30 | 2.87E-04 | 9.33E-03 | 1.29 | 77 | 2.96E-04 | 1.37E-03 |
| ILMN_1701789 | IFIT3 | 10q23.31b | 1.29 | 6.35E-08 | 7.41E-06 | 8.01E-03 | 8.94E-06 | 0.40 | 1.48 | 1.09 | 1.48 | 21 | 1.51E-07 | 8.94E-06 | 1.26 | 31 | 5.79E-04 | 3.70E-03 | 1.26 | 30 | 7.01E-06 | 4.18E-05 | 1.09 | 24 | 2.16E-04 | 8.01E-03 | 1.34 | 62 | 5.33E-04 | 2.25E-03 |
| ILMN_1772964 | CCL8 | 17q12a | 1.11 | 3.17E-06 | 1.29E-04 | 6.78E-02 | 1.40E-04 | 0.39 | 1.24 | 0.85 | 1.24 | 33 | 1.41E-05 | 2.76E-04 | 0.97 | 86 | 8.46E-03 | 3.03E-02 | 1.19 | 39 | 2.82E-05 | 1.40E-04 | 0.91 | 37 | 2.17E-03 | 2.89E-02 | 0.85 | 250 | 3.01E-02 | 6.78E-02 |
| ILMN_1690365 | USP41P | 22q11.21d | 1.40 | 2.58E-06 | 1.09E-04 | 3.06E-02 | 1.77E-04 | 0.38 | 1.51 | 1.13 | 1.49 | 20 | 2.86E-05 | 4.92E-04 | 1.22 | 32 | 8.09E-03 | 2.93E-02 | 1.47 | 17 | 3.69E-05 | 1.77E-04 | 1.13 | 17 | 2.40E-03 | 3.06E-02 | 1.51 | 41 | 2.13E-03 | 7.31E-03 |
| ILMN_2173975 | RTP4 | 3q27.3b | 0.91 | 5.10E-06 | 1.80E-04 | 4.99E-02 | 2.76E-04 | 0.38 | 1.14 | 0.75 | 1.03 | 59 | 1.40E-05 | 2.76E-04 | 1.14 | 48 | 2.47E-04 | 1.91E-03 | 0.89 | 104 | 1.68E-04 | 6.69E-04 | 0.77 | 53 | 1.99E-03 | 2.75E-02 | 0.75 | 261 | 2.08E-02 | 4.99E-02 |
| ILMN_2347798 | IFI6 | 1p35.3b | 1.11 | 2.59E-05 | 6.01E-04 | 7.02E-02 | 8.54E-04 | 0.37 | 1.31 | 0.94 | 1.27 | 30 | 5.79E-05 | 8.54E-04 | 1.31 | 27 | 1.53E-03 | 8.00E-03 | 1.06 | 70 | 7.87E-04 | 2.56E-03 | 1.00 | 29 | 2.51E-03 | 3.12E-02 | 0.94 | 240 | 3.14E-02 | 7.02E-02 |
| ILMN_1733811 | JUP | 17q21.2b | 0.91 | 1.15E-05 | 3.32E-04 | 7.83E-02 | 1.88E-04 | 0.32 | 1.02 | 0.70 | 0.97 | 66 | 9.27E-05 | 1.24E-03 | 0.75 | 102 | 1.99E-02 | 5.86E-02 | 1.02 | 84 | 3.96E-05 | 1.88E-04 | 0.70 | 62 | 6.71E-03 | 5.64E-02 | 0.72 | 265 | 3.60E-02 | 7.83E-02 |

### Online Supplemental Table S4 (Continued)

| | | | S1 | | | M1 through M5 | | | | | M1 | | | | M2 | | | | M3 | | | | M4 | | | | M5 | | | |
|---|---|---|---|---|---|---|---|---|---|---|---|---|---|---|---|---|---|---|---|---|---|---|---|---|---|---|---|---|---|---|
| ID | Gene_Symbol | Cytoband | FC | $P_{val}$ | $P_{adj}$ | Max $P_{adj}$ | Min $P_{adj}$ | ΔFC | Max FC | Min FC | FC | FC Rank | $P_{val}$ | $P_{adj}$ | FC | FC Rank | $P_{val}$ | $P_{adj}$ | FC | FC Rank | $P_{val}$ | $P_{adj}$ | FC | FC Rank | $P_{val}$ | $P_{adj}$ | FC | FC Rank | $P_{val}$ | $P_{adj}$ |
| ILMN_1715332 | TTC21A | 3p22.2a | 1.07 | 5.67E-09 | 1.40E-06 | 2.89E-03 | 3.58E-07 | 0.28 | 1.23 | 0.95 | 0.98 | 65 | 6.46E-06 | 1.52E-04 | 1.10 | 55 | 1.02E-04 | 9.55E-04 | 1.22 | 35 | 3.01E-08 | 3.58E-07 | 0.95 | 34 | 3.23E-05 | 2.89E-03 | 1.23 | 104 | 4.54E-05 | 2.67E-04 |
| ILMN_2411236 | NRCAM | 7q31.1a | -0.95 | 1.57E-15 | 2.37E-11 | 6.76E-06 | 2.26E-10 | 0.27 | -0.79 | -1.06 | -1.06 | 549 | 6.62E-14 | 2.26E-10 | -0.98 | 499 | 5.23E-08 | 2.81E-06 | -0.79 | 470 | 1.08E-08 | 1.46E-07 | -0.92 | 548 | 2.62E-10 | 3.95E-06 | -0.95 | 424 | 6.24E-07 | 6.76E-06 |
| ILMN_1736729 | OAS2 | 12q24.13b | 1.07 | 6.59E-06 | 2.22E-04 | 3.52E-02 | 5.11E-04 | 0.27 | 1.16 | 0.90 | 1.16 | 41 | 4.39E-05 | 6.82E-04 | 1.15 | 47 | 2.00E-03 | 9.95E-03 | 1.09 | 58 | 1.23E-04 | 5.11E-04 | 0.90 | 39 | 2.62E-03 | 3.20E-02 | 0.97 | 238 | 1.38E-02 | 3.52E-02 |
| ILMN_1655595 | SERPINE2 | 2q36.1d | -0.94 | 7.22E-08 | 7.82E-06 | 1.01E-02 | 7.74E-05 | 0.24 | -0.84 | -1.08 | -0.98 | 543 | 2.79E-06 | 7.74E-05 | -0.84 | 491 | 2.03E-03 | 1.01E-02 | -0.89 | 484 | 2.12E-05 | 1.09E-04 | -0.90 | 545 | 4.76E-05 | 3.48E-03 | -1.08 | 485 | 1.92E-04 | 9.37E-04 |
| ILMN_1815057 | PDGFRB | 5q33.1c | 1.19 | 3.80E-08 | 5.45E-06 | 7.22E-03 | 6.72E-06 | 0.20 | 1.26 | 1.06 | 1.23 | 34 | 1.26E-06 | 4.18E-05 | 1.06 | 66 | 1.35E-03 | 7.22E-03 | 1.26 | 32 | 8.56E-07 | 6.72E-06 | 1.09 | 22 | 4.11E-05 | 3.24E-03 | 1.11 | 150 | 1.59E-03 | 5.75E-03 |
| ILMN_1662358 | MX1 | 21q22.3a | 1.12 | 3.82E-06 | 1.49E-04 | 2.61E-02 | 6.58E-04 | 0.18 | 1.22 | 1.04 | 1.19 | 38 | 4.19E-05 | 6.58E-04 | 1.22 | 33 | 1.35E-03 | 7.24E-03 | 1.05 | 72 | 2.84E-04 | 1.06E-03 | 1.09 | 21 | 3.31E-04 | 1.01E-02 | 1.04 | 200 | 9.64E-03 | 2.61E-02 |
| ILMN_1813430 | TRIM69 | 15q21.1a | 1.00 | 2.50E-12 | 7.53E-09 | 2.12E-04 | 4.74E-08 | 0.12 | 1.07 | 0.95 | 0.95 | 69 | 1.53E-08 | 1.58E-06 | 0.99 | 85 | 5.68E-06 | 9.77E-05 | 1.00 | 88 | 2.92E-09 | 4.74E-08 | 1.07 | 25 | 1.87E-09 | 1.41E-05 | 0.95 | 239 | 3.44E-05 | 2.12E-04 |
